# A genomic history of the North Pontic Region from the Neolithic to the Bronze Age

**DOI:** 10.1101/2024.04.17.589600

**Authors:** Alexey G. Nikitin, Iosif Lazaridis, Nick Patterson, Svitlana Ivanova, Mykhailo Videiko, Valentin Dergachev, Nadiia Kotova, Malcolm Lillie, Inna Potekhina, Marta Krenz-Niedbała, Sylwia Łukasik, Serhij Makhortykh, Virginie Renson, Henry Shephard, Gennadie Sirbu, Sofiia Svyryd, Taras Tkachuk, Piotr Włodarczak, Kim Callan, Elizabeth Curtis, Eadaoin Harney, Lora Iliev, Aisling Kearns, Ann Marie Lawson, Megan Michel, Matthew Mah, Adam Micco, Jonas Oppenheimer, Lijun Qiu, J. Noah Workman, Fatma Zalzala, Swapan Mallick, Nadin Rohland, David Reich

**Affiliations:** Department of Biology, Grand Valley State University, Allendale, MI, USA; Department of Human Evolutionary Biology, Harvard University, Cambridge, MA, USA; Department of Genetics, Harvard Medical School, Boston, MA, USA; Broad Institute of MIT and Harvard, Cambridge, MA, USA; Institute of Archaeology, National Academy of Sciences of Ukraine, Kyiv, Ukraine; Scientific Research Laboratory of Archaeology, Borys Grinchenko Kyiv University, Kyiv, Ukraine; Center of Archaeology, Institute of Cultural Heritage, Academy of Science of Moldova, Chișinău, Moldova; Faculty of Biology, Adam Mickiewicz University in Poznań, Poznań, Poland; University of Missouri Research Reactor, Columbia, MO, USA; Archaeological Institute of America, Boston, MA, USA; Thracology Scientific Research Laboratory of the State University of Moldova, Department of Academic Management, Academy of Science of Moldova, Chișinău, Moldova; Museum of History of Ancient Halych, “An Ancient Halych” National Reserve, Halych, Ukraine; Institute of Archaeology and Ethnology, Polish Academy of Sciences, Kracow, Poland; Howard Hughes Medical Institute, Harvard Medical School, Boston, MA, USA

## Abstract

The north Black Sea (Pontic) Region was the nexus of the farmers of Old Europe and the foragers and pastoralists of the Eurasian steppe^1,2^, and the source of waves of migrants that expanded deep into Europe^3–5^. We report genome-wide data from 78 prehistoric North Pontic individuals to understand the genetic makeup of the people involved in these migrations and discover the reasons for their success. First, we show that native North Pontic foragers had ancestry not only from Balkan and Eastern hunter-gatherers^6^ but also from European farmers and, occasionally, Caucasus hunter-gatherers. More dramatic inflows ensued during the Eneolithic, when migrants from the Caucasus-Lower Volga area^7^ moved westward, bypassing the local foragers to mix with Trypillian farmers advancing eastward. People of the Usatove archaeological group in the Northwest Pontic were formed ca. 4500 BCE with an equal measure of ancestry from the two expanding groups. A different Caucasus-Lower Volga group, moving westward in a distinct but temporally overlapping wave, avoided the farmers altogether, and blended with the foragers instead to form the people of the Serednii Stih archaeological complex^7^. A third wave of expansion occurred when Yamna descendants of the Serednii Stih forming ca. 4000 BCE expanded during the Early Bronze Age (3300 BCE). The temporal gap between Serednii Stih and the Yamna expansion is bridged by a genetically Yamna individual from Mykhailivka in Ukraine (3635-3383 BCE), a site of uninterrupted archaeological continuity across the Eneolithic-Bronze Age transition, and the likely epicenter of Yamna formation. Each of these three waves propagated distinctive ancestries while also incorporating outsiders during its advance, a flexible strategy forged in the North Pontic region that may explain its peoples’ outsized success in spreading their genes and culture across Eurasia^3–5,8–10^.

## Introduction

The steppe and forest-steppe regions north of the Black Sea, known as the North Pontic Region (NPR, Fig. 1, Supplementary Information, section 1), have been proposed as the homeland for communities that developed core Indo-European language terminology^11^, which began spreading across Eurasia facilitated by the turn-of-the-third-millennium expansion of the Yamna archaeological complex (YAC)^10^. During the Early Metal Ages (Eneolithic and the Early Bronze Age, EBA), a diverse array of archaeological groups inhabited the NPR. In principle, important information about how these populations interacted with each other can be learned from their genetic relationships—complementing the archaeological evidence—but key aspects remain poorly understood.

**Fig. 1:**
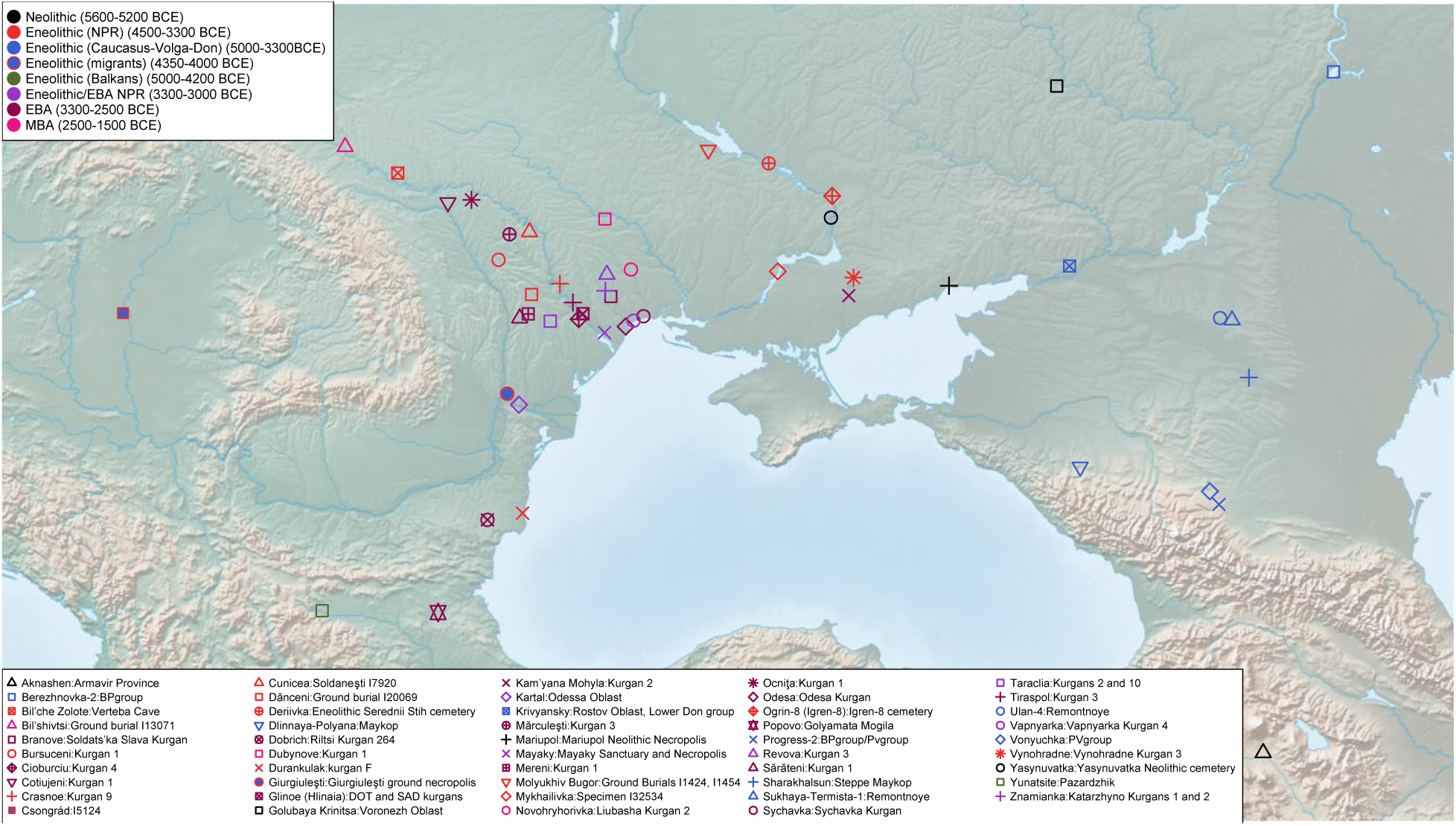
Map of sampling locations including newly generated data and key context populations.

Genome-wide ancient DNA studies have revealed that from the beginning of the Holocene to the end of the Neolithic (approximately 9200-5000 BCE), the genetic ancestry of hunter-gatherer groups in the NPR and adjacent areas was derived from a mixture of ancestral populations whose ancestry was on a genetic gradient ranging in the west from “Western Hunter-Gatherers” (WHGs) and “Balkan Hunter Gatherers” (BHGs) who lived in the Danubian Iron Gates region^6^, to “Eastern Hunter-Gatherers”^3^ (EHGs) in the east. In Ukraine, the transition from the Mesolithic to the Neolithic was marked by WHG admixture with the EHG ancestry of previously established local populations^6^.

During the Neolithic, after ca. 5800 BCE, the western NPR saw an expansion of Balkan and central European farming groups, such as Criș, Starčevo, and LBK, carrying Early European Farmer (EEF) ancestry, who, in turn, were descended from Anatolian Neolithic Farmers (ANF) with different proportions of WHG admixture^12^. In the northeastern NPR, the Neolithic populations of the Dnipro Valley continued to retain the EHG/WHG-based genetic ancestry^6^.

In the early Eneolithic (ca. 4800 BCE), farming groups of the Cucuteni-Trypillia archaeological complex (CTAC) began spreading over the forest-steppe part of the western NPR, reaching the middle Dnipro Valley by the first half of the 4^th^ millennium BCE^13^. Archaeologists trace the origin of CTAC to western Transylvania ^13,14^. The genetic ancestry of CTAC was primarily EEF-derived with admixture from WHG, EHG and Caucasus Hunter-Gatherers (CHG) ^6,15–18^.

During their eastward expansion, CTAC encountered mobile steppe communities of the Serednii Stih archaeological complex (SSAC)^13^, which likely emerged from the Azov-Dnipro-Donets area in the first half of the 5^th^ millennium BCE^19–21^. The presence of early SSAC in the Azov steppe ca. 4700-4500 BCE is supported by Sr isotope analysis of an early SSAC individual from the Mariupol necropolis (Supplementary Information, section 1). However, knowledge about the genetic ancestry of steppe populations like SSAC (referred to as “steppe ancestry”^3–6,10,13^) has been limited until now due to small sample sizes which revealed highly variable ancestry^6,13,18^.

In the 4^th^ millennium BCE, a distinctive archaeological complex known as Usatove was established in the northwestern NPR. Sampled individuals from Usatove harbored EEF and steppe ancestries, as well as a Caucasus Eneolithic/Maykop-related genetic component^5^, but the knowledge of the proximate sources of the composing ancestries has been unclear.

In the second half of the 4^th^ millennium BCE, the NPR witnessed an expanding diversity of archaeological groups, characterized by distinct burial rites and pottery types/techniques, and increased mobility, possibly including wheeled wagon transportation^2^. This diversity came to an end in the last third of the 4^th^ millennium with the expansion of the YAC, persisting into the Early Bronze Age (EBA) during the first half of the following millennium.

Genetic ancestry data on the Epipaleolithic-Early Bronze Age populations of the NPR come from a limited number of sites, hampering the understanding of population dynamics, particularly during the time that preceded a genetic turnover in Europe precipitated by YAC-related people^3,4,10,22^. This report analyses prehistoric NPR individuals from a much wider selection of archaeological sites than has previously been available, including substantially larger sample sizes from key groups, in particular CTAC, Usatove, and SSAC. Co-analyzing with the data reported in the linked paper^7^, we examine the contribution of these groups to the genetic ancestry of YAC and have a particular focus on integrating the results of the present study with the archaeological evidence to produce a holistic picture of genetic and archaeological transformations preceding and following the formation of the Yamna.

## Results

We sequenced ancient DNA for 78 ancient individuals from the NPR from the Neolithic to the Bronze Age. For 73 we report whole genome data for the first time including an individual from the Neolithic Mariupol necropolis, 11 SSAC individuals from Ukraine, 10 CTAC individuals from Moldova and Ukraine, and 23 YAC individuals from Moldova and Ukraine; for five individuals with previously reported results, we increased data quality (Online Table 1). To generate these data, we sampled 203 skeletal elements and built 452 next-generation sequencing libraries; after screening we took 235 forward into analysis (Online Table 2). We enriched our analyses by generating 51 new radiocarbon dates (Online Table 3). We co-analysed these data with that from a parallel study of steppe populations including 299 newly reported individuals and 55 individuals with improved data^7^; both studies co-analyze the full dataset.

**Table 1.**
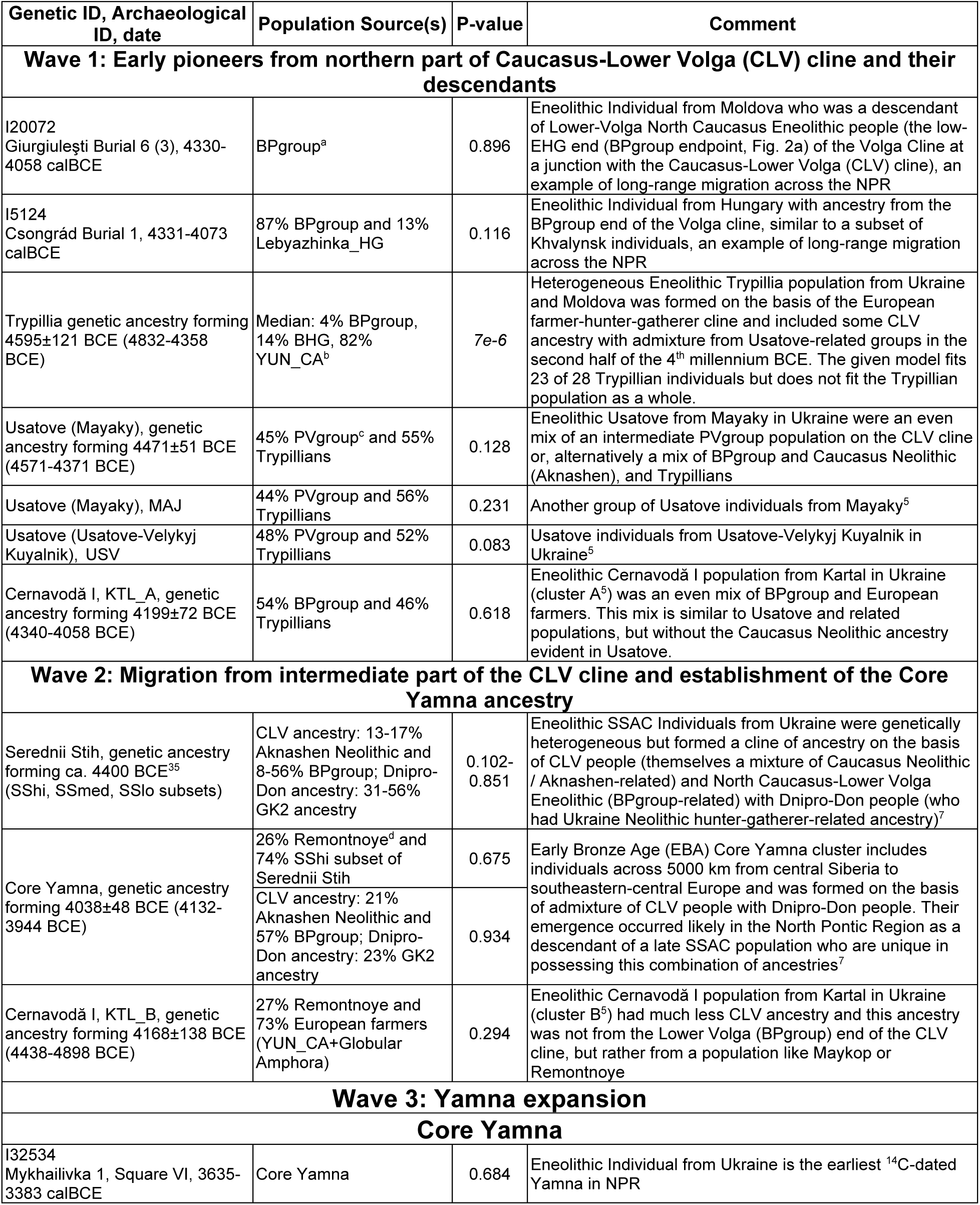
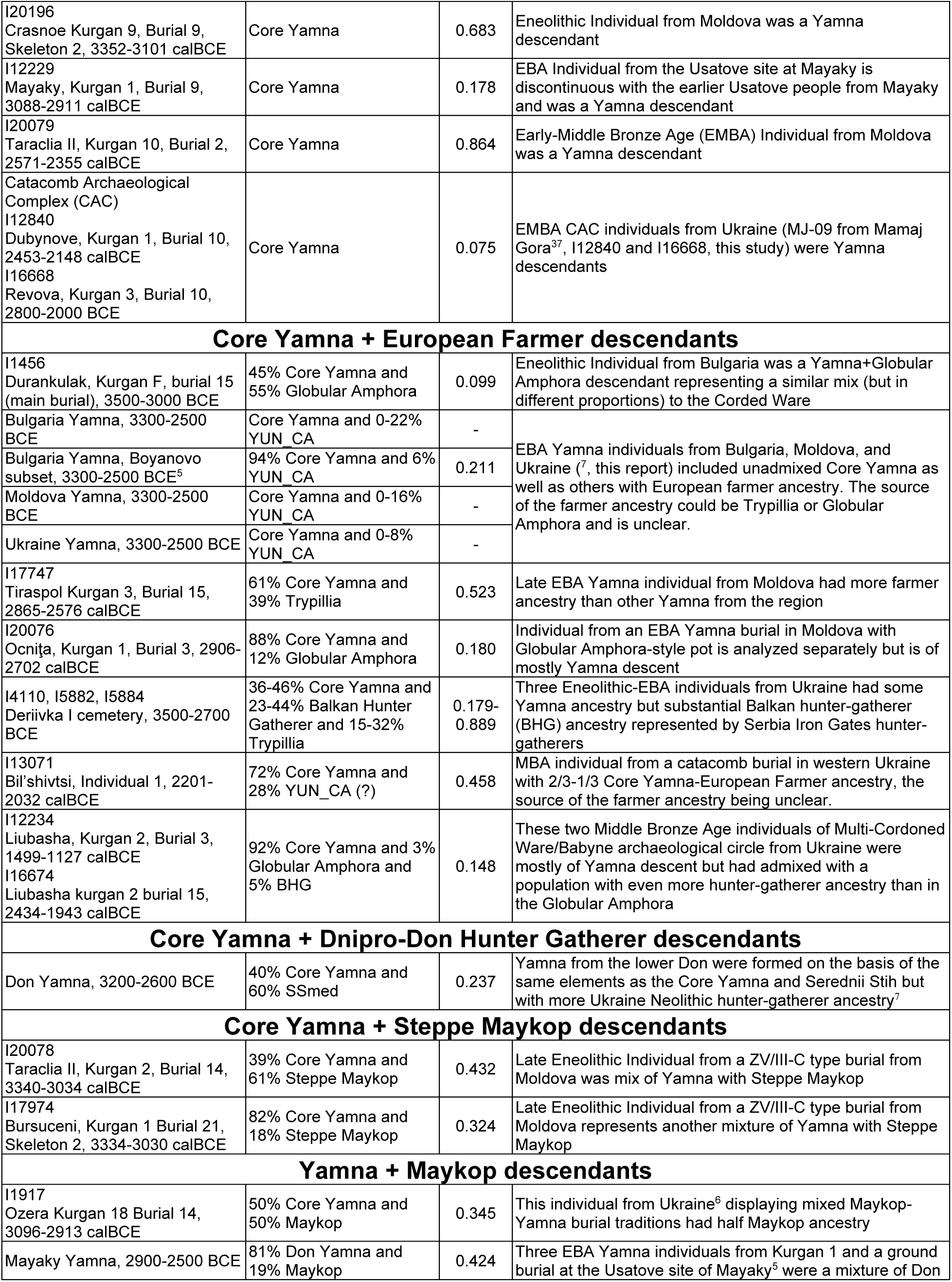
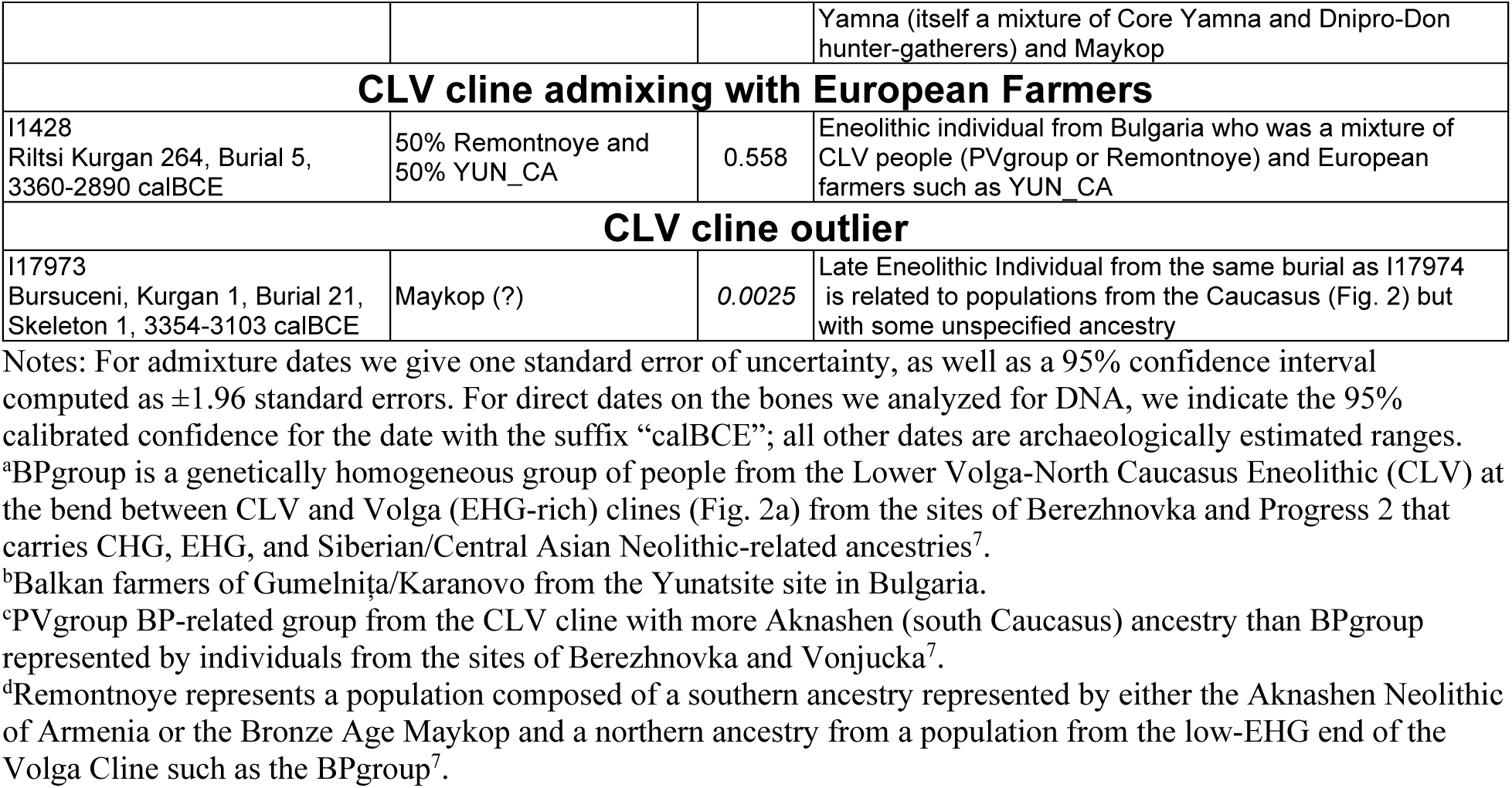
A compendium of the ancestral landscape of the North Pontic Region in the Eneolithic and Early Bronze Age (ca. 4500-2500 BCE) showing two waves of Caucasus-Lower Volga (CLV) cline ancestry migration in the NPR.

To obtain a qualitative picture of population structure in the NPR, we began by using smartpca^23^ to perform principal component analysis (PCA) using the same set of populations to form the axes as in^7^ (Fig. 2a).

**Fig. 2:**
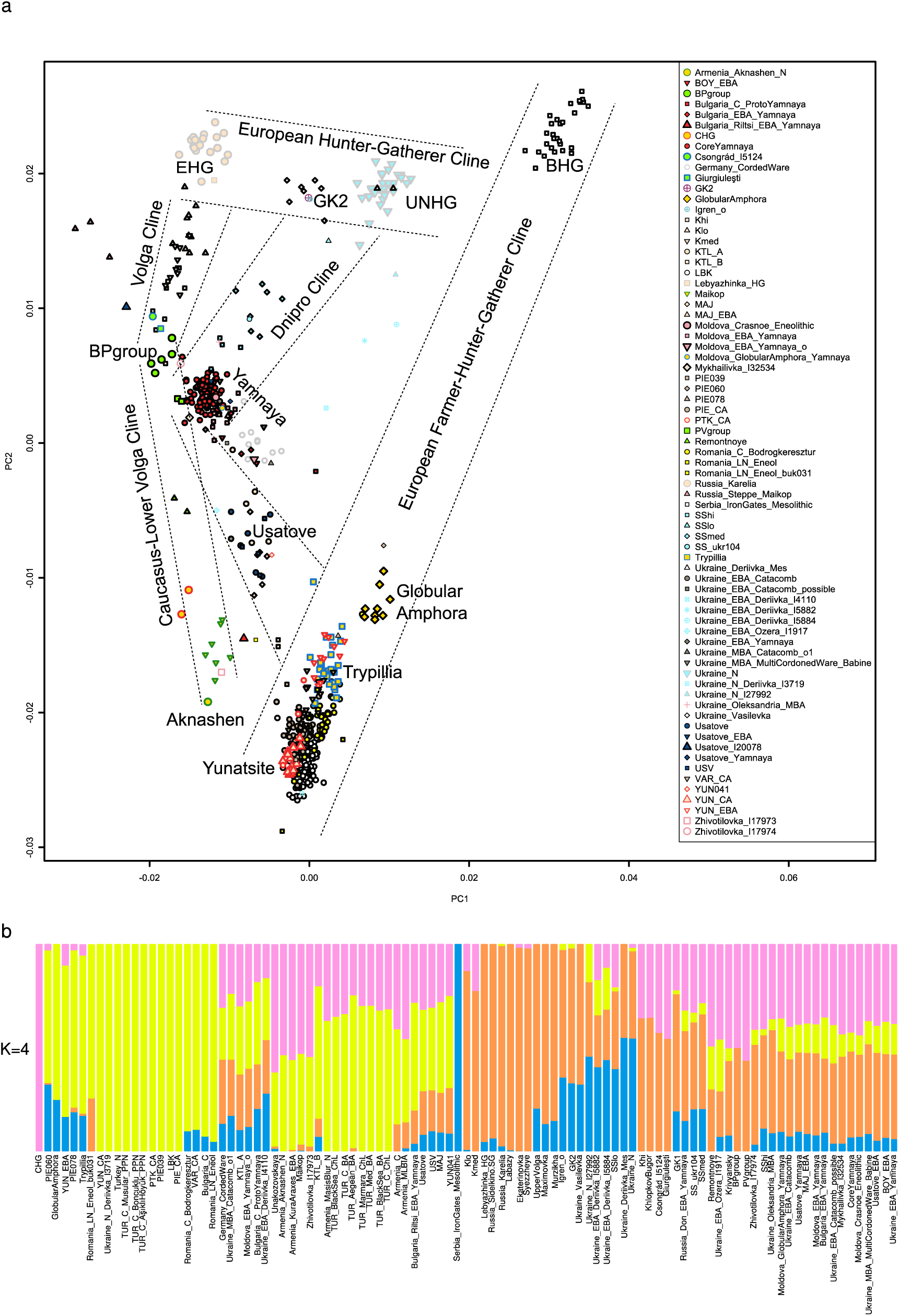
**a, PCA of the NPR samples in relation to the three steppe clines and respective samples from**^7^**. b, Unsupervised ADMIXTURE summary graph of populations and individuals from this report and**^7^.

The PCA reveals five major clines. Four of them—the Caucasus-Lower Volga (CLV) Cline, the Volga Cline, the Dnipro Cline, and the European Hunter-Gatherer (EuHG) Cline—are described formally in the accompanying paper^7^. The fourth, the European Farmer and Hunter Gatherer cline (EFHG), is formed by European farmers (central European LBK and populations related to Gumelnița/Karanovo from the Yunatsite site in Bulgaria (Yunatsite Chalcolithic, YUN_CA), on one side, and BHG, on the other (Fig. 2a).

The UNHG individuals presented in this report are located on the “eastern” end of the EuHG cline towards the BHG and at the “northern” edge of the Dnipro cline. This PCA placement suggests that UNHG contributed to the later (Eneolithic and Bronze Age (BA)) people on the Dnipro cline that are positioned along the length of that cline, with Core Yamna^7^ at the “southern” end.

The Eneolithic (apart from the SSAC) and BA individuals in Fig. 2a are mostly located towards the “farmer” end of the EFHG cline. Four NPR individuals form a cline stretching from the Core Yamna cluster towards steppe Maykop and traversing the CLV-Volga cline proximate to a key Eneolithic population represented by the Berezhnovka-2-Progress 2 individuals (BPgroup), a genetically homogeneous people between the northeast Caucasus and lower Volga that can be approximately modeled as a mixture of EHG, CHG, and Siberian/Central Asian Neolithic ancestries^7^. Two of these (I20078 and I17974) are late Eneolithic (3300-3000 BCE) individuals from Moldova. The other two, I18740 from Hungary^7^ and I20072 from Moldova, dated to ca.

4300-4000 BCE, are archaeologically associated with the Volga-Caucasus – lower Dnipro pulse of the steppe people that left “ochre graves” across the NPR and adjacent Balkan-Carpathian area^24,25^.

### Formal modeling of sources of Neolithic NPR ancestry

We computed *f3* statistics with Ukraine Neolithic as a target and a wide variety of possible sources (Supplementary Information, section 2; Extended Data Table 1). Consistent with their position in the PCA on Fig. 2a, the Ukraine Neolithic population is admixed with the most significantly negative (Z=-17.2) statistic when a sample from Karelia in northwestern Russia (EHG) and the BHG are used as sources, suggesting that the Ukraine Neolithic population is, to a first approximation, composed of sources related to the EHG and BHG populations.

However, it is evident from the PCA in Fig. 2a that the UNHG end of the EuHG cline is shifted towards populations with EEF ancestry. In unsupervised ADMIXTURE analysis (Supplementary Information, section 3; Fig. 2b), we find that the UNHG are assigned small components of Anatolian Farmer/CHG ancestry, not present in other Mesolithic Deriivka (Dnipro Valley), EHG (Karelia) or BHG (Iron Gates) groups. When samples from individuals labeled Ukraine_N (UNHGs) are modeled with other EuHG populations from^7^ only a single 2-source model (p=0.576) with 72.5±2.9% GK2 from the Golyubaya Krinitsa site on the Lower Don^7^ and 27.5±2.9% BHG ancestry, remains viable (here and in what follows, we indicate statistical uncertainty through standard errors; a 95% confidence interval corresponds to 1.96 standard errors in either direction of the point estimate). A fitting to a broader cline between EHG and BHG as a mixture of these two sources with either Lebyazhinka or Karelia as the EHG source, fails (p<1e-9) and qpAdm output suggests that these models underestimate shared genetic drift with Turkey_N (Z<-3.5).

To obtain insights into population admixture histories that could explain these patterns, we explored 3-source models detailed in Supplementary Information, section 2. The feasible models all include EHG-BHG sources (Lebyazhinka and BHG), but they all also include ∼7-9% of EEF ancestry, with the source of this ancestry (in a specific EEF-derived group) being unclear. The presence of EEF ancestry (who were largely of Anatolian Neolithic-related origin) accounts for the underestimated drift with Turkey_N in the model without such ancestry.

To test whether the inferred EEF ancestry is a population-wide feature of UNHGs, we fit a model that included central European LBK farmer ancestry representing EEF populations to 35 individuals with the Ukraine_N label (Supplementary Information, section 2; Extended Data Table 2). The results show that EEF ancestry is a general feature of UNHG populations, and thus this pattern is not driven by a few outliers.

The UNHG was modeled with significant BHG and EHG ancestry and represents an *increase* of BHG ancestry relative to the Mesolithic specimens from Vasylivka III^6^ and Vasylivka I^26^ (Fig. 2). A migration of people from the Iron Gates area in the Dnipro Valley in the 7^th^ millennium BCE^27^ may be responsible for this shift. As the BHG population from the Iron Gates has been shown to carry sporadic EEF ancestry^6^, the admixture of Iron Gates migrants could be a way to account for both BHG and EEF admixture compared to the Mesolithic Vasylivka.

Diverse hunter-gatherers of WHG-EHG mixed background in Sweden^3,28,29^ and Latvia provide no evidence for the EEF ancestry we detect in the UNHG (Supplementary Information, section 2), highlighting the uniqueness of the UNHG in that respect. As an additional control, we used the Pitted Ware/Battle Axe Culture populations from Ajvide in Sweden^30,31^ and Västerbjers^32^, finding that these populations in which EEF ancestry was incorporated into groups of predominantly hunter-gatherer background are correctly inferred by our model to have ∼1/5 EEF-related ancestry. Our finding of EEF-related ancestry in Ukraine Neolithic hunter-gatherers provides a separate and much earlier instance of the incorporation of farmer ancestry into the hunter-gatherer communities at the periphery of the Neolithic expansion in Europe.

UNHG individuals I31730 and I1738 that failed the LBK-EHG-BHG model can be modeled with CHG instead of LBK as a source (Extended Data Table 2), suggestive of CHG-related ancestry extending past the middle Don ^7,33^ to the Azov Sea coast and the Dnipro Valley during the second half of the 6^th^ millennium BCE. This sporadic and isolated CHG admixture in UNHG reflects a qualitatively different phenomenon from the generalized shift of ancestry towards the CLV cline of all Serednii Stih and Yamna individuals of the Dnipro Cline (Fig. 2a). Nonetheless, this find extends the zone of early contacts with the Caucasus that were also transforming populations of the Don and Volga rivers in the east^7^ and generating ancestry profiles radiating out of the lower Volga-Caucasus area in the Eneolithic.

### CLV cline admixture and long-range migration in the early Eneolithic Pontic steppe

Serednii Stih culture individuals had highly variable genetic ancestry—we subdivided them into “SShi”, “SSmed”, and “SSlo” subsets based on their degree of relatedness to UNHG—and their relationship with individuals on the three steppe clines are examined in detail in^7^. In that study, Serednii Stih could be modeled without European farmer populations as sources. A summary of our findings is that the Serednii Stih can be modeled with one source being the Core Yamna as the endpoint of the Dnipro cline (a proxy for earlier populations in the Eneolithic for which the Yamna descend with little or no mixture^7^), and Dnipro-Don HGs (UNHGs or GK2). Because Core Yamna themselves are formed as mixture of about 2/3 ancestry of populations of the CLV cline (proxied by PVgroup, BPgroup, Remontnoye, or Maykop) mixed with Dnipro-Don HGs^7^, the SSAC ancestry formation can be seen as the result of the fusion of CLV cline migrants with Dnipro-Don HGs.

A SSAC individual ukr104^18^ (the same as I28319 of our study) clusters with the subset of SSAC individuals with medium contribution from the UNHG group (SSmed)^7^ (Fig. 2a) and forms a clade with it using qpWave (p=0.281).

The SSAC outlier from Igren-8 (I27930), a detailed analysis of which is presented in^7^, appears to be of hunter-gatherer ancestry similar to the Neolithic GK2 individual (5610-5390 BCE) from the Middle Don. These individuals were similar in their ancestry sources to much earlier Mesolithic hunter-gatherers from Vasylivka^6,26^ (Fig. 2a) and could be modeled as having ∼2/3 EHG and ∼1/3 BHG ancestry^7^. Individual I27930 thus represents a Neolithic ancestry carry-over in a burial context of SSAC (Supplementary Information, section 1), likely appearing in the Dnipro Valley as a result of a long-range migration from the Middle Don.

In the northwest NPR, individual I20072 (4330-4058 calBCE) from Giurgiulești on the Lower Danube is cladal with the Lower Volga-North Caucasus Eneolithic groups (BPgroup and, with lower confidence, PVgroup, Supplementary Information, section 2) and, along with contemporaneous Csongrád individual from Hungary, represents an example of long-distance migration, across an even larger range than individual I27930 from Igren, spanning from the Volga to the heart of Central Europe.

### Trypillia and Usatove

Trypillian individuals^6,15–17^ are on the farmer end of the EFHG cline in the PCA (Fig. 2a). Admixture *f*3-statistics show that they are admixed Extended Data Table 1) with a highly significant negative statistic with Yunatsite Chalcolithic^5^ and BHG^6^ as sources (Z=-23.8) which parallels the PCA in suggesting that Trypillians have more hunter-gatherer ancestry than the EEF populations such as Yunatsite or LBK, but without identifying EEF ancestry sources^34^.

When we attempt to model Trypillians as a mixture of two or three sources using qpAdm, we find no fitting model for them as a whole. We explored removing the Trypillian individual that is the strongest genetic outlier (I20069 from Dănceni, 3323-2935 calBCE). However, even after excluding I20069, we still were not able to model Trypillians successfully (p<1e-5 even for *N*=3 models).

In an alternate approach to model individuals under the “Trypillia” label, 24 of 28 Trypillian individuals can be modeled in our framework. The four exceptions are discussed in the Supplementary Information, section 2. A single model (with BPgroup, YUN_CA, and BHG) is feasible for 23 of the 24 individuals. Three other models are qualitatively similar to the identified model and are feasible for 22 out of 24 individuals, all models including some CLV (Extended Data Table 3). Our calculations show that for many individuals there is no significant BPgroup-related ancestry, but this kind of ancestry is highest in the PCA outlier (I20069; 25.8±2.4%). The BP ancestry is positive for all but three, and positive by more than two standard errors for 10 of the 28 individuals. For the 23 Trypillia individuals modeled in our framework, we estimate that their genetic ancestry is, on average, 81% Balkan Eneolithic (such as in YUN_CA), 14% BHG, and the remaining 5% comes from the CLV cline (BPgroup) (Table 1). We estimate using DATES^35^ that the formative admixture of Trypillia took place 4595±121 BCE (95% C.I. 4832-4358 BCE) (Table 1, Fig. 3).

**Fig. 3:**
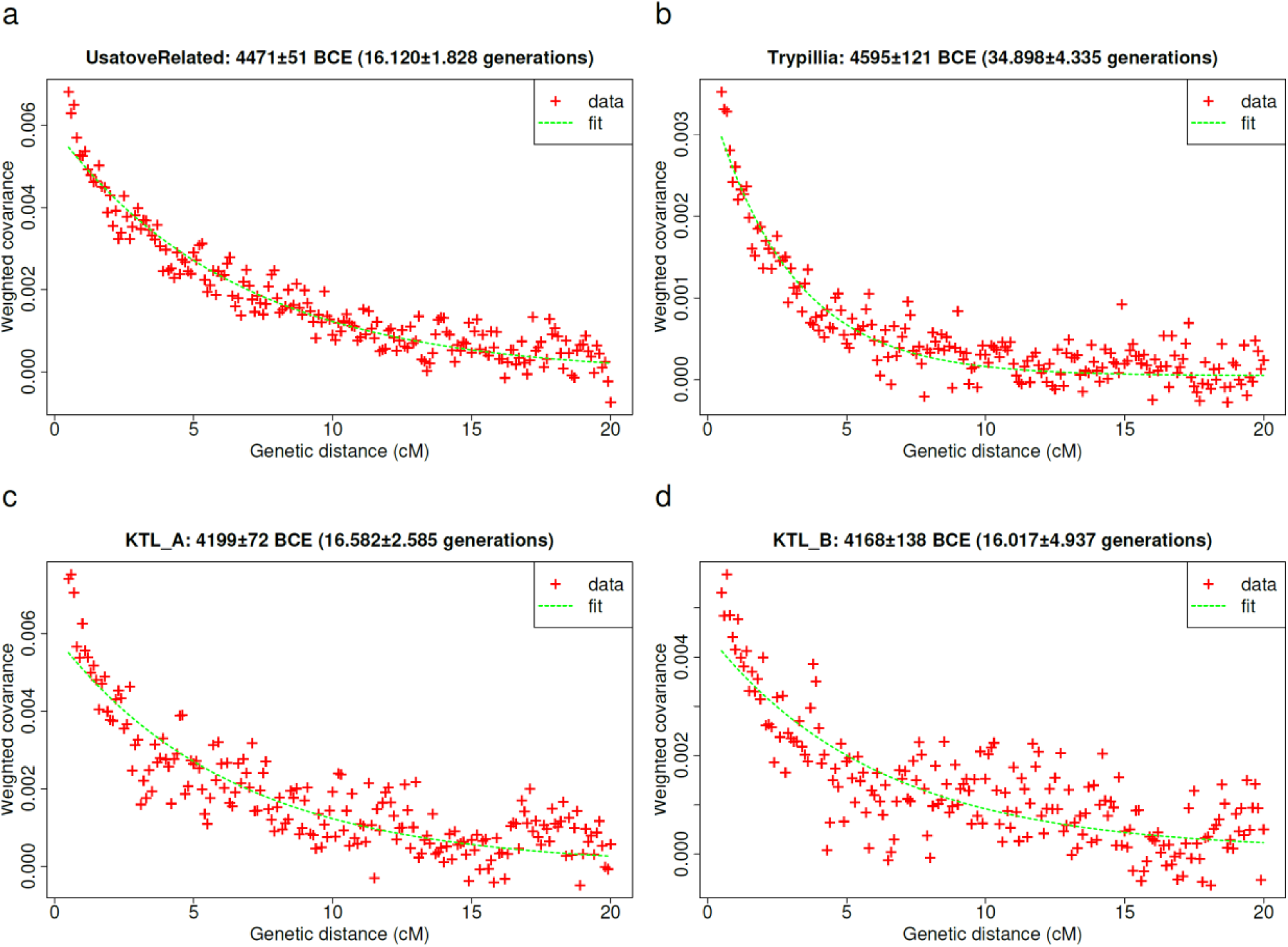
DATES estimates of admixture timing of CLV and European farmer ancestry admixture. **(a) Usatove-related individuals from this study and^5^. (b) Trypillians from this study and^17^. Kartal cluster A (c) and B (d) from^5^.**

Usatove individuals from our study, combined with previous reports to provide a substantial sample size, are genetically varied and occupy the space in the PCA between the Trypillians and the point where the three great clines of the steppe (CLV, Volga, and Dnipro) diverge from each other. Evidence of their admixed origin comes from the significantly negative admixture *f*3-statistic with Karelia (EHG) and Yunatsite Chalcolithic as sources (Z=-10.1; Extended Data Table 1) suggesting that the Usatove had ancestry from east of the NPR as well as ancestry related to European farmers. Formal modeling with qpAdm reveals that the Usatove population can be modeled uniquely (p=0.128) as a mixture of ∼45% PVgroup (and intermediate group on the CLV cline) and ∼55% Trypillians (Table 1). The model with BPgroup+Trypillians fails quite clearly (p<1e-4). A generalized 3-way model with BPgroup+Armenia_Aknashen_N (South Caucasus) ancestry, substituting PVgroup for Aknashen+BPgroup, and allowing the eastern source to vary to any position along the CLV Cline, fits (p=0.393) with an estimated 14.4±3.1% Aknashen-related ancestry (Supplementary Information, section 2), confirming that the CLV ancestry in Usatove was not from the lower Volga-centered BPgroup, but had a significant proportion of southern Caucasus Neolithic-related ancestry.

Our conclusions about the Usatove being a PVgroup+Trypillia model were confirmed on two other populations from^5^: MAJ, a different Eneolithic sample set from Mayaky (p=0.231) as well as the USV population from Usatove-Velykyj Kuyalnik (p=0.083). Neither YUN_CA nor Globular Amphora works as a source for the Usatove, which is well explained by local Trypillian origins for their farmer-related ancestry (p<1e-4). In contrast to Usatove, the CLV admixture in the Cernavodă I population from Kartal (KTL_A) in the Danube delta is best estimated as BPgroup-derived, with relatively less or no Aknashen-related ancestry (Table 1). We estimate using DATES^35^ that the formative admixture of Usatove took place 4471±51 BCE (95% C.I. 4571-4371 BCE) (Table 1, Fig. 3).

### Yamna ancestry and Maykop/steppe Maykop migrations in the Late Eneolithic NPR

Following^7^, we define a group we call “Core Yamna,” who we represent by a set of 104 individuals that are archaeologically assigned to the Yamna and Afanasievo cultures, all of whom have excellent data quality (at least 400,000 of the targeted autosomal SNPs), and that are genetically homogeneous according to qpWave (p≥0.2). In^7^ we show that these individuals descend with little or no mixture from an ancestral population that began expanding from a small founding group around 3750-3350 BCE. Core Yamna is also the largest ancestral source in all individuals carrying Yamna ancestry, who differ only in having additional admixture from local populations the Core Yamna must have encountered during their expansion^7^. In^7^ we provide multiple lines of evidence that the Core Yamna and likely the YAC itself formed in the Dnipro-Don area of the northeastern NPR region, while not being able to narrow their geographic origin further based on genetic evidence alone.

In^7^ we show that the Core Yamna can be modeled without any EEF ancestry but as a mixture of CLV and NPR hunter-gatherer groups. When we force EEF ancestry as an additional source into the Core Yamna (Supplementary Information, section 2) its proportion is not significantly greater than zero (3.2±3.1%) while that of the Caucasus Neolithic is (15.6±4.3%), suggesting that the Anatolian-related ancestry^10^ in the Core Yamna was mediated mainly from the Caucasus Neolithic populations (like Aknashen in Armenia^10^) and not from European farmers of Anatolian origin^36^. Further confirmation of this hypothesis comes from the fact that qpAdm models of exclusively CLV+NPR hunter-gatherer ancestry (Supplementary Information, section 3) conform closely with independently derived unsupervised ADMIXTURE estimates of ancestry (Fig. 2b). While EEF ancestry in the Core Yamna itself is conjectural and not necessary from the point of view of modeling this population, it was clearly present in the western Yamna from Bulgaria, Hungary, Moldova, Romania, and Serbia (^7^, Supplementary Information, section 2). In this paper, we seek to narrow down the location from which the Yamna originated, synthesizing archaeological and genetic evidence from the very beginning instead of relying on the genetic data alone and only combining with archaeological information at the end.

The chronologically earliest individuals in our sample set who are cladal with Core Yamna are I32534 (3635-3383 calBCE) from Ukraine and I20196 from Moldova (p=0.684 and p=0.683, respectively). Individual I17743 from Moldova had predominantly Core Yamna ancestry but also harbored 6.9% Balkan EEF admixture (p=0.593). Individuals I20196 and I17743 both date to ca. 3350-3100 BCE (Supplementary Information, section 2) and their contemporary I17974 from Moldova is 81.8% Core Yamna and 18.2% Steppe Maykop (p=0.324; Supplementary Information, section 2).

For individual I32534, from the second (proto-Yamna) layer of the Mykhailivka site in the lower Dnipro Valley, only the Core Yamna model remains feasible when either BPgroup or PVgroup is placed on the right set in qpAdm analysis (Supplementary Information, section 2). As a further test, when we force either EEF or UNHG as a second ancestry source (on top of the Core Yamna ancestry), neither one is significantly different from zero (|Z|<1) and both are, in fact, nominally slightly negative. Thus, there is no evidence for the presence of either the EEF or UNHG-related ancestries of the NPR region on top of the Core Yamna ancestry and I32534 is consistent with being simply a member of the Core Yamna group. Beyond qpAdm modeling, the Mykhailivka individual clusters with Core Yamna in PCA (Fig. 2a) and in unsupervised ADMIXTURE analysis (Fig. 2b). All these lines of evidence converge in showing that I32534 is indeed an early Core Yamna individual who bridges the temporal gap between the geographically proximate Late Serednii Stih populations and those of the main Yamna expansion that are sampled from south Siberia to eastern Europe and in which any associations with the locale of Yamna formation have been wiped out by thousands of kilometers of distance.

Four Yamna individuals from Ukraine are cladal with the Core Yamna group in showing no evidence of EEF admixture (Supplementary Information, section 2). Three Yamna Ukraine individuals, as well as one Catacomb Individual I12617, all from the northwest NPR, harbor significant European farmer-associated admixture from proximate sources like Bulgaria Eneolithic or Trypillia (Supplementary Information, section 2). Thus, the northwest NPR can be identified as the place in where the Yamna first received substantial EEF admixture during their western expansion.

The substantial proportion of farmer ancestry in Yamna outlier individuals I20076 and I17747 (2865-2576 calBCE) from Moldova is best fitted by Core Yamna + Trypillia or Globular Amphora models (Supplementary Information, section 2). One of the Yamna individuals from Bulgaria contained 22.3% YUN_CA admixture, while another individual from the same site was cladal with the Core Yamna (Supplementary Information, section 2). The Yamna expansion, beginning in Ukraine and reaching the South Balkans, included both individuals who maintained the Core Yamna genetic profile, as well as those who had begun to admix with local farmers, initiating the transmission of Yamna ancestry and probably Indo-European languages beyond the steppe.

We also present genetic evidence of the westward expansion of the North Caucasus/lower Volga-Don ancestry at the early stages or pre-dating the Yamna expansion. This is reflected in individuals I17974 and I20078 from Moldova, who were formed of the same Yamna+Steppe Maykop-associated admixture process, with I17974 carrying about ∼1/3 of the Steppe Maykop-associated ancestry found in I20078 (Table 1, Supplementary Information, section 2). The Caucasus affinity was also observed for individual, I17973, co-buried with I17974, who cannot be well-modeled with any of the sources available to us, but is nearest to the “southern” end of the CLV cline (Maykop of the North Caucasus (p=0.0025) or the Aknashen Neolithic of the South Caucasus (p=0.0047, Supplementary Information, section 2), which is corroborated by the position of I17973 on the PCA (Fig. 2a). In the northeastern NPR, an early Yamna individual I1917 from Ozera^6^ is best modeled as an even mix of Core Yamna and Maykop, providing, like individual I17973, a clear link to the Caucasus. More evidence for this link comes from the Early Bronze Age population from Mayaky^5^, which is discontinuous with the Usatove from the same region but represented a unique combination of 1/5 Maykop ancestry with the remainder best represented by the Yamna of the Lower Don, a population which was itself a mix of Core Yamna and NPR hunter-gatherers^7^.

### Yamna ancestry in the Bronze Age NPR

We find that individuals of the Catacomb archaeological complex (CAC), which chronologically partially overlaps and succeeds Yamna in the NPR, continued to harbor Yamna genetic ancestry. The population labeled “Ukraine_EBA_Catacomb”, including individuals I12840 and I16668 from our dataset, is cladal with the Core Yamna (p=0.075, Supplementary Information, section 1). Yamna ancestry persisted in the NPR into the second half of the 3^rd^ millennium BCE.

The Catacomb group was succeeded in the NPR by the Multi-Cordoned Ware/Babyne (MCW/B) complex (Supplementary Information, section 1). The only feasible models for the ancestry of two MCW/B individuals in our sample selection involve Core Yamna, a European farmer source, and additional hunter-gatherer ancestry above and beyond what was present in even the most hunter-gatherer admixed farmer populations of the farmer-hunter-gatherer cline (Supplementary Information, section 2). Three-way modeling with varying hunter-gatherer ancestry for BA Ukraine (Supplementary Information, section 2) confirms that MCW/B in Ukraine experienced gene flow from a population that had considerable hunter-gatherer ancestry. Such populations have been described from the BA of what is today Romania at the sites of Arman (Cârlomănești) and Târgșoru Vechi in Muntenia^10^, indicating that populations of high hunter-gatherer ancestry contributed to some post-Yamna people in the NPR and Southern Carpathians.

## Discussion

This study presents the first comprehensive reconstruction of the population dynamics in the North Pontic steppe and forest steppe, clarifying genetic transformations in this region leading up to and following the emergence of the YAC.

We demonstrate that the Neolithic populations of the Dnipro Valley were admixed, roughly with BHG and EHG sources, along with approximately ∼7-9% EEF ancestry throughout the UNHG population except for some outliers such as individual I27992 from Yasynyvatka (27±6.0% EEF, this report) and an unadmixed EEF individual I3719 from Deriivka I^6^ (103.5±1.6% EEF). CHG ancestry is also sporadically present at comparably low levels relative to the EEF ancestry (∼7-10%), including in the region most proximate to the North Caucasus in the NPR Neolithic necropolis at Mariupol. The proximal sources of EEF ancestry in UNHGs remain unclear but may have been mediated by BHG migrants in the Dnipro Valley or individuals of EEF genetic background such as individual I3719^6^ that were included in UNHG communities.

We infer that the Eneolithic Trypillia population was mainly formed from the sources along the EFHG cline that received limited (∼5%) admixture from people that had BPgroup CLV ancestry. Usatove was formed on the basis of PVgroup CLV people evenly intermixing with Trypillian ancestry. The Trypillia and Usatove populations thus both harbored ancestry from near the bend of the Volga-CLV clines (Fig. 2a) and differed from each other in that this ancestry was minor in Trypillia and more BPgroup-related, while comprising approximately half of the Usatove ancestry and being more PVgroup-related (shifted towards the Maykop-Aknashen end of the CLV cline).

The evidence from Usatove and Trypillia clarifies the process of the CLV admixture in the NPR in the Eneolithic. As people bearing the Volga-CLV ancestry moved across the North Pontic steppe and into the Balkan-Carpathian region, they encountered local farmer populations. Some carriers of Volga-CLV ancestry, as in Giurgiuleşti and Csongrád, reached the Balkans and Carpathian region with no genetic admixture with the people they encountered along the way; if some migrants did admix, they left little demographic impact, perhaps because they were small in number relative to the local populations. In contrast, their farmer counterparts in the NPR such as Trypillia were more demographically affected, incorporating the Volga-CLV incomers’ genetic ancestry as well as elements of their material culture. An intriguing possibility raised by our findings is that Usatove was formed around an outpost in the Danube-Dniester interfluve to oversee the economic interests of Trypillia, on the one hand, and early carriers of the southern Caucasus-enriched PVgroup of the CLV cline ancestry, on the other. A similar scenario is feasible for the Cernavodă I population of Kartal_A, but with BPgroup-derived carriers of CLV ancestry such as in Giurgiuleşti and Csongrád individuals. Alternatively, Usatove and Kartal A could have formed as a “commonwealth” of co-existing and interdependent cultures in which Trypillia and populations from the Caucasus-Volga both participated. A third potential scenario places egalitarian Trypillians under the dominance of hierarchically organized patriarchal societies carrying CLV ancestry, extending into the northwestern NPR.

The other great Eneolithic culture of the NPR, the Serednii Stih, also consisted of people who received varying degrees of CLV and UNHG-related ancestries^7^. The Serednii Stih cline lacks appreciable EEF ancestry in contrast to Usatove and, especially, Trypillia. The results in^7^ and the current analysis establish the Core Yamna as a late Serednii Stih-derived population that had more CLV ancestry than the sampled Serednii Stih individuals but was made of the same CLV and UNHG-GK2 derived components. CLV ancestry comprised only 5% in Trypillia and roughly 50% in Usatove, while in Yamna it was 77%^7^. In Usatove, ca. 14% of CLV ancestry was southern Caucasus Aknashen-related (Supplementary Information, section 2), while in the Core Yamna the Aknashen-related ancestry was ca. 21%, so this was not a single-source CLV migration into the steppe and Pontic region^7^.

Evidence presented in^7^ argues for a YAC origin in the Dnipro-Don area of the northeastern NPR. Yamna ancestry became a feature of almost all individuals in southeast Europe postdating the Yamna expansion, except for the southernmost corner of the Balkan Peninsula in the Aegean^10,38–40^. The expansion of the YAC eastward brought its bearers to near the foot of the Urals (where the Samara Yamna were sampled) and to west Siberia, where they formed the Afanasievo culture of the Altai.

The existence of unadmixed Core Yamna in a wide area from the Altai to Bulgaria can be seen as evidence of the rapidity of the Yamna expansion, providing little opportunity for admixture during its initial pulse. The question of whether the remarkable homogeneity of the Core Yamna cluster was a consequence of relative isolation during their formative period or a cultural avoidance of heterogamy that was later abandoned, remains to be answered.

During their western expansion, the Yamna absorbed EEF ancestry from the populations of the west-northwest edge of the NPR and southeastern Europe, while at the same time integrating individuals with ancestries from the Don Yamna or the Maykop and Steppe Maykop. Thus, the Yamna variably incorporated ancestries from nearly every encountered group during their expansion pulse. This integrative nature of their communities, coupled with their remarkable mobility, likely contributed to the Yamna’s success in disseminating their Indo-European language and culture across geographic and population boundaries.

The chronologically earliest (3635-3383 calBCE) individual with the Core Yamna ancestry comes from the Mykhailivka settlement displaying a succession of uninterrupted cultural layers from the late Eneolithic to the EBA^41,42^, without the evidence of site depopulation moving into the Yamna period seen at almost all other Eneolithic sites. In the context of the archaeological evidence, the presented results increase the plausibility of arguments that the lower Dnipro, specifically the area around the Mykhailivka site at a crossroads of ancient steppe “highway” network across the Pontic-Caspian steppe (Supplementary Information, section 1), is a place where Yamna first emerged.

The groups that succeeded Yamna in the NPR in the second half of the 3^rd^ millennium BCE continued to harbor Yamna genetic ancestry, as well as displaying a resurgence of hunter-gatherer ancestry towards the Middle Bronze Age, the latter evidenced by the MCW/B individuals from Ukraine and Romania. The geographic dispersal of individuals with MCW/B genetic ancestry may reflect high mobility of this group, like that of the Yamna but smaller in scale.

### The three waves of CLV ancestry expansion in the NPR

Our analysis suggests a history of three partially overlapping waves of CLV migrations into the NPR in the Eneolithic (Table 1). A first and, potentially earliest wave spread before ca. 4500 BCE (the DATES-estimated Trypillia and Usatove genetic ancestry formation), bringing a mostly BPgroup/PVgroup-related pulse from the genetically “northern”/Lower Volga part of the CLV cline. It was associated with Giurgiuleşti-Csongrád Suvorove-type burials, and left an admixture in Trypillia, Usatove (with participation of the Neolithic Caucasus ancestry), and Kartal_A. A second and more protracted wave carried an intermediate part of the CLV cline, involved a more genetically “intermediate” ancestry (an example of which is Remontnoye) and became associated, in its initial pulse, with the formation of Serednii Stih ca. 4500 BCE. In its westernmost reach, the second wave extended to the northwest NPR, contributing to the formation of Kartal_B, but otherwise remaining largely contained in the Lower Dnipro Valley region, notably during the steppe “hiatus” in the late 5^th^-early 4^th^ millennium BCE, characterized by a relative lack of archaeological material.

The Core Yamna genetic mixture is estimated to have taken place at 4038±48 BCE (95% C.I.: 3944-4132 BCE)^7^, which is the height of the steppe hiatus inferred from archaeological information. It is unclear whether this date corresponds to an admixture of populations that happened very rapidly, or whether it corresponds to a process that unfolded over generations, in which case the date we estimate is an average. Nonetheless, it does coincide with a sharp climatic shift towards aridity and cooler temperatures. Thus, the steppe hiatus may be a reason for the emergence of the core Yamna ancestry from a nascent SSAC-derived Yamna population that was relatively isolated due to the climatic upheaval.

It is conceivable that the steppe groups of the post-hiatus chronological period (3900-3300 BCE) such as Lower Mykhailivka, Mykhailivka 2 (proto-Yamna), and Konstantinovka in the lower Don, forming under the increasing influence of the North Caucasus^42^, stem from a SSAC-derived steppe population that became isolated during the climate-influenced hiatus, but re-emerged (now as proto-Yamna) following it. Such a scenario would explain two features in the population history of the Core Yamna: its population bottleneck prior to ca. 3750-3350 BCE^7^, potentially occurring in the context of climatic-induced isolation, and its genetic position at the low-UNHG end of the Dnipro Cline as the result of the proximity and influence of the North Caucasus. Genetically, the Core Yamna can be modeled as a mixture of ∼3/4 of the high CLV ancestry subset of Serednii Stih (SShi) and ∼1/4 of the genetically intermediate (along the CLV cline) population represented by two individuals from the Manych Depression at Remontnoye (Table 1; ^7^) who date to this key period (4152-3637BCE). In this scenario, the individual from Mykhailivka represents a proto-Yamna population near the geographical origin of the Core Yamna and sampled from the time where its genetic distinctiveness had already appeared. Other early individuals from the NPR, such as Bursuceni and Taraclia II.2.14, also carried the Core Yamna ancestry, while connecting the NPR with other populations of the North Caucasus^43,44^.

The third wave of CLV ancestry expansion is that of the YAC proper, beginning ca. 3300 BCE and lasting into the middle of the following millennium. All three expansion waves stem from different points on the geographically and genetically diverse CLV cline.

It is remarkable that the three genetic waves of CLV ancestry expansion align, spatially and temporally, with the three waves of Kurgan People proposed by Marija Gimbutas in the 1950s to explain the spread of Indo-European influences and the fall of “Old Europe” (summarized in^1,45^). In Gimbutas’ theory and in our genetic analysis, the three waves originated in the Lower Volga-North Caucasus area and acted as constituent elements of a single process that unfolded in time and space throughout the Eneolithic and into the Bronze Age, transforming the cultural landscape of Western Eurasia. We must note, however, that Gimbutas envisioned the spread of Kurgan ancestry as a military conquest and emphasized *cultural* transformation of the conquered people encountered by the Kurgan culture bearers. Our results present evidence of massive *genetic* transformations effected by the spread of CLV ancestry during Waves 1 and 2, and especially, with the spread of the Yamna during Wave 3. Such genetic changes must have involved complex cultural dynamics, in which both conflict and peaceful synthesis may have played a role. Future studies can further explore the cultural impact these three expansion waves brought about, informed by the new understanding of the immense genetic impacts that accompanied them.

## Conclusion

Our detailed survey of individuals from the Neolithic to Bronze Age in the NPR shows a shifting landscape of ancestry. The earliest inhabitants of the NPR that carried BHG/WHG-EHG-EEF ancestry components had lived there from Paleolithic to Neolithic times. Caucasus ancestry, having made its tentative and sporadic appearance already among the UNHG of the NPR, appears in large proportion among all the Eneolithic NPR populations that followed, derived from the diverse people of the Caucasus-Lower Volga cline who appear in the Lower Danube and its tributaries in central Europe unadmixed, and leaving traces of their ancestry in larger NPR-area populations (as in Trypillia), or as equipotent ancestry contributors (as in Usatove and Kartal_A), and as highly variable clinal populations (as in the Serednii Stih). Many diverse blends of autochthonous NPR inhabitants and CLV newcomers were formed in which both farmers and hunter-gatherers of the NPR contributed and in which people from different sections of the CLV cline participated (Table 1). The distinctive Serednii Stih-descended population ancestral to the Core Yamna dominates, after its 4^th^ millennium BCE appearance and subsequent expansion, absorbing and incorporating, in diverse blends of its own (Table 1), ancestries of the people they encountered along the way. The later history of the NPR, in which Cimmerians, Scythians, Greeks, Sarmatians, Turks, Bulgars, and Slavs, and numerous others who made their mark on the cultural and genetic landscape, mirrors the region’s more distant past that we study here: a continuous process of transformation and change that not only shaped its modern inheritors, but also played a central role in shaping events across the wider Eurasian continent.

## Materials and Methods

### Wet laboratory work

In clean rooms where the goal was to protect bones and teeth from contamination by the individuals handling them, we processed human skeletal remains into powder^46^, extracted DNA using a method designed to retain short molecules^46–48^ in some cases using automated liquid handlers^49^, and converted the extracts into double-stranded^50^ and single-stranded^51^ libraries, which were molecularly barcoded with appended dual barcodes (for double-stranded libraries) and dual indices (for both double-stranded and single-stranded libraries) to allow them to be pooled together and then bioinformatically deconvoluted at the analysis stage. We enriched the libraries for sequences overlapping more than 1.2 million SNPs as well as the mitochondrial genome^52^, and then sequenced on NextSeq500, HiSeqX, or NovaSeq instruments, targeting on the order of a hundred thousand molecules for unenriched libraries and on the order of 30 million molecules for enriched one. Online Table 2 provides information on each library we analyzed.

### Bioinformatic analysis

Following sequencing, we used the identification markers (barcodes and indices) to demultipex reads into the to the appropriate library, before trimming these and sequence adapters. Paired-end reads were then merged requiring an overlap of at least 15 base pairs (allowing for 1 mismatch), using a modified version of SeqPrep 1.1 (https://github.com/jstjohn/SeqPrep); at overlapping bases, we selected the highest quality nucleotide to represent the sequence at that position. We aligned sequences to both the human reference genome sequence (hg19) (https://www.internationalgenome.org/category/grch37/) and to the inferred ancestral Reconstructed Sapiens Reference Sequence (RSRS) mitochondrial sequence^53^, using BWA’s samse command^54^. We removed duplicated molecules based on having the same start/stop positions and orientation in their alignment and the same barcodes. The computational pipelines we used are publicly available on GitHub at https://github.com/dReichLab/ADNA-Tools and https://github.com/dReichLab/adna-workflow. We call variants by using a ‘pseudohaploid genotyping’ approach, where a single base is randomly selected from a pool of possible bases at each SNP, filtering by a minimum mapping quality of least 10, and base quality of at least 20, trimming each read by two base pairs to remove damage artifacts. To assess ancient DNA authenticity, we used both *contamMix-1.0.1051*^55^ to search for heterogeneity in mitochondrial DNA sequences which are expected to be non-variable in uncontaminated individuals, and ANGSD (ref) to search for heterogeneity in X chromosome sequences which should be non-variable in contaminated male individuals^56^. We also evaluated authenticity by searching for an increase in cytosine-to-thymine errors in the final nucleotide (in untrimmed reads) which is expected for genuine ancient DNA^57^ and by computing the ratio of Y chromosome to sum of X and Y chromosome sequences which is expected to be very low for females and to have a very much higher value for males. We determined a consensus sequence for mitochondrial DNA using *bcftools* (https://github.com/samtools/bcftools) and *SAMTools*^58^ requiring a minimum of 2-fold coverage to call the nucleotide and a majority rule to determine its value. We used *HaploGrep2* to determine the mitochondrial haplogroups based on this consensus sequence, leveraging the phylotree database (mtDNA tree build 17)^59^.

### Population genetic analysis

We performed principal components in smartpca^23^ using lsqproject: YES and newshrink: YES parameters and the populations OberkasselCluster (set of trans-Alpine WHG individuals identified in^26^), Russia_Firsovo_N, Iran_HajjiFiruz_C^9^, Iran_C_SehGabi^60^, Iran_C_TepeHissar^61^, Israel_C^62^, Germany_EN_LBK^3,12,28,63^ to form the axes (Fig. 2).

We used qpWave and qpAdm^3,64^ to test whether *n*+1 “left” populations (one Test and *n* sources) are consistent with descending from *n* ancestral sources with respect to a set of Right populations as in^7^ (OldAfrica^65–67^, Russia_AfontovaGora3^68^, CHG^69^, Iran_GanjDareh_N^60^, Italy_Villabruna^68^, Russia_Sidelkino.SG^8^, Turkey_N^28^).

We performed unsupervised ADMIXTURE analysis^70^ using a new methodology of “summary individuals” (SI) that prevents the formation of population-specific ancestry components, as a complementary approach (other than qpAdm) to assess the ancestry of diverse population from the NPR and neighboring regions (Fig. 2b).

We dated the admixture time of Usatove-related populations (individuals from Mayaky presented in this report and from Mayaky (MAJ) and Usatove-Velykyj Kuyalnik (USV) from^5^) and Trypillians, using DATES^35^ to infer the number of generations prior to the ^14^C date of the studied individuals, and converted to a calendar date assuming 28 years per generation^71^. Uncertainty ranges reflect the standard error computed by DATES and not the uncertainty of the average ^14^C date of admixed individuals.

## Acknowledgments

The authors thank David Anthony for a critical review of a manuscript draft; Natalya Burdo, Yuri Rassamakin, and Sergey Razumov for stimulating discussions; Sergey Agulnikov, Joachim Burger, Elke Kaiser, Vitali Sinica, Mykhailo Sokhatsky, and Evegenii Yarovoy for sharing samples, Iñigo Olalde for bioinformatic support, and Rebecca Bernardos, Nasreen Broomandkhoshbacht, Nicole Adamski, Matthew Ferry, Ilana Greenslade, Zhao Zhang, and Kristen Stewardson for technical support. We acknowledge the contribution of Ukrainian archaeologists Mykola Makarenko (1877-1938) and Dmytro Telegin (1919-2011) as leaders of the excavations that produced many of the samples featured in this report and for providing the theoretical groundwork that inspired many of the hypotheses tested here.

## Data Availability

Genotype data for individuals included in this study can be obtained from the Harvard Dataverse repository through the following link (XXX). The DNA sequences reported in this paper have been deposited in the European Nucleotide Archive under the accession number XXX. Other newly reported data such as radiocarbon dates and archaeological context information are included in the manuscript and supplementary files.

## Author Contributions

AGN, IL, SI, VD, ML, IP, and DR conceived the study. AGN, IL, NP, and DR supervised data analysis. AGN, SS, VR, and DR secured funding for the study. AGN, SI, MV, VD, NK, ML, IP, MK-N, SL, SM, HS, GS, and TT provided samples for the study. IL, NP, and DR supervised or performed statistical analyses. AGN, VR, SS, KC, EC, EH, LI, AML, MeM, MaM, AM, JO, LQ, JNW, FZ, SwM, and NR performed laboratory and bioinformatic analyses. AGN and AK curated the samples. NP, ML, NK, SM, SL, HS, SS, PW, and DR critically reviewed and edited manuscript files. AGN and IL wrote the manuscript with input from all co-authors.

## Ethics Statement

All applicable regulations were followed when handling human remains both in the lab and in the field. All samples originating from Ukraine were excavated or sampled from museum or archival collections in Ukraine prior to 2022. Authors obtained consent, when available, from the individuals who conducted the excavations, who are either co-authors of the study or are acknowledged for their contribution. Human remains were processed using a minimal amount of skeletal material with the goal of minimizing damage. The open-access publication of the results of this study ensures unrestricted access to the results by specialists as well as the general public. Geographic names as well as names of archaeological groups were transliterated following their spelling in the countries from which samples originate. Geographic boundaries of political entities were respected following international law.

---

**Extended Data Table 1.** Statistics of the form *f*_3_(Source_1_, Source_2_; Test). The statistic with the lowest Z-score of all the considered pairs is shown. This is the same as Table S3 in the supplement.

| Test | Source1 | Source2 | $f_3(\text{Source}_1, \text{Source}_2; \text{Test})$ | Z-score |
| --- | --- | --- | --- | --- |
| BOY_EBA | TTK | Trypillia | -0.016097 | -7.0 |
| Bulgaria_EBA_Yamna | Russia_Karelia | YUN_CA | -0.011836 | -9.2 |
| CoreYamna | Maykop | Russia_Karelia | -0.006310 | -13.6 |
| GlobularAmphora | Serbia_IronGates_Mesolithic | YUN_CA | -0.005914 | -8.2 |
| KTL_A | Russia_Karelia | YUN_CA | -0.014186 | -17.5 |
| KTL_B | Russia_Karelia | YUN_CA | -0.009922 | -9.1 |
| MAJ | Russia_Karelia | YUN_CA | -0.009438 | -12.7 |
| MAJ_EBA | GlobularAmphora | TTK | 0.004403 | 1.6 |
| Moldova_EBA_Yamna | Maykop | Russia_Karelia | -0.007198 | -10.0 |
| PIE_CA | Serbia_IronGates_Mesolithic | YUN_CA | -0.002351 | -6.9 |
| PTK_CA | TTK | YUN_CA | 0.001444 | 0.3 |
| Romania_LN_Eneol | Armenia_Aknashen_N | Serbia_IronGates_Mesolithic | 0.002525 | 0.5 |
| SShi | Armenia_Aknashen_N | Russia_Karelia | -0.010140 | -6.3 |
| SSmed | BPgroup | Serbia_IronGates_Mesolithic | -0.012501 | -10.6 |
| Trypillia | Serbia_IronGates_Mesolithic | YUN_CA | -0.008350 | -23.8 |
| Ukraine_Deriivka_Mes | Russia_Karelia | Serbia_IronGates_Mesolithic | -0.003244 | -1.3 |
| Ukraine_EBA_Catacomb | Armenia_Aknashen_N | Russia_Karelia | -0.022783 | -1.7 |
| Ukraine_EBA_Yamna | Maykop | Russia_Karelia | -0.009610 | -8.1 |
| Ukraine_MBA_MultiCordonedWare_Babyne | GK2 | YUN_CA | -0.017018 | -2.5 |
| Ukraine_N | Russia_Karelia | Serbia_IronGates_Mesolithic | -0.007871 | -17.2 |
| Ukraine_Vasilevka | Serbia_IronGates_Mesolithic | TTK | -0.005716 | -3.0 |
| Usatove | Russia_Karelia | YUN_CA | -0.008941 | -10.1 |
| USV | Russia_Karelia | YUN_CA | -0.011918 | -12.0 |
| VAR_CA | Serbia_IronGates_Mesolithic | YUN_CA | -0.003861 | -9.3 |
| YUN_EBA | Serbia_IronGates_Mesolithic | YUN_CA | -0.001677 | -2.6 |

**Extended Data Table 2.**
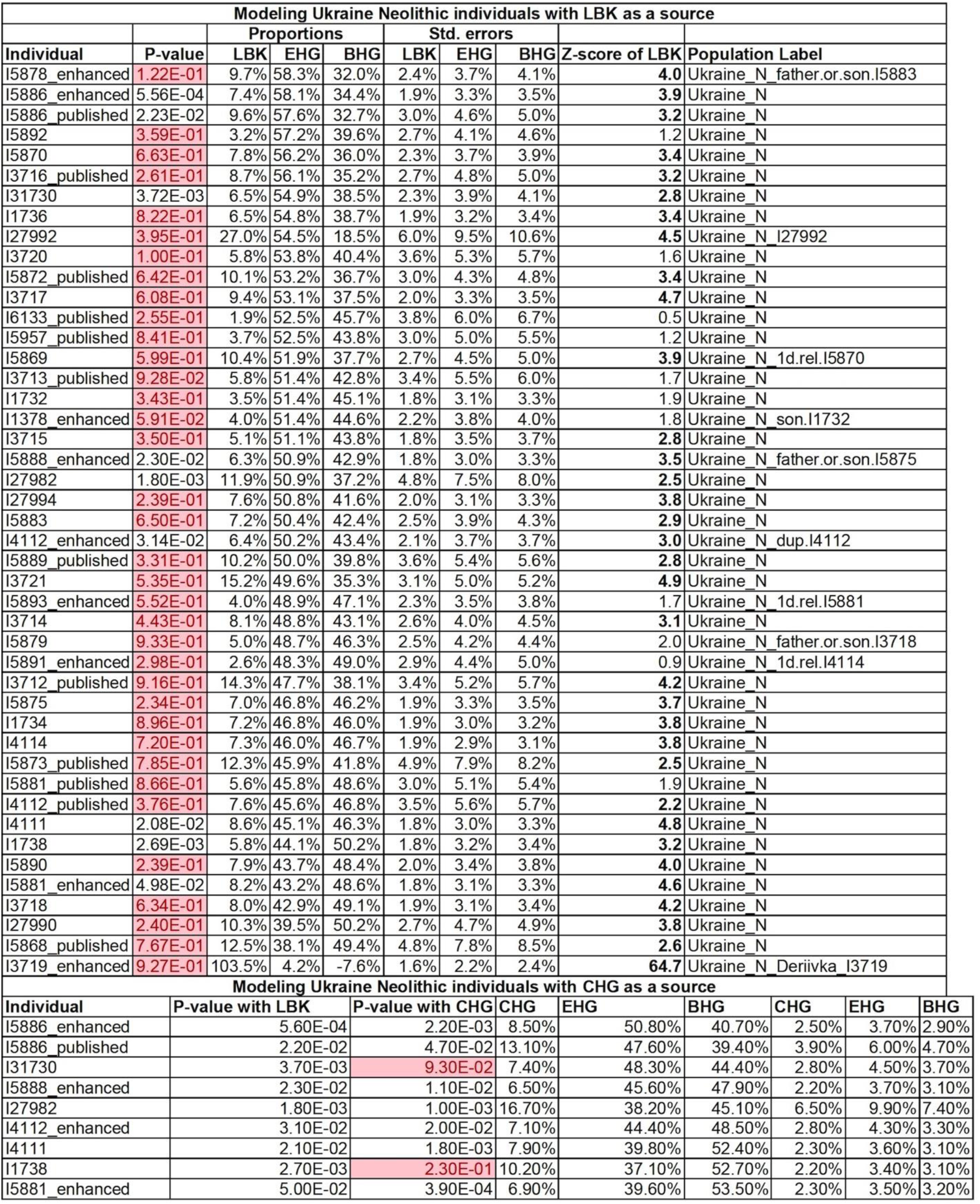
Ancestry of Ukraine Neolithic individuals. EHG=Lebyazhinka_HG; BHG=Serbia_IronGates_Mesolithic; CHG=Caucasus_Hunter_Gatherer. We include close relatives and outliers. These are the same as Tables S46 and S47 in the supplement.

**Extended Data Table 3.**
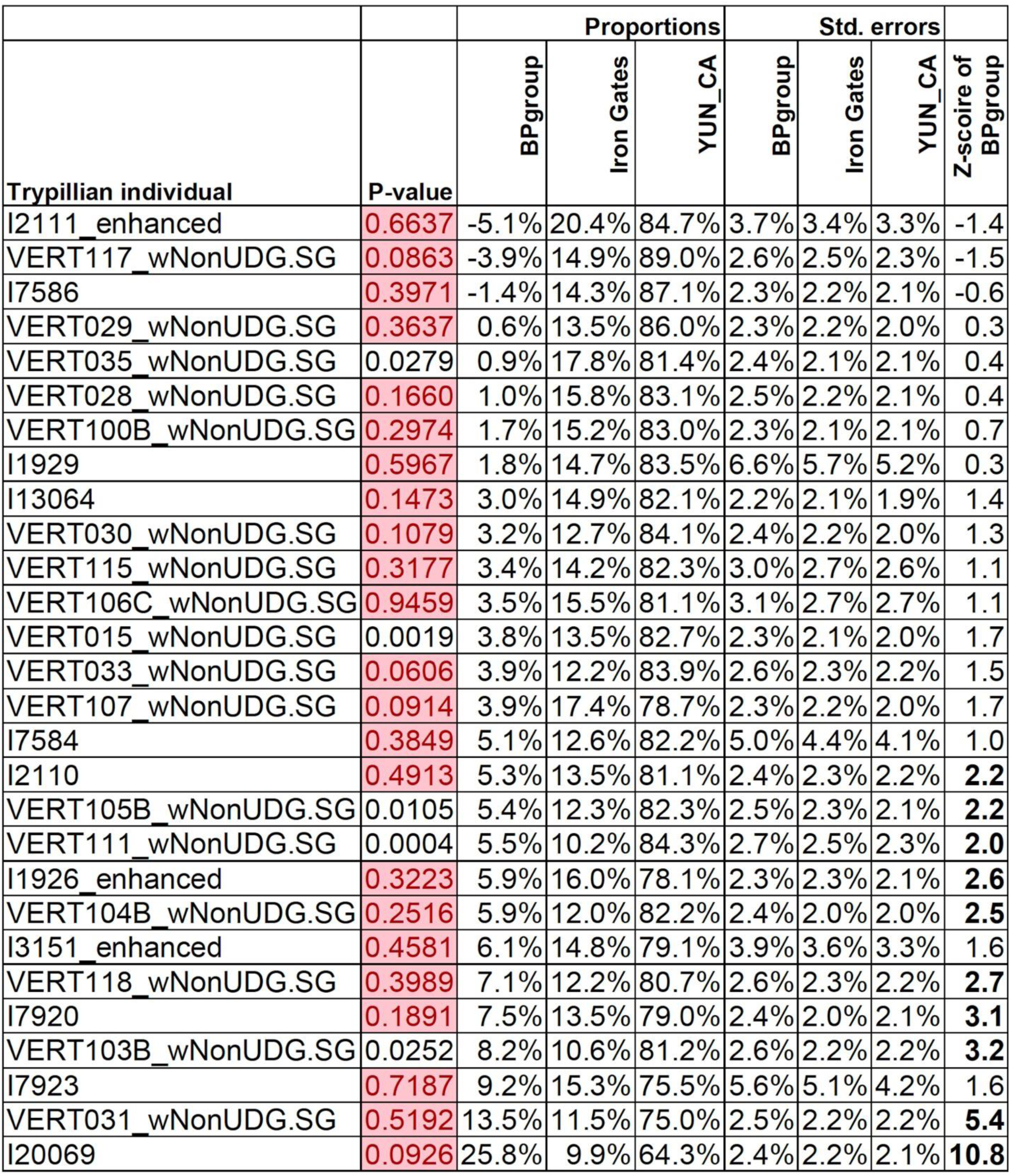
By-individual modeling of Trypillians. This is the same as Table S19 in the supplement.

**Extended Data Figure 1.**
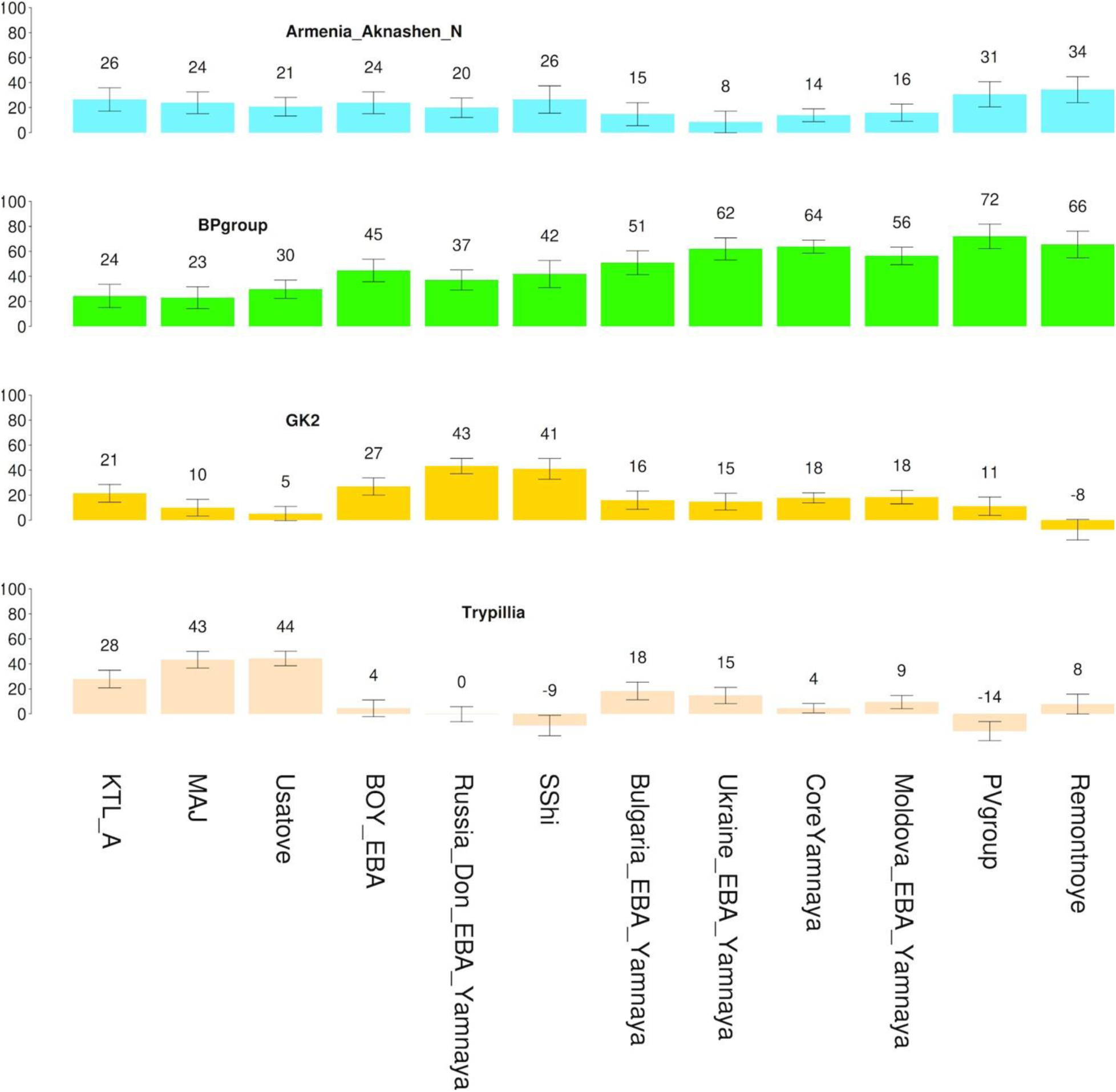
Admixture proportions of 4-source model with Trypillians as the 4th source. Plotted populations fit the model (p>0.05) and have RMSE of standard errors ≤10%. ±1S.E. are shown.

**Extended Data Figure 2.**
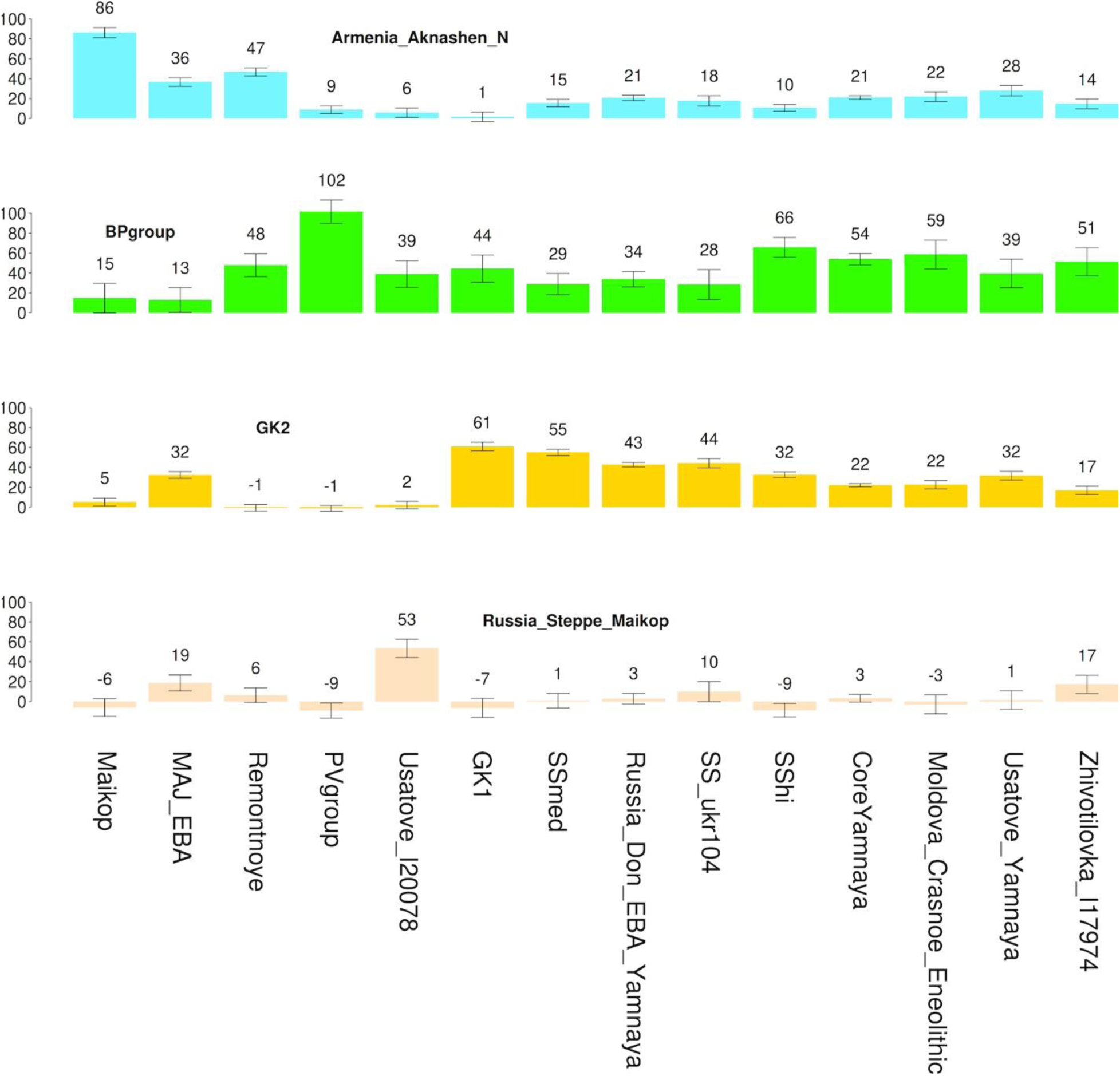
Admixture proportions of 4-source model with Steppe Maykop as the 4th source. Plotted populations fit the model (p>0.05) and have RMSE of standard errors ≤10%. ±1 S.E. are shown.

## References

1. Gimbutas, M. A. Three Waves of the Kurgan People into Old Europe, 4500-2500 B.C. Journal of Indo-European Studies 18, 240–268 (1997).

2. Anthony, D. W. The Horse, the Wheel, and Language: How Bronze-Age Riders from the Eurasian Steppes Shaped the Modern World. (Princeton University Press, Princeton and Oxford, 2007).

3. Haak, W. et al. Massive migration from the steppe was a source for Indo-European languages in Europe. Nature 522, 207–11 (2015).

4. Allentoft, M. E. et al. Population genomics of Bronze Age Eurasia. Nature 522, 167–172 (2015).

5. Penske, S. et al. Early contact between late farming and pastoralist societies in southeastern Europe. Nature 620, 358–365 (2023).

6. Mathieson, I. et al. The genomic history of southeastern Europe. Nature 555, 197–203 (2018).

7. Lazaridis, I., Patterson, N., Anthony, D. & & others. The Genetic Origin of the Indo-Europeans. in Submission. (2024).

8. de Barros Damgaard, P., et al. The first horse herders and the impact of early Bronze Age steppe expansions into Asia. Science (1979) eaar7711 (2018) doi:10.1126/science.aar7711.

9. Narasimhan, V. M. et al. The formation of human populations in South and Central Asia. Science (1979) 365, (2019).

10. Lazaridis, I. et al. The genetic history of the Southern Arc: A bridge between West Asia and Europe. Science (1979) 377, (2022).

11. Kroonen, G., Jakob, A., Palmér, A. I., van Sluis, P. & Wigman, A. Indo-European cereal terminology suggests a Northwest Pontic homeland for the core Indo-European languages. PLoS One 17, e0275744 (2022).

12. Lipson, M. et al. Parallel palaeogenomic transects reveal complex genetic history of early European farmers. Nature 551, 368–372 (2017).

13. Nikitin, A. G., Videiko, M., Patterson, N., Renson, V. & Reich, D. Interactions between Trypillian farmers and North Pontic forager-pastoralists in Eneolithic central Ukraine. PLoS One 18, e0285449 (2023).

14. Burdo, N. B. Kul’turno-istoricheskiye kontakty ranne-tripol’skikh plemen. In Drevneyshiye obshchnosti zemledel’tsev i skotovodov Severnogo Prichernomor’ya (V tys. do n.e. — V vek n.e.) (ed. Yarovoy, E. V.) 49–51 (Nauchno-issledovatel’skaya laboratoriya «Arkheologiya» PGU im. T.G. Shevchenko, Tiraspol’, 2002).

15. Nikitin, A. G. et al. Mitochondrial DNA analysis of Eneolithic Trypillians from Ukraine reveals Neolithic farming genetic roots. PLoS One 12, e0172952 (2017).

16. Nikitin, A. G., Sokhatsky, M. P., Kovaliukh, M. M. & Videiko, M. Y. Comprehensive Site Chronology and Ancient Mitochondrial DNA Analysis from Verteba Cave – a Trypillian Culture Site of Eneolithic Ukraine. Interdisciplinaria Archaeologica. Natural Sciences in Archaeology. 1, 9–18 (2010).

17. Gelabert, P. et al. Genomes from Verteba cave suggest diversity within the Trypillians in Ukraine. Sci Rep 12, 7242 (2022).

18. Mattila, T. M. et al. Genetic continuity, isolation, and gene flow in Stone Age Central and Eastern Europe. Commun Biol 6, 793 (2023).

19. Kotova, N. S. Early Eneolithic in the Pontic Steppes. (British Archaeological Reports, Oxford, UK, 2008).

20. Telegin, D. Y. Neoliticheskiye Mogil’niki Mariupol’skogo Tipa. (Naukova Dumka, Kiev, 1991).

21. Telegin, D. Y. Keramika rannʹoho eneolitu typu Zasukha v lisostepovomu Livoberezhzhi Ukrayiny. Arkheolohiya 64, 73–84 (1988).

22. Nielsen, R. et al. Tracing the peopling of the world through genomics. Nature 541, 302– 310 (2017).

23. Patterson, N., Price, A. L. & Reich, D. Population structure and eigenanalysis. PLoS Genet (2006) doi:10.1371/journal.pgen.0020190.

24. Ecsedy, I. The People of the Pit-Grave Kurgans in Eastern Hungary. (Akadémiai Kiadó, Budapest, 1979).

25. Govedarica, B. Zepterträger, Herrscher Der Steppen: Die Frühen Ockergräber Des Älteren Äneolithikums Im Karpatenbalkanischen Gebiet Und Im Steppenraum Südost-Und Osteuropas. (Verlag Philipp von Zabern, Mainz am Rhein, 2004).

26. Posth, C. et al. Palaeogenomics of Upper Palaeolithic to Neolithic European hunter-gatherers. Nature 615, 117–126 (2023).

27. Haskevych, D. Late Mesolithic Individuals of the Danube Iron Gates Origin on the Dnipro River Rapids (Ukraine)? Archaeological and Bioarchaeological Records. Open Archaeology 8, 1138–1169 (2022).

28. Mathieson, I. et al. Genome-wide patterns of selection in 230 ancient Eurasians. Nature 528, 499–503 (2015).

29. Lazaridis, I. et al. Ancient DNA from Mesopotamia suggests distinct Pre-Pottery and Pottery Neolithic migrations into Anatolia. Science (1979) 377, 982–987 (2022).

30. Skoglund, P. et al. Genomic Diversity and Admixture Differs for Stone-Age Scandinavian Foragers and Farmers. Science (1979) 344, 747–750 (2014).

31. Malmström, H. et al. The genomic ancestry of the Scandinavian Battle Axe Culture people and their relation to the broader Corded Ware horizon. Proceedings of the Royal Society B: Biological Sciences 286, 20191528 (2019).

32. Coutinho, A. et al. The Neolithic Pitted Ware culture foragers were culturally but not genetically influenced by the Battle Axe culture herders. Am J Phys Anthropol 172, 638– 649 (2020).

33. Allentoft, M. E. et al. Population genomics of post-glacial western Eurasia. Nature 625, 301–311 (2024).

34. Patterson, N. et al. Ancient Admixture in Human History. Genetics 192, 1065–1093 (2012).

35. Chintalapati, M., Patterson, N. & Moorjani, P. The spatiotemporal patterns of major human admixture events during the European Holocene. Elife 11, (2022).

36. Wang, C.-C. et al. Ancient human genome-wide data from a 3000-year interval in the Caucasus corresponds with eco-geographic regions. Nat Commun 10, 590 (2019).

37. Järve, M. et al. Shifts in the Genetic Landscape of the Western Eurasian Steppe Associated with the Beginning and End of the Scythian Dominance. Current Biology 29, 2430–2441.e10 (2019).

38. Lazaridis, I. et al. Genetic origins of the Minoans and Mycenaeans. Nature (2017) doi:10.1038/nature23310.

39. Skourtanioti, E. et al. Ancient DNA reveals admixture history and endogamy in the prehistoric Aegean. Nat Ecol Evol (2023) doi:10.1038/s41559-022-01952-3.

40. Clemente, F. et al. The genomic history of the Aegean palatial civilizations. Cell 184, 2565–2586.e21 (2021).

41. Korobkova, G. F. & Shaposhnikova, O. G. Poselenie Mikhailovka: Etalonnyj Pamyatnik Drevneyamnoj Kultury. (Evropejskij Dom, St. Petersburg, 2005).

42. Kotova, N. S. Dereivskaya Kul’tura i Pamyatniki Nizhnemikhaylovskogo Tipa. (Maidan, Kiev, Kharkov, 2013).

43. Ivanova, S., Nikitin, A. G. & Kiosak, D. Pendulum migrations in the Circum-Pontic steppe and Central Europe during the Paleometal Epoch and the problem of genesis of the Yamna culture. Archeologia 26, 101–146 (2018).

44. Nikitin, A. G. & Ivanova, S. Long-distance exchanges along the Black Sea coast in the Eneolithic and the steppe genetic ancestry problem. in Steppe Transmissions (eds. Preda-Bălănică, B. & Ahola, M.) 9–27 (Archaeolingua, Budapest, 2023). doi:10.33774/coe-2022-7m315.

45. Gimbutas, M. The Indo-Europeanization of Europe: the intrusion of steppe pastoralists from south Russia and the transformation of Old Europe. WORD 44, 205–222 (1993).

46. Dabney, J. et al. Complete mitochondrial genome sequence of a Middle Pleistocene cave bear reconstructed from ultrashort DNA fragments. Proceedings of the National Academy of Sciences 110, 15758–15763 (2013).

47. Korlević, P. et al. Reducing microbial and human contamination in DNA extractions from ancient bones and teeth. Biotechniques 59, (2015).

48. Rohland, N., Harney, E., Mallick, S., Nordenfelt, S. & Reich, D. Partial uracil-DNA-glycosylase treatment for screening of ancient DNA. Philosophical Transactions of the Royal Society B: Biological Sciences 370, 20130624–20130624 (2014).

49. Rohland, N., Glocke, I., Aximu-Petri, A. & Meyer, M. Extraction of highly degraded DNA from ancient bones, teeth and sediments for high-throughput sequencing. Nat Protoc 13, 2447–2461 (2018).

50. Prendergast, M. E. et al. Ancient DNA reveals a multistep spread of the first herders into sub-Saharan Africa. Science (1979) 365, (2019).

51. Gansauge, M.-T., Aximu-Petri, A., Nagel, S. & Meyer, M. Manual and automated preparation of single-stranded DNA libraries for the sequencing of DNA from ancient biological remains and other sources of highly degraded DNA. Nat Protoc 15, 2279–2300 (2020).

52. Fu, Q. et al. An early modern human from Romania with a recent Neanderthal ancestor. Nature 524, 216–219 (2015).

53. Behar, D. M. et al. A “Copernican” Reassessment of the Human Mitochondrial DNA Tree from its Root. The American Journal of Human Genetics 90, 675–684 (2012).

54. Li, H. & Durbin, R. Fast and accurate long-read alignment with Burrows-Wheeler transform. Bioinformatics (2010) doi:10.1093/bioinformatics/btp698.

55. Fu, Q. et al. A revised timescale for human evolution based on ancient mitochondrial genomes. Current Biology 23, 553–559 (2013).

56. Korneliussen, T. S., Albrechtsen, A. & Nielsen, R. ANGSD: Analysis of Next Generation Sequencing Data. BMC Bioinformatics 15, 356 (2014).

57. Briggs, A. W. et al. Removal of deaminated cytosines and detection of in vivo methylation in ancient DNA. Nucleic Acids Res 38, e87–e87 (2010).

58. Li, H. et al. The Sequence Alignment/Map format and SAMtools. Bioinformatics 25, 2078–2079 (2009).

59. Weissensteiner, H. et al. HaploGrep 2: mitochondrial haplogroup classification in the era of high-throughput sequencing. Nucleic Acids Res (2016) doi:10.1093/nar/gkw233.

60. Lazaridis, I. et al. Genomic insights into the origin of farming in the ancient Near East. Nature 536, 419–424 (2016).

61. Shinde, V. et al. An Ancient Harappan Genome Lacks Ancestry from Steppe Pastoralists or Iranian Farmers. Cell 179, 729–735.e10 (2019).

62. Harney, É. et al. Ancient DNA from Chalcolithic Israel reveals the role of population mixture in cultural transformation. Nat Commun 9, 3336 (2018).

63. Rivollat, M. et al. Ancient genome-wide DNA from France highlights the complexity of interactions between Mesolithic hunter-gatherers and Neolithic farmers. Sci Adv 6, (2020).

64. Reich, D. et al. Reconstructing Native American population history. Nature 488, 370–374 (2012).

65. Skoglund, P. et al. Reconstructing Prehistoric African Population Structure. Cell 171, 59–71.e21 (2017).

66. Wang, K. et al. Ancient genomes reveal complex patterns of population movement, interaction, and replacement in sub-Saharan Africa. Sci Adv 6, (2020).

67. Lipson, M. et al. Ancient DNA and deep population structure in sub-Saharan African foragers. Nature 603, 290–296 (2022).

68. Fu, Q. et al. The genetic history of Ice Age Europe. Nature 534, 200–205 (2016).

69. Jones, E. R. et al. Upper Palaeolithic genomes reveal deep roots of modern Eurasians. Nat Commun 6, 8912 (2015).

70. Alexander, D. H., Novembre, J. & Lange, K. Fast model-based estimation of ancestry in unrelated individuals. Genome Res 19, 1655–1664 (2009).

71. Fenner, J. N. Cross-cultural estimation of the human generation interval for use in genetics-based population divergence studies. Am J Phys Anthropol 128, 415–423 (2005).

