## Supplementary Information for "A genomic history of the North Pontic Region from the Neolithic to the Bronze Age"

**Table of Contents**

**SI 1** – Archaeological Background and Burial Descriptions 1-103

**SI 2** – Population genetic analysis of populations from the North Pontic and adjacent regions 104-151

**SI 3** – Unsupervised admixture modeling with summary individuals 152-166

### A genomic history of the North Pontic Region from the Neolithic to the Bronze Age

*Supplementary Information, section 1*

#### Archaeological Background and Burial Descriptions

#### Table S1. Data summary report for specimens used in the analysis.

| № | Genetic ID | Specimen code | Site, Burial | $^{14}$C uncalBP (Lab Code) | BCE or Calibrated BCE, 2σ (95.4%)[a] | Genetic sex | MtDNA haplogroup | Y chromosome haplogroup |
|---|---|---|---|---|---|---|---|---|
| | **Mariupol type cemeteries, Ukraine (Neolithic)** | | | | | | | |
| 1 | I31730 | MM.50a | Mariupol, Burial 50 (adult)[1] | 6395±35 (PSUAMS-10373) | 5474-5236 calBCE | M | U5a1+@16192 | I2a1b1a2a2a |
| 2 | I31731 | MM.50b | Mariupol, Burial 50 (immature) | 3720±30 (BETA-681064)[b] | 2204-1986 calBCE[c] | F(?) | U5b2 (<2X) [d] | |
| 3 | I27982 | Yas23a/Ya42 | Yasynuvatka, Burial 23a, Pit Б-1 [2,3] | n/a | 5560-4750 BCE | M | U4a1 | Q1b |
| 4 | I27983 | Yas24/Ya40 | Yasynuvatka, Burial 24, Pit Б-1 [2,3] | n/a | 5560-4750 BCE | M | H | R1b1a1b1a1a2c1a3a2 |
| 5 | I27986 | Ya8 | Yasynuvatka, Burial 35, Pit Б-1 [2–5] | 6320±30 BP, BETA-441244/445800)[5]; 5810±60 (Ki-3162)[4] | 5363-5216 calBCE; 4797-4502 calBCE | M | U5a2 | P1 |
| 6 | I27990 | Yas44 | Yasynuvatka, Burial 44, Pit Б-1[2,3] | n/a | 5560-4750 BCE | M | U5a1 | Q |
| 7 | I27992[e] | Yas50 | Yasynuvatka, Burial 50, Pit Б-1[2,3] | n/a | 5560-4750 BCE | F | K1a3 | |
| 8 | I27994 | Yas55 | Yasynuvatka, Burial 55, Pit A2[2,3] | 6650±30 (BETA-445802)[5] | 5630-5484 calBCE | M | T2a1b | I2a1b2 (I-S2555) |
| 9 | I27995 | Yas58 | Yasynuvatka, Burial 58, Pit A2 [2,3] | n/a | 5650-5450 BCE | M | U4b1b1 | n/a |
| | **Cucuteni-Trypillia, Moldova (Eneolithic)** | | | | | | | |
| 10 | I7920 | 12M1 | Cunicea-Prişanscaia Gora (Soldaneşti), Section 4, Layer 3, burial 4 | 4505±25 (PSUAMS-4872) | 3350-3099 calBCE | M | K1b2 | I2a1b1a2a2a2 (I-S12195) |
| 11 | I20069[g] | DII.301 | Dănceni II, Ground Burial 2 (301) | 4435±20 (PSUAMS-8832) | 3323-2935 calBCE | M | H | I2a1b1a2a2a (I-L699) |
| | **Cucuteni-Trypillia, Ukraine (Eneolithic)** | | | | | | | |
| 12 | I1926 [h] | 1V1a-H1 | Verteba Cave, Site 7[6], individual 3.17.1[7–9] | 4906±23 (weighted mean[10] of 4890±30 (Beta-432808[8]) and 4925±33 (OxA-26207[7]) | 3756-3638 calBCE | M | H5a | G2a2b2a (G-P303) |
| 13 | I2109 | 3V3a-H3 | Verteba Cave, Site 7[6], Individual V1-2SE (3.18.1) | 4863±33 (OxA-26202)[7] | 3710-3531 calBCE | M | J1c | G2a (G-L1259) |
| 14 | I2110 [h] | 4V4a-H4 | Verteba Cave, Site 7[6], Individual 3.14.1[7–9] | 4976±33 (OxA-26203)[7] | 3911-3650 calBCE | M | T2b | G2a2b2a (G-P303) |

| | | | | | | | | |
|---|---|---|---|---|---|---|---|---|
| 15 | I2111 [h] | 5V5a-H5 | Verteba Cave, Site 7[6], Individual 3.16.1[7–9] | 4888±32 (OxA-26204)[7] | 3765-3543 calBCE | M | H+16311 | G2a (G-P15) |
| 16 | I3151 [h] | 6V6a-H6 | Verteba Cave, Site 7, individual 3.15.1[7–9] | 4807±33 (OxA-26201)[7] | 3644-3527 calBCE | M | U8b1b | E1b1b1a1b1 (E-L618) |
| 17 | I1929 | 9 V9a-A22.2 | Verteba Cave Site 6, Individual 2 | 4910±400 (Ki-14308)[6] | 3900-3500 calBCE | F | H | |
| 18 | I7921 | A22.P3 | Verteba Cave, Site 6[6], Individual 3[6,7] | n/a | 3900-3500 BCE | F | H2a | |
| 19 | I7923 | RL2.05 (VRT-LR1) | Verteba Cave, Site 2[6], Individual 1 | 4955±30 (PSUAMS-8692) | 3789-3649 calBCE | M | T2b+16362 | J2a1 |
| 20 | I7584 | VCS1.CL.05 | Verteba Cave Site 1[6], Individual 1 | 4875±25 (PSUAMS-10769) | 3708-3543 calBCE | F | H5b | |
| 21 | I7586 | VCS1.TBL.05 | Verteba Cave, Site 5[6], Individual 2 | 4920±25 (PSUAMS-9527) | 3766-3642 calBCE | F | J1c2 | |
| 22 | I13068 | VRT1.1.1 | Verteba Cave, Site 7[6], cranium 1 (west) | n/a | 3950-3500 BCE | M | T2 | n/a |
| 23 | I13064 | VRT-VSE.22N, VRT-VNWL3 | Verteba Cave, Site 7[6], Individual 2 | 4820±30 (PSUAMS-7807) | 3648-3528 calBCE | M | T2 | I2a1b1a2b (I-Z161) |
| | **Usatove, Ukraine (Eneolithic-Early Bronze Age)** | | | | | | | |
| 24 | I12615 | Ma6 | Mayaky Sanctuary, Burial 1 | 6100±30 (BETA-432410)[5] | 5208-4907 calBCE | F | U4b1b2 | |
| 25 | I12228 | Ma13 | Mayaky Sanctuary, Burial 2 | 5580±30 (BETA-432406)[5] | 4486-4350 calBCE | F | U4[f] | |
| 26 | I12229 | Mau1 | Mayaky, Kurgan 1, Burial 9[11] | 4375±25 (PSUAMS-7865) | 3088-2911 calBCE | F | U5a1a1 | |
| 27 | I12704 | Mau6 | Mayaky, Kurgan 7, Burial 2[11] | 5144±20 (weighted mean[10] of 5530±32 (OxA-22959)[12], 5390±30 (BETA-441235)[5], and 5545±40 (PSUAMS-7793, this report) | 3620-3030 calBCE[g] | M | T2h2 | E1b1b1a1 |
| 28 | I12710 | Mau12 | Mayaky Kurgan 8, Burial 4[11] | 5295±30 (PSUAMS-7862) | 3769-3196 calBCE[g] | F | W1 | |
| 29 | I12707 | Mau9 | Mayaky Kurgan 8, Burial 6, Skeleton 2[11] | 5330±40 (PSUAMS-7794) | 3511-2917 calBCE[g] | F | U5a1d1 | |
| 30 | I12706 | Mau8 | Mayaky Kurgan 9, Burial 2[11] | 5444±19 (weighted mean[10] of 5471±24, 0xA-22960[12], and 5400±30, PSUAMS-7846) | 3947-3519 calBCE[g] | F | X2d | |
| 31 | n/a | Ma12 | Mayaky, Kurgan 10, Burial 2, Skeleton 1[11] | 5530±30 (BETA-432413)[5] | 4033-3534 calBCE[g] | F | U4[b] | n/a |
| 32 | I1423 (I12705) | Mau7/MAJ1 | Mayaky, Kurgan 10, Burial 2, Skeleton 2[11] | 5390±30 (PSUAMS-7845) | 3936-3372 calBCE[g] | M | H+195 | R1a |
| 33 | I7929 | REV3.19 | Revova, Kurgan 3, Burial 19[13] | 4905±20 (PSUAMS-4763) | 3711-3639 calBCE | M | U4d3 | J1 (J-FT265222) |

| | | | | | | | | |
|---|---|---|---|---|---|---|---|---|
| | **Serednii Stih, Ukraine (Eneolithic)** | | | | | | | |
| 34 | I27282 | Der15 | Deriivka II, Skeleton 8[14] | 5345±30 (PSUAMS-10370) | 4322-4042 calBCE | M | U4b1a | I2a1b1a2 |
| 35 | I28319 | Der16 | Deriivka II, Skeleton 5[14] | 5344±29 (weighted mean of 5380±90 (OxA-5032)[15] and 5340±30 (BETA-445808)[5] | 4321-4052 calBCE | F | T2a1b | |
| 36 | I27283 | Der41 | Deriivka II, Skeleton 11[14] | 5500±30 (PSUAMS-10371) | 4444-4262 calBCE | M | U4c | I2a1b1a2a2a |
| 37 | I2108 | Igren-8 9 | Ogrin-8, Burial 20 | 5415±25 (PSUAMS-8220) | 4340-4178 calBCE | F | U2e | |
| 38 | I1924 | Igren 8-78 | Ogrin-8, Burial 24 | 5105±25 (PSUAMS-8219) | 3971-3802 calBCE | F | U5b2 | |
| 39 | I27930 | Igren 8-74 | Ogrin-8, Burial 3a | n/a | 4400-4000 BCE | M | U4 | R1b1b (R-V88) |
| 40 | I12167 (I31733) | K1.10 | Katarzhyno, Kurgan 1, Burial 10[13] | 5039±27 (weighted mean of 4970±110 (Ki-11376), 4950±70 (Ki-9523)[13], and 5060±30 (PSUAMS-10375) | 3951-3715 calBCE | F | U5a2a2 (<2X)[d] | |
| 41 | I7585[h] | KST07 | Kolomiytsiv Yar Tract, Burial 1[16] | 5170±30 (BETA-523816) | 4049-3820 calBCE | F | U4b1b2 | |
| 42 | I31732 | MM.21 | Mariupol Necropolis, Burial 21[1] | 5790±30 (PSUAMS-10374) | 4715-4548 calBCE | F | n/a | |
| 43 | I1424 | MOB1 | Molyukhiv Bugor, Burial 4, Skeleton 3[15,17] | 4943±28 (weighted mean[10] of 5020±50 (OxA-17502), and 4909±33 (OxA-17503)[15]) | 3777-3648 calBCE | M | U5a1[17] | I2a1b1a2a2a (I-L699) |
| 44 | I1454 | MOB3 | Molyukhiv Bugor, Burial 4, Skeleton 2[17] | n/a | 4000-3600 BCE | F | U/K[17] | |
| 45 | I1430 | VIN1 | Vynogradne, Kurgan 3, Burial 15 [17–20] | 5230±60 (Ki-14726)[18] | 4241-3951 calBCE | M | H13 | I2a1b1a2a2a (I-L699) |
| | **Steppe Eneolithic/Kvityana/Post-Mariupol, Moldova** | | | | | | | |
| 46 | I20196 | Kr/K9/B9/S2 | Crasnoe, Kurgan 9, Burial 9, Skeleton 2[21] | 4510±25 BP, PSUAMS-8835 | 3352-3101 calBCE | F | J1b1a1 | |
| 47 | I17977 | Kr/K9/B9/S2(d) | Crasnoe, Kurgan 9, Burial 9, Skeleton 3 (child)[21] | n/a | 3352-3101 calBCE (based on I20196 – 1st degree kin) | F | J1b1a (<2X)[d] | |
| | **Kvityana/Post Mariupol, Ukraine (Eneolithic)** | | | | | | | |
| 48 | I12618 | V4.4 | Vapnyarka, Kurgan 4, Burial 4[22] | 4480±30 (BETA-647661); 4100±80 (Ki-15013)[22] | 3341-3031 calBCE; 2878-2476 calBCE | F(?) | U8b1a2a (<2X)[d] | |
| | **Zhyvotylivka, Moldova (Eneolithic)** | | | | | | | |
| 49 | I17973 | Br/K1/B21/S1 | Bursuceni, Kurgan 1, Burial 21, Skeleton 1[23] | 4516±13 (weighted mean[10] of 4530±20 (PSUAMS-8723), 4520±25 (PSUAMS-8310) and 4500±20 (PSUAMS-8722) | 3354-3103 calBCE | M | U1a1a | J2b2b2~ (J-Z42942) |

| | | | | | | | | |
|---|---|---|---|---|---|---|---|---|
| 50 | I17974 | Br/K1/B21/S2 | Bursuceni, Kurgan 1 Burial 21, Skeleton 2[23] | 4470±20 (PSUAMS-8724) | 3334-3030 calBCE | F | U5a1g1 | |
| 51 | I17975 (I20195) | Br/K1/B21/S3 | Bursuceni, Kurgan 1 Burial 21, Skeleton 3[23] | n/a | 3334-3030 calBCE (based on I17974 twin) | F | U5a (<2x)[d] | |
| 52 | I20078 | TII.K2.14 | Taraclia II, Kurgan 2, Burial 14 | 4480±25 (PSUAMS-8833) | 3340-3035 calBCE | M | H2a1 | Q1b2b1b2b~ (Q-BZ1466) |
| 53 | I20079 | TII.K10.2 | Taraclia II, Kurgan 10, Burial 2 | 3965±20 (PSUAMS-8834) | 2566-2347 calBCE | M | K1b2b | R1b1a1b (R-M269) |
| | **Suvorove, Moldova (Eneolithic)** | | | | | | | |
| 54 | I20072 | GK2.6 | Giurgiulești, Burial 6 (3)[24] | 5370±26 (MAMS-23175)[24] | 4330-4058 calBCE | M | H13 | Q1a2 |
| 55 | I20073 | GK2.11 | Giurgiulești, Burial 11 (4)[24] | 5571±32 (MAMS-28088)[24] | 4348-4248 calBCE | M | K1b2b (<2X) | CF |
| | **Proto-Yamna, Bulgaria (Eneolithic)** | | | | | | | |
| 56 | I1456 | DUR1 | Durankulak, Kurgan F, burial 15 (main burial)[25] | n/a | 3500-3000 BCE | M | U5a2a | R1a1a1 (R-M417) |
| | **Yamna, Ukraine (Eneolithic-Early Bronze Age)** | | | | | | | |
| 57 | I32534 | M1P1.6 | Mykhailivka 1, square VI | 4755±25 (PSUAMS-10750) | 3635-3383 calBCE | F | H6a1 | |
| 58 | I12170 | K2.1 | Katarzhyno, Kurgan 2, Burial 1[13] | 4490±30 (BETA-647662); 4270±90 (Ki-17331)[13] | 3348-3038 calBCE; 3314-2537 calBCE | M(?) | U4c1 | |
| 59 | I20975 | KM17P2 | Kamyana Mohyla, Kurgan 2, Burial 2[26] | 4141±21 (BE-8040.1.1)[26] | 2872-2626 calBCE | M | N1b1a2 | R1b1a1b1b (R-M12149) |
| 60 | I12168 | KTR-1.13 | Katarzhyno, Kurgan 1, Burial 13[13] | 4359±16 (weighted mean of 4375±20 (PSUAMS-4762) and 4335±25 (PSUAMS-9815) | 3018-2911 calBCE | M | U4a | R1b1a1b1b3 (R-Z2108) |
| 61 | I11999 | KTR-1.1 | Katarzhyno, Kurgan 1, Burial 1[13] | 4045±20 (PSUAMS-4903) | 2626-2476 calBCE | M | U4a2 | R1b1a1b1b3 (R-Z2108) |
| 62 | I12233 | L2.16 | Liubasha, Kurgan 2, Burial 16[13] | 4330±35 (PSUAMS-7790) | 3074-2888 calBCE | M | U4a1 | R1b1a1 (R-P297) |
| | I7927 | R3.7 | Revova Kurgan 3, Burial 7[13] | 3910±60 (Ki-11058) | 2568-2206 calBCE | n/a | U5a (>2X)[d] | n/a |
| 63 | I7928 | R3.16 | Revova Kurgan 3, Burial 16[13] | 4135±60 (Ki-11059)[13] | 2885-2501 calBCE | M (?) | U+152, 16311 (>2X)[27,d] | n/a |
| 64 | I12005 | SL-10 | Soldats'ka Slava Kurgan, Burial 10[13] | n/a | 3300-2500 BCE | M | T2c1a2 | n/a |
| 65 | I12843 | S15 | Sychavka Kurgan, Burial 15[28] | 4087±19 (weighted mean of 3960±80 (Ki-16610)[28], 4070±30 (BETA-441246)[5], and 4110±25 (PSUAMS-9556) | 2847-2501 calBCE | M | U5a1a1 | R1b1a1b1b (R-M12149) |

| | | | | | | | | |
|---|---|---|---|---|---|---|---|---|
| 66 | I12619 | V4.16 | Vapnyarka, Kurgan 4, Burial 16[22] | 4155±27 (weighted mean of 4050±60 (Ki-15014)[22], and 4180±30 (BETA- 441245)[5] | 2876-2631 calBCE | M | W3a1a | R1b1a1b1b3 (R-Z2108) |
| 67 | I3141 [h] | Yamna 5 | Shevchenko (Shakhtar), Kurgan 28, Burial 12[8] | n/a | 3300-2500 BCE | F | H | |
| | **Yamna, Bulgaria (Early Bronze Age)** | | | | | | | |
| 68 | I1448 | POP1 | Popovo, Golyamata Mogila, Burial 29[29] | n/a | 3300-2500 BCE | F | H6a1b | |
| 69 | I1427 | POP4 | Popovo, Golyamata Mogila, Burial 37[29] | n/a | 3300-2500 BCE | M | U5a1a1 | R1b1a1b1b3 (R-Z2108) |
| 70 | I1428 | RIL3 | Riltsi, Riltsi Kurgan 264, Burial 5[30] | n/a | 3360-2890 BCE | F | K | |
| | **Yamna, Moldova (Eneolithic, Early Bronze Age)** | | | | | | | |
| 71 | I20067 | CiK4.3 | Cioburciu, Kurgan 4, Burial 3 | n/a | 3300-2500 BCE | F | I4 | |
| 72 | I20068 | CoK1.6 | Cotiujeni, Kurgan 1 Burial 6 | 4175±20 (PSUAMS-10740) | 2881-2671 calBCE | M | H13a2b2a | R1b1a1b1b (R-M12149) |
| 73 | I12507 | Glinoe DOT K1B9 | Glinoe (Hlinaia), DOT Kurgan 1, Burial 9 | 4275±20 (PSUAMS-10777) | 2914-2882 calBCE | F | U5a1a1 | |
| 74 | I12512 | Glinoe SAD K1B25 | Glinoe (Hlinaia), SAD Kurgan 1, Burial 25 | 4165±25 (PSUAMS-10778) | 2881-2633 calBCE | M | H2a1 | R1 |
| 75 | I12637 | Glinoe SAD K1B26 | Glinoe (Hlinaia), SAD Kurgan 1, Burial 26 | n/a | 3300-2500 BCE | M | U2e3 | R1b1a1b1b (R-M12149) |
| 76 | I17743 | MeII1.15 | Mereni II, Kurgan 1, Burial 15 | 4515±30 (PSUAMS-7822) | 3358-3100 calBCE | F | H+16311 | |
| 77 | I17742 | MK3.15 | Mărculeşti, Kurgan 3, burial 15 | n/a | 3300-2500 BCE | F | n/a | |
| 78 | I20076 | OcK1.3 | Ocniţa, Kurgan 1, Burial 3 | 4230±25 (PSUAMS-11947) | 2906-2702 calBCE | F | U5a2a1 | |
| 79 | I17744 | SK1.1 | Sărăteni, Kurgan 1, Burial 1 | 4115±20 (PSUAMS-8442) | 2860-2578 calBCE | M | H6a1a | R1b1a1b1b3 (R-Z2108) |
| 80 | I17745 | SK1.9 | Sărăteni, Kurgan 1, Burial 9 | 4105±25 (PSUAMS-8309) | 2862-2573 calBCE | M | H13a2b2a | R1b1a1b (R-M269) |
| 81 | I17747 | TiK3.15 | Tiraspol, Kurgan 3, Burial 15 | 4115±25 (PSUAMS-7864) | 2865-2576 calBCE | M | U4d3 | R1b1a1b1b (R-M12149) |
| | **Catacomb, Ukraine (Early Bronze Age)** | | | | | | | |
| 82 | I13071 | BLV1 | Bil'shivtsi, Individual 1[31] | 3741±20, weighted mean[10] of 3770±30 (BETA-446187) and 3720±2 (PSUAMS-7848) | 2271-2039 calBCE | F | T2c1d1a | |
| 83 | I12621 | DBN1.9 | Dubynove, Kurgan 1, Burial 9[13] | 3970±35 (PSUAMS-7791) | 2575-2348 calBCE | M | U5b | n/a |

| | | | | | | | | |
|---|---|---|---|---|---|---|---|---|
| 84 | I12840 | DBN1.10 | Dubynove, Kurgan 1, Burial 10[13] | 3830±30 (PSUAMS-7805) | 2453-2148 calBCE | F | H2b | |
| 85 | I11850 | ODK1 | Odesa Kurgan, Burial 10 | 4150±25 (PSUAMS-11224) | 2874-2630 calBCE | F | H+16311 | |
| 86 | I16668 | REV3.10 | Revova, Kurgan 3, Burial 10[13] | n/a | 2800-2000 BCE | M | U3a (<2X) [d] | CF |
| 87 | I13072 | SL-1 | Soldats'ka Slava kurgan Burial 1[13] | n/a | 2800-2000 BCE | M | T1a1 (<2X) [d] | n/a |
| 88 | I12617 | V4.3 | Vapnyarka, Kurgan 4 Burial 3[22] | 4089±24 (weighted mean[10] of 3960±70 (Ki-15230)[22], and 4105±25 BP (PSUAMS-10808) | 2851-2500 calBCE | F | R1a1a1 | |
| | **Multi-Cordoned Ware/ Babyne, Ukraine (Bronze Age)** | | | | | | | |
| 89 | I12234 | L3 | Liubasha, Kurgan 2, Burial 3[13] | 3080±70 (Ki-11176)[13] | 1499-1127 calBCE | M | R1a (<2X) [d] | R1a1a (R-M198) |
| 90 | I16674 | L2.15 | Liubasha kurgan 2 Burial 15[13] | 3740±70 (Ki-11201)[13] | 2434-1943 calBCE | M | H15a1a1 | R1b1a1b1b (R-M12149) |

[a] Radiocarbon dates were calibrated using the IntCal20 atmospheric curve[32], unless otherwise indicated.
[b] Radiocarbon date from enamel carbonate.
[c] Calibration: BetaCal4.20; HPD method: INTCAL20.
[d] Mitochondrial coverage below 2X is considered unreliable by the standards used by the Reich's lab.
[e] The specimen has a high contamination rate mtcontam=[0.717,0.942]. It is not cladal with Ukrainian Neolithic hunter-gatherers, which is a further consequence of contamination.
[f] Mitochondrial DNA haplogroup assignment by low-resolution PCR at GVSU, followed by Sanger sequencing.
[g] Calibrated using reservoir offset of 545±70 BP.
[h] This study reports enhanced coverage.

#### 1. Archaeological background (with archaeogenetic notes)

In this section, we will use a tiered structure for the designation and taxonomic grouping of archaeological type units, namely archaeological circle/horizon/complex – archaeological group – regional subgroup – local archaeological type. An archaeological circle (horizon, or complex) is composed of archaeological groups, connected by isomorphism of the structure, typological series, and similarity of features to such an extent that it allows us to assume kinship among these groups through common origin or the presence of common elements, which are distinctly different from similar taxonomic units across the geographic and chronological scale; or a group of related archaeological groups characterized by the same typological types. An archaeological group is defined by a recurrent assemblage of similar artifacts and features found within a defined geographic area over a particular time period, and which can be further subdivided into regional subgroups. A local archaeological type is defined by a regionally restricted set of artifacts and features that are variably represented within an archaeological group/subgroup.

##### 1.1. Ponto-Caspian Steppe from the Early Holocene to the Neolithic period (9700-5000 BCE)

The Eurasian steppe is an extensive temperate grassland, stretching across 8000 km from Hungary to Manchuria, covering nearly one-fifth of the Earth's circumference. The Ural Mountains divide this vast expanse into two distinct ecosystems: the Pontic-Caspian steppe to the west, and the East Eurasian steppe to the east. The Pontic-Caspian steppe encompasses coastal Bulgaria and Romania, southern Moldova, and Ukraine, the Kuma-Manych depression in the Northern Caucasus foreland, and the lower Volga region. In the North Pontic region (NPR), a transitional forest-steppe vegetation zone exists between the southern steppe and the forest north of the Black Sea. Climatic features, vegetation types, and faunal compositions vary in the NPR based on proximity to major waterways such as the Dnipro River, which bisects the region from northwest to southeast, and the southern border formed by the Black Sea.

Historically, the Eurasian steppe has played a crucial role as a major thoroughfare for travel and trade. Its expansive flat terrain provides an optimal route connecting Asia, the North Caucasus, and Europe.

During the last Ice Age, the southern part of the NPR was a peri-glacial tundra steppe. It likely served as a glacial refugial zone for human populations of central and eastern Europe during the Last Glacial Maximum (LGM)[33]. These migrations brought the carriers of Western and Eastern Hunter Gatherer (WHG and EHG) ancestry closer together. These groups remained genetically separated until the inflow of WHG ancestry in the gene pool of hunter-gatherers of the Dnipro Valley in the early Neolithic[8]. During the LGM, humans settled along the river valleys such as the Danube, Dnipro, and Dniester rivers, which formed a shared alluvial plain since the late Pleistocene[34], and their river mouths extended ca. 100 km into the now submerged coastline of the brackish Neoeuxinian Lake that existed in the place of the Black Sea at the time.

Following the LGM, the melting of glaciers resulted in the rise of the Caspian Sea levels, causing the Khvalynian transgression. The Caspian Sea subsequently spilled into the

Neoeuxinian Lake via the Manych depression, inundating the coastal shelf, raising the water levels of the lake by 20 meters during 10000-9000 BCE[35] and drowning the coastal glacial refugia such as the Black/Azov Sea Lowland[36]. The prevalence of grasses and broad-leaved trees in the area during the Early Holocene indicates the warming of the climate compared to the LGM, when the existing forest was dominated by pine suggesting of cool and dry conditions at that time[37].

In the Early Holocene, human habitation in the NPR continued to concentrate along the river valleys. The Neoeuxinian Lake remained disconnected from the World Ocean until ca. 7600 BCE. The two-way connection between the Black Sea and the Sea of Marmara became re-established by the onset of the Holocene Climatic Optimum ca. 7000 BCE[37,38]. The presence of a deciduous forest in the northeast NPR During the 6th – late 5th millennium (ca. 6000-4200) BCE[37] corroborates the establishment of an overall warming trend and humid conditions[37] during the late part of the Stone Age.

The mid-7th – 6th millennium BCE that outlines the Neolithic period of the NPR chronology is primarily defined by a coexistence, with various levels of interaction, between the Neolithic agricultural communities of the Balkans and fisher-hunter-foragers living along the tributaries to the Black Sea. The agricultural revolution in the NPR begins in its western part with the establishment of Karanovo-Starčevo-Körös-Criş farming communities in the late 7th - early 6th millennium BCE, thus coinciding with the earliest stages of the Neolithization of Europe. The LBK archaeological complex, which formed in central Europe in the early 6th millennium, reached the Prut-Dniester interfluve in the northwestern forest steppe area of the NPR shortly thereafter. Over 300 LBK sites are now known in the area east of the Carpathians. The site of Kamyane-Zavallia in south-central Ukraine (48.579, 30.059) is the easternmost LBK site in the NPR region excavated to date[39]. A detailed review of the presence of LBK in west-northwest NPR can be found in[40].

The central and eastern NPR in the Neolithic were settled by populations that followed Mesolithic subsistence practices and who had limited interactions with the farming world of the Balkans. The archaeological space of these groups is generally defined by extended supine cemeteries of Mariupol type (M-t cemeteries) named after the eponymic Neolithic Necropolis at Mariupol[1]. Ukrainian archaeology scholars have different views on archaeological group divisions within the populations that left M-t cemeteries. Genetically, the M-t Neolithic groups studied to date display homogeneity irrespective of cultural subdivisions[5,8]. The Neolithic of the North Pontic is further detailed in[41].

We also note that two published Mesolithic samples from the Deriivka 1 archaeological site, I5876 (7040-6703 calBCE) and I5885 (6392-5927 calBCE) cluster with the Neolithic Ukraine samples on the PCA in this report (Figure 2a). Their radiocarbon dates also postdate the other four Mesolithic Ukraine samples published to date. If we consider the Ukrainian hunter-gather ancestry transformation to be associated with the processes that led to the Neolithic transition, these genetic and radiocarbon dating results, taken together, suggest considering the two Deriivka samples as Neolithic, thus extending the Mesolithic-Neolithic boundary in the Dnipro Valley to the first half of the 7th millennium BCE.

#### 1.2. The Eneolithic period (5000-3300 BCE)

The examination of archaeological materials of the Eneolithic North Pontic region suggests that the formation of the steppe archaeological groups in the NPR in the 5th - 4th millennium BCE was affected by the dynamics of the economic and cultural development of neighboring agricultural societies, represented by the Cucuteni-Trypillia archaeological complex (CTAC) in the North Pontic forest-steppe area, the Darkveti-Meshoko archaeological complex, and, later, the Maykop-Novosvobodnaya complex of the North Caucasus. Archaeological sources show that the general stability of the development of these agricultural societies, primarily the Cucuteni-Trypillia, also ensured the stability of the development of their steppe neighbors.

Interactions between the steppe populations and farmers of the Gumelniţa-Karanovo, Cucuteni-Trypillia, and Darkveti-Meshoko complexes become evident since the beginning of the 5th millennium BCE. These integrations lasted through the end of the early Eneolithic, ca. 4200 BCE. It has been suggested that the switch from the collective extended supine burials of the Neolithic to a new steppe burial tradition of the individual contracted on-the-back burials (Chapli-type burials, Kryvyi Rih, Suvorove, etc.) is the reaction of a stratum of steppe populations to first direct contacts with agricultural societies, reflecting significant changes in the economic, social, and spiritual spheres of life of the steppe inhabitants[42].

In the early Eneolithic (ca. 5000-4200 BCE), decorative elements associated with the Serednii Stih Archaeological Complex (SSAC, the Skelya phase), collectively known as Cucuteni C ceramics, are commonly found on Trypillian ceramics beginning with Trypillia A ([43] and references therein). In connection with the Cucuteni C ceramics, assumptions were made about the influx of the SSAC populations into Trypillia. Some scholars linked the spread of the steppe-derived type of ceramics in Trypillia with “steppe wives”, who made these ceramics according to the traditions of their tribes [16,43].

The early phase of SSAC included archaeological sites such as Oleksandriia (third layer), Serednii Stih II, Stril’cha Skelya, as well as burial complexes of the Novodanylivka type [44,45]. Novodanylivka burials contain copper items of Balkan provenance[45], indicating contacts between Novodanylivka and the west Pontic, which can be traced to the operation of the circum-Pontic trade network ca. 4700-4200 BCE[16,46]. In the 4th millennium BCE, early SSAC was succeeded by the Katarzhyno-type post-Stih communities in the steppe, which continued the SSAC burial tradition of flexed on the back inhumations, as well as Post-Mariupol/Kvytyana, with extended supine burials accompanied by ochre and funerary feast pottery, and other steppe Eneolithic groups, some without clear cultural designation[13,47]. In the forest-steppe region of the Dnipro Valley, the SSAC-derived cultural circle of the late 5th – first third of the 4th millennium BCE is represented by Deriivka II-type communities such as Deriivka II, Ogrin (Igren)-8 and Molyukhiv Bugor (the Eneolithic layer). Interactions between SSAC and CTAC intensified in the 4th millennium. Deriivka II-type groups engaged in reciprocal economic and cultural exchanges with CTAC [16].

Prior to this report, only three individuals of the SSAC horizon, from the Serednii Stih cemetery at Oleksandriia in east Ukraine, the Deriivka II cemetery of the Middle Dnipro Valley, and the CTAC settlement at the Kolomiytsiv Yar Tract (KYT) in central Ukraine have undergone an archaeogenetic evaluation. The Oleksandriia individual (I6561) carried early European farmer (EEF) ancestry, potentially from interactions with farming groups such as CTAC [8]. Two-thirds of the genetic ancestry of the Deriivka II individual was inferred to be derived from EHG and Caucasus Hunter-Gatherer (CHG)/Iranian Neolithic (a genetic mix broadly defining the "steppe ancestry"), and one-third was from the Mesolithic and Neolithic populations of the Dnipro Valley [5]. The KYT individual's genetic ancestry was primarily EHG and CHG/Iranian Neolithic-related, with some admixture of WHG ancestry best proxied by Iron Gates Mesolithic[5], potentially mediated through Neolithic populations of the Dnipro Valley [16]. Lastly, the genetic ancestry of an individual from the Decea Mureșului cemetery in Transylvania, Romania, displaying features of elite steppe "Ochre Grave" burials of the Novodanylivka type, was determined to be EEF-derived [5].

The middle Eneolithic in the steppe (ca. 3900-3500 BCE) follows the so-called hiatus period (ca. 4200-3900 BCE)[44] with less clearly defined chronological sequence of steppe monuments and likely reflecting regional transformations precipitated by atmospheric cooling and fluctuations of the Black Sea level[46,48]. The climatic conditions in the steppe are thought to have returned to more optimal for human habitation after 3900 BCE, which facilitated subsequent cultural transformations[49]. The Middle Eneolithic corresponds to the time of the height of Trypillian development (the BII-CI and CI period of Trypillian chronology) and the formation of a new system of stable archaeological groups in the steppe and, partly, forest-steppe zones, in particular, the Lower Mykhailivka, Deriivka, Kvityana and Konstantinovka, which are characterized by different forms of complex economy. It was at this time that the first complex burial structures appeared in the steppe. These include cromlechs, stone rings, sanctuaries with pillar structures, ring ditches, and a complex combination of black earth and clay in the structure of the embankment. Ritual steppe architecture of this period acquires considerable monumentality, and its main structural features resemble European funeral megalithic architecture of central and northwestern Europe. The earliest such structures in the NPR appear in the steppe-Trypillia contact zone of Pobuzhzhia (the Southern Buh basin) and the Dniester-Prut interfluve[42]. These complexes chronologically correspond to Trypillia BII-Cl and CI, as well as CI-II in burials of Lower Mykhailivka and post-Stih burials in the northwest steppe. Ritual monumental structures from the Eneolithic period are also known in the lower Dnipro and the Molochna River basins. Similar burial structures, containing ditches and cromlechs, accompany burials containing Maykop-Novosvobodnaya burial goods in Siverskyi Donets, Lower Don, and the Transcaucasia.

In the middle Eneolithic, the active advance of local Trypillian types into the steppe zone takes place, coinciding with the appearance of Trypillian mega-sites along the forest steppe-steppe boundary in south-central Ukraine. In this advance, Trypillians reached not only the more southern regions of Pobuzhzhia (seen a series of sites with painted Trypillian ceramics such as Novorozanivka, Tashlyk 2, Pugach, etc.) but also the Dnipro (exemplified by finds of clay Trypillian figurines and a flint axe on the east bank of the former Kakhovskiy reservoir near the village of Velyka Lepetykha, in the Kherson region). In contrast to the previous period (Trypillia B1, B1-2), when there was a certain balance between Trypillia and the steppe, the presence of steppe elements and imports in the material culture of Trypillian farmers at the end of the middle and the beginning of the late Eneolithic appears rather limited. Such as, steppe elements are absent at Trypillian settlements of this period, including the mega-sites. At the same time, Trypillian imports and influences are evident at cemeteries and settlements of the

Deriivka-type forest-steppe communities of the Stih horizon in the Dnipro Valley during this chronological period.

It is considered that steppe groups of the Serednii Stih horizon such as Post-Stih/Katarzhyno type contributed to the formation of the Cernavodă I archaeological group through interactions with Gumelnița and, potentially, remnants of Hamangia, ca. 4100-3900 BCE, and, eventually, contributed to the formation of Usatove after 3800 BCE[50]. By the end of the middle Eneolithic, Trypillian influences on Usatove become pronounced. In our analysis, Usatove is modeled as roughly even mix of Trypillia and Caucasus-Lower Volga (CLV) cline ancestries ([51], Supplementary Information, section 2). Individuals I20069 from Dănceni, representing the Vykhvatyntsi group of late Trypillia, and I7920 from Cunicea, belonging to the Gordinești late Trypillian group, are placed between the Ukraine Trypillia and Usatove clusters on the PCA in Figure 2. The Dănceni individual is a mixture of 53.7±5.5% Usatove and Trypillia in our analysis (Supplementary Information, section 2). Both individuals had the same diet, similar to that of Trypillian population from Verteba and distinct from that of the Usatove population from Mayaky (Table S4.3).

The late Eneolithic period (3500-3000 BCE) corresponds to Trypillia CII and represents a transition to the Early Bronze Age. This period is characterized by the fragmentation of the Cucuteni-Trypillia complex into a number of regional groups. This fragmentation also affected the steppe zone, where archaeological groups of the middle Eneolithic took part in the formation of separate local types, which, as a rule, were syncretic in their composition. Trypillia and Maykop-Novosvobodnaya either influenced or took a direct part in the formation of these steppe subgroups[42].

The intertwining of traditions is evident at the Lower Mykhailivka monuments from the second half of 4$^{th}$ millennium BCE in the eastern part of the Pontic steppe (to the south of Zaporizhzhia and the north of Tavria). Their graves, under kurgans, consisted of several separate burials, in which the dead were buried in a contracted position on their side. Quite often these burials were accompanied by small round-bottomed and flat-bottomed pots with river shell admixtures in clay. The surface of most of the pots was well smoothed - almost to a shine. Some, mostly flat-bottomed, pots contained thin rope imprints. This flat-bottomed ware imitated the kitchen ware traditions of late Trypillia, while polished ware reflected the influence of late Maykop. The Lower Mykhailivka burials were accompanied by typical late Maykop pots as well[52]. Bearers of the Lower Mykhailivka traditions were among the first full pastoralists in the North Pontic steppe with a developed understanding of domestication and the use of secondary animal products such as milk. Faunal assemblages of Lower Mykhailivka are dominated by domestic ruminants such as sheep and goats, followed by cattle, with a very minor inclusion of non-ruminant herbivores such as horses[53].

Further to the north, in the territory between Southern Buh and the Dnipro, the coexistence and displacement of different traditions, both in ritual practices and in material culture was even more pronounced. Such as, late Eneolithic populations of the Dnipro-Buh group, exemplified by kurgans near Pokrov and Shyroke in the Dnipropertrovsk Region[54], combined features of Lower Mykhailivka, Kvityana, Post-Stih and Trypillia. It is not uncommon for the Dnipro-Buh group to house, under the same kurgan, a group of several burials in which the dead were buried in extended supine, as well as contracted on the side and back burial positions.

The syncretic composition of steppe communities of the late Eneolithic is exemplified by the monuments of the Zhyvotylivka-Volchans'k (ZV), or III-C, burial type[55,56]. These burials are characterized by the contracted on the side burial position with palms in front of the face, heads in the southern direction, accompanied by arcuate staff-shaped bone and metal Bolgrad-type pins imitating the staff-shaped pins of Maykop-Novosvobodnaya, as well as late Trypillia painted ceramics and Maykop-type incised ("parquet"-style) and glazed pottery[57–59]. People associated with ZV/III-C monuments utilized ceramic vessel forms, such as amphoras, goblets, and bowls, made using atypical technology for the steppe.

The ZV/III-C burial type is represented in the steppe zone exclusively by burials in kurgans. The spread of ZV/III-C burials over an entire North Pontic area between the Danube and the Prut in the west to the Lower Don in the east, as well as the absence of settlements, reflects the nomadic and highly mobile lifestyle of ZV/III-C. Most ZV/III-C burials in the Pontic steppe are dated to ca. 3300-3000 BCE (this report). Taking into account two Bronze-Age burials from Taraclia II.10.2 (2566-2347 calBCE) and Tiraspol 3.15 (2865-2576 calBCE) with the III-C burial features, the terminal phase of ZV/III-C burials may have extended into the middle 3rd millennium BCE.

The ZV/III-C monuments are considered by some archaeologists in the context of the "Gordinești-Late Maykop phenomenon", in which six groups are distinguished: I - Bursuceni (Carpatho-Dniester region), II - Zhyvotylivka (Dnipro region), III – Volchans'k (Azov region), IV - Crimea (Crimean Peninsula), V - Lower Don, VI - Kuban (North Caucasus)[60]. Other ZV/III-C group subdivisions have also been proposed[57]. Thus, the ZV/III-C territory encompasses the entire North Pontic steppe and their existence overlaps with the initial expansion of the Yamna Archaeological Complex (YAC). In fact, the present report shows that people of the ZV/III-C group where among the earliest carriers of Yamna genetic ancestry, admixed with steppe Maykop, in the NPR.

The profound impact of Maykop on Late Trypillia, Usatove, and ZV/III-C groups resonates throughout various facets of their existence, evident in an array of archaeological discoveries from the late-4th millennium NPR steppe and forest-steppe. Notably, typical Maykop artifacts, such as metal implements like socketed axes and flat-handled daggers, alongside flint artifacts like diamond (trapezoid)-shaped arrowheads, asymmetrical with one shortened spiked and flag-shaped, or symmetrical/asymmetrical dart tips/knives, showcase this influence. The archaeological findings also include woodworking tools like adzes with asymmetrical blades and grooved chisels, jewelry like pendant rings made of silver, gold, or copper with one, one and a half, or several turns, as well as staff-shaped pins made from various metals or bone. Moreover, unique ceramics, distinguished by their shapes and ornamentation techniques, further attest to the far-reaching influence of Maykop on these archaeological groups[61].

Some scholars suggested the appearance of burial monuments displaying the III-C type features, which also include a specially designated place for the burial ritual involving fire and sacrificial offerings and the contacted on the side burial position, as reflecting the emergence of an institution of social, ideological and trade leadership in the NPR[60].

The highly mobile bearers of ZV/III-C traditions were in contact with communities interspersed throughout the Ponto-Caspian steppe. The system of relations, built on kinship and socio-economic ties, was accompanied by the reception and transfer of information and new technologies, thus facilitating the formation of a unified system, in which late Trypillia and Maykop might have acted as catalysts for the creation of the cultural and ideological basis for the emergence of a new socio-economic system manifested by the YAC.

In the second half of the 4$^{th}$ millennium BCE, late Trypillian groups such as Gordinești in the middle Dniester (3400-3100 BCE) show an increase of the CHG-rich "steppe ancestry" component[62]. At least some individuals of late Trypillian groups retained their mostly EEF-derived ancestry (the Vykhvatyntsi individual I7920 from this report, and the Pocrovca 2 individual from[62] of Starčevo-related genetic ancestry), while others displayed variable levels of "steppe ancestry". The emergence of highly mobile steppe groups such as those that left ZV/III-C type burials, connected the Carpatho-Dniester region with the North Caucasus and both became linked to a vast steppe area, where the process of synthesis and integration of diverse cultural formations took place, culminating in the emergence of the Yamna phenomenon, the core genetic ancestry of which appeared in the NPR around 3600-3300 BCE.

While the Eneolithic Trypillian population initially remained marginally affected by the CLV migrants, Serednii Stih were shaped by an intermixture of people from the CLV cline with the descendants of the Neolithic hunter-gathers in the NPR steppe, resulting in all SSAC individuals sampled being of an admixed CLV-hunter-gatherer background. Trypillian resilience to the eastern admixture finally ended in the second part of the 4$^{th}$ millennium BCE[62], on the eve of the Yamna expansion.

##### *1.2.1. West-Northwest Pontic coast from the 4$^{th}$ millennium BCE to the Early Bronze Age transition (3300-3000 BCE)*

In the early Eneolithic, Gumelniţa-Karanovo VI-Kodžadermen (GKK) archaeological complex and the related Varna archaeological group developed from the preceding Neolithic farming communities of the Balkans. Both GKK and Varna show a decline starting around 4200 BCE. The upper layers of GKK tells contain evidence of destruction. Scholars attributed this destruction to internal conflicts as well as conquest by the steppe groups of the Serednii Stih complex from the north Pontic[46,48]. As the northeastern part of the Balkan Peninsula becomes depopulated, settlements connected with GKK begin to appear in the Danube Delta and further northeast[46]. Such as, the Kartal archaeological complex on the Danube contains layers of GKK followed by Cernavodă I. In the Dniester Delta, a population center of Mayaky associated with the Usatove archaeological group and displaying influences from Cernavodă I (the Khadzhider type), CTAC, and the steppe, begins to function. The Cernavodă/ Khadzhider population at Kartal displays a genetic continuum from GKK (Kartal B) to steppe (Kartal A). Kartal A genetically overlaps with the Usatove population from Mayaky and Usatove-Velykyj Kuyalnik[63].

The Usatove archaeological group stands out for its unique ceramic style, impressive burial structures, and established metalworking practices. Concentrated primarily within the Dniester-Danube interfluve in Ukraine, approximately 35 sites bearing Usatove characteristics have been identified [50], although the total count could reach as high as 100 [64]. Notably, the largest Usatove sites in Ukraine—Usatove-Velykyj Kuyalnik and Mayaky—comprise both kurgan and ground cemeteries. The Mayaky site, in particular, features an elaborate system of ditches, variously interpreted by scholars as a sanctuary, a hillfort ("*horodyshche*"), or a causewayed camp [65].

Usatove can be categorized as “Steppe Trypillia” [43], reflecting the influence of Trypillia and the steppe groups of the Serednii Stih archaeological complex on the material culture of the Usatove archaeological group. Usatove is considered synchronous with the CII period of the Trypillia, which corresponds to *ca.* 3550-2750 BCE [6,66], marking the transition from the Eneolithic to the Bronze Age in the NPR chronology. Pottery of Eneolithic steppe (Serednii Stih, Lower Mykhailivka) origin and the steppe-derived elements in Usatove-specific pottery, cranial morphology parallels between Usatove and steppe forager-nomads, and signs of social hierarchy recognized by some scholars in burials attributed to Usatove [67–69], reflects steppe influences on Usatove.

The burial mounds of the Usatove elite are distinguished by complex funeral structures such as cromlechs, stone shells, anthropomorphic steles, ditches, cult pits with pots and animal bones, accompanying the burials of people and animals. Among the inventory there are often items of weapons and/or insignia of power. However, such a common type of offensive weapon such as a bow and arrow, is practically not represented in the Usatove complexes. Apart from isolated finds in kurgans and mixed layers of the settlements of Usatovo- Velykyj Kuyalnik and Mayaky, bifacial triangular arrowheads characteristic of the Eneolithic–Early Bronze Age were found in only two burials. In both cases, their location indicates that they were lodged within the body.

Usatove has material culture connections to Gumelnița, Cernavodă I, and Trypillia and Suvorove-Novodanylivka or their Post-Stih successors in the northwest Pontic steppe [11,12,43,46,50,65,70–72]. The presence of burials of the Yamna archaeological complex (YAC) at Usatove culture sites[12] indicates potential interactions between late Usatove and the YAC. In a series of burial mounds on the east bank of the Dnipro from the rapids to the river delta and in the Azov region, burials corresponding to the Usatove canon and containing Usatove ceramics and figurines, are stratigraphically located either under early YAC burials or sites with Mykhailivka-type features. Individual finds or imitations of Usatove sculpture are known even from the Crimea and the North Caucasus. Maykop influences can be traced in ceramic and metal finds from the Usatove graves at Usatovo- Velykyj Kuyalnik[73].

Among the burial goods of Usatove, the uncharacteristic for the Northern Black Sea region of the Early Bronze Age finds such as microlithic trapezoids, have been reported. Based on traceology analysis, it was concluded that almost all trapezoids were inserts for composite coarse-toothed sickles. Only one item was recognized as a hunting weapon insert. It should be noted that in the Usatove culture sickle inserts are known on massive flint plates. Analysis of the context of microlith finds in burials led to the conclusion that they were transversal (cross-blade) arrowheads[74]. These arrowheads, as well as other items of material culture, link Usatove with the eastern Mediterranean and Mesopotamia[46] (further discussed in Section 1.2.2).

Using comparative analysis of ceramic imports from securely dated periods and backed by radiometric radiocarbon dating on charcoal, pottery, and animal bone from Usatove sites, the current chronology of the Usatove archaeological group aligns with the period approximately ranging from 3650 to 2700 BCE.[12,71,75]. However, dates as early as the end of the 6$^{th}$ millennium BCE and as late as the last quarter of the 3$^{rd}$ millennium BCE have been reported for Usatove sites as well [12,71]. Radiocarbon dates on human remains from the Mayaky necropolis assigned to the Usatove period obtained by Accelerator Mass Spectrometry (AMS) do not align with radiometric dates from charcoal and pottery from the same site [5,12,75]. The discrepancy was attributed to radiocarbon reservoir effect (RE), inferred from elevated ratio of stable isotopes of nitrogen [5,12,75]. High $\delta^{15}N$ ratios accompany aquatic-resource-based diet, leading to a reservoir offset (RO) (e.g., [15,76–82]).

The RO at the Mayaky site has previously been quantified using ROs from Meso-Neolithic sites in the middle and lower Dnipro Valley, based on the assumption that radiocarbon dates obtained on human remains of populations living along the same drainage basin and consuming foods largely derived from freshwater resources will be under the influence of comparable ROs[5]. Archaeological reports indicate a significant presence of remains of aquatic animals at the Mayaky and Usatove-Velykyi Kuialnyk sites. For Mayaky, this evidence is summarized in[75]. At Usatove-Velykyi Kuialnyk, it was determined that fishing in the Khadzhybey Estuary, but also the Black Sea, occupied a prominent place in the daily activities of its inhabitants. The Khadzhybey Estuary was significantly desalinated by rivers, making it habitable for catfish, sturgeon and other fish, the remains of which have been found at the Usatove-Velykyi Kuialnyk site. Stone sinkers were found at the main territory of the Usatove-Velykyi Kuialnyk site, while bone fishhooks were found in the kurgans, suggesting that fish were caught using nets and fishing rods. Large quantities of mollusk shells were also found at Usatove-Velykyi Kuialnyk, dominated by *Mytilus* and *Cardium (Cerastoderma) edule* saltwater bivalves[69(p.148)].

Diet based on aquatic resources from the Black Sea or its brackish estuaries can offset radiocarbon dates by 545±70 years[83]. This offset was calculated based on human remains and contextual fauna from the Iron Gates area on the Danube. Comparing the stable isotopes of carbon and nitrogen from the Iron Gates area (N=140 [8,84,85]) with those from Eneolithic Mayaky samples attributed to Usatove (N=16 [5,12,75], this report) reveals that they are within the same distribution (2-tailed Mann-Whitney test $p$=01928, $z$-score=2.34494 for $\delta^{13}C$, p= 0.06724, $z$-score=-1.83128 for $\delta^{15}N$, significance level 0.01). At the same time, diet isotopes from Mayaky are outside of diet isotope distribution of hunter-fisher-forager populations of the Dnipro Valley[16]. Therefore, in this report we used the 545±70 year offset calculated for the Iron Gates, to calculate the RO for the Eneolithic burials at Mayaky where stable isotope ratios suggest aquatic resource input in diet. Using the Iron Gates offset, the RE-adjusted average date for the majority of Eneolithic burials at Mayaky came to 3708-3109 BCE (Table S1).

The upper date of the Mayaky site chronology correlates well with the date obtained on a genetically Usatove individual from burial 19 in the Revova kurgan, the date from which does not bear evidence of RE influence (Table S4.3).

Other Usatove populations such as at the eponymic Usatove-Velykyi Kuialnyk, as well as the population at the Cernavodă I site at Kartal on the lower Danube, likely used similar aquatic subsistence resources. Therefore, the RO at Usatove-Velykyi Kuialnyk and Kartal should be comparable to that at Mayaky. Adjusted for the Iron Gates RO, the chronological average for the Usatove-Velykyi Kuialnyk date comes to 3361-2783 calBCE, and the average date for the Cernavodă I burials at Kartal is 3363-2784 calBCE, based on the absolute $^{14}C$ dates for these two sites reported in[63].

Combining RE-adjusted dates for human remains at Mayaky and Usatove-Velykyi Kuialnyk, the chronological timespan of Usatove comes to ca. 3700-2800 BCE, in where Mayaky begins to function ca. 300 years before Usatove-Velykyi Kuialnyk. Overall, the RO-adjusted $^{14}C$ AMS-generated date on human remains of Usatove is in full agreement with the 3650-2740 BCE radiometric date from pottery and animal bone from Mayaky, obtained by the scintillation $^{14}C$ technique[75], and thus fits the currently accepted Usatove chronology. The RO-adjusted chronology of Cernavodă I, based on human remains at Kartal, is within the 3400-2800 BCE time period, thus postdating the beginning of Usatove by ca. 300 years.

At the same time, the formative phase of Usatove genetic ancestry using DATES[86] is estimated to be 4471±51 BCE (Table 1), which correlates with the earliest $^{14}C$ dates (not RE-adjusted) from Mayaky (Table S1, [75]). Therefore, the RO calculations presented here may reflect the main phase of the Usatove existence, while the formative phase ("archaic" Usatove[43,75]) likely dates to the middle of the 5th millennium BCE, coinciding with (or shortly following) the formation of Trypillian ancestry at 4595±121 BCE (Table 1).

##### *1.2.2. Contacts between the NPR, the Caucasus, and the Near East*

Several lines of archaeological evidence point to connections between the populations of the North Pontic, Near East, and the Caucasus, beginning in the Neolithic.

The shape of bracelets found at the Neolithic Mariupol Necropolis shows analogies with finds from the early burial phase at the Nal'chik Necropolis in the North Caucasus[1]. The style of Mariupol mace heads and bracelets was further suggested to originate from Mesopotamia[1,87]. Shell beads found at the Mariupol Necropolis show analogies with the material from the Neolithic burials at Kaylu in the southeastern Caspian, which, correspondently, are similar to those from the Neolithic sites in the southwestern Caspian Sea area[88]. Early Serednii Stih ceramics display analogies with pre-Maykop ceramics of Svobodnoe, pointing to the existence of trade/exchange relationships between the Serednii Stih of the Dnipro Valley and populations of the North Caucasus Piedmont[89].

Mesopotamian influences on CTAC plastics and ornaments on pottery are evident on artifacts dated to ca. 4100-3500 BCE[90]. Images on Trypillian ceramics of figures in dresses with fringe-trim edges[91], also found on Sumerian dress depictions (see images below), appear after 3500 BCE.

Individuals I17973 from Bursuceni and I20079 from Taraclia in the sample selection presented in this report were interred with staff-shaped pins made of bone. Several late Trypillia, Usatove and Zhyvotylivka-Volchans'k (ZV) burials across the north Pontic (Group III-C burials according to Y. Rassamakin[56]) contain similar pins[61,92], often accompanied by beads[57,60], like in the case of Individual I20079.

Stylistically similar pins made of gold, silver and copper alloy and accompanied by suspended beads and cylinder seals have been uncovered at the Early Metal Age sites in Mesopotamia such as the royal Early Dynastic period cemetery at Ur[93,94]. Copper beads resembling Mesopotamian cylinder seals are extensively found in late Trypillian metal hoards[95]. It is conceivable that the hammer-top pins of the YAC, sometimes found in association with stung beads[96] were patterned after the Mesopotamia pins and symbolized a particular social status.

Burials of the III-C type are often accompanied by glazed and pierced Maykop-style pottery. The appearance of pottery with pierced ornaments in Maykop ceramics is considered to be an influence of southern Mesopotamian Uruk tradition. In eastern and southeastern Anatolia, northern and western Syria, ceramics with this type of ornament remained in use during the Late Chalcolithic IIb - Early Bronze Age IA period, or approximately the second half of the 4th millennium BCE. North of Mesopotamia, similar pottery has been found in significant quantities at settlements along the Upper Euphrates.

Another group of finds potentially linking the Eneolithic NPR to the Near East are trapezoid flint microliths. Traceology analysis of these artefacts from the Usatove culture monuments revealed they had a multifunctional purpose, serving as compound sickle and knife components and, possibly, as arrowheads. Many of these, however, did not show any signs of utilization, suggesting that they had a ritual or ceremonial purpose[97]. The association of such microliths with elite Usatove graves such as burial 18/1 at the Usatove site of Mayaky[98] strengthens the latter assertion. In at least three elite Usatove burials trapezoid transversal microliths were found in compact clusters and single trapezoid microliths were found in close association with human remains[74]. Transversal flint microliths, along with a staff-like pin, were found in a III-C type burial 28 of the Slobodzeya Kurgan 1[99]. A microlithic trapezium was also found in the main burial (#18) of the YAC at Glinoe-DOT Kurgan 1, extending the presence of such microliths in the NPR into the EBA. The calcified trapezium was located close to the femur, suggesting its association with the remains to be a result of a healed injury[74].

The use of arrows with transversal microliths as tips is known in Egypt of the predynastic period, the Old and Middle Kingdoms, traced both from images and finds of arrows in tombs, as well as images of arrows with transversal tips on reliefs and cylinder seals from Uruk (Sumer, late Uruk period, last quarter of the 4th millennium BCE) and Knossos (Crete)[74]. It is notable that the use of microlithic transversal arrowheads in the NPR ceased almost 1500 years before the appearance of the Usatove archaeological group[74]. Late Trypillia cultural groups and the cultures of the steppe pastoralists of the NPR used bifacial triangular points. At the same time, a trapezoid microlith has been found at the Trypillian megasite of Nebelivka[100].

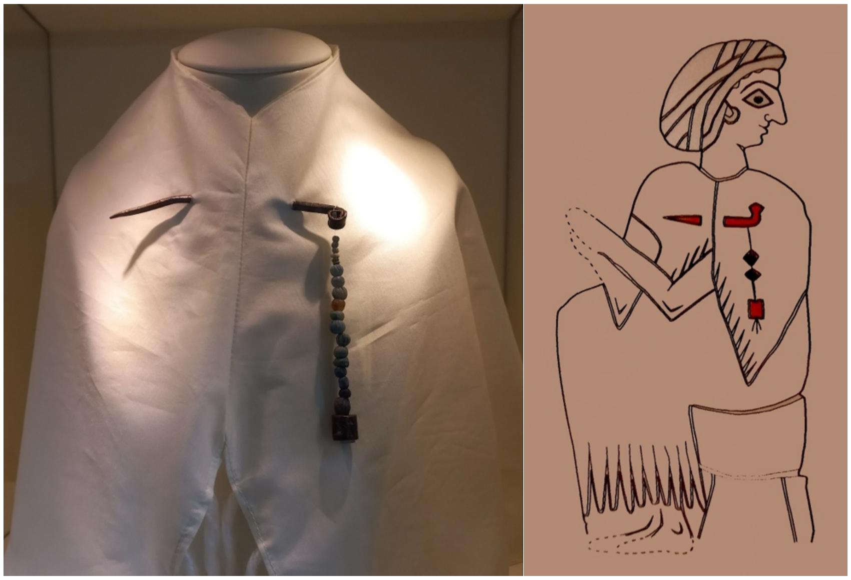

*A cloak pin with beads and cylinder seals suspended from it. Uruk, 4th millennium BCE. Neues Museum, Berlin. Photo: M. Y. Videiko, 2023.*

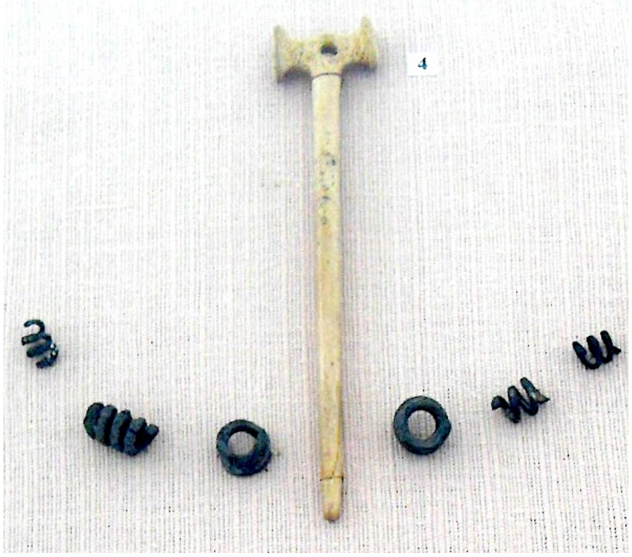

*Hammer-top pin of the Yamna archeological complex. Stari Bilyari Kuragn 1 Burial 4.*
*Photo: S. Ivanova, undated.*

#### 1.3. The Early Bronze Age (3300/3000-2400 BCE)

In the last third of the 4th millennium BCE, the transition to the Early Bronze Age (EBA) in the Pontic steppe is marked by the appearance and rapid spread of the traditions of the YAC. At their maximum extent, the Yamna groups occupied the area from the middle Danube and the northern slopes of the Balkans in the west to Ural Mountains in the east. Several archaeological groups are variably recognized by archaeologists throughout the Yamna domain. In the NPR, these include the Donetsk, Middle Dnipro (Northwestern), Southwestern, Lower Dnipro, Azov-Crimean and Southern Buh groups, with additional divisions proposed within each region. The differences between Yamna regional groups are observed primarily in the material culture, especially, in ceramics and its typology. At the same time, the regional divisions within the Yamna complex that were established by the degree of similarity of ceramics often do not coincide with traditional local variants. Furthermore, comparisons of published ceramic series from individual Yamna regions often show more differences than similarities[101].

Yamna is known predominantly from burial monuments and funeral rites; settlements are rare. In fact, the Yamna archaeological complex is defined primarily by burial traditions, rather than characteristics of the material complex, as it is traditionally done in archaeology, where ceramics provides the most distinguishing features[101].

Yamna burials utilized kurgans constructed in the Eneolithic, but the Yamna also erected kurgans of their own. In the lower Don, there were more than 300 Eneolithic-EBA kurgans and 1,500 Yamna burials excavated by the end of the 20th century[102]. In Romanian Transylvania, the number of the Yamna-age kurgans is estimated to be over 600[103]. As of 2020, 177 kurgans containing 714 Yamna burials have been investigated in Romania. There is information of about 75 investigated Yamna kurgans in Hungary, and about 15 Yamna kurgans in Serbia. According to the latest publications, 80 kurgans containing 460 Yamna burials have been excavated in

Bulgaria[101]. The vast majority of Yamna kurgans are located along the river valleys of the Ponto-Caspian steppe, with their highest concentration in the northwest Pontic[104,105].

As of the late 1980s, the number of excavated Yamna burials in Ukraine exceeded 10,000 (M. Y. Videiko, personal communication, 2020). Just between the Prut and Dniester rivers there are over 700 kurgans and over 5,000 burials that were excavated and published, but it is also estimated is that there are over 10,000 kurgans in the Prut-Dniester interfluve, with over 50% erected during the Eneolithic-EBA[105]. The highest density of Yamna graves is documented in the Southern Buh area, while the lowest density is found in the Lower Don region. The Yamna burial density as well as the number of stratigraphic phases attributed to the Yamna period pronouncedly decreases towards the Volga-Ural periphery of the Yamna domain[104].

Yamna kurgans typically exhibit a complex stratigraphy. In the mounds constructed above the primary (main) burial, additional (inlet) burials were subsequently added. Larger kurgans can contain several subsequent mounds or local fills associated with a group of burials or a specific complex. Burials within each mound layer, especially if numerous, are arranged in arcs or circles, possibly linked to ancient beliefs about celestial body movements[106]. Elements of mound architecture include ditches, cromlechs, and retaining walls. Burial chambers, mostly rectangular, consist of simple ground graves and those with ledges, with the latter comprising about 30%. Some grave pits have ceilings made of logs or stone slabs. Anthropomorphic steles are occasionally found as part of the stone ceiling, and wooden carts or parts thereof are rare elements in the funeral ritual. One Yamna burial in the northwest NPR dated to 3315-2916 calBCE contained a stylized boat made of wooden planks as part of the funerary installation[107].

The position of the deceased in Yamna graves varies considerably. Some researchers identify five main types based on the position of the arms, legs, head, and body inclination, with each type having numerous subvariants[108]. These five types include (1) flexed on the back with bent legs (knees up), arms extended along the body or slightly bent, hands on hips, sometimes legs arranged in a diamond shape (57.2% of the burials); (2) tilted to the right, with left arm bent at the elbow, hand near the pelvis, abdomen or chest; the right arm extending along the body (16.3%); (3) tilted to the left, the right hand is located at the pelvis (13.1%); (4) on the right side, with different positions of the hands (7.3%); (5) on the left side, with different positions of the arms (6.1%). Some researchers propose three (back, the right or left side), or two groups (contracted on the back and contracted on the side)[64,105].

Analyzing main Yamna burials reveals a predominance of the flexed on the back group, possibly indicating social stratification[109]. The Moldovan archaeology scholars Valentin Dergachev and Evgeni Yarovoy consider that early Yamna burials before 2500 BCE were flexed on the back, while flexed on the side and tilted burials occurred thereafter. However, exceptions exist, challenging a clear chronological pattern. Other archaeologists suggests that burial postures of Yamna burials may be a social factor rather than chronological indicator[106]. Some argue that the flexed on the back position characterizes the early chronological stage, while flexed on the side to be the feature of the late stage[108,110,111]. The supporters of the idea of correlating burial position with chronology emphasize that the flexed on the side position mostly present in inlet burials. Yet, when planography data and stratigraphic layers of the Yamna kurgans are considered, the simultaneous occurrence of various positions within one horizon becomes evident[112]. Such as, in 95% of multi-layer Bugeac kurgans, there are horizons with mixed ritual traditions.

Arguments against a clear chronological dependence on burial position include the fact that all burials in an existing kurgan are chronologically later than the primary burial. However, the relative simultaneity of subsequent burials and their formation into single complexes with the main burial raises questions about the time elapsed between them. Speculation remains regarding how Yamna burial traditions reflect societal structure, the deceased's status, and their connection to the kurgan's symbolism as a sacred object in this stage of understanding prehistory[113].

The funeral tradition in the EBA became more unified across the Yamna territory, compared to burial practices of steppe populations from the preceding chronological period. At the same time, differences, primarily in ceramics, as well as in the number and assortment of metal artifacts, persisted among different Yamna groups. This was due, on the one hand, to different Eneolithic substrates in different territories of the YAC, and on the other, to cultural contacts.

One notable example of regional YAC specifics is the Bugeac archaeological group in northwest NPR. Bugeac ceramics, unlike in other YAC regions, have a flat bottom. Some of the vessels demonstrate the influence of archaeological complexes and groups of the Balkan-Carpathian area and Central Europe (Corded Ware, Globular Amphora, Cernavodă II, Kotsofeni, Glina III-Schneckenberg, etc.), and some are imports. At the same time, specific forms of vessels were created, characteristic only of the Bugeac ceramic complex. In addition, in contrast to synchronous archeological groups of the North Pontic and the greater Carpatho-Balkan area, a concentration of silver jewelry is observed in the Bugeac region in the Usatove and YAC Bugeac archaeological contexts.

The distinctive funeral rituals of the Bugeac group are characterized by a kurgan, featuring the main burial at its center in a rectangular burial chamber covered with either wooden or stone slab ceilings (one-third of the burials incorporated ledges), a plant mat beneath the body, and the use of ocher. The deceased is often placed in a contracted position on their back or side, with a prevailing western orientation. Main and inlet burials are often arranged in circular formations, and intricate mounds with multiple stratigraphic horizons are occasionally observed. These elements, observed in diverse forms and combinations, are shared with Late Eneolithic groups across the region, amalgamated into a cohesive complex within the Bugeac group.

The cultural exchanges between the Bugeac Yamna and populations in the Danube region are evident in the ornamentation of pottery, as well as the introduction of askos-type vessels and cups. Furthermore, in the northern part of the Bugeac group territory, the influence of the Globular Amphorae archaeological complex is discernible through the import and imitation of ceramics. Additionally, there is speculation that the Bugeac group played a role in transmitting the ceramic traditions of Usatove to the Złota group within the Corded Ware archaeological complex[114]. Occasional imports from the Balkan-Carpathian area can be traced in YAC groups further to the east, such as at Sofyivka 10/1, the Kherson region, and the western bank of the Dnipro[115].

Ceramics such as cups and amphorae, reflecting connections with the Corded Ware archaeological complex, are distributed throughout the northwest Pontic, concentrating along the Dniester and in the south of the steppe zone. Amphoras are often imported, while cups are imitations, their ornamentation is different from the classical canons of CWC and distorted. Probably, it was the Bugeac population that was responsible for transferring the Corded Ware features to the west and east, evidenced by a corded cup found in the Crimea (Istochnoe 12/5)[116] as well as cups of the same type from Romania, Bulgaria and Hungary. The discovery in 2023 of a similar cup near the village of Orlivka, located at the crossing of the Danube, is likely part of the same transfer process (the authors are grateful to Dr. Igor Bruyako for access to the information about the Orlivka excavations).

Around 2700 BCE, the Catacomb Archaeological Complex (CAC) emerged in the north-eastern Azov and the lower Don region[117] and later expanded across a significant part of Ukraine. Researchers date the entire existence of the CAC to 2700–2000 BCE, with the early stage, marking the formation of Catacomb groups, occurring between 2700–2500 BCE[118]. Excavations in the 1890-1990s uncovered over 1200 CAC burials.

West of the Ingulets River, only isolated burials from the early stage of CAC are known[119]. The CAC tribes' settlement involved interactions with the local, predominantly Yamna population, evident in the material culture inventory[117,120]. Even in its early stage, the CAC lacked unity, exhibiting territorial features that later led to the formation of distinct Catacomb groups during the developed stage (2500–2300 BC), characterized by relatively stable sets of characteristics[118].

The CAC comprises several regional groups characterized by different burial rites, including the Donetsk and Ingul groups in the Pontic steppe. The Donetsk group features a flexed and tilted-to-the-right body position in graves, while the Ingul group is characterized by an extended supine position. The catacomb shapes and ceramic complexes also differ between these groups.

The eastward expansion of CAC reached beyond the Danube, displacing the Yamna from the archaeological scene by the mid-3rd millennium BCE. Radiocarbon dates from a small sample of the CAC in the northwest Pontic align with the ca. 2900-2000 BCE range. The CAC' final stage in the region coincides with the initial stage of the Babyne cultural circle.

In our sample selection, CAC individuals I11850 (ODK1, 2874-2630 calBCE, the Odesa Kurgan) and I12840 (DBN1.10, Dubynove Kurgan 1 Burial 10, 2453-2148 calBCE), cluster with the main Yamna group. Successful models for the I11850 ancestry include some CLV Cline ancestry, and the best fit model is PVgroup+Trypillia (Supplementary Information, section 2). Individual I12621 (DBN1.9, 2575-2348 calBCE) aligns with the main Yamna cluster (Figure 2a) but leans towards the Caucasus. Catacomb individual I13072 from Soldats'ka Slava is on the opposite side of the main Yamna cluster, near Serednii Stih specimens.

Catacomb Individual I16668 from Revove stands alone on the PCA in Figure 2a, leaning towards the main Usatove cluster. This position is intriguing, as the main burial in Revove Kurgan 3 genetically identified as part of the Usatove population, suggesting a genetic ancestry continuum despite the change in cultural traditions.

##### 1.4. The Middle Bronze Age (2400-2000 BCE)

The Middle Bronze Age in south-eastern Europe was marked by drastic climatic shifts, likely leading to the disintegration of the Catacomb cultural complex and the formation of a number of new archaeological units in its place.

Among the post-Catacomb formations, the Babyne archaeological circle (BAC), which covers most of the northern Azov-Black Sea basin, stands out for its scale and a series of distinguishing features. The formation and development of the Babyne cultural region affected the content and direction of cultural and historical processes in south-eastern Europe during the entire transition period from the Middle to the Late Bronze Age.

The Babyne circle includes the homogenous Dnipro-Don and the more fragmentary Dnipro-Prut archaeological groups. Within the latter, Dnipro-Dniester, Dnipro-Buh (forest-steppe), and Dniester-Prut (steppe) regional subgroups are distinguished. Babyne burial sites are represented by 1140 burials from 592 kurgan and 8 ground (kurgan-less) cemeteries, containing predominantly contracted on the side inhumations. A qualitatively new stage of kurgan construction in the Black Sea region is associated with the Dnipro-Don Babyne, marked by a rapid increase in scale and innovations in architecture, which, sometimes, included three to six Babyne building horizons. Furthermore, 79% of Babyne burials were accompanied by grave embankments and backfill. The Dnipro-Don group started the practice of erecting long graves by lengthening and unifying fillings. A striking feature of the Babyne kurgan architecture in the Donetsk region is the presence of a variety of megalithic elements. Around 270 settlements are associated with the Dnipro-Don Babyne archaeological group[121].

The Dnipro-Prut Babyne group stretched from the Dnipro Valley and the Crimea in the east to the west bank of the Prut and the lower Danube in the west. The Dnipro-Prut group is represented by about 170 settlements, up to 2500 burials, and several treasure hoards, potentially associated with Babyne[121].

A considerable part of Babyne material culture features, having no local roots, finds striking correspondences in the populations of the Bronze-Age Caucasus and the Carpatho-Danubian region, implying transregional culture synthesis to be part of Babyne's cultural identity. Material culture parallels with the NPR Babyne are recognized in Bronze Age populations of lower Don and Northern Caucasus. The Bronze Age Don-Volga Abashevo, Syntashta, and Potapovo “chariot culture” groups are partially synchronous with the Babyno circle, based on the relative stratigraphy, as well as analogies in the material culture. To the west, in the Transcarpathia and the Middle-Upper Danube, the Middle Rhine, and the Alps, post-CWC/Beaker groups such as Mierzanowice, Polada, and Únětice demonstrate a complex of analogies in funeral rite and material complex with groups of the Babyne circle. In the basin of the Middle-Lower Danube, several archaeological groups of the Middle Bronze Age of Hungary and the Early Bronze Age of Romania appear to be simultaneous with different stages of Babyne. The chronological successor of Babyne in the NPR is the Sabatynivka-Noua archaeological complex[121].

##### 1.5. Flat-top kurgan sanctuaries in the NPR and the origin of Yamna

Kurgans stand out as the hallmark features of the Yamna archaeological complex. Most surviving Yamna kurgans exhibit a rounded shape, although a substantial number also display a conical form. The origins of conical kurgan mounds date back to the Eneolithic period.

Throughout the Yamna period, truncated conical mounds with flat horizontal upper platforms were likely common in the Ukrainian steppe zone, though accurately estimating their prevalence remains difficult[96]. The compacted upper platforms of some of such kurgans in the NPR are coated with a thick layer of clay. These include such kurgans as Bakumova Mohyla near Mayaky with a clay horizontal platform 10 m in diameter, and Kurgan 22 near the village of Hlukhe with a 50 cm thick clay platform layer. Both kurgans had a height of 7 m in their finished form.

Flat-top kurgans are also known in the North Caucasus. These include Maykop kurgans such as Zamankul Kurgan 2, Klady Kurgan G and the massive Kurgan 11 that covered it, the Nalchik Kurgan, the Bolshoy Ipatovsky Kurgan (5th embarkment), etc. As in the NPR, the surface of embankment platforms of most flat-top kurgans in the Caucasus was finished with a layer of bright yellow clay coating[96].

One of the most architecturally complex flat-top kurgans in the NPR is the Shakhtar (Shevchenko) Kurgan 29. In addition to the horizontal platform at the top of the 4th embarkment, the Shakhtar Kurgan also had a ledge. The shape and structure of the embarkments of kurgans such as Shakhtar Kurgan 29 likely relate to their use not only as burial places but also as sanctuaries associated with the performance of ceremonial activities. The Lower Dnipro Valley houses a chain of kurgans with ramps and altar platforms with a pavement of turf-mud rolls, and with roads radiating from them. One such kurgan is the Vynohradne Kurgan 3 located in the Molochna River Valley (see Burial Descriptions), from which came one of the closest to the Core Yamna Serednii Stih individual in Burial 15[51].

An estimated 46 sanctuaries like Vynohradne Kurgan 3 with radially extending roadways are concentrated in the Dnipropetrovsk and Kherson regions of southern Ukraine, along the lower stretches of the Dnipro and Ingulets rivers[122] (Fig. S1). A particular dense concentration of such monuments can be found within ca. 60-km radius from the Eneolithic-EBA Mykhailivka and Kapulivka sites, the latter featuring Mykhailivka-type early Yamna ceramics[123] (Fig. 1). On the Ingulets River, kurgan sanctuaries extend from a river crossing at Davydiv Brid upstream to Kryvjy Rih, and from a river crossing at Kakhovka upstream to the city of Dnipro on the Dnipro. A flat-top kurgan near Pishchanka at the northern edge of the Dnipro River kurgan chain contains an overlapping sequence of Serednii Stih and Yamna burials, in which the earliest Yamna burial dates to 3626-3106 calBCE[124], the upper range of the date of which overlapping with the $^{14}C$ date from the earliest Core Yamna individual I32534 from Mykhailivka, presented in this report. The Pishchanka Kurgan is also in the vicinity of Orgin-8, from which one other closest to the Core Yamna Serednii Stih individual originates[51]. A line of sanctuaries with radially extending roads or erected at branching points of dried-up river valleys such as Forois (Zhovtneve) Kurgan 11 in the Tokmak District extends from the main concentration in the Dnipro-Ingulets interfluve northeast towards the Siverskyi Donets River (Fig. S1).

Taken together, the flat-top kurgan sanctuaries found in southern Ukraine establish a connection with the Maykop kurgans located in the North Caucasus. These sites in Ukraine encapsulate a continuous archaeological record spanning from the Serednii Stih era to the early Yamna period. Notably, they harbor the earliest chronologically dated Yamna individuals within the NPR. These complex architectural structures served a polyfunctional purpose, acting as both burial grounds and venues for communal ceremonial rituals. This ceremonial function, extending beyond mortuary practices, aligns with the notion that a fundamental aspect of the Yamna culture complex was the shared worldview embraced by its communities[46,101].

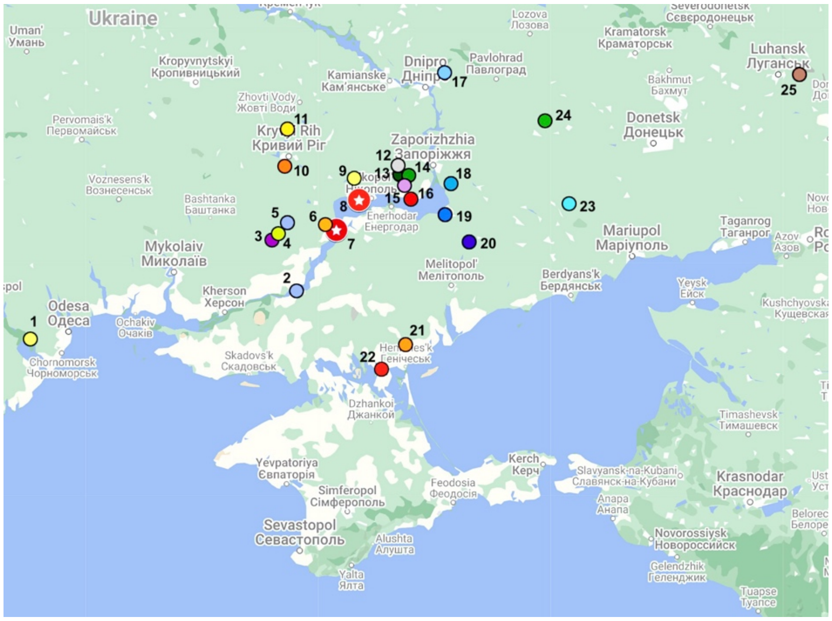

**Figure S1**. Flat-top kurgan sanctuaries and ancient river crossings in southern Ukraine. Base map image: Google Maps, maps.google.com
1, Mayaky (Bakumova Mohyla); 2, Kakhovka (Dnipro River crossing); 3, Davydiv Brid (Ingulets River crossing and a kurgan); 4, Mala Oleksandrivka; 5, Starosillya (Velyka Mohyla); 6, Shevchenkivka (Kurgan 14); 7, Mykhailivka; 8, Kapulivka; 9, Shakhtar (Kurgan 29); 10, Ryadovi Mohyly; 11, Nedaivoda (kurgan group "Three brothers"); 12, Kyslychuvata (Hola Mohyla); 13, Myrove; 14; Vysoke; 15, Hlukhe; 16, Vyshchetarasivka; 17, Pishchanka; 18, Hryhorivka; 19, Vasylivka (Kurgan 1); 20, Vynohradne (Kurgan 3); 21, Hordienkivtsi; 22, Vesnyanka; 23 Forois (Zhovtneve Kurgan 11); 24, Volodymirivka (Kurgan 13); 25, Sykhodil (Pioners'ke).

#### 2. Uniparental lineage connections in the Ponto-Caspian region

##### 2.1. MtDNA lineages

Our study uncovered an ample amount of mitochondrial lineage diversity in most populations studied, with some lineage sharing among the groups. Trypillian lineage diversity in our sample group is limited to the Anatolian Neolithic farmer (ANF)/EEF-derived haplogroups and includes members of previously reported mitochondrial lineages in Trypillia. The finding of additional lineages such as K1b2, H2a and H5b not previously identified in Trypillia adds to the substantial lineage diversity in Trypillia reported to date, suggesting a large population size of the founding population, as well as reflecting the overall size of Anatolian farmer-derived mating network and the absence of population bottlenecks [125].

Usatove mitochondrial haplogroup composition consisted of lineages of predominantly ANF/EEF origin (H*, K1b*, T2*, W1, X2d), as well as steppe and European hunter-gatherer-derived U4* and U5a1* lineages. The Mesolithic distribution of U5a1 lineages is primarily confined to the Iron Gates area, as well as Norway and the Baltic region [8,126]. In the Neolithic, U5a1 lineage distribution extends to the North Pontic, likely accompanying the migration of carriers of WHG ancestry to the Dnipro Valley [8]. Carriers of U5a1 in our study were identified in the Neolithic Mariupol-type cemeteries at Mariupol (individual I31730) and Yasynuvatka (individual I27982). Individual I17974 from Bursuceni carried the U5a1-derived haplogroup U5a1g1, previously identified in a Catacomb individual from Burial 11 at Dubynove Kurgan 1[5,13]. The U5a1g subgroup was previously reported in a CWC individual from Germany[127], roughly contemporaneous to the individual from Dubynove.

Mitochondrial DNA of individuals I2108 from Ogrin-8 (Serednii Stih) and I12637 from Glinoe (Yamna) belonged to the U2e* lineage. Haplogroup U2 has been present in the Don River Valley in the northeastern NPR since the Upper Paleolithic[128,129]. Carriers of U2e* have been identified in the Mesolithic Baltic, Scandinavia and Neolithic Ukraine[8,130,131].

The U4-derived lineages found in our study are shared with prehistoric groups regionally and throughout Europe. Phylogeographic distribution of the U4 node and its derivatives in pre-Eneolithic Europe includes the Iron Gates area of the Danube, the Baltic region, and the Middle Dnipro Valley [8,126,132]. Chronologically earliest representatives of the U4 haplogroup in Europe date to the Epipaleolithic and come from the Dnipro Valley [5,133].

The chronologically earliest individual I12615 from the Mayaky archaeological site shared mtDNA lineage U4b1b2 with the individual of Serednii Stih ancestry (SSmed) from Kolomiytsiv Yar Tract (KYT) [16]. It has been suggested that the U4b1b1~ lineage represents the pre-Neolithic mitochondrial lineage diversity, expanding from the Mesolithic Balkans and not being directly influenced by the steppe[132], at least in the initial stages of the lineage expansion. Both the Mayaky and the Serednii Stih sites of the Dnipro Valley are located along the route of the Eneolithic circum-Pontic trade network connecting the Balkans and the Eastern Steppe[16,46]. The presence of a Balkan-derived mitochondrial lineage in an individual of steppe ancestry supports the existence of mating interactions between the steppe and their trade allies from the eastern Balkans.

The post-Stih DNA from the individual K1.10 (specimens I12167/I31733) did not produce quality nuclear DNA data. The initial mitochondrial DNA analysis of K1.10 was reported in[27]. Subsequent sequencing of additional PCR amplicons of the mtDNA coding region from two additional extractions of K1.10 produced a recurring transition at nucleotide position 12308, suggesting the placement of the specimen in the U clade. Considering the previously reported lack of diagnostic polymorphic sites in the control region of mtDNA, no further refinement of the haplogroup designation for K1.10 can be made.

Mitochondrial DNA lineage U4c1 of the main burial of the Katarzhyno 2 kurgan has been identified in a Yamna burial from the Prydnystryanske kurgan 4 of the Yampil kurgan sequence from the middle Dniester area, as well as in one of the two early Usatove individuals from burial 6 of Kurgan 8[5,134]. The former lineage sharing adds support to the involvement of the Eneolithic communities from the greater northwest Pontic in the formation of the local forest-steppe Yamna group of the middle Dniester area[135]. The Post-Katarzhyno-Usatove haplogroup sharing supports the involvement of the Katarzhyno-type group or its ancestral population in the formation of Usatove[72]. On the other hand, mitochondrial DNA lineage sharing between the Trypillian individual from Soldanești and the ZV individual from Taraclia (I20079) carrying

Usatove genetic ancestry provides additional support for the genetic connection between Trypillia, Usatove, and ZV.

Two individuals of Yamna ancestry in Katarzhyno 1 kurgan shared their mtDNA (U4a) as well as Y chromosomal lineages. No IBD segments have been identified between the two, probably because of the large time gap between them. The uniparental lineage match between two individuals in the same kurgan raises the possibility of a familial relationship among the interred in EBA kurgans, especially considering that the mtDNA lineage of the main burial in the Katarzhyno 1 kurgan belongs to the U clade as well. However, we do not have enough data to test this relationship further.

Mitochondrial lineages of burials I13071 (Bil'shivtsi Burial 1, northwestern forest-steppe) I12005 and (Soldats'ka Slava Burial 10, northwest Pontic steppe) belong to the T2c1 haplogroup. Subdivisions of this haplogroup are found throughout Europe in the Neolithic-Bronze Age. The T2c1a2 sub-lineage carried by the Yamna I12005 individual has also been identified in a Yamna individual I0429 from Samara in Russia[127], suggesting maternal lineage ancestry sharing among the Yamna individuals across the Pontic-Caspian steppe in the EBA.

MtDNA lineage H13 of the Giurgiulești individual I20072 matches that of the Mesolithic Iron Gate individuals from Lepenski Vir and Ostrovul Corbului[8], suggesting potential genetic admixture between the Serednii Stih II/Skelya/Novodanylivka migrants from the Dnipro Valley and local populations of the Lower Danube. On the other hand, H13 could have arrived at the Dnipro Valley as part of the WHG Iron Gates ancestry in the Neolithic[8], thus becoming part of the Dnipro Cline ancestry from which Serednii Stih was 1/5$^{th}$ derived[51]. An H13-derived lineage H13a1 was recorded in a Lower Volga individual I22199 from Berezhnovka II[51], as well as in a Yamna individual from Ishkinovka in pre-Ural steppe[130]. Genetic ancestries of both I22199 and I20072 are derived from Lower Volga populations of the BP group[51].

Two contemporaneous Yamna individuals from Moldova from Sărăteni and Cotiujeni shared an H13-derived lineage H13a2b2a. Outside of the Pontic steppe, this lineage was found in Bronze Age Germany[136].

Sublineages of the H6a1* haplogroup of the Eneolithic early Yamna individual from Mykhailivka and an EBA Yamna individual from Sărăteni have widespread distribution in Bronze Age Europe. The earliest find of H6a1* outside of the NPR is in an EBA Yamna individual from Kutuluk River near Samara[130]. The distribution of the H6a1* lineages in Europe and Central Asia suggests their spread is linked to the Yamna expansion.

#### 2.2. Y lineages

Our study shows a diverse array of Y lineages of our sample selection of the North Pontic area, represented by autochthonous European and Anatolia-derived lineages, as well as lineages that are found in prehistoric populations of North Caucasus.

Y chromosomal lineages identified in CTAC specimens in the present study include lineages of I2a1b, G2a, E1b1b, and J2a1. Considering the Y chromosome lineage diversity reported in previous studies of CTAC remains from Verteba Cave, the Y chromosome lineage diversity in CTAC appears to be quite substantial and on par with the mtDNA lineage diversity.

Two Y chromosomal lineages, G2a and J2a, found in CTAC specimens from Verteba Cave, are considered to have originated in Anatolia and are associated with the spread of agriculture out of Anatolia into Europe in the Neolithic [137]. The G2a2b2a lineage identified in two CTAC specimens in this study has been found in Neolithic northwest Anatolia, Neolithic Austria, Bulgaria, and Hungary, as well as early Eneolithic Bulgaria (Ivanovo) and Romania (Curătești) [5,8,138]. The latter two sites are located in the Lower Danube, either within or bordering the southeastern part of the Romanian historical region of Muntenia, also home to the Eneolithic Bodrogkeresztúr culture, with which CTAC has been previously shown to share genetic ancestry[8].

Individual I17973 of the North Caucasus genetic ancestry from Bursuceni carried Y haplogroup J2b2b2~ (J-Z42942). Subineage J2b2* (J-M241) shows up at low diversity in the Balkans during the Neolithic; its origins remain uncertain[137].

Three of the CTAC specimens carried derivatives of the I2a1b Y haplogroup. Both CTAC specimens from Moldova carried the I2a1b1a2a2a lineage and one specimen from Verteba Cave in Ukraine carried the I2a1b1a2b lineage. I2a1b is part of the Y lineage diversity of the European Mesolithic and Neolithic, but it is not found in post-Neolithic samples from Europe, except for the Orkney Islands of Scotland [139]. It is possible that the group became part of the European Neolithic ancestry as some of the European Mesolithic hunter-gatherers became integrated into the ANF-derived EEF communities [139,140].

Among the specimens in our selection, a CTAC individual from Verteba Cave and a proto-Usatove individual from Mayaky carried E1b1b1a1b* Y chromosomal lineages. The proto-Usatove designation of the Mayaky individual is based on the $^{14}$C date that predates Usatove even after adjusting for reservoir effect[5] (Section 1.2.1). The E1b1b1a1b lineage is not associated with the Balkan-Danubian route of farming expansion from Anatolia to central Europe in the Neolithic[137]. On the other hand, the E1b1b1a1b~ expansion in the Mediterranean, which is timed to the late Mesolithic in southeast Europe [137], raises a possibility that it coincided with the spread of agriculture out of the Fertile Crescent into southern Europe via the Mediterranean coast beginning ca. 9000 years ago [141]. Such as, Y chromosomal lineage diversity in the Epicardial culture of the Cardial Ware complex of the Neolithic Mediterranean shows Near Eastern influence and Cardial/Epicardial mtDNA lineage frequencies and diversity are comparable to those from Near Eastern Pre-Pottery Neolithic B sites[142]. A bearer of the E1b1b1a1b1 haplogroup belonging to the Cardial Ware archaeological complex was reported from the Croatian Zemunica Cave in western Balkans [8]. Cardial Ware complex is considered to have influenced the Late Neolithic-Eneolithic Hamangia culture of the west-northwest Pontic [143]. Hamangia, in turn, is considered to have influenced the formation of Precucuteni-Trypillia A [144]. The presence of the E1b1b1a1b1 lineage in Trypillia and a proto-Usaove individual strengthens the link between Hamangia and CTAC as well as CTAC and Usatove and connects the genetic ancestry of CTAC and Usatove with Cardial Ware.

Lineages of the Y chromosome in two individuals from Yasynyvatka (Neolithic) and Taraclia (ZV) belong to the Q1b clade. Q1b1 has a general Eurasian distribution. Q1b2b1b2b (Taraclia) has been identified in his second cousin, a steppe Maykop individual from Sharakhalsun (SA6004, Q1b2b1b2b~).

The J-Y6313-deived J-FT265222 lineage of Y haplogroup J1 identified in a genetically Usatove individual from the Revova kurgan (Burial 19) is present in modern populations of

Europe, as well as Saudi Arabia and Iraq. Another Usatove male from Mayaky carried the R1a lineage, has a widespread Eurasian distribution, but its initial diversification is thought to have started in Iran[145].

#### 3. Archaeogenetic notes on reported individuals

Here we present some notes on selected individuals of particular archaeological interest according to the methodology of Supplementary Information, section 2.

##### 3.1. Trypillia outliers

Both Trypillia specimens from Moldova (Dănceni, I20069, and Soldanești, I7920) are placed between the Ukraine Trypillia and Usatove clusters on the PCA. Individual I20069 (3323-2935 calBCE) has a significant amount of CLV ancestry and is composed of ~1/4 PVgroup and ~3/4 Trypillia ancestry (using the remaining 27 Trypillians as a source). The individual can also be modeled as a mixture of 53.7±5.5% Usatove and Trypillia (as Usatove itself is a mixture of PVgroup and Trypillia, p=0.161).

##### 3.2. Genetic ancestry of Zhyovotylivka-Volchans'k/III-C burial type (ZV/III-C) individuals

Four individuals in our database come from burials archeologically classified as belonging the ZV/III-C burial type. Three of them came from the quadruple burial at Bursuceni and one, I20078, came from a double burial at Taraclia II Kurgan 2. All four burials are contemporaneous and date to the chronological interval of 3350-3030 calBCE. A fifth individual (I20079) from a ZV/III-C type burial from Moldova is dated to the middle 3$^{rd}$ millennium BCE and is a Yamna descendant (Table 1).

Individual I20078 from Taraclia can be modeled as Core Yamna Steppe Maykop ancestry (Supplementary Information, section 2). The IBD analysis uncovered that Individual I20078 is a second cousin of Steppe Maykop specimen SA6004 from Sharakhalsun Kurgan 2 burial 18[146]. The two individuals have four IBD blocks longer than 20 cM, one of which is 45.9 cM[51]. The I20078- SA6004 kinship provides genetic support for the link between the communities of the ZV/III-C burial type from the North Pontic and those from the Kumo-Manych periphery of the Maykop-Novosvobodnaya domain[57].

Individual I17974 (one of the female twins from Bursuceni) has much lower amount of Steppe Maykop-related ancestry than the I20078 individual. A parsimonious interpretation of the modeling presented in Supplementary Information, section 2 suggests that this individual, like I20078, was formed of the same Yamna+Steppe Maykop admixture process but with about ~1/3 of the Steppe Maykop ancestry found in I20078.

Individual 17973 (the male adolescent from Bursuiceni) has ancestry from the Caucasus based on the position in PCA. Modeling suggests that over 90% of ancestry of this individual comes from an Aknashen-derived source with an admixture of steppe ancestry, similar to that of the Maykop population, but with a higher proportion of the south Caucasus ancestry compared to that of Maykop.

##### 3.3. Catacomb outliers

Individual I11850 (ODK1, 2874-2630 calBCE, the Odesa Kurgan) and I12840 (DBN1.10, Dubynove Kurgan 1 Burial 10), both archaeologically Catacomb, cluster with the Yamna main cluster. Individuals I12621 (DBN1.9, contaminated) and I17742 (MK3.15, Moldova, Yamna archeologically) occupy positions on the PCA next to the main Yamna cluster, but towards the Caucasus. Both score low on SNP hits (9788 and 3980, respectively). I13072 (SL-1, Catacomb archaeologically, 4001 SNP hits, contaminated) is on the other side of the main Yamna cluster, near Serednii Stih specimens. All successful models for the Individual I11850 include some CLV Cline ancestry.

The Catacomb Individual I16668 (REV3.10) stands alone on the PCA, towards the main Usatove cluster. Its resolution is likely affected by a low SNP hit score (15071), but the PCA position is interesting considering the main burial in Revove Kurgan 3 genetically identified as part of the Usatove population. Here an individual retaining the genetic ancestry of a previous archaeological group is buried in the cultural tradition of that chronological period. Or, perhaps, this has something to do with burying genetically related individuals in the same kurgan, or some local groups retaining more archaic genetic features than others.

Individual I13071 (2201-2032 calBCE) from Bil'shivtsi is another Catacomb outlier and is discussed in the Corded Ware Connection section.

##### 3.4. A Corded Ware connection

Within our sample selection, YAC individual I12005 from northwest Pontic (Soldats'ka Slava Burial 10) and an individual from a catacomb burial with an ambiguous cultural placement from the Podillya region in west Ukraine (Bil'shyvtsi Burial 1, I13071, 2201-2032 calBCE) demonstrate potential connections to Corded Ware. Their mtDNA lineages belong to distinct sublineages of the T2c1 clade, likely of steppe origin. On the PCA (Figure 2a), they cluster together, with Individual I27983 (Yasinyvatka burial 24) positioned in close proximity. In PCA iterations involving central European Bronze Age specimens, this location is populated by central European Únětice and Corded Ware. While I27983's contamination raises concerns about its use in comparative analysis, its proximity to the central European Bronze Age is noteworthy. Individual I13071 fits multiple models, most of which suggest a significant North Caucasus genetic ancestry. When modeled with Core Yamna and EEF ancestry, approximately a third of I13071's ancestry appears to originate from EEF (Supplementary Information, section 2).

##### 3.5. Hunter-gatherer genetic ancestry resurgence in Babyne/Multi-Cordoned Ware

Modeling shows that both the Yamna of Ukraine of the EBA and the later Multi-Cordoned Ware of the MBA were largely descended from the Core Yamna but the latter experienced gene flow from a population that seems to have had more hunter-gatherer ancestry.

It was previously observed that Bronze Age populations from neighboring Romania included those that had a high proportion of hunter-gatherer ancestry[147]. These individuals, together with those of Ukraine both point to the existence of Chalcolithic populations of high hunter-gatherer ancestry that bequeathed it to those of the Bronze Age. A previously reported

Chalcolithic individual from Gura Baciului of >50% hunter-gatherer ancestry[147,148] provides direct evidence for the existence of such populations during the Chalcolithic.

#### 4. Strontium isotope analysis of the Neolithic Mariupol Necropolis

Two individuals from the sample selection reported in this study came from the Neolithic Mariupol Necropolis. One of the specimens, ANB002 (I31731) came from the Neolithic Burial 50 (double burial of an adult and a child (sampled)). Another specimen, ANB003 (I31732) came from Burial 21 (from one of the two skeletons in this tripe burial, dated to the Eneolithic period). These were excavated by M. Makarenko in 1930 prior to the construction of the Azovstal teel plant and subsequently transferred to the Museum of Local History of Mariupol[1], where they remained until the Russian invasion of Ukraine in 2022. The third sample, ANB005, came from the Neolithic layer (257 $m^2$ of total excavated area) of the Kalmius settlement in the vicinity of the necropolis[149]. The Kalmius settlement archaeological site was discovered in 2010. Excavations were carried out by the Archaeological Expedition of the Mariupol Museum of Local History under the direction of V.M. Gorbov in 2010-2012, 2014, and 2015. The settlement is considered to be associated with the necropolis.

The Museum of Local History of Mariupol and its collections including the excavated artifacts from the Neolithic Mariupol Necropolis, as well as the Kalmius settlement site, were destroyed by the Russian bombardment during the siege of Mariupol in the spring of 2022[150].

Samples of bone and teeth for the current analysis were collected in the fall of 2021. The ANB002 and ANB003 samples, to our knowledge, are the only human biological artefacts from the Mariupol Necropolis that survived the Russian invasion. In addition, these two samples have not been contaminated by heavy metals in the result of modern warfare (WWII, 1941-45, and the Russian invasion of 2022) as well as environmental pollution caused by the operation of the Azovstal steel plant.

The tooth enamel and bone samples were prepared and analyzed at the University of Missouri Research Reactor (MURR). The samples were mechanically cleaned using a microdrill equipped with a bristle brush and the bone samples further cleaned with a diamond bit. A small fraction of the samples was taken for analysis. In the case of the tooth, a small corner was cut using a diamond disk mounted on a Dremel. All materials were rinsed with mQ water. The bone samples were soaked in a solution of 2% NaOCl for eight hours and rinsed three times with mQ water, then soaked in 0.1N acetic acid for 30 minutes, rinsed five times with mQ water, soaked in 0.1N acetic acid for about seven hours, rinsed five times with mQ, and the last two steps were repeated once more. The leaching cycles for the enamel were shorter and as follows: one hour in NaOCl, then two hours and one hour in 0.1N acetic acid. The number of rinsing cycles was similar. All samples were then dried at room temperature. The remaining weights were 4.4 mg, 9.9 mg and 2.3 mg for ANB002, ANB003 and ANB005, respectively. The three samples were dissolved in 2ml of 7N $HNO_3$ in a PFA vial at 110˚C for 24 hours. The dissolution was followed by an evaporation cycle at 90˚C. The dry residues were re-digested in 7N $HNO_3$ prior to the Sr separation using a protocol adapted from[151]. The Sr eluates were evaporated at 90˚C, and the residue dissolved in 0.05N $HNO_3$ before conducting the strontium isotopic analysis following the procedure described in[16]. The average and standard deviation calculated for the SRM987 were 0.71022 ±0.00003 (2sd) (n=16).

The enamel sample (ANB002) has a $^{87}Sr/^{86}Sr$ ratio of 0.70950 and the bone samples have a $^{87}Sr/^{86}Sr$ ratio of 0.70990 and 0.70993, for ANB003 and ANB005 respectively (Table S2).

The strontium isotopic signature of the Mariupol samples was compared with values from[152], which provide estimated ranges of $^{87}Sr/^{86}Sr$ ratios for the main geological ages in Ukraine. The site of Mariupol is located in a geological context that corresponds mainly to Miocene and Pliocene sediments and sedimentary rocks with a $^{87}Sr/^{86}Sr$ ratio estimated between 0.709 and 0.711[152] and a diversity of Precambrian metamorphic rocks with a $^{87}Sr/^{86}Sr$ ratio estimated between 0.712 and 0.780[152]. An overview of the lithology of Ukraine is presented in Fig. S2.

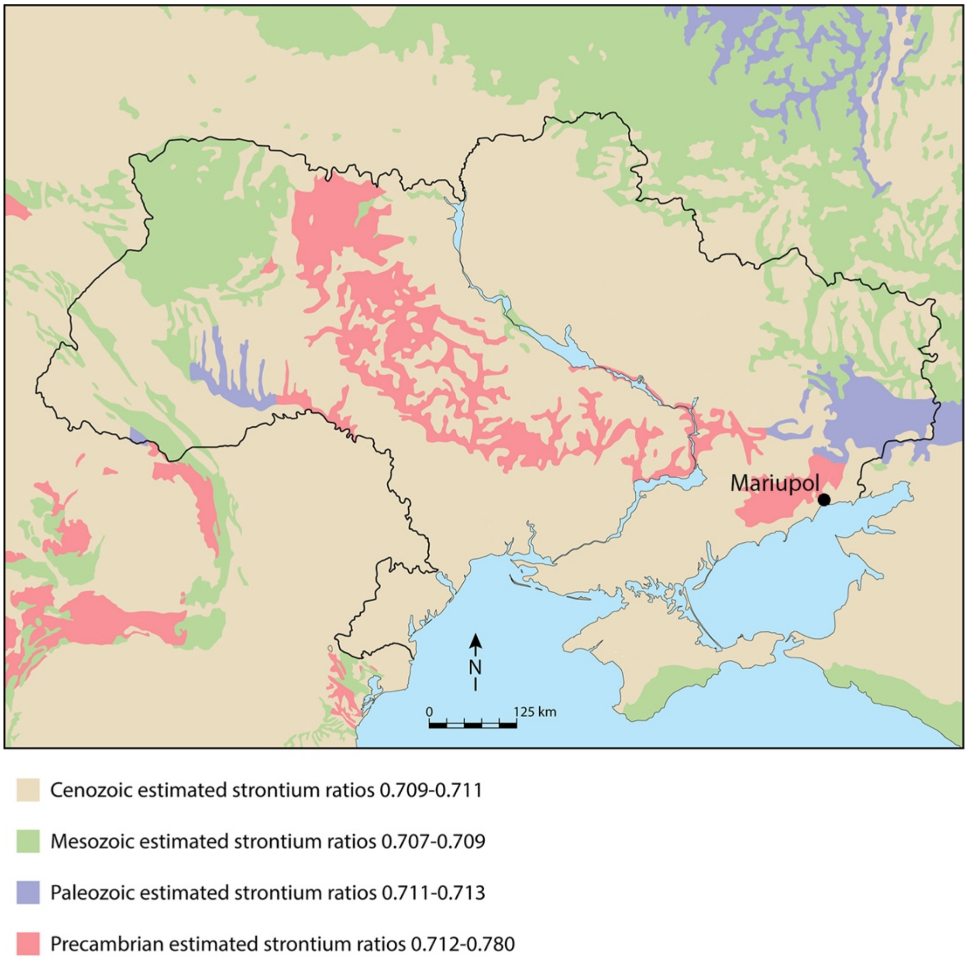

**Fig. S2**. Simplified geological map of Ukraine (modified after[152]) and the location of Mariupol.

All three samples have a signature that is compatible with the range of expected local $^{87}Sr/^{86}Sr$ ratio. These substrates are, however, largely distributed and the same range of 87Sr/86Sr ratio can be expected for a large portion of Ukraine. The $^{87}Sr/^{86}Sr$ ratio of seawater (0.7092) falls within the range estimated for Miocene and Pliocene sediments and sedimentary rocks.

The samples were compared in more detail with data available for local baselines and human remains for a number of sites in Ukraine and published in[7,152,153]. We use the local baseline ranges as defined by[152]. The results are presented in Fig. S3 and Table S2.

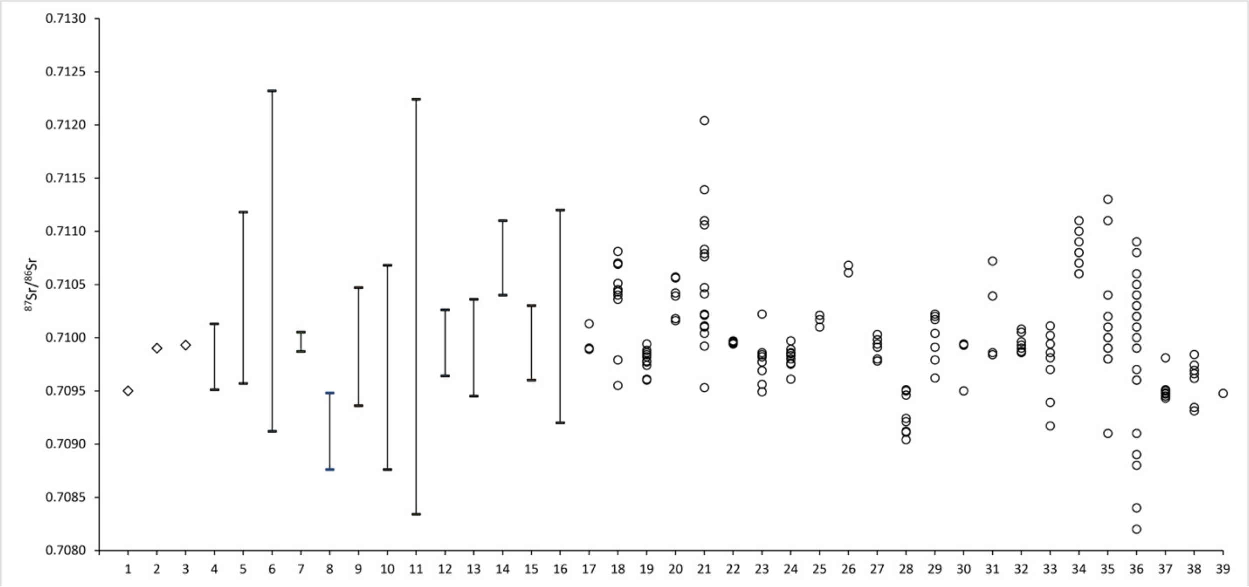

**Fig. S3**. Comparison of the $^{87}Sr/^{86}Sr$ ratio of the three samples from Mariupol with ratios for baselines and human remains (including enamel, dentin, and bone samples) from other sites in Ukraine and Olenii (Russia). The baseline ranges are reported as defined in[152]. The values for human remains are from[7,152,153] and are reported by site and by culture. Data used to construct the graph are presented in Table S4.4.
Samples from Mariupol: 1. ANB002, 2. ANB003, 3. ANB005. Baselines: 4. Vynohradne, 5. Pishtchanka, 6. Kirovohrad, 7. Shakhta Stepna, 8. Nevs'ke, 9. Alexandropol, 10. Babyna Mohyla, 11. Drana Kokhta, 12. Ordzhonikidze, 13. Zolota Balka, 14. Medvin, 15. Mamai-Gora, 16. Bel'sk. Human remains (including enamel, dentine and bone): 17. Vynohradne - Eneolithic, 18. Pishtchanka - Eneolithic, 19. Vynohradne - Yamna, 20. Pishtchanka - Yamna, 21. Kirovohrad - Yamna, 22. Shakhta Stepna - Yamna, 23. Vynohradne - Early Catacomb, 24. Vynohradne - Catacomb, 25. Pishtchanka - Catacomb, 26. Kirovohrad - Catacomb, 27. Shakhta Stepna - Catacomb, 28. Nevs'ke - Catacomb, 29. Alexandropol - Iron Age, 30. Babyna Mohyla - Iron Age, 31. Drana Kokhta - Iron Age, 32. Ordzhonikidze - Iron Age, 33. Zolota Balka - Iron Age, 34. Medvin - Iron Age, 35. Mamai-Gora - Iron Age, 36. Bel'sk - Iron Age, 37. Olenii - Eneolithic and/or Bronze Age, 38. Verteba Cave - Eneolithic, 39. Verteba Cave - Iron Age.

The $^{87}Sr/^{86}Sr$ ratio of sample ANB002, 0.70950 ±1 (2se), is within the local baseline ranges of the sites of Kirovograd, Babyna Mogyla, Drana Kokhta, Zolota Balka and Bel'sk, and close to that of Vynohradne and Nevs'ke. It is different from the local baseline ranges for the sites of Pishtchanka, Shakhta Stepna, Alexandropol, Ordzhonikidze, Medvin, and Mamai-Gora.

The $^{87}Sr/^{86}Sr$ ratios of samples ANB003 and ANB005, 0.70990 ±1 (2se) and 0.70993 ±1 (2se), are within the local baseline ranges of the sites of Vynohradne, Pishtchanka, Kirovograd, Shakhta Stepna, Alexandropol, Babyna Mogyla, Drana Kokhta, Ordzhonikidze, Zolota Balka, Mamai-Gora, and Bel'sk. It is different from the local baseline ranges for the sites of Nevs'ke and Medvin.

**Table S2**. Strontium isotope ratios for samples from the Mariupol Necropolis and the Kalmius Settlement.

| **Lab ID material** | **Sample ID** | **Sample Description** | **$^{87}Sr/^{86}Sr$** | **2*se*** |
|---|---|---|---|---|
| ANB002 (I31731) Tooth enamel | I31731, Mariupol Necropolis V2 | Neolithic double Burial 50 (immature), permanent maxillary left incisor | 0.70950 | 0.00001 |
| ANB003 (I31732) bone | I31732, Mariupol Necropolis V3 | Eneolithic triple Burial 21 (female) | 0.70990 | 0.00001 |
| ANB005 bone | Kalmius Neolithic Settlement V5 | Burial K2 (immature) | 0.70993 | 0.00001 |

The individuals presenting the $^{87}Sr/^{86}Sr$ ratio the closest to sample ANB002 (I31731) are one from Verteba Cave (Scythian), one from Kirovograd (Yamna), one from Vynohradne (Early Catacomb), two from Nevs'ke (Developed Catacomb culture), one from Babyna Mogyla (Scythian) and six from Olennii (Eneolithic and/or Bronze Age).

The individuals presenting the $^{87}Sr/^{86}Sr$ ratio the closest to both samples ANB003 and ANB005 are one from Vynohradne (Eneolithic), one from Kirovograd (Yamna), one from Shakhta Stepna (Catacomb), seven from Mamai-Gora (Iron Age), two from Bel'sk (Iron Age), one from Olexandropol (Scythian), two from Babyna Mogyla (Scythian), and two from Ordzhonikidze (Scythian).

Sample ANB005 is also similar to three individuals from Shakhta Stepna (two from the Yamnaya culture and one from the Catacomb culture), one from Vynohradne (Yamna), one from Babyna Mogyla (Scythian), one from Ordzhonikidze (Scythian) and one from Zolota Balka (Scythian).

Sample ANB003 (I31732) is also similar to three individuals from Vynohradne (one from the Eneolithic, one from the Yamnaya culture, and one from the Developed Catacomb culture) and two from Ordzhonikidze (Scythian).

In addition to Sr isotope ratios, stable isotopes ratios of carbon and oxygen were obtained on tooth enamel of specimen ANB002. The obtained values are as follows: $\delta^{13}C$: -10.4 ‰, IRMS $\delta^{18}O$: -8.7 ‰ (IRMS). Since the radiocarbon date obtained on enamel carbonate of I31731 indicates possible contamination with soil carbonates (Table S1), it is not clear how diagnostic are the $\delta^{13}C$ and $\delta^{18}O$ measurements from the same enamel sample.

In conclusion, strontium ratios of all three individuals from Table S2 have Sr isotope ratio signature compatible with that of the environment in which they were buried, but this signature is also compatible with a large portion of Ukraine and that of the baselines and human remain samples from multiple sites in Ukraine as well as proximate to the Sea of Azov sites outside of Ukraine.

#### 5. Burial descriptions

### Bulgaria

(Contact: D. Reich)

##### *Durankulak Kurgan F*

Durankulak, Tolbukhin district, Southern Dobruja Region (43.693, 28.527)
The cemetery of Durankulak is the largest Late Neolithic-Early Bronze Age cemetery in the Pontic region, containing 1204 burials. In 1980, 39 prehistoric graves (burials 10-49), including complexes with remains of a funeral meal, were uncovered on the border between loess and black earth[25]. One of these complexes was kurgan F, containing burials 12, 15, and 29. Kurgan F was one of seven kurgans at the Durankulak cemetery. The 17 burials from the Durankulak kurgans document the earliest presence in Southern Dobruja of a population carrying the traditions of the Usatove, Proto-Yamna, and Cernavodă I populations. These individuals represent the population that came to Dobruja from the North Pontic steppe after 4200 BCE and filled the prolonged cultural vacuum after the collapse of the Eneolithic cultural system of Varna- Kodjadermen-Gumelnița-Karanovo (KGK) VI [154].

### I1456

DUR1, Kurgan F, Burial 15 (main burial), male, adult, 3700-3300 BCE
Contracted on the right side, head to north, facing east.

##### *Golyamata Mogila (Popovo)*

Popovo, Bolyarovo Municipality, Yambol Region (42.204, 26.728)

Archaeological rescue excavations of the Big Popovo kurgan 4 km south of the village of Popovo took place in 2006 [29]. The kurgan had the shape of a segment of a sphere. Thirty nine graves were discovered in the kurgan, 12 which (graves 10, 12, 18, 28, 29, 30, 31, 33, 36, 37, 38 and an empty burial pit 39) dated to the second phase of the Early Bronze Age. Four of them belonged to the Yamna complex (graves 28, 36, 37, 39). The dead were laid in the middle of a deep burial pit along its length so that the orientation of the pit was the same as the orientation of the skeletons. The skeletons were oriented in most cases in the east-west direction with the head to the west. The dead were laid on their backs, head slightly bent forward, and legs bent at the knees. The arms were extended down to the body. Ocher was found most around the skull. Four of the pits had a wooden cover. On the short walls of the pits, three or four thinner beams (with a circular cross-section) were placed transversely. Six or seven flat beams were placed on top of them, longitudinally of the burial pit.

Pit 39 and grave 37 are the only structures in the kurgan that were dug into the matrix. It is possible that these pits were built together and were in some kind of relationship. After the completion of the funerary rites, the primary mound embankment was poured over them.

The remaining graves from the Early Bronze Age were dug into the already existing mound embankment. These graves lack a wooden covering. The skeletons were oriented in most cases in the east-west direction with the head to the west. The positions in which the dead were buried were supine, head slightly bent forward, and legs bent at the knees, which later fell in different directions. The hands were placed on the pelvis. Red ocher was found on the skeletons, with the skull, feet, and wrists being the most heavily colored. Pieces of red ocher were mostly placed near the head and on the bottom of the burial pit.

The inventory in the graves consisted of hair spirals made of silver and gold. The spirals found in the graves from Popovo find parallels among grave goods from present-day Bulgaria, Serbia, Romania, Moldova and Ukraine, all dated to the second phase of the Early Bronze Age.

### I1448

POP1, Golyamata Mogila, Burial 29, female, 3050-2550 BCE

### I1427

POP4, Golyamata Mogila, Burial 37, male, 3050-2550 BCE

Grave 37 was located in the NE sector of the mound, dug into the mainland. It was oriented NW-SE. A badly crushed skull was found in it, missing the upper jaw and the entire face. A humerus without epiphyses containing traces of ocher was discovered at the SE under the skull. Eight stake holes were found along the inner periphery of the burial pit. The presence of these holes suggests the possibility that the pit had another wooden structure before being covered with wooden beams and filled in.

##### *Riltsi Kurgan 264*

Dobrich, Southern Dobruja Region (43.592, 27.779)

The Riltsi necropolis consists of more than 30 kurgans arranged in a semi-circle in the west suburb of Dobrich. The kurgans are one to five meters in height. Rescue excavations of Kurgan 264 in 2000 uncovered five graves, archaeologically dated to the Early Bronze Age. Some of the graves studied in the Dobruja region showed parallels with the Bursuceni-Zhyvotylivka type burials north of the Danube River[30].

### I1428

RIL3, Riltsi Kurgan 264, Burial 5, female, 3360-2890 BCE

Excavated by S. Alexandrov in 2000. Semi-supine inhumation to the right, oriented southwest-northeast. Red ocher over the skull and pelvis. Burial inventory included a bovine bone placed near the skull. The burial pit had no covering. The burial is archaeologically attributed to the second stage of the Yamna archaeological complex [155]. Radiocarbon date estimate is based on dates from Riltsi Kurgan 264, Burial 4 (3361-3099 calBCE (4520±34 BP, SUERC- 86512)) and Kurgan 260, Burial 1 (3011-2887 calBCE (4314±26 BP, SUERC- 84759))[156].

#### Moldova

(Contact: H. Shephard, S. Łukasik, G. Sibru, V. Dergachev)

##### *Bursuceni Kurgan 1*

Bursuceni, Sîngerei district (47.488, 28.016)

In 1977, the Eneolithic Archaeological Expedition of the Institute of History of the Academy of Sciences of Moldova directed by E. V. Yarovoy conducted research on a single kurgan on the west bank of the Middle Prut River, on the route of a dirt road under construction near the village of Bursuceni. The southern half of the mound fell under the construction of the roadway.

The kurgan was located on the route of the local road Koshkodeny-Sloboda Megura, 400 m west–southwest of the outskirts of the village of Bursuceni. It was situated in the center of a valley, almost in the middle of a medium-sized upland, the width of which reached 2 km. Within a few kilometers, the mound was clearly visible from anywhere in the valley.

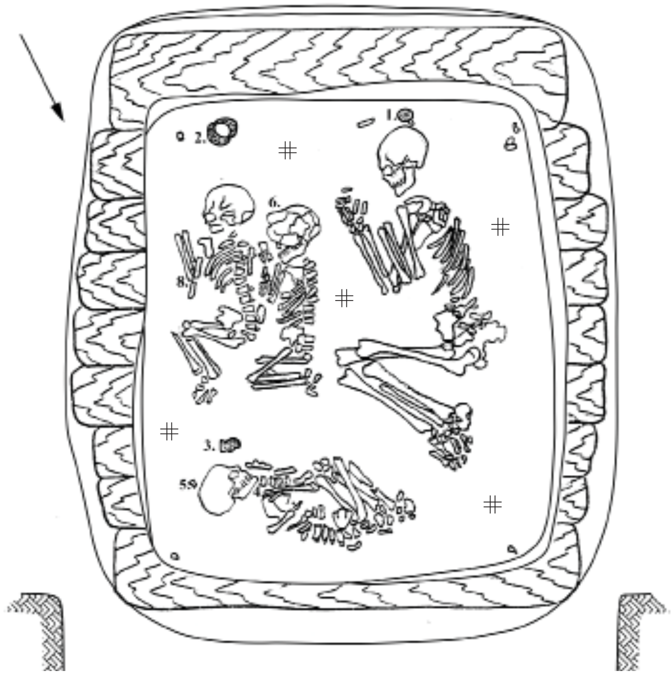

*Bursuceni Kurgan 1 burial 21. Image from* [23].

### I17973

Br/K1/B21/S1

Bursuceni Kurgan 1 Burial 21, Skeleton 1, male (Individual 5 according to the field numeration), 3354-3103 calBCE (4516±13 BP) [R_combine: (4520±25 BP, PSUAMS-8310); (4530±20 BP, PSUAMS-8723); (4500±20 BP, PSUAMS-8722)]

### I17974

Br/K1/B21/S2, Bursuceni Kurgan 1 Burial 21, Skeleton 2, female (Individual 6 according to the field numeration), 3334-3030 calBCE (4470±20 BP, PSUAMS-8724)

### I17975 (S20195)

Br/K1/B21/S3, Bursuceni Kurgan 1 burial 21, Skeleton 3, female (Individual 8 according to the field numeration), 3334-3030 calBCE (4470±20 BP, PSUAMS-8724).

Burial 21 was located in the center of the kurgan, at a depth of 3.9 m from the ground and 3.7 m from the surface of the mound. The burial chamber was of irregular rectangular shape with uneven walls and rounded corners, 1.75 m long, 1.4–1.55 m wide, and 0.6 m deep from the ledge, oriented along the line north-north-east - south-south-west. Its southern and eastern walls were sloping to the bottom by 0.05–0.08 m. At the level of the ledge, the pit had a transverse wooden ceiling of nine oak logs with a diameter of 0.18–0.2 to 0.35 m and a length of 1.65 to 1.95 m.

The burial contained skeletons of four individuals, an adult, and three children, all in excellent state of preservation.

The skeleton of an adult rested at the western wall of the burial chamber, contracted on the right side, head to the south, face to the east. The arms were sharply bent, with the hands placed at the front of the skull. The legs were sharply bent.

The skeleton of a child (I17974, Individual #6), 2–3 years old, rested in front of the skeleton of the adult, between the adult and another child, contracted on the right side, head to the south-southwest, facing east. The arms were sharply bent, with the hands placed at the front of the skull. The legs were sharply bent.

The skeleton of another child (I17975, Individual #8), 3–4 years old, was laid near the eastern wall of the pit, with the back close to the first child, contracted on the right side with a slight inclination to the chest, head to the south-southwest, facing southeast. The arms were sharply bent, with the hands placed at the front of the skull. The legs were moderately bent.

The skeleton of the third child, 7–8 years old (I17973, Individual #5), was located at the northern wall of the pit, at the feet of the adult and the first two children, contracted on the left side, head to the southeast, facing southwest. The arms were sharply bent, pressed to the chest, with the hands placed at the mandible. The legs were sharply bent, the knees were pressed to the elbows, heels to the pelvis.

All skeletons, especially the frontal parts of the skulls, were intensely colored with crimson ocher. In the southwestern corner of the pit, 0.25 m from the back of the head of an adult, there was a piece of crimson ocher 0.05 m in diameter.

Burial inventory included three clay vessels, a bone pin with a curved end and a pointed opposite end, and a 0.2 cm round hole in the central part (found under the mandible on the chest of the third child (individual #5), flint insert of a sickle (at the parietal part of the skull of the third child), beads made from shell valves (found on the remnants of the plant mat under the skeleton of the third child), gold temporal ring (under the right temple of the first child (individual #6)), and a flint flake without retouching (laid 0.05 m from the radius bones of the second child (individual #8)).

S17974 and S20195 were considered duplicates during DNA amplification and appeared genetically identical. Thus, individuals 6 and 8 (first and second child) are identical twins.

##### *Cioburciu Kurgan 4*

Cioburciu, Ștefan Vodă District (46.599, 29.722)

The Cioburciu kurgan group was located 7 km south of the Cioburciu village, 200 km west of the Cioburciu – Ștefan Vodă highway, on the edge of the plateau of the west bank of the Dniester, limited by a gully-beam system and stream tributaries of the Dniester. The group consisted of five kurgans and was stretched along a north-south line.

Kurgan 4, the largest in the group, 2.7 m high and 60 m in diameter, was located 50 m northwest of Kurgan 3 and 80 m north of Kurgan 2. Kurgan's embankment was subjected to intensive plowing. Thirty-eight burials, eight of them belonging to the Yamna and four belonging to the Catacomb archaeological complexes, were discovered in the kurgan[157].

### I20067

CiK4.3, Cioburciu, Kurgan 4, Burial 3, female, 3300-2500 BCE

Burial 3 (Yamna archaeological complex) was discovered in the southeastern sector of the mound, 2.1 m from the benchmark at a depth of 1.7 m. The contours of the burial chamber were not traced. The skeleton of an adult lay contracted on the back, tilted to the left, with the head to the south-southwest. The skull was facing left. The left arm was extended forward along the body, the humerus and radius bones were partially preserved, the right arm was slightly bent, and the hand rested on the pelvis. The legs were bent to the left, the degree of skeletal contraction was average. The skeleton was painted with bright red ocher, most intensely the skull. Dark brown decay was traced under the interred.

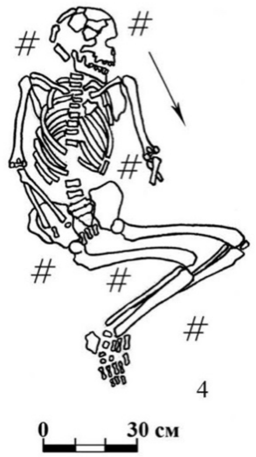

*Cioburciu Kurgan 4, Burial 3. Image from*[157].

##### *Cotiujeni Kurgan 1*

Cotiujeni, Briceni District (48.346, 26.938)

The kurgan chain of five kurgans near the village of Cotiujeni was studied in 1986 [158]. Kurgan 1 was the largest in the group. It was 44 m in diameter and 1.5 m in height. The kurgan contained eight burials.

### I20068

CoK1.6, Cotiujeni Kurgan 1 Burial 6, male, 2881-2671 calBCE (4175±20 BP, PSUAMS-10740).

Burial 6 (double burial, Yamna archaeological complex) was discovered 6.3 m west of the central benchmark at the depth of 1.67 m. The rectangular, 1.7 × 1 × 0.6 m burial chamber with rounded corners was oriented along the WNW-ESE line. Both skeletons were laid contracted on their backs, legs to the right, head to WNW. The skeletons were covered with red ocher, more intensely in the head and pelvic area.

##### *Crasnoe Kurgan 9*

Crasnoe, Grigoriopol District (47.131, 29.320)

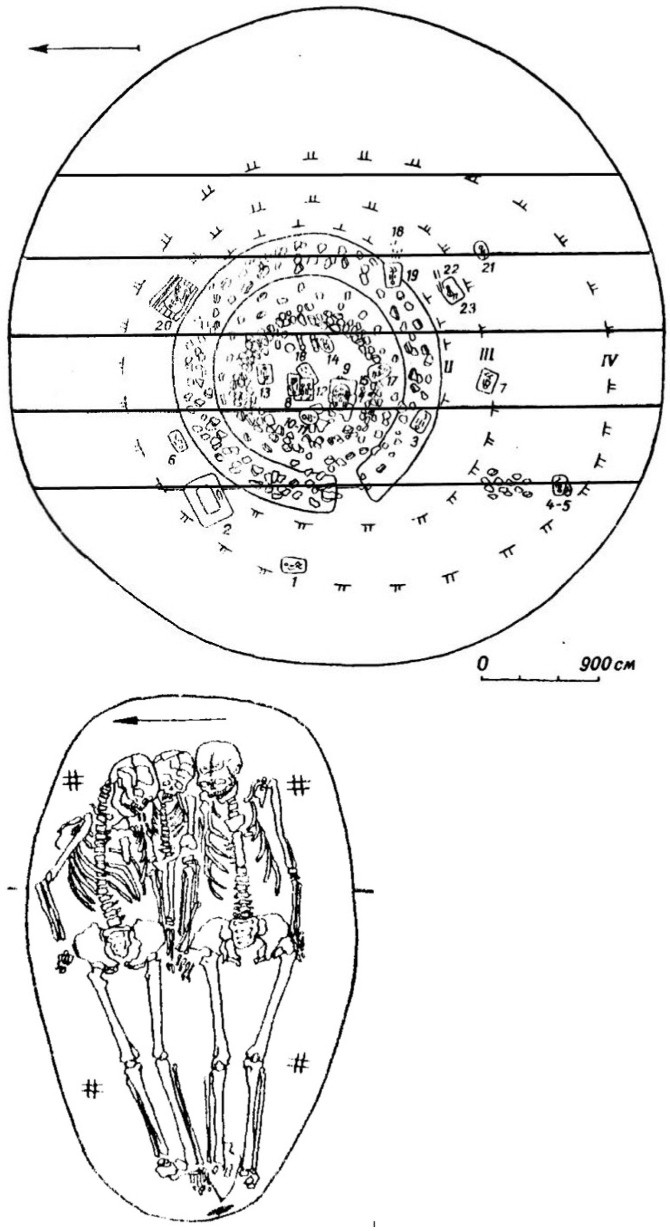

*General plan of the Crasnoe kurgan 9 and burial 9. Image from* [21].

Kurgan 9 was located on the west bank of the Lower Dniester near the village of Crasnoe, northeast of Grigoriopol, 2.5 km northeast of the Grigoriopol-Crasnaia Gorca highway. The kurgan was located on a plateau, on an arable field. It was the largest in the kurgan group. The northern foundation of the kurgan was steeper and 10 m shorter than the southern one. The preserved height of the mound from the buried soil was 4.8 m, from the mainland - 5.4 m, diameter - 60 m. The length of the northern part of the mound was 20 m, the southern part was 30 m in length. The kurgan contained 23 burials. The kurgan was investigated by E. V. Yarovoy in 1979 [21].

I20196

Kr/K9/B9/S2, Crasnoe Kurgan 9, Burial 9, Skeleton 2, female, 3352-3101 calBCE (4510±25 BP, PSUAMS-8835)

I17977

Kr/K9/B9/S2(d), Crasnoe Kurgan 9, Burial 9, Skeleton 3, female, 3350-3100 BCE

A triple Burial 9 (female, male and child) was found in a pit covered with three massive stone slabs in the southwestern sector of the mound at a depth of 3.4 m from the ground to the stone slab cover. On the western side of the grave cover there was a massive slab of irregular rectangular shape with an oval protrusion on one side, with clearly visible traces of processing. Placed on the side, the slab resembled a generalized image of a bull in profile.

The individuals were buried in extended supine position. All three skeletons were intensively painted with crimson ochre. The male skull has ornamented with bands of ocher. An animal shoulder blade was placed under the right foot of the male skeleton.

The skeleton of a woman (sampled for this report) laid in an extended position on her back with slight tilt to the left, head to the east-southeast. The skull was inclined towards the left shoulder, facing southwest. The left arm was straight, pressed against the body. The right arm was bent at a right angle, the hand was placed at the pelvis. The spine was bent to the left side - towards the child (sampled in this report).

Burial Inventory: a 7 × 2.4 × 0.6 cm adze made of local Dniester flint was found at the western wall of the pit in the filling 0.1 m west of the man's feet.

*Cunicea*

Prişanscaia Gora, Soldaneşti District (47.904, 28.674)

The cemetery and settlement at Cunicea-Prişanscaia Gora is a Cucuteni-Trypillia Vykhvatyntsi-type site (Trypillia CII chronological period). The site was excavated in 2012 [159].

I7920

12M1, Cunicea-Prişanscaia Gora, Section 4, Layer 3, Burial 4, male, adult, 3350-3099 calBCE (4505±25 BP, PSUAMS-4872)

Human remains included maxilla with teeth and long bone fragments.

*Dănceni II*

Ialoveni District (46.968, 28.717)

The late Trypillian ground cemetery at Dănceni II was located on the edge of a low (5-6 m) terrace above the floodplain on the east bank of the Ishnovets River, about 1.5 km NW from the village. Investigated in 1976 by I.A. Rafalovich and V.A. Dergachev. The burial ground consisted of four or five burials, which were organized in a semicircle with a radius of 6-7 m. To the east of the conditional center of the circle were two adult burials, to the south there was one child’s burial, and to the north, destroyed burials of one or two more children were located. The distance between the burials was 1.5-4 m.

The Dănceni site is considered poly-cultural by archaeologists. It reflects interpenetrating traditions of the late Trypillian Gordinești and Usatove cultural groups [160,161]. The site is located approximately halfway between the major Usatove culture sites in the northwest Pontic such as Mayaky and Usatove-Velykyj Kuyalnik, and the Gordinești sites of Cucuteni-Trypillia Archaeological Complex in the middle Dniester area such as Pocrovca and Prydnystryasnske.

### I20069

DIIGB301, Dănceni II Ground Burial 2 (301), male, 3323-2935 calBCE (4435±20 BP, PSUAMS-8832)

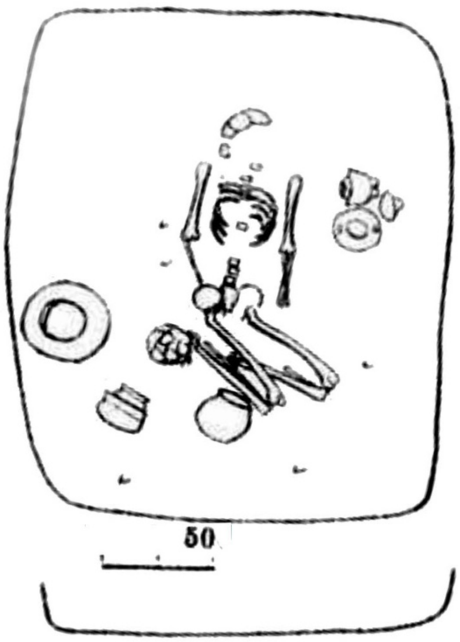

*Dănceni II Ground Burial Grave 2 (301). Image from*[186].

The rectangular burial chamber of Burial 2 (301) was located at a depth of 0.47 m. It was oriented SSE-NNW. The interred was laid in a contracted position on the back, knees turned to the left, head to N, arms extended along the body.

To the left of the interred, at the level of the elbow, there were two amphorae with handles-ears. One of the amphorae (H 10 cm) was equipped with four legs (broken off in antiquity) and covered with paintings. The second amphora (H 12.7 cm) was equipped with a lid with a conical ledge with a horizontal hole and two stop handles with vertical holes (H 6.6 cm). The amphora was apparently painted, but the painting has not been preserved. At the feet of the interred, closer to the southwestern corner of the burial pit, there were two bowls made of clay containing shattered burnt shards (chamotte). One of the bowls was ornamented with sickles, cord imprints, and paired knobs. The second bowl (H 10 cm) was decorated with double pinpricks and cord impressions filled with white paste. There was a jar (H 18.5 cm) near the knees of the interred. The upper part of the second similar vessel was found at the feet (H 14.4 cm). Another large vessel, decorated with "sickles" and cord impressions (H 31.4 cm), was located under the eastern wall of the burial chamber. Dispersed flint flakes and coals were found at the bottom of the chamber.

##### *Giurgiuleşti*

Cahul District (45.483, 28.2)

The Giurgiuleşti sanctuary and necropolis of the Suvorove-Novodanylivka archaeological group was located on the east bank of the Danube. Detailed descriptions of the Giurgiuleşti archaeological complex and associated finds, as well as burial descriptions, can be found in [24,162].

## I20072

GK2.6, Giurgiuleşti Kurgan 2 Burial 6, male, 4430-4058 calBCE, 5370±26 BP, MAMS-23175

Burial 6 (Gave 3 according to the burial sequence in[24]): a 2–3-year-old child buried in a catacomb with a round vertical shaft, narrow entrance into a chamber, and a domed chamber.

## I20073

GK2.11, Giurgiuleşti Kurgan 2 Burial 11, male, 20-25, 4484-4346 calBCE, 5571±32 BP, MAMS-28088

Burial 11 (Grave 4 according to the burial sequence in[24]): a 20–25-year-old male buried in a 5 m-deep pit, accompanied by weapons, including a composite sword and spear, jewelry, and sacrificial offerings.

#### *Glinoe (Hlinaia)*

Slobodzeya District (46.6797, 29.8099)

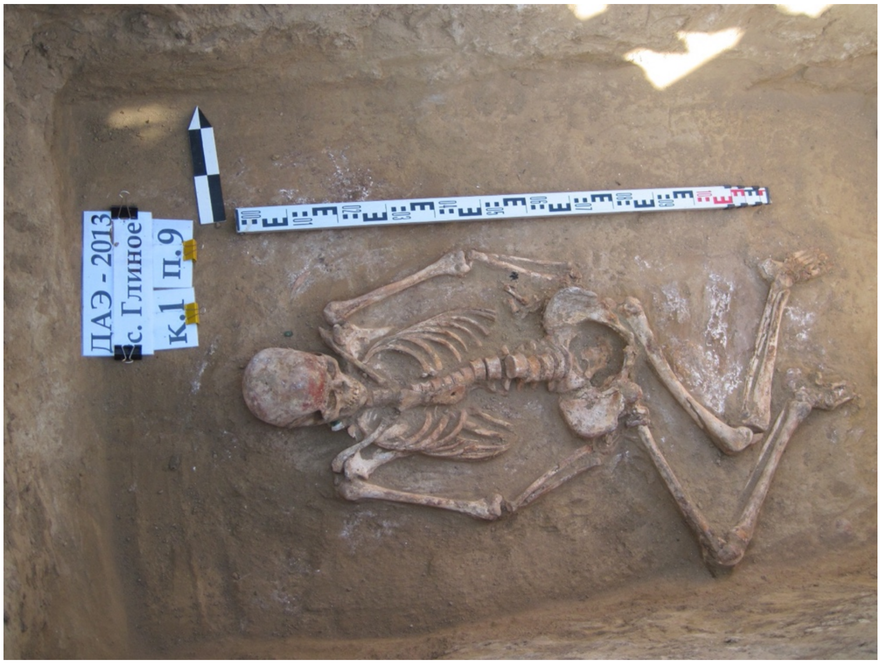

*I12507, DOT K1B9. Photo by V. Sinica.*

Two groups of kurgans near the village of Glinoe (Hlinaia) on the east bank of the Lower Dniester were investigated in 2013 by V.S. Sinica. The kurgan group "DOT" is located 1.82 km north-northeast of the northern outskirts of the village, adjacent to the Tiraspol-Dnestrovsk highway. The “SAD” kurgan group is located 2.6 km northeast of the northern outskirts of the village[163].

## I12507

DOT kurgan 1, burial 9, DOT K1B9, female, 2914-2882 calBCE (4275±20 BP, PSUAMS-10777)

Kurgan 1 was located on the slope of a floodplain terrace. On the southern side the height difference from the center to the bottom was about 1.3 m, and on the northern side - about 0.2-0.3 m. The kurgan was erected over Burial 18 of the Yamna archaeological complex. In total, 19 burials of different periods were recorded in the kurgan.

Burial 9 (Yamna archaeological complex) was made in a pit with a ledge. The buried was laid on the back, with the upper extremities along the body, the lower extremities flexed, and the head to the west. The skeletal remains were well-preserved and rather complete. Bones from all parts of the skeleton were present.  On the frontal part bone there was a strip of red ochre. The individual was estimated to be an adult, 35-50 years (age-at-death was estimated based on cranial suture closure and tooth wear). Morphological traits of the skull (frontal bone, mastoid process, supraorbital ridge, zygomatic bone, and gonial angle) indicated female sex, as confirmed by the aDNA results. Based on long bone measurements, the body height of the individual was 166 cm. The burial inventory consisted of five copper tubular piercings and a mollusk shell piercing as part of a necklace around the neck.

### I12512

SAD Kurgan 1, Burial 25, SAD K1B25, male (45-50), 2881-2633 calBCE (4165±25 BP, PSUAMS-10778)

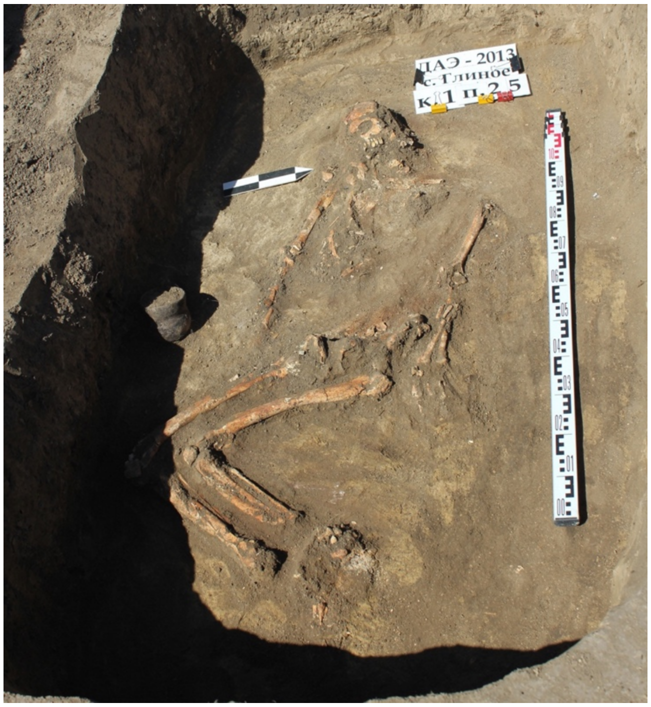

*I12512, Glinoe SAD K1B25. Photo by V. Sinica.*

Kurgan 1 of the “SAD” group was actually two kurgans with fused foundations. The northwestern kurgan (Kurgan 1A) was built in two stages. The first mound with a diameter of up to 19 m, consisting of dark gray chernozem, was built over the main burial 3 of the Yamna archaeological complex. At the time of excavations, the mound was heavily plowed, its maximum height above the ancient horizon was 0.8 m. Later, southeast of the first mound of Kurgan 1A, at a distance of about 20 m from its center, Kurgan 1B, with a diameter of up to 25 m, was built over the main Yamna burial 15, and the center of the mound was shifted to the northwest. In total, 27 burials from different periods were recorded in the mound.

Burial 25 was made in a pit. The interred was laid on the back, with the upper limbs along the body, the lower limbs flexed, and the head to the west. The skeletal remains were poorly preserved and incomplete. Fragments from all parts of the skeleton were present.

The individual was estimated to be an adult, 40-50 years (age-at-death was estimated based on tooth wear and pubic symphysis morphology). Morphological traits of the skull and pelvis (mastoid process, supraorbital ridge, orbits, and subpubic angle) indicated male sex, as confirmed by the aDNA results. Reconstruction of the body height of the individual was not possible owing to the insufficient degree of bone preservation. The burial inventory consisted of a molded goblet to the right side of the skeleton.

### I12637

SAD Kurgan 1, Burial 26, K1B26, male, 3300-2500 BCE

Burial 26 was made in a pit. The interred (non-adult) was laid on the back, with the upper extremities along the body, the lower extremities flexed, and the head to the west. The skeletal remains were moderately preserved and partially incomplete. Fragments from all parts of the skeleton were present. The individual was determined to be an infant, 0.5-1 years (age-at-death was estimated based on the dental development and eruption and long bone measurements). The male sex of the individual was genetically determined.
The burial was without an inventory.

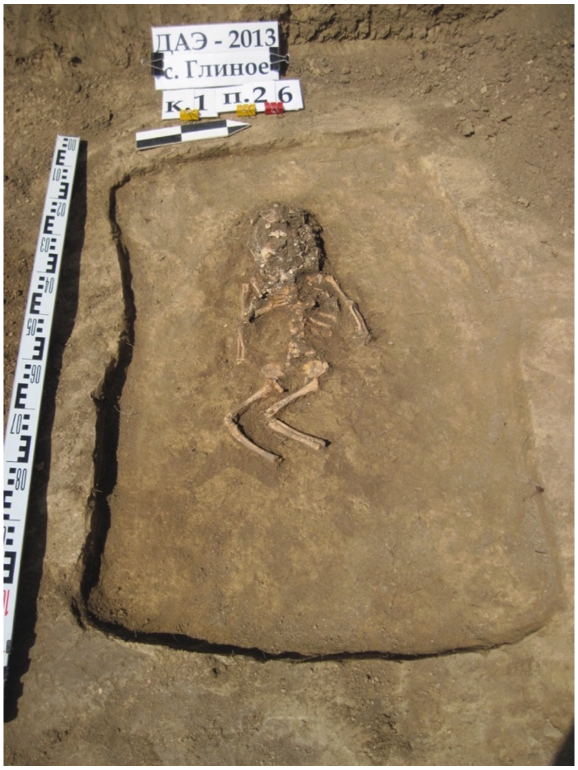

*I12637, Glinoe SAD K1B26. Photo by V. Sinica.*

##### *Mărculeşti Kurgan 3*

Mărculeşti, Florești District (47.869, 28.248)

Kurgan 3 was excavated in 1986 [164]. It was the largest of the kurgan group, with the diameter of 62 m and 3 m in height.

### I17742

MK3.15, Mărculeşti Kurgan 3 burial 15, female, 3300-2500 BCE

Burial 15 (Yamna archaeological complex) was discovered 13.5 m south of the central benchmark, at a depth of 4 m. The rectangular burial chamber 1-1.1 x 1.67 m was oriented along the ENE-WSW line. The chamber floor was covered with a plant mat. Brown-yellow ochre was observed under the skeleton. The interred was laid contracted on the back, legs to the left head to the ENE. The entire skeleton was covered in ochre.

*Mereni Kurgan 1*

Mereni, Anenii Noi district (46.677, 28.646)

Kurgan 1 was excavated in 1986-87. It was located on a high promontory approximately 3.2 km to the east of the Mereni village. The kurgan stood 1.2 m above the ancient surface and was about 35 m in diameter. A total of seven fill layers have been identified in the kurgan. The kurgan contained 16 burials[165].

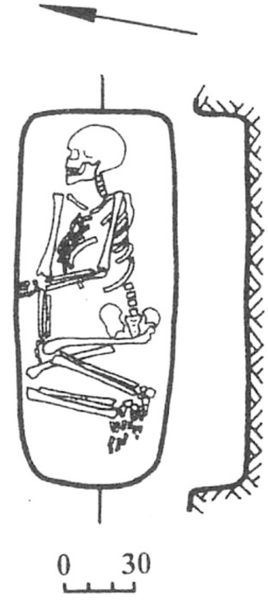

*Mereni Kurgan 1, Burial 15. Image from*[165].

I17743

MeII1.15, Mereni II Kurgan 1, Burial 15, female, 3358-3100 calBCE (4515±30 BP, PSUAMS-7822)

Burial 15 (Yamna archaeological complex) was discovered 2.70 m south-west of the marker at a depth of 2.1 m. The burial was located directly under burials 10 and 13. The burial pit was rectangular in shape (0.6 × 1.47 m) with the walls slightly narrowing towards the even bottom. The interred lay contracted on the right, skull towards northeast, facing north. The right hand was extended along the body with the wrist resting under the knees. The left hand was bent at the elbow. There was no inventory. The Mereni Kurgan 1 Burial 15 is the chronologically earliest burial archaeologically and genetically identified as Core Yamna in northwest NPR.

*Ocniţa Kurgan 1*

Ocniţa District, west bank of the Lower Dniester (48.385, 27.439)

I20076

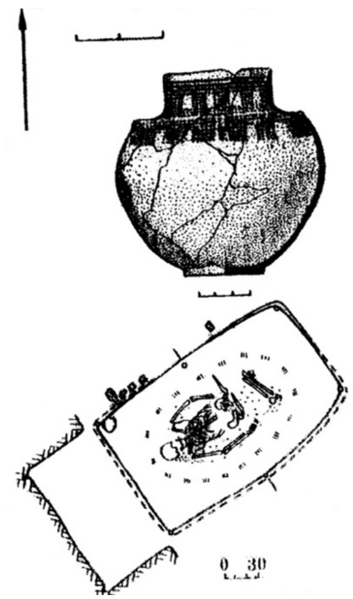

*Ocniţa Kurgan 1, Burial 3. Image from*[166].

OcK1.3, Ocniţa Kurgan 3, Burial 14, female, adult, 2906-2702 calBCE (4230±25 BP, PSUAMS-11947)

A kurgan group located 1.5 km southeast from the Ocniţa village was investigated in 1988 [166]. Kurgan 3, based on stratigraphic profiles, consisted of three overlapping mounds. The original mound was constructed over Burial 17 of the Yamna archaeological complex. Subsequently, Burial 14 of the Yamna archaeological complex was incorporated into the northern floor of the second mound. The remaining Yamna burials in the kurgan were situated within the third mound, forming a circle relative to the mound's center, where one group of five burials formed the inner circle, and a group of four burials formed the outer circle.

The burial structure of Burial 14 comprised a 0.9 m deep inlet pit and a rectangular burial chamber measuring 2.1 × 1.32 × 1.06 m, with a depth of 0.95 m. The chamber's northeastern part was oriented along the northeast-southeast direction. The fill consisted of mixed loose soil with remnants of a transverse wooden ceiling, part of which was preserved at the edge of the northwestern wall of the chamber. The adult interred in a flexed position on the back, head to the southwest at 240 degrees, face up, with arms slightly moved to the sides and left hand resting on the pelvis. Initially raised with knees up, the legs had fallen to the left. The bones were covered in red ocher, most intensely on the right calcaneus and humerus of the right hand. Dark brown decay from a mat interspersed with ocher was traced beneath the interred.

In the western corner of the chamber, a molded amphora of the Globular Amphora style measuring 24 cm in height, with a rim diameter of 12.5 cm, body diameter of 23.4 cm, and bottom diameter of 8.8 cm, was discovered. The vessel, made of a clay mix with ample sand and occasional chamotte grain inclusions, exhibited high-quality firing, with a dense, reddish shard. The outer and inner surfaces were carefully polished, with a gray-black color on the outer surface and a grey color on the inner surface. The vessel had a round-conical body with convex sides and a flat, well-defined bottom. The high and cylindrical neck featured a straight and rounded corolla, with a noticeable low ridge at the neckline and shoulders. Four small vertical handles with round horizontal holes adorned the shoulders, and the edges of these handles resembled low sides. The amphora displayed rich ornamentation, including four parallel horizontal lines of imprints resembling a twisted cord under the rim. Below, short horizontal sections of the same prints formed vertical fields, followed by rows of sickle-shaped depressions, mirroring a similar ornamental system located below the sides at the shoulders.

##### *Sărăteni Kurgan 1*

Sărăteni, Leova District (46.613, 28.465)

The Sărăteni kurgan was located approximately 0.9 km east of the southern periphery of the village of Sărăteni. The kurgan occupied the northernmost position within a group made of four kurgans arranged in a row on the edge of a plateau rising above the Prut valley. The mound of the kurgan was plowed annually, and, at the time of the research, it had an irregular circular configuration in plan. The kurgan was about 2 m high and 25 m in diameter. The kurgan contained 14 asynchronous burials.

#### I17744

SK1.1, Sărăteni Kurgan 1 Burial 1, male, 3300-2500 BCE

Burial 1 (collective, Yamna archaeological complex) was detected in the SW sector of the kurgan, 6.3 m from the center and at a depth of 2.42 m. The grave contained the remains of four individuals, two children and two mature adults. Burial descriptions are presented in[167].

#### I17745

SK1.9, Sărăteni Kurgan 1 Burial 9, male, 3300-2500 BCE

Burial 9 (late Yamna archaeological complex, Middle Bronze Age period) was discovered in the SE sector of the kurgan, at a distance of 7.6 m from the center and a depth of 3.4 m. The skeleton of a child, moderately twisted on the left side, oriented with the skull to the NE at 70$^{o}$, facing south. The arms, bent at the elbows at an angle of almost 90$^{o}$, were placed with forearms forward. The preserved leg was bent at a right angle to the body, and the knee at an angle of 180$^{o}$, so the tibia was parallel to the femur. The bones of the skeleton were moderately covered with ochre. At the bottom of the pit, in some places, remnants of a plant mat were discovered.

##### *Taraclia II Kurgans 2 and 10*

Taraclia, Căușeni District (46.569, 29.116)

A group of 20 kurgans was located on the watershed between the Yalpug and Lungutsa rivers and was a continuation of the kurgan group near the Kazaklia village. The kurgans within the group were located quite compactly in the form of a strip elongated in the meridional direction. In 1980 and 1982-1984, 19 kurgans were investigated by C. M. Agulnikov, I.V. Manzura and E.N. Sava. Burials of the Late Eneolithic and Late Trypillian chronological period were found in kurgans 2, 10 and 18 [168].

Kurgan 2 (H. 1.2 m, D. 36 m) was located at the southern tip of the kurgan group, 0.15 km south from the Taraclia-Novoselovka road. The kurgan contained 15 burials. The kurgan was built over burial 14 (Zhyvotylivka type). Subsequently, burials of the Yamna complex were deposited into it, in the result of which the original center of the mound shifted to the northwest.

Kurgan 10 (H. 9.9 m, D. 80 m) was the largest of the kurgan group. It was located in the southern part of the group. The kurgan contained 21 burials. The first mound (H. 2.7 m, D. 46 m) was built over burials 16 and 17 of the late Trypillian period. The original kurgan was surrounded by a ring ditch (47 m × 1 m × 0.6 m). During the Early Bronze Age, several burials of the Yamna archaeological complex were deposited into the original mound. Additional mounds built over these burials increased the size of the kurgan.

### I20078

TIIK2.14, Taraclia II Kurgan 2 Burial 14, male, 3340-3034 calBCE (4480±25 BP, PSUAMS-8833)

The double burial 14 in a square pit (1.7 m × 1.7 m × 1 m) was located 7.5 m to SE from the presumed center of the kurgan. Two skeletons of adult individuals rested in a contracted position on the left side, head to the NE (80°). Arms were bent, hands in front of the face. The bones of both skeletons were painted with crimson ochre, especially the skulls. At the bottom of the pit, dark brown decay from a plant mat was traced.

Inventory: A Trypillian vessel (H. 9.5 cm), decorated with a painted net ornament, stood in front of the face of the right (southern) skeleton (presumably the one sampled for this report).

### I20079

TIIK10.2, Taraclia II Kurgan 10 Burial 2, male, 2571-2355 calBCE (3965±20 BP, PSUAMS-8834)

Burial 2 was located outside of the kurgan ditch, 31 m to the south from the presumed center of the kurgan, at a depth of 9.9 m. The pit was rectangular in shape, oriented NNW-SSE (1.5 m × 0.65 m × 0.35 m). The poorly preserved skeleton of a child rested in a likely contracted position on the left side, with its head to the north.

Inventory: 1) A Trypillian amphora made of well-elutriated clay (H. 22 cm), painted with brown paint, stood at the feet. 2) A dark-gray goblet made of well-elutriated clay (H. 5.6 cm) laid on its side near the eastern wall, opposite the skeleton. 3) A bone hook-shaped pin with a hole (L. 3.5 cm) laid near the skull. 4) Cylindrical beads made of bone, a brownish mineral, and a shell (D. 0.5 cm), as well as a tubular copper thread, laid near the skull next to the pin. 5) Three beads made of a reddish and white mineral (D. 0.7-0.5 cm), as well as a copper bead, laid at the feet.

##### *Tiraspol Kurgan 3*

Tiraspol (46.848, 29.597)

Kurgan 3 near the city of Tiraspol was investigated in 1983 [169]. The kurgan was located 500 m to the south of the Tiraspol-Odesa highway. The kurgan was 3.5 m in height from the buried soil and 60 by 60 m in diameter at the base. The kurgan contained 31 burials.

### I17747

TiK3.15, Kurgan 3 Burial 15, male, 2865-2576 calBCE (4115±25 BP, PSUAMS-7864)

Burial 15, containing Usatove features, was discovered by a round spot of red-colored fired clay in the western sector of the kurgan, 7.7 from the central benchmark, at a depth of 3.55-3.6 m. The 1 × 06 × 4.03 m rectangular burial chamber had rounded corners and was oriented SWW-NEE. A child skeleton was laid on the back, slightly tilted to the left, skull to NEE at 75$^{o}$. Legs were bent, likely up to the chest, with knees to the left. Fragments of a molded jar-shaped vessel were found next to the skull. The vessel was made from clay mixed with sand.

#### Ukraine

(Contact: A. G. Nikitin, S. Ivanova, I. Potekhina, M. Lillie, N. Kotova, T. Tkachuk)

##### *Bil'shivtsi*

Ivano-Frankivsk Region, Ivano-Frankivsk District (49.181, 24.75)

The multi-layer archaeological site at Bil'shivtsi (Kuty) is located on a loess hill on the west bank of the Hnyla Lypa River, the east tributary of the Dniester River.

### I13071

BLV1, Bil'shivtsi-Kyty Burial 1, female (55-60), 2271-2039 calBCE (3741±20 BP, weighted mean[10] of 3770±30 BP (BETA-446187) and 3720±25 BP (PSUAMS-7848))

An oval pit, 1.28 × 1.52 m wide and 2.57 m deep, was uncovered at Bil'shivtsi at level V stratigraphic sequence associated with the local Koshylyvtsi group of Trypillia[31]. A dog skeleton in an extended supine position on the back with hind legs in the shape of a rhomb, skull to the east, facing north, was found at the bottom of the pit, 2.54 m deep. At the depth of 1.5 m three human crania without mandibles were excavated in the south-east part of the pit. Sherds of LBK and Trypillian pottery and a fragment of a clay female figurine were uncovered in the fill of the pit. The dog skeleton was dated to 3793-3377 calBCE (4840±80 BP, Ki 8273).

At around 4 m to the southeast from the eastern edge of the burial of the dog and the three crania, a two-part burial chamber was excavated. The chamber consisted of a western rectangular 1.65 × 0.8 m wide and 1.9 m deep pit and an oval eastern pit in the shape of a catacomb, 2 × 1.65 m wide, 2 m deep, and 0.7 m in height. A sherd of a Funnel Beaker pot with cord print ornamentation was found in the fill of the catacomb.

The skeleton of a mature (55-60) woman was found at a depth of 1.3 m in contracted position faced down, legs to the right, arms stretched along the body, left wrist missing, oriented south-east. The bones of a juvenile sheep were found under the torso. A bovine mandible, laid out on ashes, was found in the east part of the pit, at the entrance to the burial chamber near the right foot of the interred. Grave goods included four polished boar tusks, a cylindrical 0.5 × 0.25 mm copper pin, and a triangular cooper knife 5.7 cm long and 3.2 cm wide.

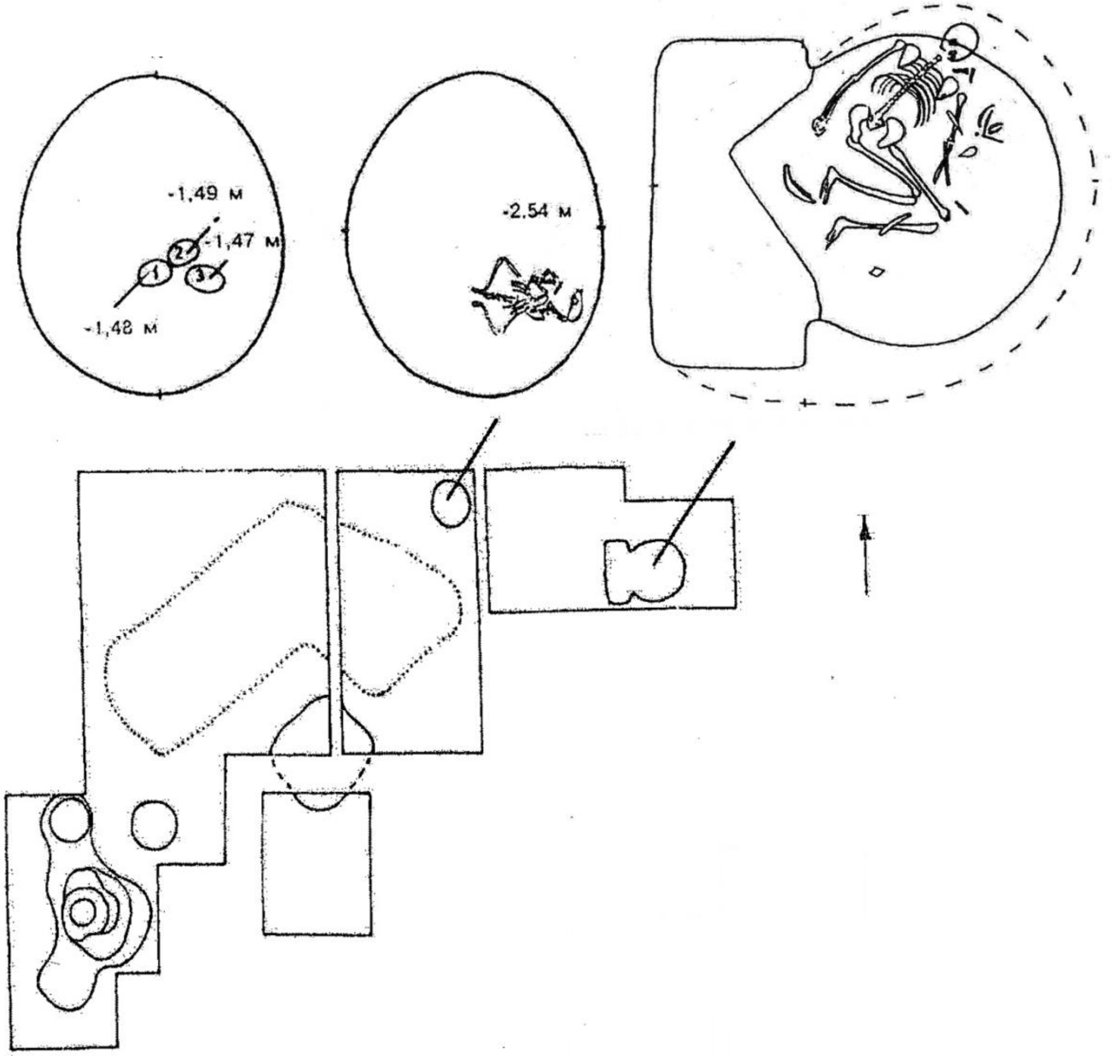

*Bi'lshivtsi (Kuty) burial. Image from[31], modified.*

#### *Deriivka II*

Onufryiv District, Kirovohrad Region (48.933, 33.766)

The Eneolithic settlement of Deriivka II was discovered by an archaeological survey in 1959. The site was located on the west promontory of the Omelnyk River. The Eneolithic cemetery was located 400-500 m from the western outskirts of the settlement. The site was excavated in 1960-1967 and 1983 by D. Ya. Telegin. The Deriivka II anthropological collection is housed in the Bioarchaeological Stores of the Institute of Archaeology of the National Academy of Sciences of Ukraine in Kyiv. Burial descriptions are from [14].

### I27282

Der15, Burial 8, male, 4322-4052 calBCE (5345±30 BP, PSUAMS-10370)
A mature individual in a contracted on the back position, legs flexed upward, hands on the pelvis, eastern orientation.

### I27283

Der41, Burial 11, Skeleton 1, male, 4444-4262 calBCE (5500±30 BP, PSUAMS-10371)
Paired burial with a child. Half-sitting position, legs flexed, hands placed near the abdomen. A polished stone axe was placed near the left half of the pelvis.

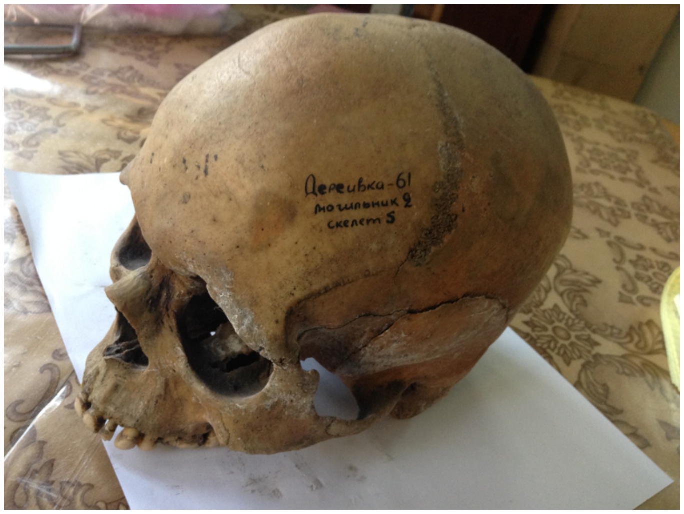

*Deriivka 2 Burial 5. Photo by A. G. Nikitin, 2014.*

### I28319

Der16, Burial 5, female, 4321-4052 calBCE (5344±29 BP, Combination R_Combine ([5380±90 BP, OxA-5032 [15]]; [5340±30 BP, Beta-445808]))
Burial of a mature female in a contracted on the back position, legs flexed and fallen to the right, oriented to the north. Arms bent at the elbows, hands on the pelvis.

##### *Dubynove Kurgan 1*

Dubynove, Odesa Region, Podilsk District, (48.1, 30.28)

The kurgan was located on a plateau on the west bank of the Southern Buh River, 200 m east of the village Dubinovo, at the eastern side of the Kyiv-Odesa highway. An extensive hollow was observed around the kurgan, on the site of an ancient excavation of bulk soil. From the level of this depression, the height of the kurgan was 2.0–2.2 m with a diameter of the

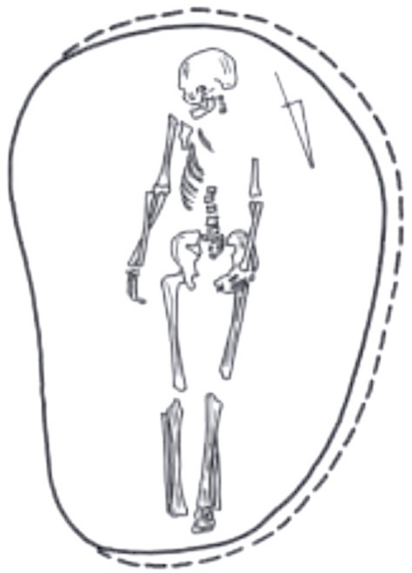

*Dubynove Kurgan 1, Burial 5. Image from*[13].

embankment of 36 m. A total of 15 burials were identified in the kurgan, including one Eneolithic (Burial 7), two Yamna (Burials 5, 13), five Catacomb (Burials 8–12), one Babyne (Burial 14) and one Late Bronze Age burial (Burial 3).

### I12621

DBN1.9, Dubynove Kurgan 1, Burial 9, male, adult, 2575-2348 calBCE (3970±35 BP, PSUAMS-7791)

The oval 2.1 × 1.4 m burial chamber of Burial 9 was discovered 8 m to the southeast of the benchmark, at a depth of 2.55 m. It was oriented with its long axis in the south-southwest - north-northeast direction. The entrance well was located at the eastern edge of the chamber. The interred was lying in extended supine position on the back, with his limbs straight, head to the south-southwest. The left hand was placed on the pelvis. The face was turned to the east. There was no inventory.

### I12840

DBN1.10, Dubynove Kurgan 1, Burial 10, female, juvenile, 2453-2148 calBCE (3830±30 BP, PSUAMS-7805)

Burial 10 was discovered 8.3 m southeast of the benchmark, at a depth of 2.85 m. The burial structure was a 1.4 × 0.7 m elongated pit, with a lining in the western wall. The interred lay on the back, head to the south-southeast. The skull was tilted towards the left shoulder, the arms extending along the body with the wrists resting on the pelvis. There was no inventory.

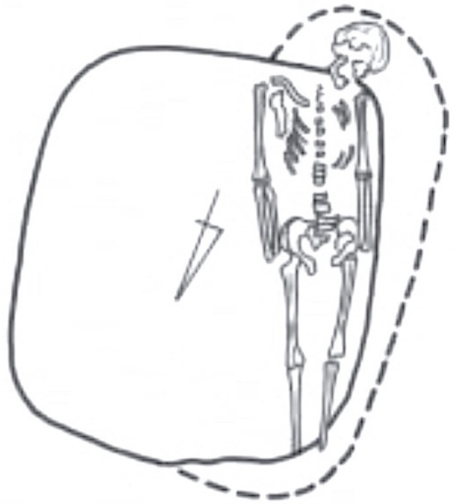

*Dubynove Kurgan 1, Burial 10. Image from*[13].

##### *Kam'yana Mohyla Kurgan 2*

Myrne, Zaporizhzhia Region (46.951, 35.470)

Kam'yana Mohyla is a Ukrainian National Historical and Archaeological Preserve in the Steppe of Azov, 16 km north of from Melitopol, on the west bank of the Molochna River. Kam'yana Mohyla is a natural formation, the height of which reaches 12 m. It occupies an area of 1.6 hectares, on the territory of which there are about 3000 sandstone slabs. Geologically, the monument is a remnant of Sarmatian Sea sandstone (the Tertiary period), with cracks and faults that led to the accumulation of slabs and the formation of a large number of grottoes.

Long-term research of Kam'yana Mohyla led to the discovery of 65 grottoes and caves, on the ceilings of which several thousand petroglyphs from the Late Paleolithic and Mesolithic to the Middle Ages were discovered. The grottoes are mainly small (up to 5-8 m), low (1.2-1.5 m) rock cracks with flat (or close to them) ceiling slabs. Nowadays, a large part of the grottoes is filled with sand (conserved) to ensure the preservation of ancient petroglyphs.

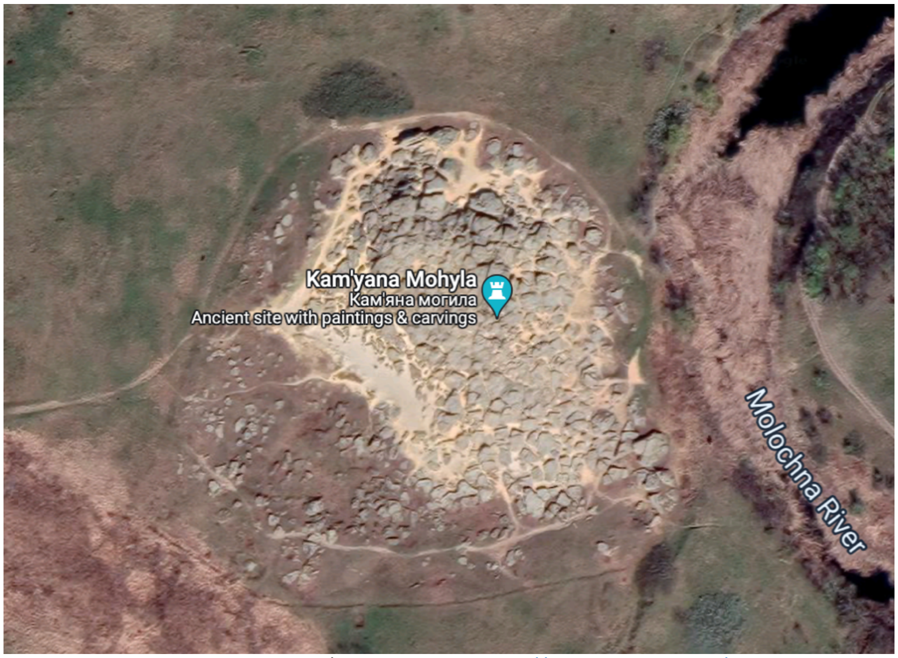

*Kam'yana Mohyla. Google Maps/Google Earth, https://www.google.com/maps, 2022.*

I20975

KM17P2; KM-2017 Kurgan 2, Burial 2, male, 2872-2627 calBCE (4141±21 BP, BE-8040.1.1)

Kurgan 2 was excavated in September 2017 by the Azov Archaeological Expedition of the Institute of Archeology, National Academy of Sciences of Ukraine, led by S. V. Makhortykh[26,170]. The investigated monument was located 200 m north of Kam'yana Mohyla. No visible traces of the mound were recorded at the location of Kurgan 2 before the excavations began. The uneven, bumpy surface was planted with trees and covered with grass. During the excavations, the stratigraphy of the studied area became quite clear. The upper layer (from 0.19 to 0.25 m) consisted of chernozem. A layer of light gray loam, in many places disturbed by burrows, lay under it. The thickness of this layer varied from 0.35 to 0.50 m. Below the light gray loam was a thick layer of brown loam with numerous crushed shells. Its traced thickness was at least 0.4-0.6 m.

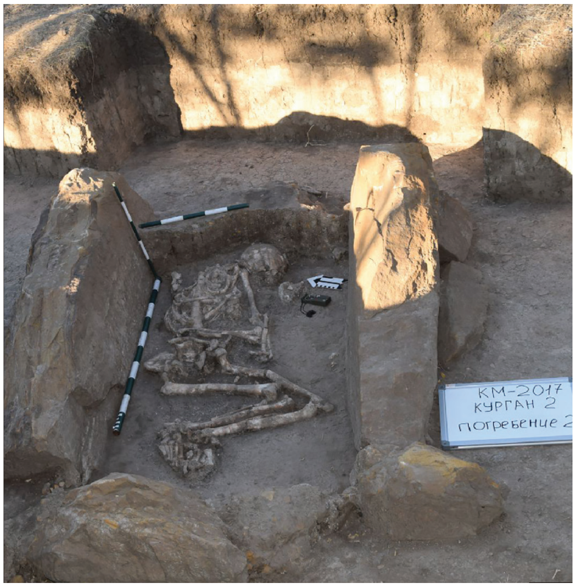

*Kam'yana Mohyla Kurgan 2 Burial 2. Photo: S. Makhortykh.*

During the excavation of the kurgan, three burials and two presumed ritual complexes were discovered. They were located in a layer of light gray loam. Burials 1 and 3 were located in the kurgan's middle and upper parts, burial 2 in a stone tomb, and two ritual complexes were in its lower part.

Burial 2 was made in a Globular Amphora-style stone sarcophagus, which was located on an ancient horizon consisting of brown loam with numerous admixtures of crushed shell. It had a rectangular shape and was built from rough-hewn sandstone slabs placed on an edge. The burial structure was oriented with its long sides along the northeast-southwest line. No stone covering over the tomb was found. Individual bones of animals, such as the rib of an adult bull, and the hoof of a horse, were associated with the burial. In the central part of the tomb, next to the southern wall, a Unio shell was found.

The skeleton of a young male (< 25) discovered in the tomb was lying contracted on the left, with the head to the east, tilted to the north. The mandible was displaced and located in front of the face of the interred. The clavicle, revealed between the ribs, was also displaced. The hands of the deceased were also partially displaced. The left hand was placed extended towards the knees. The right arm lay perpendicular to the left and appeared to be separated from the body (the elbow and shoulder bones laid parallel to each other). The legs were bent at the knees and sunk to the left. The phalanx of the finger lying next to the right femur had a red articular surface.

##### *Katarzhyno Kurgans 1 and 2*

Znamianka, Ivanovsky district, Odesa Region (47.025, 30.288)

The Katarzhyno kurgan group was excavated in 1990-1991 by the expeditions of the Odesa Protective Archaeological Center under the Ukrainian Society for the Protection of Historical and Cultural Monuments. The kurgan group was located 4 km south of the village Znamianka (until 1932 - the village of Katarzhyno), on the plateau of the east bank of the Malyi Kuyalnyk river. The burial ground consisted of five kurgans. Four of them, including Kurgan 2 with a height of 2.4 m and kurgans 3–5, plowed up to the foundation, stood in a row from north to south. The largest kurgan 1, 6.45 m high, was located somewhat to the side. At the same time, the group closed a rarefied kurgan chain from the north, stretched along the meridional watershed up to 120 m high above the thalweg of the river. Kurgan 1 of the Katarzhyno group is one of the five largest kurgans excavated in the northwestern Black Sea region.

Kurgan 1 was 80–85 m diameter. Its height at the beginning of the excavations was 6.45 m, while according to the topographic survey of 1950, the height was 7.3 m (the absolute mark is 134.7 m in the Baltic system of heights). The slopes of the kurgan were heavily raked and partially cut off. The summit had also been cut off and disturbed by modern digging. In total, 20 burials were found in the kurgan: one Eneolithic (burial 10), Yamna — 11 burials (1, 2, 6–8, 9 (?), 11–14, 16, 21), Catacomb — two burials (3, 5), Babyne (Cordoned Ware) – three burials (9, 17, 20), two burials of the Bronze Age (4, 18) and one indeterminate burial 15.

Kurgan 2, 2.4 m high and about 50 m in diameter, was located 20 m northwest of Kurgan 1. The kurgan was heavily plowed, and its top was cut off. In total, six burials were found in the kurgan. Of these, two (Burial 1 and Burial 6) belonged to the Yamna archaeological complex. The other four burials belonged to the Babyne archaeological complex (Burial 5), late nomadic (Burial 2), and two of indeterminate cultural association (Burials 3–4).

### I11999

KTR1.1, Katarzhyno Kurgan 1 burial 1, male, adult, 2626-2475 calBCE (4045±20 BP, PSUAMS-4903)

Burial 1 (YAC) was found 25.5 m west-southwest of the central benchmark. At a depth of 6.25 m, in a section of 3.2 to 3 m, stones smeared with green clay and large unworked floor slabs of a stone tomb emerged. The southern slab of the stone cyst burst and sank. The tomb was a rectangular stone box of four slabs placed in a ground pit. After the burial, the pit with the tomb was filled in. The burial box, 1.9 × 1.2 × 0.9 m, was oriented in the north-south direction. On the northern, eastern, and southern slabs, traces of ornamentation with pale pink ochre have been preserved. The buried individual was laid with the head to the north, contracted on the back. The legs were bent and turned to the left but may have originally stood with their knees up. The arms extended along the body, the left arm was straightened, the right arm was slightly bent at the elbow, and its wrist almost touched the pelvis (the hand was not preserved). The skull was tilted towards the left shoulder. A limestone triangular slab, without traces of processing, was laid on the chest of the interred. Considering that the roof of the tomb over this part of the burial was not broken, it can be assumed that the slab was placed deliberately. Small sharp-angled stones were placed in the upper left part of the chest, between the femurs, and on the right tibia. At the head, at the left forearm, and the corners of the tomb, scatterings of pink ochre powder were traced. On the chest bones on the right were traces of fire in the form of charcoal and burnt pieces of bark. At the elbow of the right hand and the right wrist, two copper or bronze tubular piercings were found. Other than these, no other funeral offerings were found in the tomb.

I31733
KTR1.10, Katarzhyno Kurgan 1 burial 10, female, 3955-3783 calBCE (5060±30 BP, PSUAMS-10375)

Burial 10, main, Eneolithic, was found 6.5 m southeast of the benchmark, under an embankment of several layers of earth and a layer of stone. At the level of the ancient day surface, the burial pit was covered with a reed mat with a preserved direction of the fibers and a single layer of limestone. Among the stone slabs was a fragment of an anthropomorphic stelae with a chiseled human head. The reed mat near the stellae was covered with a layer of 5–8 mm organic matter in the form of a dense yellowish-white streak (honey?), covered with a large fragment of a ceramic pot. Two other slabs without signs of processing, were laid on the sides of the stone syst on the east side. The bottom of the tomb was covered with a dark brown decay of the bedding, on which the interred was laid contracted on the back. The legs were bent, and the knees turned to the right, the arms were straightened and laid along the body with the palms down, the skull was tilted to the left shoulder, and the orientation was southwestern. The reconstructed height of the individual was 178-193 cm[27]

Ochre was noted: a thin deposit on the bones and massive flat cakes about 2.5 cm thick. A whole cake, 6 cm in diameter, laid near the left hand at the level of the elbow. On the chest, near the right clavicle, a segment of a larger cake, about 10 cm in diameter, was found; coals were found under the cake. At the left hand, closer to the shoulder, there was a vessel placed on a layer of coal. Other finds are represented by flint tools. One of them, with retouched ends, was found under the skull, the other was in the filling.

Burial inventory: 1) A molded askos-type vessel with a characteristic asymmetry of shape and an oval-shaped broken-off handle. 2) Half of a molded clay pot (without a bottom) of rough dressing with an ample admixture of crushed shells and a corded ornament in the form of a horizontal four-row belt and bows attached from below. 3) A chisel-type greyish-smoky flint tool with two working sides. 4) The same type of tool (gray flint), but more massive and with one working side.

The specimen failed amplification at Reich's Lab. Low-resolution mtDNA typing conducted in the Molecular Archaeology Lab at GVSU following the protocol detailed in [9]

## I12168

KTR1.13, Katarzhyno Kurgan 1 Burial 13, male, adult, 3024-2908 calBCE (weighed average of 4335±25 BP, PSUAMS-9815, and 4375±20 BP, PSUAMS-4762)

Burial 13 (early Yamna) was found 5.6 m south of the central benchmark, at a depth of 2.35 m. It was not possible to trace whether there was a ledge near the burial chamber. The 2.0 × 0.9 m chamber was covered with four large limestone slabs without traces of processing. The chamber was rectangular with rounded corners, oriented in length in the northeast-southwest direction. The skeleton was poorly preserved. The humerus of the right hand was missing, the skull was destroyed. The interred was laid on the back, with the head to the northeast. The arms were extended along the torso. Bent legs were placed with knees up. At the bottom of the grave was a bedding of bark sprinkled with chalk. A layer of ochre was deposited at the left shoulder.

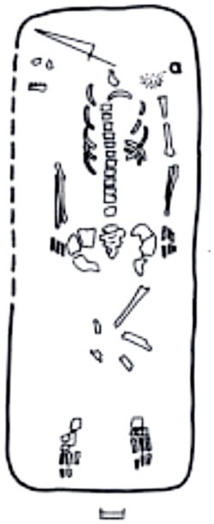

*Katarzhyno Kurgan 1 Burial 13. Image from[13].*

#### I12170, KTR-2.1

Katarzhyno Kurgan 2 burial 1, female, adolescent, 3311-2578 calBCE (4270±90 BP, Ki-17331), mtDNA U4c1.

The main Burial 1 (YAC) was found 1.4 m south of the central benchmark, at a depth of 2.3 m. At this level, a deposit of small stones (0.2–0.3 m in size) was recorded, which lay flat in one or two tiers within the grave. Under the stones, the dust of the wooden floor was traced. The burial chamber was rectangular in plan, 1.7 × 1.1 m, with rounded corners, 0.6 m deep from the level of the ancient day surface. The buried individual was laid on the back, with the head to the northeast. The arms were slightly spread apart, bent at the elbows, and extended along the body. The legs were turned with the knees to the right. Between the right side of the chest and the pelvis was a stone tool. A small piece of limestone rested on the sacrum. Under the pelvic bones and above, at the level of the lumbar spine, a layer of 1 cm of green clay was traced. Burial inventory: 1) Gray sandstone slab, prismatic shape, 11.7 × 6.3 × 3.4 cm, used as a pestle and grater stone. As a result of use, the upper plane of the tool was polished, and the lower and all side faces were uniformly covered with characteristic potholes. 2) Sturdy limestone tile in half-segment shape, 6.6 X 3.6 × 2 cm, with sharp edges. One edge was blunted during the use of the stone, possibly as a scraper or adze.

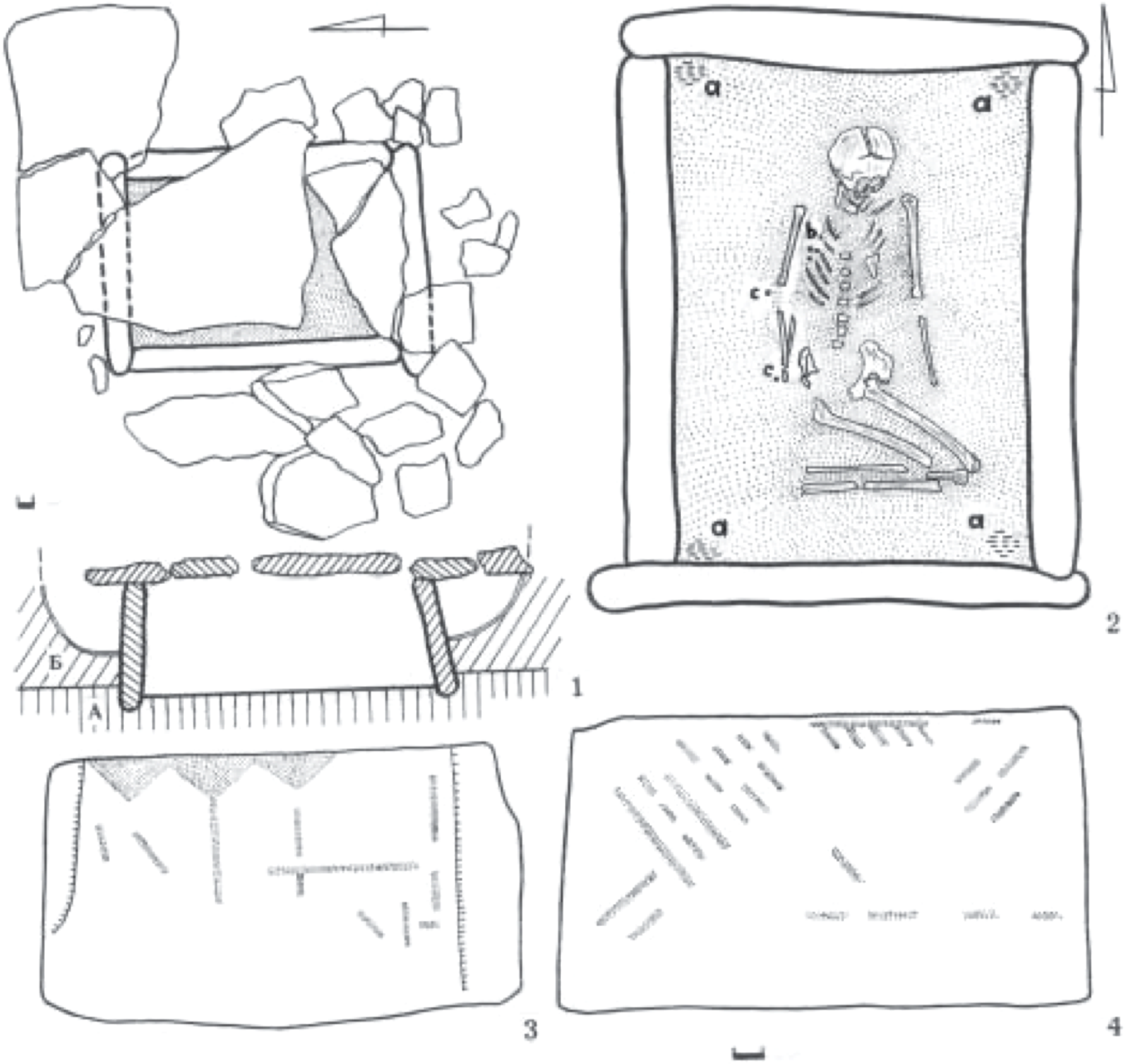

*Katarzhyno Kurgan 1 Burial 1. Image from* [13].

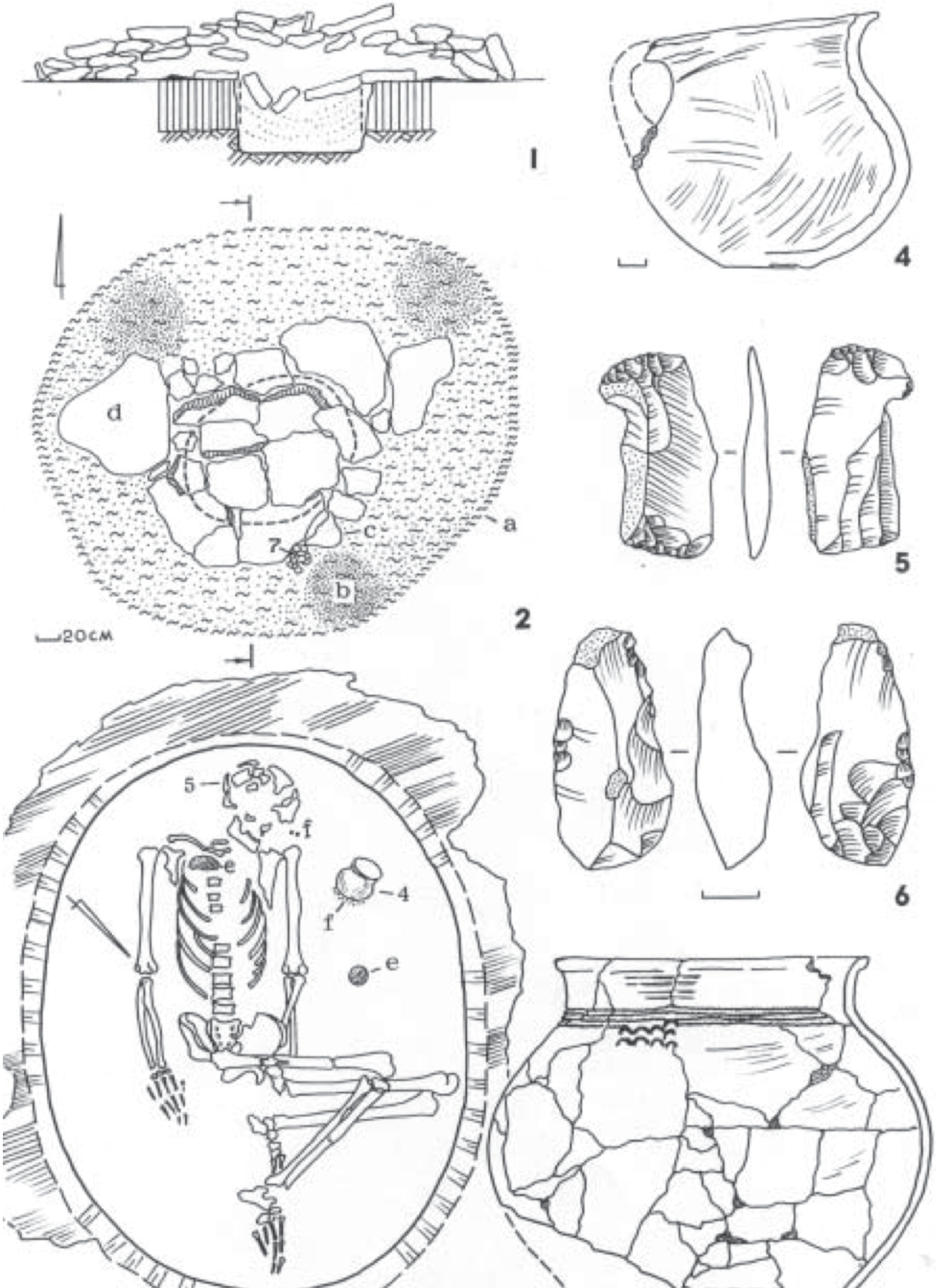

*Katarzhyno Kurgan 1 Burial 10. Image from* [13].

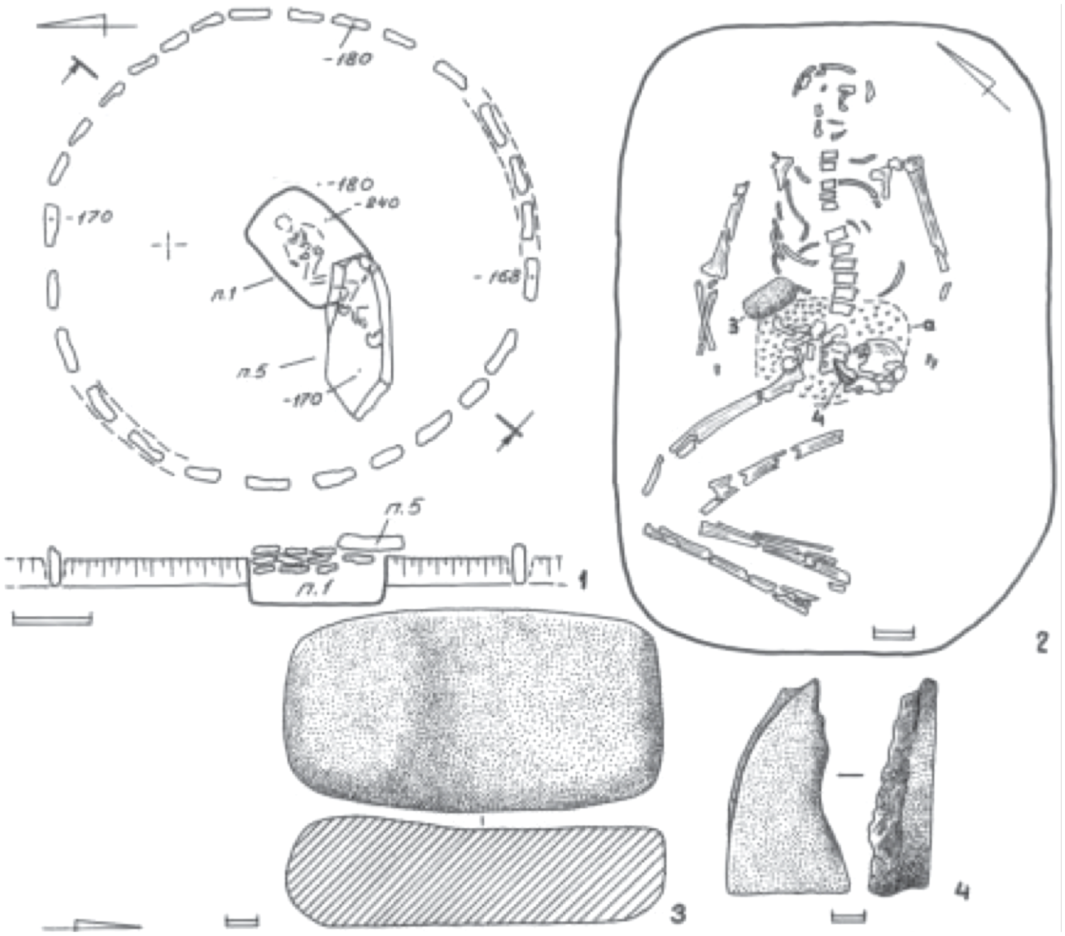

*Katarzhyno Kurgan 2 Burial 1. Image from* [13].

#### *Kolomiytsiv Yar Tract*

Kopachiv, Kyiv Region**,** Obukhiv District (50.130, 30.480)

The Kolomiytsiv Yar Tract archaeological site and burial are detailed in[16].

## I7585

KST07, female, 4049-3820 calBCE (5170±30 BP, BETA-523816).

#### *Mariupol Necropolis*

Mariupol, Mariupol District, Donetsk Region (47.0998, 37.6003)

The necropolis was located on the east bank of the Kalmius River, on a peninsula between the shores of the Sea of Azov and the bend of the Kalmius River, opposite the city of Mariupol. The blast furnace of the Azovstal steel plant was built on the site of the necropolis in the summer of 1931. The necropolis was excavated by Mykola Makarenko in 1930, August 10 through October 15[1].

The necropolis was distinguished in the ground by a strip of red clay 28 meters in length and 2 meters wide. On the peninsula where the necropolis was located and up to the gully that separates it from the general coastal plain, such clays were not present. Deposits of similar clays have been found in ravines and cliffs of the Kalmius River at a significant distance upstream from the burial site. The burials, all in extended supine burial position except for the triple burial 21, were located in this clay strip 0.40 to 0.75 meters from the surface, with a maximum depth of 1.5 meters. A total of 124 burials were excavated.

Extended burials were archaeologically assigned to the Neolithic period. Burial 21 was considered by Makarenko to be of a later period. The Neolithic double burial 50 and the triple burial 21 were cut out of the matrix in the field by Makarenko and brought to the Mariupol Museum of Local History, where they were sampled in the fall of 2021 for the current analysis. The museum and its collections were destroyed by the Russian military during the siege of Mariupol in the spring of 2022[150].

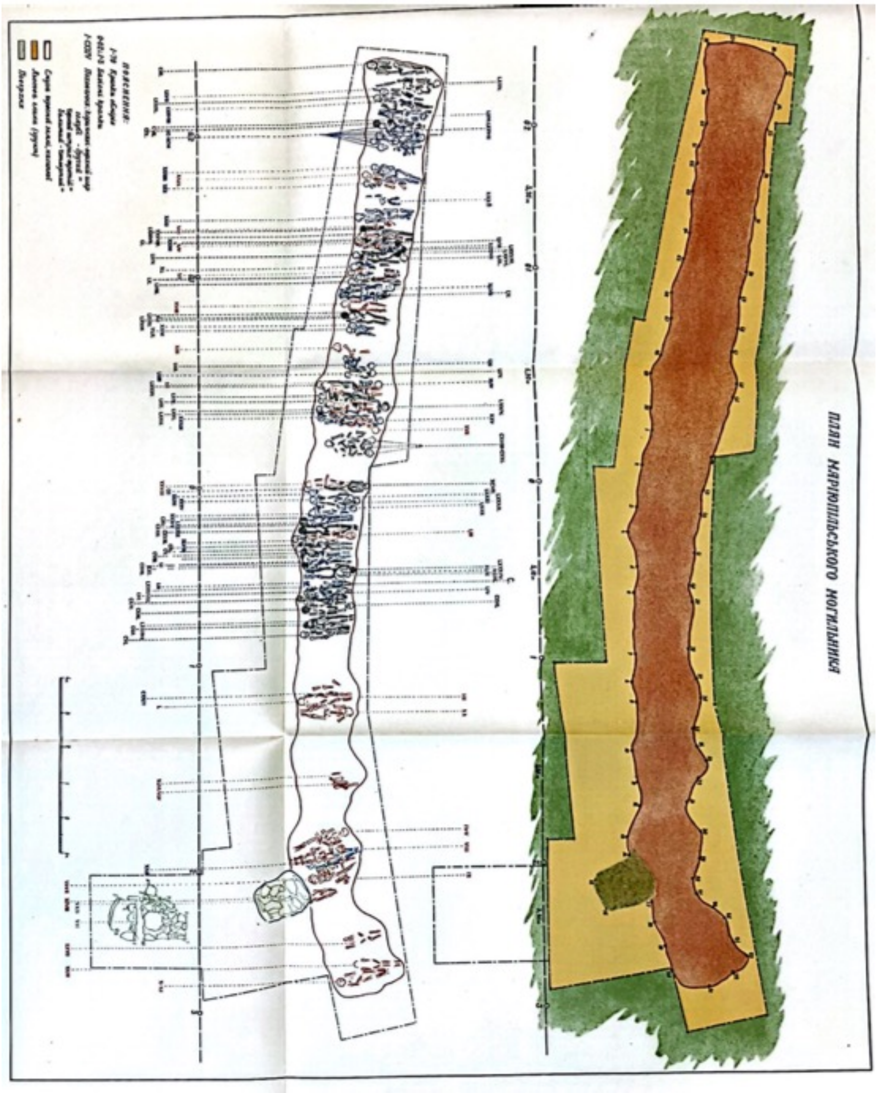

*Mariupol Necropolis layout. Image from*[1].

I31730

MM50a, Mariupol Necropolis, Burial 50, Skeleton 50a, male (adult), 5474-5236 calBCE (6395±35 BP, PSUAMS-10373)

I31731

MM50b, Mariupol Necropolis, Burial 50, Skeleton 50b, immature, 2204-1986 calBCE (3720±30 BP, BETA-681064, radiocarbon date from tooth enamel carbonate).

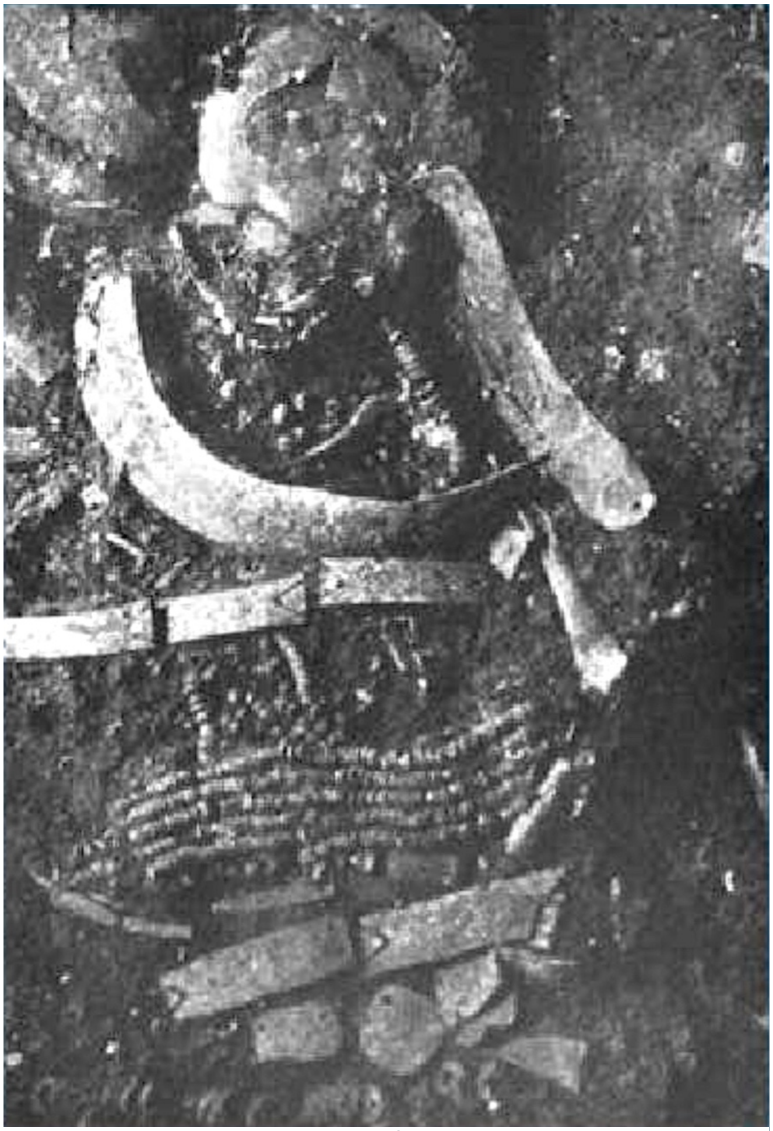

*Mariupol Necropolis, Burial 50b. Photo from*[1].

Burial 50 contained a double inhumation of an adult (Burial 50a) and a child (Burial 50b) in an extended supine position, heads to the west. Starting from the sacrum and going down the legs, both skeletons were covered with a mass in the form of clay of a pure purple color. The head of an adult was decorated with a pair of boar tusks so that the sharp ends of the fangs protruded above the forehead. A large boar tusk and a necklace made of mother of pearl cut into disks were found around the neck. The child skeleton, laid on top of the adult, was adorned with boar tusks over the head and neck. A row of narrow plates lay across the chest and around the body. Five necklaces made from mother-of-pearl disks laid on the stomach. The necklaces laid tightly across the skeleton like a belt. Several narrow plates made of boar tusks laid vertically around the right hand, on the line of the strings of the necklace. A belt made of the same narrow boar tusk plates laid below the belt with the necklace. Between them and the rows of small mother-of-pearl necklaces laid a skillfully polished miniature wedge made of green porphyrite, partly covered with rows of necklaces and partly with a row of plates. Below the row of these plates lay wider plates made of the same material, and in between lay a pear-shaped flat pendant with a hole at the narrow end. Further, down to the knees, laid a row of shell beads. Each bead was shaped like a circle with a cut-off segment. On the knees laid two rows of wide boar tusk plates and a scattered mother-of-pearl disks necklace.

## I31732

MM.21, Mariupol Necropolis, Burial 21, female, 4715-4548 calBCE (5790±30 BP, PSUAMS-10374)

The burial consisted of a cremation and a double inhumation. The burial was found within a square covered with limestone stones of various sizes, 0.4 m from the sod surface, and between 0.3 m and 0.6 m above the first row of the Neolithic burials. Stones covered the surface with a layer up to 0.4 m in the shape of a crescent. A pile of ashes containing cremated human bones, approximately 0.6 m in diameter, was found in the center of the grave, 0.45 m from the surface. Stones beneath the ashes were burned and covered in soot. A finely knapped arrow or spearhead point with a wide base and traces of fire, an elongated scraper in the form of an arrowhead with finely retouched edges, several boar tusk plates with transverse double incised lines along the shorter edges and without peripheral holes, and a black oblong stone with polished and worn-out edges, were found among the ashes.

The grave was outlined with upright limestone slabs. One large slab was positioned at the eastern edge, surrounded by smaller slabs on the sides. The burial pit was cut into the upper part of the Neolithic burial 98. The two interred individuals were laid contracted on their right side, heads facing east, arms extended over the sacrum bones. The skulls were significantly elevated relative to the postcranial remains and close to each other. A long flint blade was found in the hands of one of the skeletons.

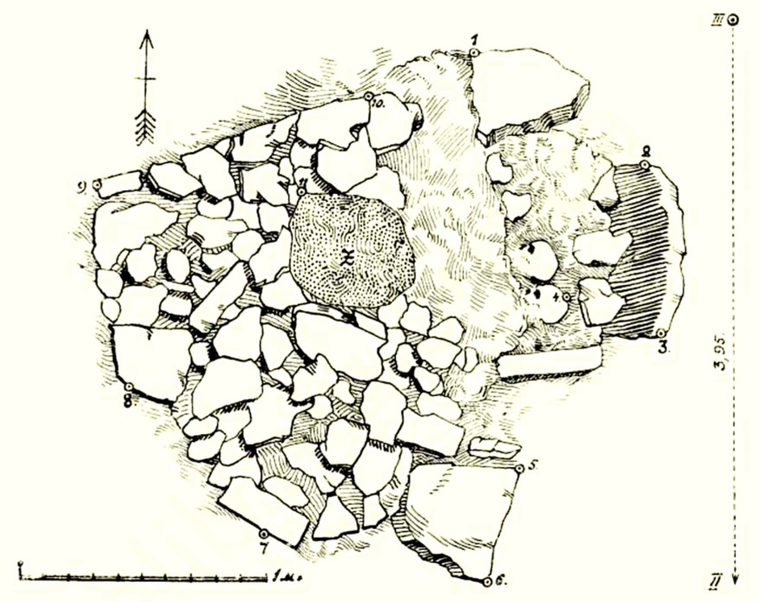

Рис. 27. Загальний вигляд кам'яної кладки поховання XXI.
Fig. 27. General aspect of a stone layer in grave XXI.

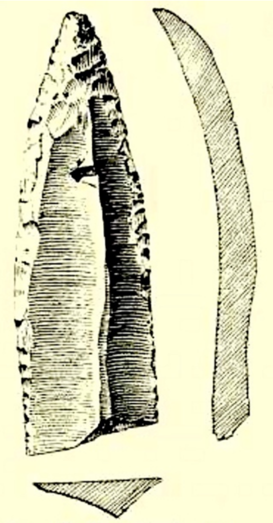

Рис. 28. Кремінний наконечник списа чи стріли над похованням XXI.
Fig. 28. Flint spearhead over grave XXI.

*Mariupol Necropolis Burial 21 (left) and the flint arrow- or spearhead from the pile of cremated bones on top of the burial. Images from*[1].

##### *Mayaky*

Belyaiivsky District, Odesa Region (46.397, 30.272)

The Mayaky archaeological complex is located on the southwestern fringe of the Mayaky village on a promontory along the eastern bank of the Dniester River near its convergence with the Dniester Estuary. [171]. The 300 × 100 m ancient cemetery situated 200 m north of the network of ditches enclosing a prehistoric sanctuary. Nineteen sites within the cemetery were designated as kurgan or ground-level burial plots [11,71]. Burial positions at the Mayaky necropolis, for the most part, featured left-sided positioning with heads oriented northeastward[11,69]. Artefacts from the Mayaky necropolis reported here were unearthed during archeological in 1965, 1974, and 1986 [11,171,172]. Sanctuary burials 1 and 2 were excavated by V. G. Zbenovich [171]. The Mayaky anthropological collection is housed in the Bioarchaeological Stores of the Institute of Archaeology, National Academy of Sciences of Ukraine, Kyiv, Ukraine, from which it was sampled in 2014. Burial descriptions are from [11,171,173]. Detailed descriptions of burials I12229, I12707, I12705, I12706, and I12710 are also presented in [75].

#### I12615

Ma6, Mayaki sanctuary, burial 1, female, 5208-4907calBCE (6100±30 BP, BETA-432410)

#### I12228

Ma13, Mayaki sanctuary, burial 2, female, 4486-4350 calBCE (5580±30 BP, BETA-432406)

#### I12229

Mau1, Kurgan 1, Burial 9, female, 40-45, 3088-2911 calBCE (4375±25 BP, PSUAMS-7865)

Kurgan 1 was erected over the main burial 9. The burial, containing ceramics of the Usatove type, was centrally located in the kurgan. The burial was made in a rectangular pit with rounded corners, 2.30 × 1.30 m in size, let into the matrix. The depth of the burial pit from the modern surface was 1.65 $m^2$. The walls of the burial pit were vertical, and the bottom was leveled. The pit was oriented from west to east. The western edge of the pit was destroyed by a depression with a diameter of 0.50 m. The northeastern part of the grave was damaged by a robbers' trench. The remnants of wooden planks found in the filling of the burial pit suggest that the pit had a cover. A stain, formed by organic decay, probably from the bedding on which the deceased was placed, was found at the bottom of the pit.

Only the bones of the legs along with the remnants of the bedding were found *in situ*. Pelvic bones appeared to have been shifted. The cranium was placed on top of the pelvic bones. Disarticulated postcranial skeletal remains (Burial 9.1, female, 92 calBCE - 66 calCE (2020±25 BP, PSUAMS-7792[75]).

Judging by the degree of obliteration of cranial sutures and the wear of the teeth, the individual was 40-45 years old at the time of death. The abrasion of the teeth was continuous. The powerful relief of the cranium and bones of the skeleton led to the male sex assignment by anthropology.

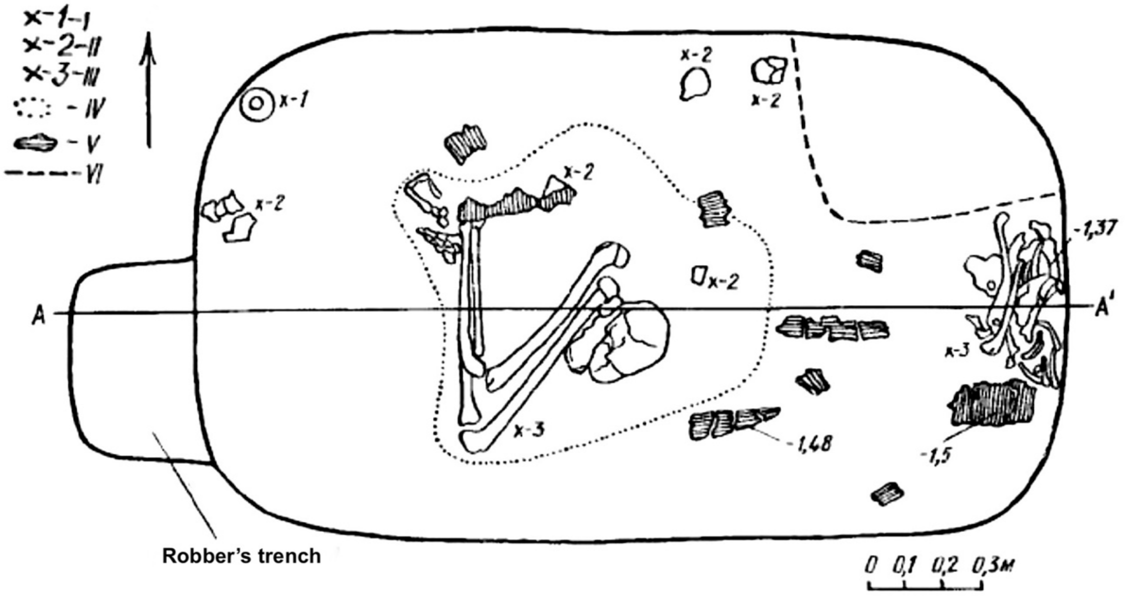

*Mayaky Kurgan 1 Burial 9. Image from [173], modified. I, unornamented pot; II, amphora fragments, III, human skeletal remains; IV, traces of the bedding; V, remnants of wooden cover; VI, contours of a modern dugout.*

On the right side of the braincase of the skull there were a series of stripes of dark violet paint. A similar stripe, crossing the left part of the arch from the back of the head, descended to the eyebrows and under the right eye socket. The left side of the cerebral part of the skull was deformed (plagiocephaly), the right part of the facial skeleton is much smaller than the left. The deformity was created by systematically placing the subject as an infant on his left side. The cracks on the left parietal bone of the skull are traces of blows inflicted on the subject, possibly causing death.

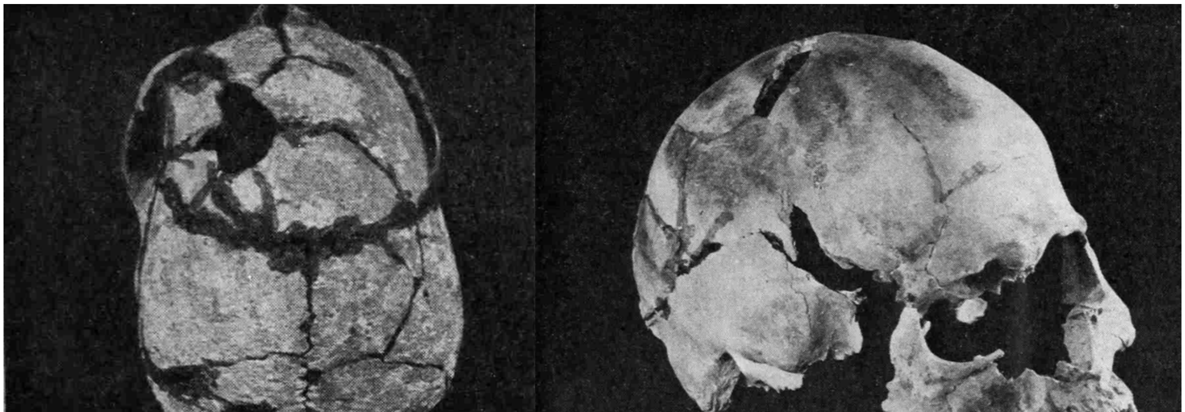

*Mayaky Kurgan 1 Burial 9 cranium, showing traces of paint and a perimortem injury. Photos from[174] .*

### I12704

Mayaky, Kurgan 7, Burial 2, male (25-30), 3620-3030 calBCE (RE-adjusted) (5144±20 BP, weighted mean[10] of 5530±32 (OxA-22959[12]), 5390±30 (BETA-441235[5]), and 5545±40 (PSUAMS-7793, this report)).

Burial arrangements 7, 8, and 9, identified as kurgans with missing mounds, were found in the northern part of the Mayaky burial complex, situated next to each other. The southeast side of each kurgan was surrounded by a 0.4-0.5 × 0.6-0.65 m ditch. It is likely that the three kurgans were erected in a chronological sequence, with kurgan 7 appearing first, followed by kurgan 9, and, subsequently, kurgan 8[75].

Burial 2 of Kurgan 7 was determined to be of Usatove origin through archaeological analysis. The grave was discerned as a discoloration in the transition to sandy loam, situated 0.65 m beneath the current surface. Its depth extended to 1.25 meters, with traceable walls reaching 0.6 meters. The oval-shaped burial pit, measuring 1.4 × 1.15 m, exhibited a slight elongation at the bottom (1.55 × 1.15 m), aligned from southwest to northeast. The backfill comprised three distinct layers: black earth with sandy loam lenses up to a depth of 1.05 m, followed by a thin 0.1-m layer of sandy loam, and black earth with numerous lime inclusions at the base.

Within the grave lay the robust skeleton, positioned in a contracted manner on the left side, with the skull oriented towards the east and facing upwards. The arms were bent and crossed over the chest, with the left hand under the skull and the right hand resting by the facial bones. Traces of raspberry-colored ocher remained on the skull, joints of the thighs, and lower legs, accompanied by remnants of organic material resembling brown mold on the lower legs. Dark red ocher crumbs were discovered between the upper ribs.

Burial inventory consisted of two vessels, found next to the arms of the interred. One, a beige beaker standing 46 mm tall with a rim diameter of 45 mm, featured a short body and a low neck. The vessel was ornamented with four horizontal rows of fine cord coated with red ocher around the neck, while incised lines decorated the body, with a crisscross pattern forming a six-pointed star in the upper region and three parallel grooves at the bottom. The clay composition included finely crushed shells. The second vessel, a heavily fragmented miniature bowl measuring 72 mm in height with a rim diameter of 114 mm and a bottom diameter of 38 mm, typified Usatove pottery. It boasted an S-shaped profile, thin walls, and round cord indentations along the rim, complemented by three rows of thin cord indentations underneath. The interior was coated with red ocher, and the shard exhibited a gray-brown hue, with the temper comprising finely crushed shells and grains of sand.

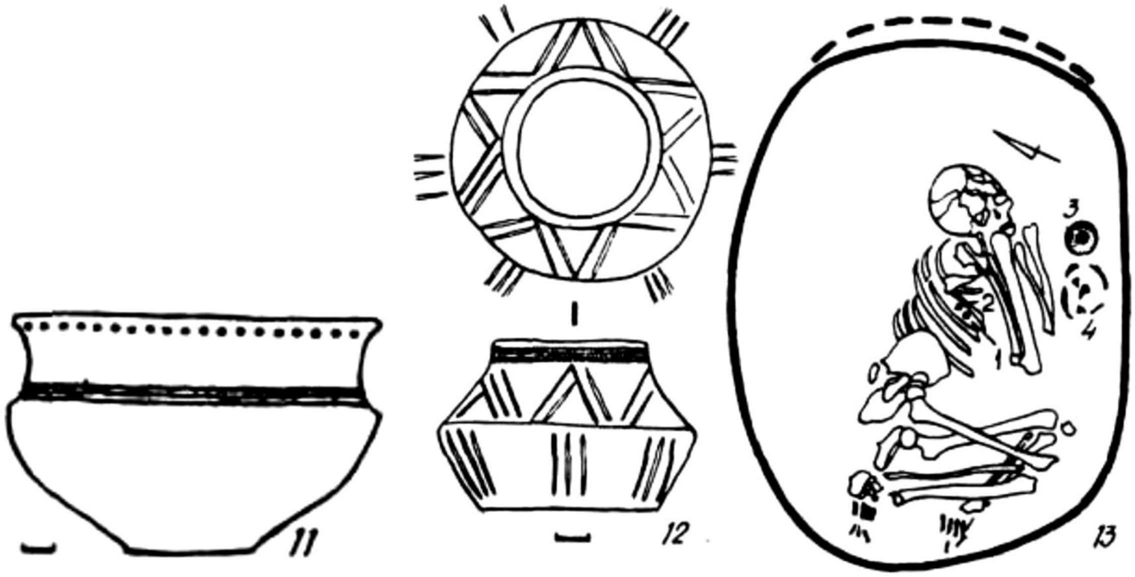

*Mayaky Kurgan 7 Burial 2. Image from*[11].

## I12710

Mau12, Kurgan 8, Burial 4, female (*ca.* 60), 3769-3196 calBCE, RE-adjusted (5295±30 BP, PSUAMS-7862)

Burial 4 of Kurgan 8 was discovered within an oval pit, featuring a 0.30-m-high lining and depths ranging from 0.8 to 1.6 m. The dimensions of the grave measured 1.2 by 0.9 m at the top and 1.3 by 0.9 m at the bottom. The grave fill consisted of two layers: compact sierozem containing calcareous nodules overlaying a base layer of continental loess-like loam.

Within the grave, approximately 0.4 to 0.5 m above the bottom, were discovered fragments of horse, cow, and sheep bones covered with dark red ocher, along with two shards and a fragmented light brown Usatove bowl. The bowl displayed coarse molding with notches, impressions of a round-ended stamp, and cord markings, with sand grains mixed into the clay.

At the bottom of the pit lay the contracted skeleton, positioned on the left side with the skull facing north-northeast at an angle of 33 degrees, and hands bent. The left hand was near the face, while the right hand rested under the cervical vertebrae. Crimson ocher stripes marked the skull vault, with similar markings found on the pelvic bones. Evidence of brown decay was noted on the wrist bones and lower epiphyses of the forearm on the right hand, as well as in the region of the left elbow joint.

Burial inventory consisted of five ceramic vessels located alongside the deceased. These included the lower part of a vessel crafted from clay mixed with crushed shells, exhibiting soot stains on the walls; a polished pot featuring imprints of a cord and a pipe, with holes below the rim and sand grains within the clay; a polished bowl containing finely ground shells and sand grains in the clay; a polished amphora adorned with holes beneath the rim and a corded design; and a pot made from a crumbly clay mixture with crushed shells, displaying a pattern of pipe imprints.

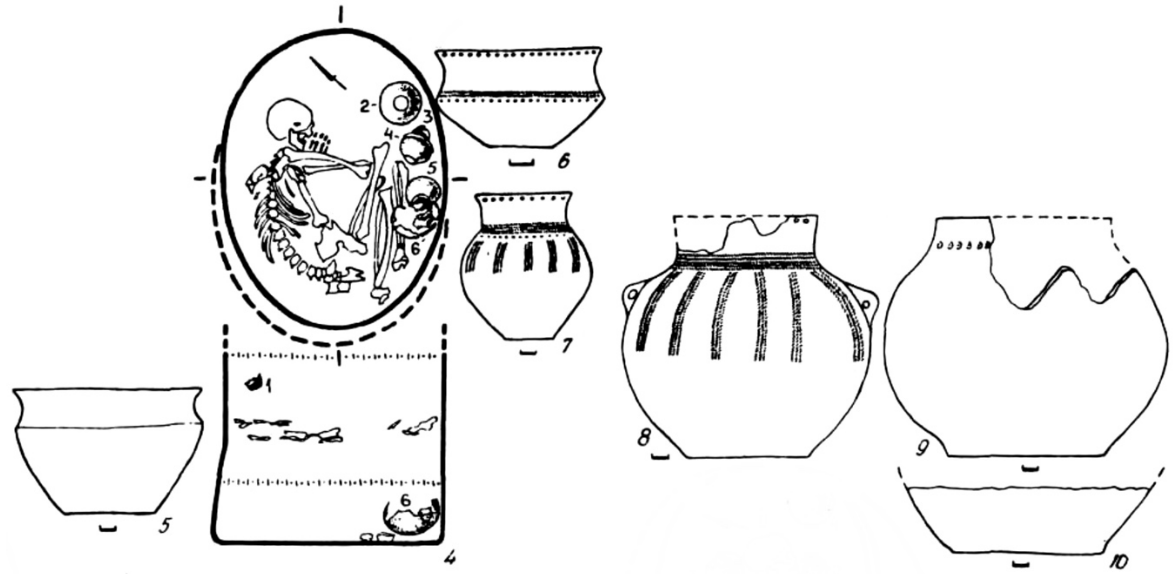

*Kurgan 8 Burial 4 and associated finds. Images from*[11].

## I12707

Mau9, Kurgan 8, Burial 6, Skeleton 2, female (25-35), 3511-2917 calBCE, RE-adjusted (5330±40 BP, PSUAMS-7794)

The burial pit for Burial 6 reached depths of 0.75 to 1.50 meters, featuring a lining and a ledge. The oval-shaped burial chamber measured 1.3 × 0.9 m at the top and 1.35 × 1.10 m at the bottom. Within this grave, two individuals were interred: a 40 to 55-year-old male (Skeleton 1) and a 25 to 35-year-old female (Skeleton 2).

Skeleton 1, dated to 4229-3656 calBCE (5670±30 BP, BETA-4324085[5]), was found at the upper layer of the grave. Beneath a 0.3 m-thick layer of yellow loess-like loam, indistinguishable from the matrix, lay Skeleton 2. This second individual was positioned in a contracted manner on the left side, with the skull facing eastward at an angle of 85 degrees. Notably, the spine exhibited a pronounced curvature, the femurs pressed against the chest, knees brought towards the shoulder joints, and feet positioned near the pelvis. A pot was placed within the hands of the buried individual. Decay stripes with black paint spots were observed on the postcranial skeleton and the ground beneath them, with bright crimson ocher traces noted on the right parietal bone.

The burial inventory associated with Skeleton 2 included a pot crafted from gray crumbly clay mix, featuring flat stamp impressions and containing a flint flake; a vessel made of a loose clay mix with weak firing featuring two protruding protrusions on the body, adorned with cord and round-ended stamp imprints; and a bowl coated on the outside with a layer of greasy soot. These latter two vessels displayed gray and light brown hues, with crushed shells incorporated into the clay mix.

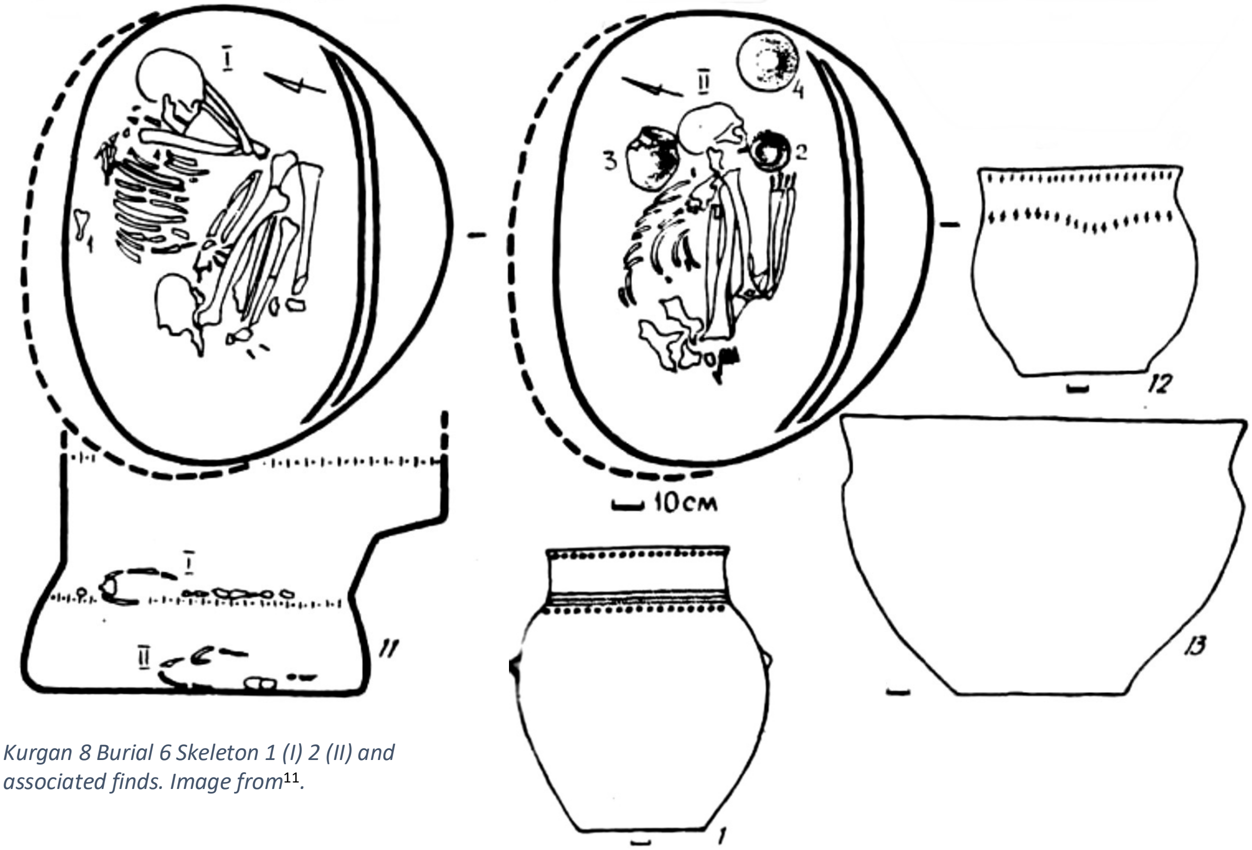

*Kurgan 8 Burial 6 Skeleton 1 (I) 2 (II) and associated finds. Image from*[11].

## I12706

Mau8, Kurgan 9, Burial 2, female (mature), 3947-3519 calBCE, RE-adjusted (5444±19 BP, weighted mean[10]

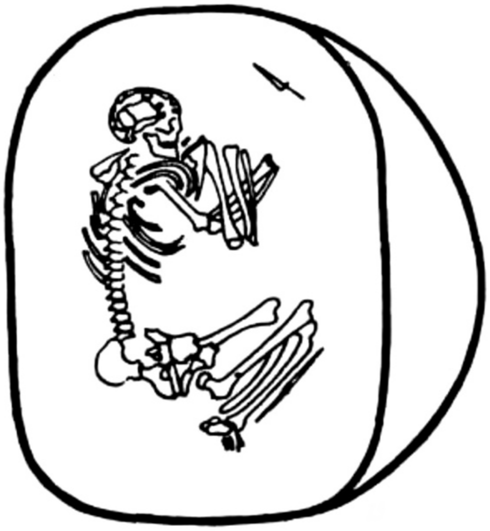

*Mayaky Kurgan 9 Burial 2. Image from*[11].

During the 1986 excavations of Kurgan 9, two burials were unearthed. Burial 2 was tentatively linked to Usatove due to the absence of burial inventory, while Burial 1 was associated with a different time period. Subsequent excavation efforts revealed two additional burials, designated as 3 and 4, with the latter being a double interment.

Burial 2 was identified as a discoloration located at the interface with sandy loam, approximately 0.45 m below the present-day surface. The rectangular pit exhibited rounded corners, almost resembling an oval shape, measuring 1.6 × 1.5 m along a southwest-northeast axis. The walls of the pit were discernible up to a height of 0.6 m, with a bottom depth of 1.05 m. Notably, the southeast wall of the pit featured a 0.5 m-deep and 0.3 m-wide step. The backfill material was compact, gray-brown, sandy-loamy, with sprinklings of lime. A mature adult in a contracted position lay on the left side with the skull oriented northeastward, arms flexed, and hands positioned in front of the face. Traces of raspberry-colored ocher were discovered on various skeletal elements, including the right shoulder bone, lower epiphysis of the right shoulder bone, pelvis, and legs. Additionally, patches of brown-colored organic material were detected on the long bones.

### Ma12

Kurgan 10, Burial 2, Skeleton 1, female (>60), 4033-3534 calBCE, RE-adjusted (5530±30 BP, BETA-432413[5]

### I1423/ I12705

MAJ1/Mau7, Kurgan 10, Burial 2, Skeleton 2, male (50-55), 3936-3372 calBCE, RE-adjusted (5390±30 BP, PSUAMS-7845)

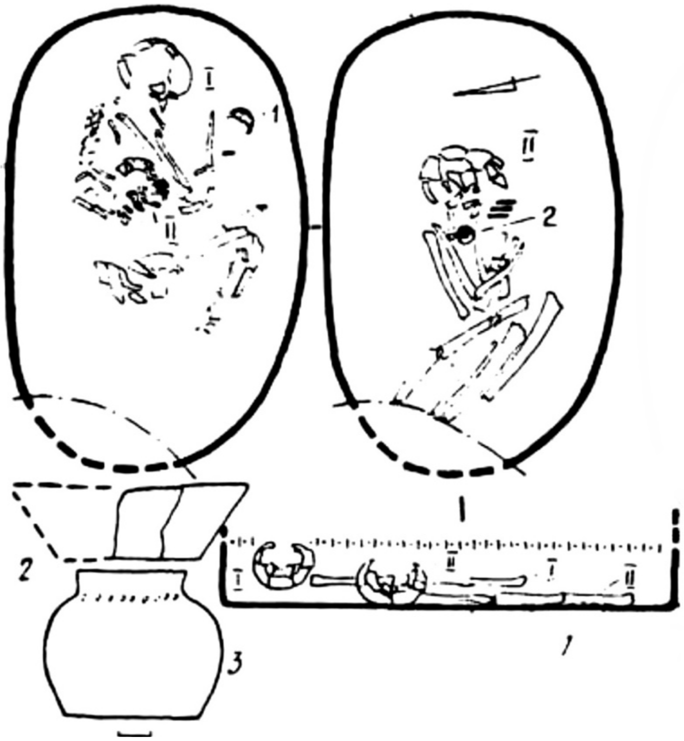

*Mayaky Kurgan 10 Burial 2. Image from*[11].

Kurgan 10 presented as a mound of chernozem soil measuring 0.15 m in height and approximately 20 m in diameter. Among the five burials designated as Usatove, two had incurred damage from pits dating back to the early centuries AD. These graves were filled with dense humus loam.

Burial 2 was housed within an oval-shaped grave pit, measuring 1.3-0.85 × 0.5-0.65 × 0.8 m. Within this burial site lay two individuals. Skeleton 1, identified anthropologically as a female aged over 60 years, was positioned on the left side, with Skeleton 2, a male aged 50–55 years, laid atop Skeleton 1, with the skull resting between the female's shoulder bones and chest. Both skeletons were oriented with their skulls facing east-southeast. The female's arms were folded, with her left hand positioned in front of her face and her right hand near her knees. The male's arms were bent, and his hands were raised to his face.

The inventory associated with Skeleton 1 included a fragment of a coarse bowl crafted from dark gray clay with sand. Skeleton 2's inventory comprised a brown vessel featuring imprints of a semicircular stamp, along with a loose ceramic mix containing grains of sand.

Low-resolution mtDNA typing conducted in the Molecular Archaeology Lab at GVSU, following the protocol detailed in [9].

##### *Molyukhiv Bugor*

Chygyryn district, Cherkasy region (49.134, 32.479)

Molyukhiv Bugor is a Neolithic-Eneolithic settlement and cemetery. It is located in the floodplain of the Tyasmin River to the north of the Novoselytsia village. The settlement is located on a sand dune, which has the appearance of an irregular acute triangle, stretched almost exactly from south to north, its length is about 90 m, and the width of the base is about 60 m. It was first excavated in 1950 by O.I. Terenozhkin and subsequently researched in 1955-1956 by V.M. Danylenko. Since 1992, the site has been excavated by T. M. Neradenko [1756].

I1424
MOB1, burial 4, skeleton 3, male, 3952-3657 calBCE (5020±50BP, OxA-17502)[15]

I1454
MOB3, burial 4, skeleton 2, 4000-3600 BCE

*Mykhailivka*
Kherson Region, Beryslavskyi District (47.315, 33.959)

The Mykhailivka multi-layer settlement is located 4 km south of the village with an eponymous name in the Beryslavskyi district of the Kherson region. It occupied two hills and partly a third, located across the ravine, as well as the marginal part of the adjacent plateau on the west bank of the Pidpilna River, a western tributary of the Dnipro. The slopes of the plateau were cut by deep ravines. On the high bank, two adjacent hills stood out, rising about 30 m above the level of the Dnipro. The largest (central) hill designated by researchers as Mykhailivka I came close to the bank of the river. One of the ravines separated the central hill from another, smaller one, on which the second settlement, Mykhailivka II, was located, at a distance of 50 m from Mykhailivka I. The area of the central hill was more than 0.5 ha, the height above the level of the Dnipro in the southeastern part was 23–25 m and almost 30 m in the northwestern part. Its greatest length is 100 m, the width on the north side is 50 m, south - 20 m.

The Mykhailivka archaeological site was studied by 1952-55, 1960, and 1963 expeditions of the Institute of Archaeology, National Academy of Science of Ukraine, led by E.F. Lagodovskaya and O.G. Shaposhnikova [20,176].

The total reconstructed area of the entire Mykhailivka settlement site is about two hectares. Geological cross-sections carried out in different places and directions of the site uncovered three cultural layers, of which the lower (third) was separated from the two upper (second and first) by a sterile horizon. The upper two layers, associated with the different phases of the Yamna archaeological complex, formed a continuous chronological sequence. Cultural remains occurred mainly in chernozem and humus loam. However, deepened ground dwellings and dugouts were also traced in the upper layers of light loam.

The upper layer of Mykhailivka contained artifacts from Yamna and Catacomb archaeological complexes of the Early Bronze Age. The lower horizon of the middle layer is dated to 3646-3348 calBCE (4710±80 BP, Ki-8012) and the upper horizon is dated to 3371-3026 calBCE (4519±53 BP, R-combine 4480±70 BP, Ki-8186, 4570±80 BP, Ki-8010). The lower layer of Mykhailivka is dated to 4247-3988 calBCE (5290±43 BP, R-combine5320±60 BP, Ki-9486, 5260±60 BP, Ki-9487, dwelling 4) and 3912-3540 calBCE (4921±53 BP, R_combine 4890±80 BP,

Ki-8011; 4945±70 BP, Ki-8182, dwelling 3) is associated with the Lower Mykhailivka archaeological culture [177].

### I32534

M1P1.6, Mykhailivka I, square VI, female, 3635-3383 calBCE (4755±25 BP, PSUAMS-10750)
A single molar from excavation square VI of the Mykhailivka I (central) settlement. Excavations 1952. Based on the $^{14}$C dating of the settlement layers, the specimen comes from the lower horizon of the second (middle) layer, associated with the proto-Yamna phase of the site's occupation.

##### *Liubasha Kurgan 2*

Novohryhorivka, Odesa Region, Mykolaivskyi District (47.343, 30.833)

Archaeological excavations near the Novohryhorivka village were conducted by the expedition of the Department of Archeology of the Northwestern Black Sea Region of the Institute of Archeology of the National Academy of Sciences of Ukraine. The "Liubasha" kurgan, or Kurgan 2, was located 2 km north of the village Novohryhorivka. The kurgan was located on the plateau of the east bank of the Tiligul River, at an altitude of 118 m above the river thalweg (167.7 m in the Baltic System of Heights). This location represents the edge of the highest watershed plane in the area. It stretches from the southeast to the northwest between Tiligul and the Gluboky Yar ravine. Other kurgans on the same section of the plateau are located 1.13 km southeast and 2.5 km northwest of Liubasha. In total, 17 burials were found in the Liubasha kurgan. Six belonged to the Yamna archaeological complex (Burials 2, 7, 8, 10, 16, 19), and three were of the Catacomb archaeological complex (Burials 3, 9, 11, 12, 15). Burial 5 belonged to the Sabatynivka group. Burial 14 was from the Sarmatian period and Burials 6 and 13 were of an indeterminate cultural affiliation. Periodically, processed flints with a patina, of a Paleolithic appearance, were found in the kurgan's embarkment[13].

### I12234

Liubasha, Kurgan 2, Burial 3, male, adult, 1499-1127 calBCE (3080±70 BP, Ki-11176)

Burial 3 (Babyne archaeological complex) was discovered 10.5 m southwest of the benchmark, at a depth of 1.5 m. The walls of the burial pit have not been traced. The burial was covered with a platform of planks, laid longitudinally directly on the body or, less likely, settled on the skeleton and bottom without any intermediate layer. There were five or six planks on the platform, the ends of three have been preserved. The semicircular bark (unpeeled) side was facing down. The interred (adult) lay contracted on the left side, with the head to the east. The legs were strongly bent at the knee and hip joints, the left knee was pulled towards the chest. The arms were also bent, the hands were in front of the face. Bone tissue was brittle, grey-blue in color on the limb bones, possibly as a result of thermal exposure. The level at which the skeleton was found was distinguished by a greenish clay coating with one well-preserved smooth edge, interspersed with coals and light firing. Outside the coating, traces of three pits with a diameter of 2–3 cm with wood decay were found, possibly from the racks of some then a light canopy. The radiocarbon age of the burial was established from fragments of bones of the lower extremities.

## I16674

Liubasha kurgan 2, Burial 15, male, 2434-1943 calBCE (3740±70 (Ki-11201)

Burial 15 (Babyne archaeological complex) was situated 24 m southeast of the central benchmark, at a depth of 4.43 meters. The oval pit, measuring 1.2 by 0.8 m, was filled with chernozem and oriented in a northeast-southwest direction. A disarticulated skeleton was located in the middle of the pit, densely folded in a compact "package". The skull was positioned in the southwestern part of this arrangement, resting atop the cervical vertebrae in an upright position before being crushed; the occipital portion of the skull was absent. Amidst the central cluster were the ribs, with one arm bone beneath them. Adjacent to the eastern wall, the bones of the arms were neatly folded, followed by the legs and then the pelvis atop them. Close to the skull at the pit's base, grains of pink ocher and a flint flake were discovered. The burial inventory comprised a light brown flint flake, lacking patina.

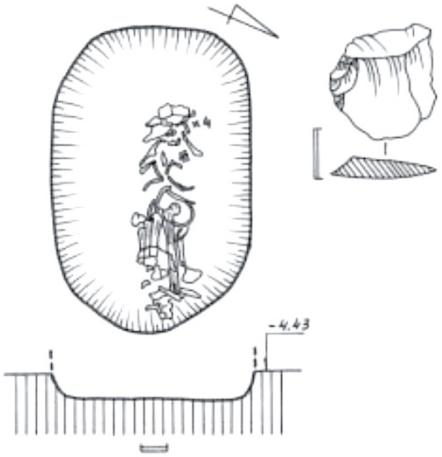

*Liubasha Kurgan 2 Burial 16. Image from*[13].

## I12233

L2.16, Liubasha Kurgan 2, Burial 16, male, adult, 3074-2888 calBCE (43330±35 BP, PSUAMS-7790)

The burial chamber of Burial 16 (Yamna archaeological complex) was 0.88 m wide, was covered by timber planks. In the decay under the skeleton, longitudinal and transverse fibers (stems of some plants) were clearly visible. The interred lay on the back, head to the west, arms extended along the body. Legs were preserved above the knees. It could be determined that they were bent with the knees to the right. The skeleton was poorly preserved. To the right of the skull, a lump of dark ochre, egg-shaped, about 3 cm high, was found.

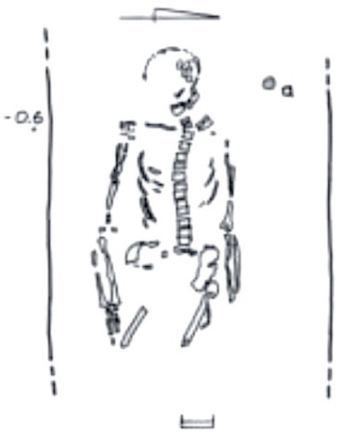

*Liubasha Kurgan 2 Burial 16. Image from*[13].

#### *Odesa Kurgan*

Odesa (46.488, 30.721)

The Odesa Kurgan was studied in 1912-13 by M. F. Shkadyshko and A. V. Dobrovolsky. A detailed analysis of kurgan’s stratigraphy was published by V. G. Zbenovich and A. M. Leskov in 1965. Analysis of the stratigraphy of the Odesa kurgan shows the presence of the main and three additional embankments. The main mound was erected over Burial 11. Burial 10 and Burial 7, burials in stone cists, later destroyed, were subsequently added into the main mound.

Yamna burials located both in the center and along the kurgan's circumference are probably associated with two subsequent embarkments. Catacomb and Babyne burials, as well as burials of the late Bronze Age period, were subsequently added into the formed mound of the kurgan[178].

I11850

ODK1, Odesa kurgan, Burial 10, female, 2874-2630 calBCE (4150±25 BP, PSUAMS-11224).

Burial 10, possibly of Catacomb archaeological complex, was lowered into the main mound of the kurgan. The burial pit was covered by a stone slab. The interred was in a contracted position on the back with legs falling to the right side. A stone circle was arranged around the burial.

*Ogrin-8* (Igren-8)

Dnipro, Dnipropetrovsk Region (48.444, 35.116)

The archaeological site of Ogrin-8 was settled since the Mesolithic. The earliest occupation phase at the site is connected with the Kukrek archaeological group and dates to the early Holocene [179]. Ogrin' 8 ground multi-layer cemetery (Neolithic-Late Bronze Age) was located on the northern side of the Ogrin' Peninsula at the confluence of the Dnieper and Samara rivers. Excavated in 1932 by M. O. Miller. The next phase of excavations at the cemetery took place in 1945-46 by A. V. Dobrovol'sky. In 1974-76, 1978, and 1986, excavations at the cemetery were carried out by D.Ya. Telegin [1801].

I2108

Igren-8 9, Burial 20, infant, female, 4340-4178 calBCE (5415±25 BP, PSUAMS-8220)

I1924/ I27935

Igren-8, bag 11, burial 24, female, 30-40, 3971-3802 calBCE (5105±25 BP, PSUAMS-8219).

Excavations 1978.

I27930

Igren 8-74 - Individual 3a, male, young adult, 4400-4000 BCE.

Excavations 1974-78. One of the two adult individuals in a double burial at the depth of 0.9-1.0 m, on the back with raised knees, ESE orientation, ochre, and three flint blades between the shoulders.

*Revova Kurgan 3*

Odesa Region, Shyriaivskyi District (47.268, 30.321)

In the vicinity of the village of Revova, two kurgans, 3 and 4, were investigated in 2003 by the expedition of the Department of Archeology of the Northwestern Black Sea Region of the Institute of Archeology of the National Academy of Sciences of Ukraine. The kurgans were located on the east side of the Velykyij Kuyalnik river valley, at an altitude of 84 m above the thalweg of the river (125 m in the Baltic system of heights). They occupied the upper part of the slope of the southwestern exposure, gently descending to the river, which is 1.5 km from the kurgans.

Kurgan 3 had a complex organization. It started as an Eneolithic ritual complex (sanctuary). Prior to building the initial sanctuary, the topsoil was cleared and the ground leveled, leaving the center of construction rising slightly compared to the rest of the surface. Around this elevation, a semicircular ditch about 12 m in diameter was dug. The earth from the ditch was used to build a platform, 12 m in diameter, and 1 m high. It appears that the pit for the main Eneolithic burial 19 was already laid out prior to erecting the platform. In the next construction step, the platform was tiled around with limestone, quarried, apparently, 4 km away from the site. Limestone was sometimes laid in two or three layers; the largest pieces were found surrounding the eastern side of the platform. The final stone mosaic resembled in its outline the shape of a turtle. The whole construction was eventually covered with soil, making it difficult to tell whether it was intentional or was the result of a natural process.

Two ritual pits/postholes 9 and 11 to the west of the primary mound likely belonged to the same Eneolithic complex. According to stratigraphy, the complex was completed by an addition of loam, or mound 2, about 18–22 m in diameter.

Burials 3 and 16 (Yamna archaeological complex) were lowered into the central part of the mound from the level of mound 2, later almost completely destroyed by digging. Burials 3 and 16 were covered with large stone slabs (stelae), two in each burial. There was a central cover slab over burial 3, which was likely destroyed by a robber trench. The slabs from burials 3 and 16 were possibly part of a dolmen erected on top of burial 19, and subsequently re-purposed in later burials [181]. The original dolmen's capstone may have been used as a central slab in burial 3.

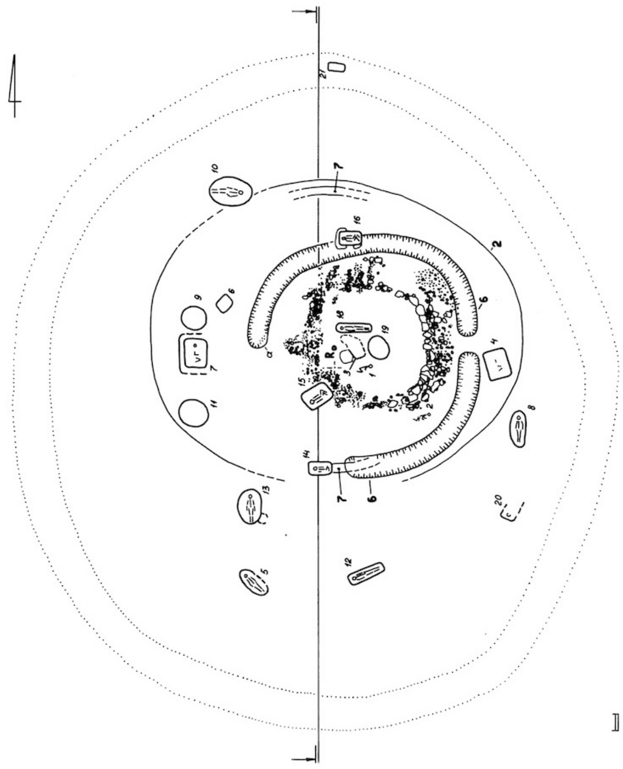

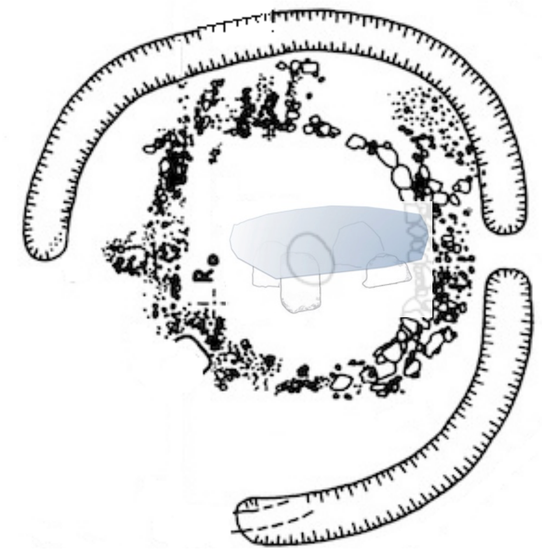

Reconstruction of a dolmen over the central part of the Revova Kurgan 3. Drawing by S. Ivanova and A. G. Nikitin.

*Revova Kurgan 3. Drawing by S. Ivanova.*

## I16668

REV3.10, Revova Kurgan 3, Burial 10, male, 2700-2000 BCE

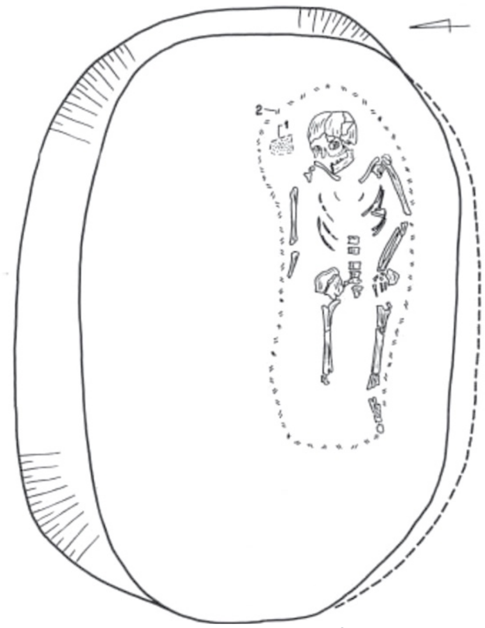

*Revova Kurgan 3, Burial 10. Image from[13]. 1, stone chips, 2, remnants of a burial mat.*

Burial 10 (Catacomb archaeological complex) was found 11 m northwest of the benchmark, at a depth of 1.5 m. It was identified by a dark oval-shaped spot. The dimensions of the grave pit at the top were 2.6 × 1.8 m, at the bottom – 2.5 × 1.68 m, the traced depth is 0.93–0.95 m (2.43–2.45 m). The southern wall was made in the form of a lining, the northern wall was beveled towards the bottom at an angle. Stone chips were found at the bottom of the burial chamber. The interred was located closer to the southern wall, in the extended supine position on the back, with the head oriented to the east. The right arm extended along the body, the left arm was bent at the elbow, the hand covering the left hip joint. All that was left of the legs were the femurs and part of the left tibia. There was a faint decay from an organic mat under the skeleton.

## I7928

R3.16, Revova Kurgan 3, Burial 16, 2885-2501 calBCE (4135 ± 60 BP, Ki-11059 [13]

Burial 16 (Yamna archaeological complex) was found at a distance of 7 m to the north of the central benchmark, along the edges of limestone anthropomorphic stelae protruding through the ground. The lower faces of both stelae were not processed. The eastern stela, 0.9 × 0.6 m and 20–25 cm thick, had stocky proportions. Half of the slab was outlined by a semicircular protrusion indicating the head (partially knocked down). The western stelae was oblong (1.2 × 0.8 m, thickness 0.12–0.2 m). The head on the anthropomorphic silhouette of this slab was displaced from the middle axial line.

The burial chamber was found under the stelae at a depth of 0.8 m from the central benchmark. It was a rectangular pit with rounded corners, 1.4 x 0.75 m, filled with dense loam. The buried (adolescent) was lying on the back, with his head to the west. The arms were slightly bent at the elbows and laid along the body, palms down. The legs were bent and placed with the knees up; long bones are partially preserved *in situ*. The skull was covered with lilac-colored ochre, the hands – with a scarlet-colored ochre. To the right of the skull was a layer of chalk and a lilac ochre pellet in the shape of an egg, about 3 cm long. To the left of the skull was a piece of chalk or lime. The area under the skull was sprinkled with grains of chalk. The bottom of the pit was smeared with light clay.

The specimen failed amplification at Reich's Lab. Low-resolution mtDNA typing conducted in the Molecular Archaeology Lab at GVSU, following the protocol detailed in [9].

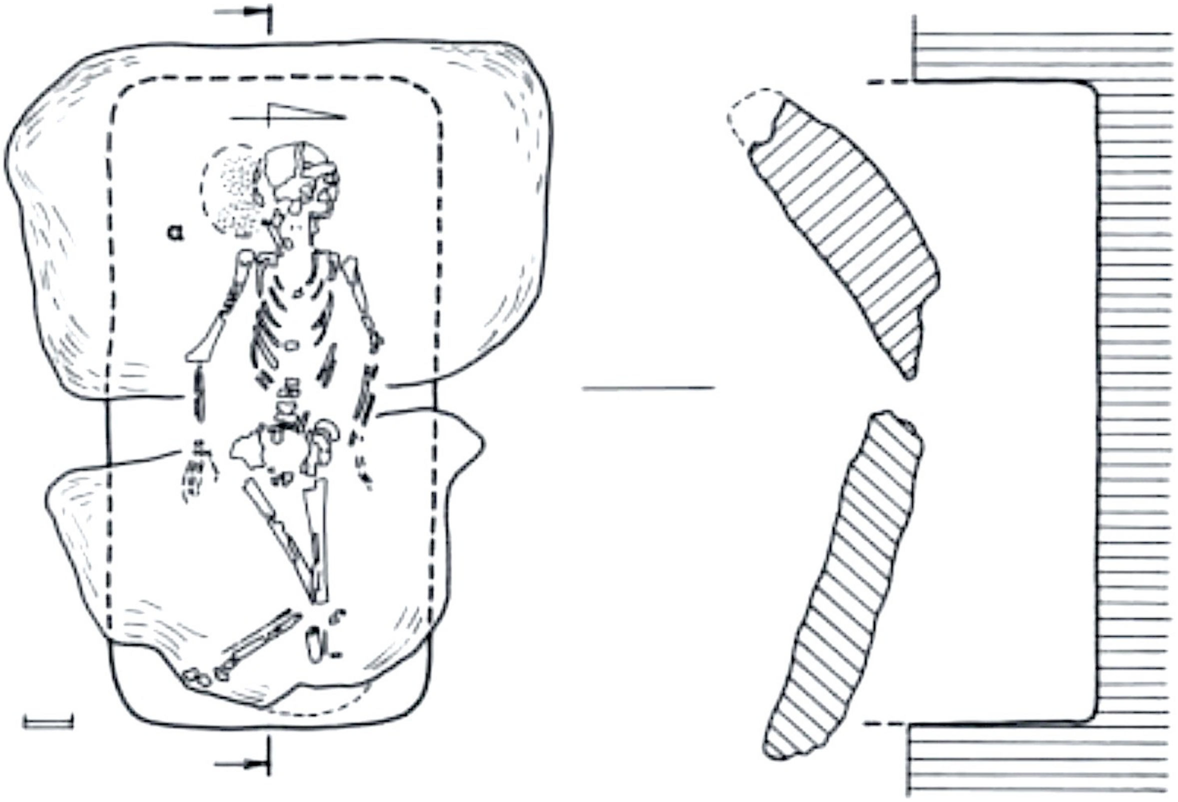

*Kurgan 3 Burial 16. Image from* [13].

## I7929

REV-3.19, Revova Kurgan 3, Burial 19, male, 3711-3639 calBCE (4905±20 BP, PSUAMS-4763)

Burial 19 was the main burial in the kurgan. The burial was found at a distance of 2.1 m to the east-northeast of the central benchmark, at a depth of 1.05 m. Scattered and mostly fragmentary human bones lay throughout the fill area in the eastern part of the pit. The layer-by-layer analysis of the remains suggests that the state and placement of the bone remains were consistent with the rite of secondary burial, perhaps with an imitation of the sitting position of the buried, possibly wrapped in cloth. The upper level of the bone “package” was occupied by the remains of the skull (parietal bones) and cervical vertebrae, the middle level consisted of the ribs, and the lumbar vertebrae and the sacrum lay in the lower tier. The bones of the left leg were completely missing, including the foot. The tibia of the right leg, also without a foot, was in a vertical position, while the femur lay horizontally with its head turned towards the main body of bones. The vertical position was occupied by the radius of the right hand. Other bones of the arms, including the hands, were absent. The zygomatic bones were located at the bottom of the pit. A patella rested somewhat away from the main assembly. Some fragments of the skull were painted with ochre. At the bottom of the burial chamber, near the bones, grains of scarlet ochre were noted. The dust from burnt timber planks extended throughout the chamber floor.

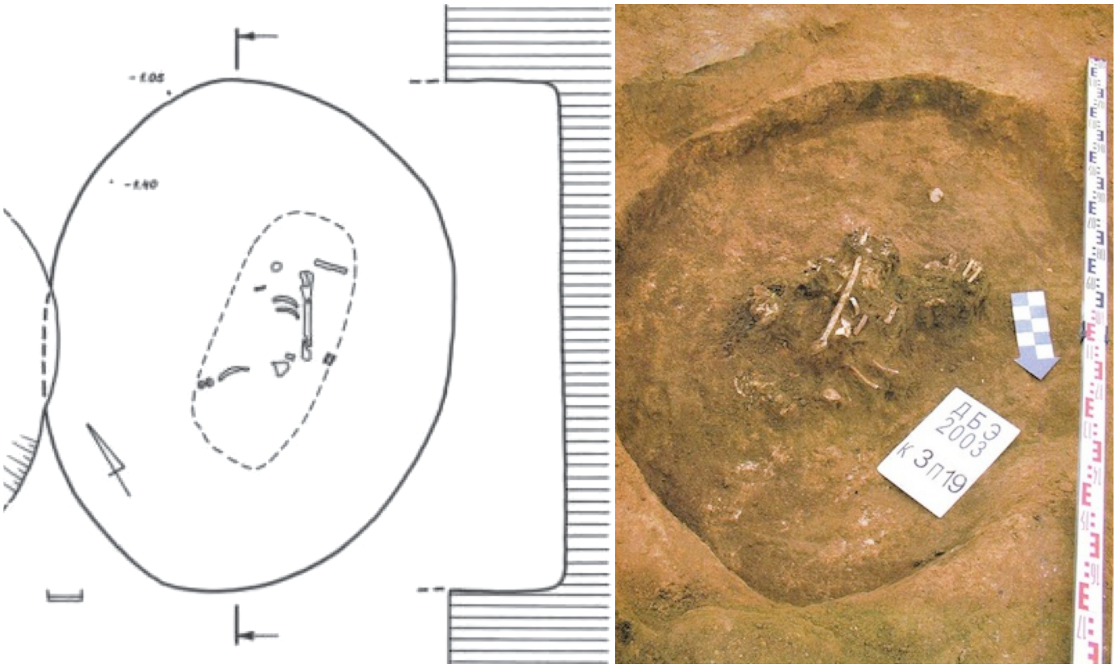

*Revova Kurgan 3, Burial 19. Image from*[13].

#### *Shakhtar (Shevchenko) kurgan group*

Shakhtar (Shevchenko) Hamlet, Dnipropetrovsk Region Nikopol’ District (47.739, 34.179)

Kurgans 28 and 29 of the Shakhtar (Shevchenko) kurgan group, studied by the expedition of the Institute of Archeology of the Academy of Sciences of Ukraine under the leadership of S.V. Polin in 2003, were situated on the first above-floodplain terrace of the east bank of the Solena River, 200 m from the shore edge, 500 m east of the hamlet. The kurgans were 2.8 and 4.5 m high, respectively, located at a distance of 50 m from each other. In terms of size and relative position, these mounds represent a characteristic type of kurgans called "twins".

At the time of excavations, the upper half of Kurgan 28 was covered with turf, and the lower part of the slopes was plowed. The embankment had an asymmetrical shape: the northern slope was noticeably steeper than the southern one, which was greatly stretched and deformed by later diggings. The floors of the embankment along the circumference within a radius of 10–12 m from the top of the kurgan were cut by long-term plowing of the field. The height of the kurgan from the level of the field surface from the north was 2.8 m, from the south - 2.5 m. The apparent diameter of the embankment was 42 m. The kurgan contained 17 burials of the Yamna and Catacomb archaeological complexes as well inlet burials dated to later chronological periods.

I3141

Shakhtar (Shevchenko) Kurgan 28, Burial 12, female 35-45, 3300-2700 BCE[8,96]

Burial 12 (Yamna archaeological complex) was situated 12.8 m south-southwest of the reference point, in a rectangular pit with a wide ledge along the NW – SE axis, 1.2 m deep from the mound's surface. The ledge displayed white decay with plant fiber imprints (reeds?) across the pit. The burial chamber's bottom, 1.25 m below the ledge, sloped into the mainland by 0.95 m, narrowing slightly. Upper dimensions: length 1.4 m; center width 0.83 m, SE wall 0.75 m, NW wall 0.9 m. Bottom dimensions: length 13.8 cm; center width 0.78 m, SE wall 0.73 m, NW wall 0.7 m. Filling: loose mixed soil with reed floor decay. Matrix walls exhibited vertical grooves from digging tools, up to 2 cm wide.

The interred was a female, contracted on her back, with her head to the southeast. Her legs, knees bent upward, tilted to the right, her knees rested against the northeastern wall of the chamber. The arms were extended and slightly away from the body. From the cervical vertebrae along the right humerus stretched a low necklace of bone piercings, ending at the elbow with a hammer-head pin lying perpendicular to the arm, with the head on the elbow joint. In the middle of the ulna bones, on the inside of the right arm, opposite the pelvis of the interred, there was the wing of a pelvis of a fetal child. To the left of it, several more very small bones of a skeleton of an infant were located. At the north-eastern wall, under the knees, there was a cracked cake made of a dense but fragile substance of violet color (ocher?), about 10 cm in diameter and 4 cm thick. The left foot had two hooves (sheep-goat?), next to them was one small animal bone. At the bottom there were traces of a white plant decay covered by a layer of red decay 0.5–1.0 cm thick (from felt?), covered by brown decay of a similar thickness in the pelvic area.

The necklace comprised eight tubular bone piercings with screw threads and two smooth beads from bird or small animal bones. Thread dimensions: length 2.7–5.7 cm, width 1.1–1.3 cm. Bead dimensions: 1.1 x 1.4 cm, thickness 0.8 cm; 1 x 1 cm, thickness 0.4 cm. The hammer-head pin had a cylindrical shaft, T-shaped head with a hole, shields at ends, length 21.8 cm, base diameter 1.1 cm, head length 3.9 cm, shield diameter 2.1 cm, hole diameter 0.5 cm.

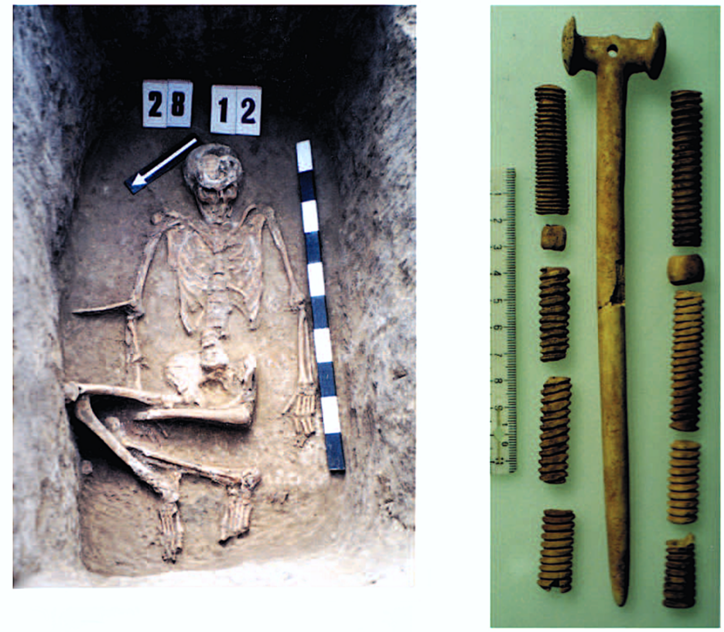

*Shakhtar (Shevchenko) Kurgan 28, Burial 12. Image from [96].*

#### *Soldats'ka Slava Kurgan*

Baranove, Odesa Region, Ivanivskyi District (46.938, 30.407)

Archaeological excavations near the village of Baranove were conducted in 1990-1991 by the expeditions of the Odesa Protective Archaeological Center under the Ukrainian Society for the Protection of Historical and Cultural Monuments. The "Soldats'ka Slava" kurgan was located on the lands of the Baranove village council of the Ivanovsky district. The kurgan was erected on the watershed between the Maly Kuyalnik River and the Krivaya arroyo, which flows from the left into the valley of the Svinaya River, a tributary of the Khadzhibey Estuary. The top of the watershed is a flat steppe plateau, stretched from north to south and southeast and marked in these directions by tall kurgans. According to measurements before the excavations, the height of the "Soldats'ka Slava" kurgan was 2.55 m. The diameter of the base of the embankment was 53–55 m. In total, 8 burials of the Bronze Age were found in the kurgan, including three of the Yamna archaeological complex (Burials 3, 4, 10), one each of the Kemi-Oba type (Burial 9), and the Babyne archaeological complex (Burial 2), two presumably Catacomb archaeological complex (Burials 1, 11), and Burial 5 without a clearly defined cultural affiliation.

### I12005

SL-10, Soldats'ka Slava Kurgan, Burial 10, male, 3300-2500 BCE

Burial 10, of the Yamna archaeological complex, was arranged in a 2.25 × 1.40 m rectangular burial chamber. The chamber had a depth of 1.3 m from the ledge (5.65 m) and was covered by bark at its level. In the eastern part of the chamber, in the filling, at a depth of 0.8 m, a 0.8 × 0.5 × 0.15 m limestone cinder was found. The interred was laid contracted on the back, head to the west-south-west, legs bent at the knees and falling to the left, the arms were extended along the body. A thoroughly polished stone axe with no traces of a hilt was placed at the left shoulder. A pair of silver pendants rested at the temples.

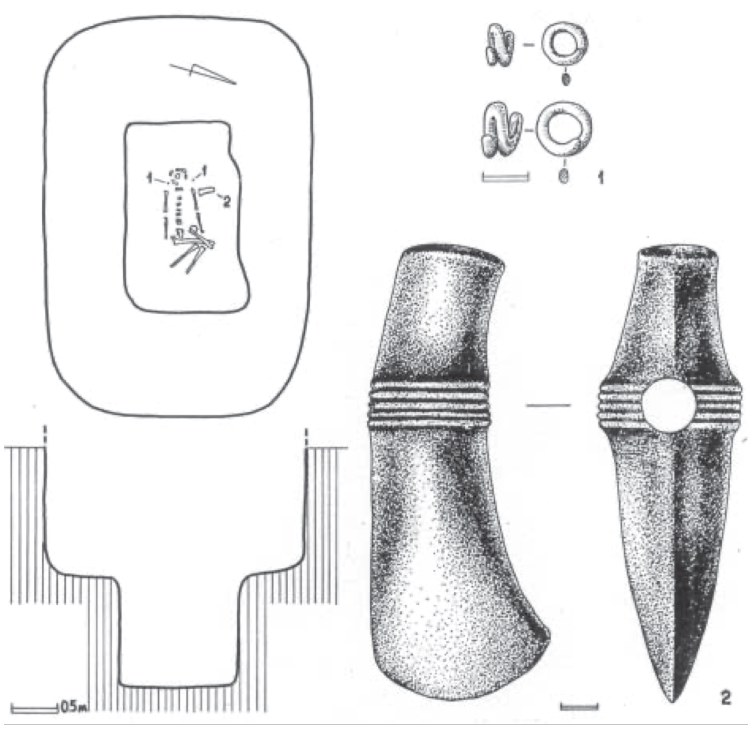
*Soldats'ka Slava Kurgan Burial 10. Image from [13].*

#### *Sychavka Kurgan*

Odesa Region, Kominternivskyi District (46.643, 31.101)

The expedition of the Department of Archeology of the North-Western Black Sea Coast of the Institute of Archeology of the National Academy of Sciences of Ukraine conducted a study of a kurgan near the village Sychavka (former Malaya Aleksandrovka) in 2010.

Three mounds and a cromlech were found in the kurgan. The construction of the kurgan began in the Eneolithic era. Most likely, the anthropomorphic stelae that were used in Burial 15 of the Yamna archaeological complex came from the cromlech. The kurgan was used in the Bronze Age by the tribes of the Yamna archaeological complex (Burials 9, 10, 13, 15, 19), the Catacomb archaeological complex (Burial 12), the Babyne archaeological complex (Burials 17, 18, 21, 23), and the Sabatynivka archaeological group (Burial 6).

## I12843

S15, Sychavka Kurgan, Burial 15, male (40-45), 2847-2501 calBCE (4087±19 BP, weighted mean of 3960±80 BP (Ki-16610)[28]

Burial 15 (Yamna archaeological complex) was discovered 5.2 m to the southeast of the central benchmark at a depth of 0.68 m behind a stone embankment. The part of the embarkment consisted of a large limestone slab, oriented along the west-east axis. The 1.48 × 1.78 × 0.95 × 0.28 m slab was sub-rectangular in shape. The lower side of the slab is slightly wider than the upper side. It was surrounded by masonry on three sides. Human rib fragments as well as a fragment of a stucco vessel were found under the main slab.

The skeleton of a 45-50-year-old male was lying in a very contracted position on his left side with his head tilted to the west. The left arm was extended to the knees, the right arm was bent at the elbow and the wrist was probably placed by the face (only the epiphyses of the ulnar and radial bones have been preserved). A stucco vessel was placed by the head. A stone grinder was located near the elbow. Ochre was not noticed. Under the pelvis, the floor was coated with green clay.

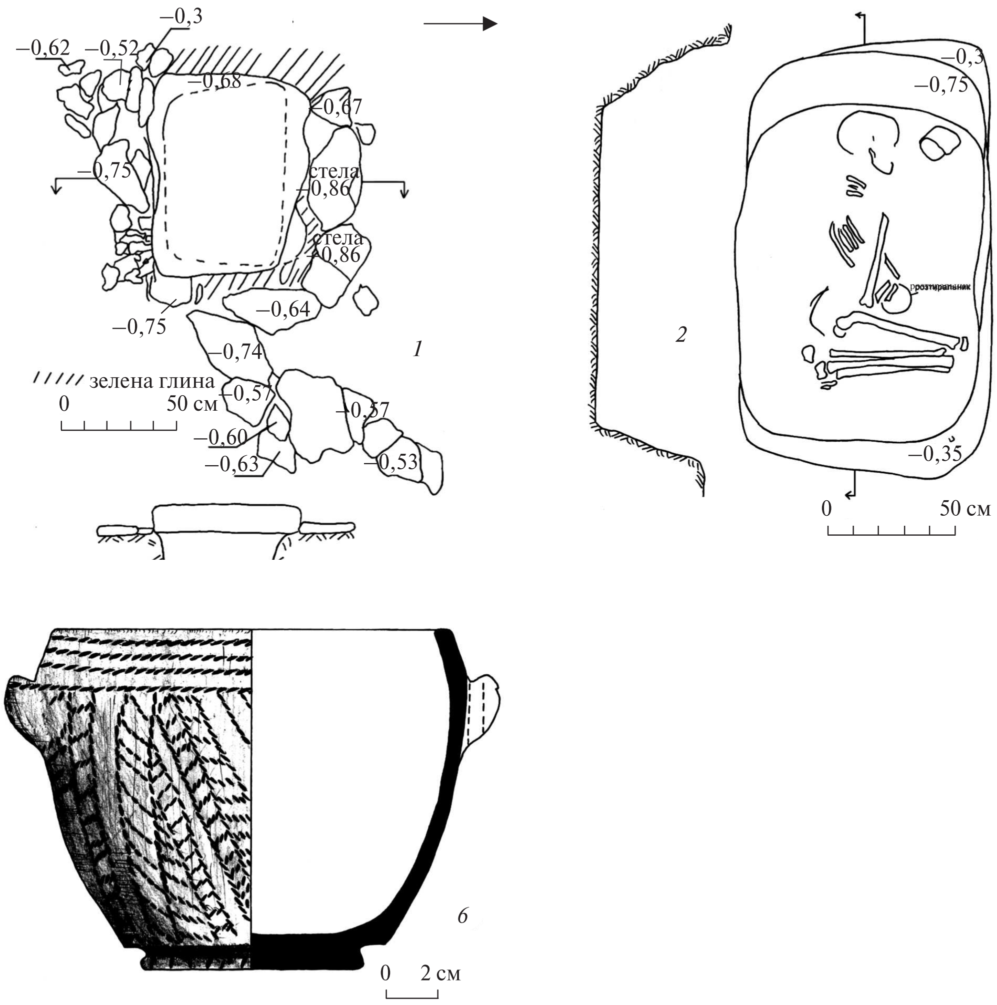

*Kurgan Sychavka Burial 15. Images from* [28].

#### *Vapnyarka Kurgan 4*

Odesa District, Odesa Region (46.576, 30.889)

Kurgans near the village of Vapnyarka comprise a burial ground known since the middle of the 19th century. Kurgan 4 was excavated by a team from the Institute of Archaeology, National Academy of Sciences of Ukraine, in 2008. The kurgan contained 20 interments. Two belonged to the Eneolithic period, 15 – to the Bronze Age, one was Scythian, and two others came from an undetermined period [182].

I12617

V4.3, Vapnyarka Kurgan 4 burial 3, female, 12-14, 4089±24 (weighted mean[10] of 3960±70 (Ki-15230)[22], and 4105±25 BP (PSUAMS-10808)

Burial 3 belonged to the Catacomb archaeological complex. The skeleton was laid on the back in a slightly bent position, knees bent to the left, head facing southwest. Arms were slightly bent at the elbows, lying along the torso. A cluster of red ochre 3 × 4 cm was found by the right shoulder. A Catacomb pot was placed behind the head.

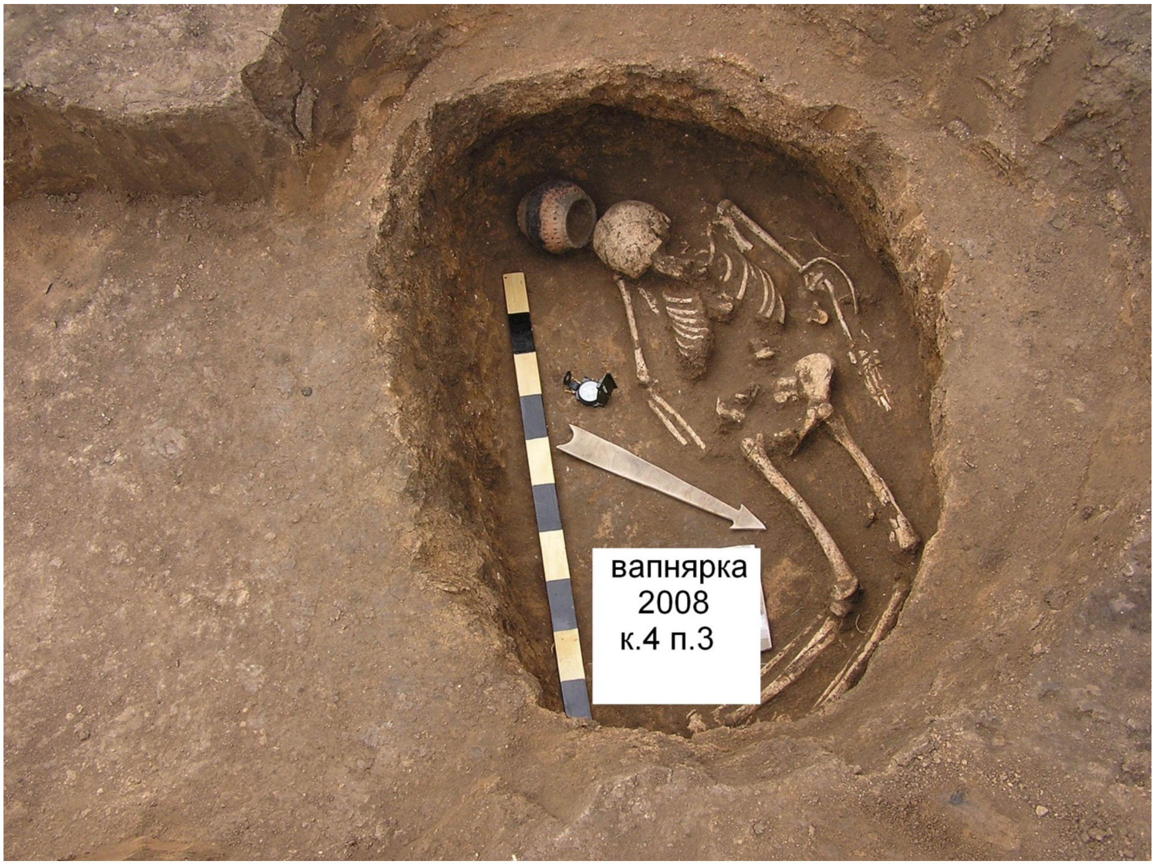

*Vapnyarka Kurgan 4 Burial 3. Photo by S. Ivanova, 2008.*

I12618

V4.4, Vapnyarka Kurgan 4 burial 4, female, 3341-3031 calBCE (4480±30 BP, BETA-647661)

Burial 4 (main burial, Kvityana/post-Mariupol archaeological group) was discovered at 2.8 m north of the central benchmark at a depth of 0.45 m. The pit was covered transversely by three large polygonal, torn layer by layer, rough-hewn limestone slabs that split and sank into the grave in ancient times (some of the slabs were almost vertical). The length of the

*Vapnyarka Kurgan 4 Burial 4. Image from[13].*

arrangement was 1.6 m, and the width was from 0.5 m (in the west) to 1.0 m (in the east). The dimensions of the slab fragments: 0.80 × 0.65 × 0.15 m, 0.62 × 0.46 × 0.21 m, 0.38 × 0.38 × 0.18 m. The grave was sub-trapezoidal in plan with rounded corners ("bean-shaped"), tapering to the feet of the buried. It was 1.95 m long (2.0 m along the bottom), 0.95 m wide at the head, 0.70 m at the feet, 0.75 m in the middle part, 0.32 m deep from overlap (1.07 m from the central benchmark). It was filled with mixed soil (chernozem interspersed with fine limestone), identical to the structure of the first embankment. The edges of the pit were slightly beveled. The eastern edge was damaged during the deposition of Burial 9.

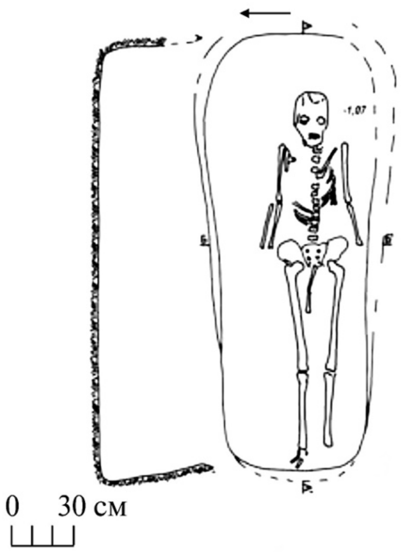

The skeleton was lying straight on the back with the head to the east. The radius of the left arm rested between the femurs. The right part of the chest was damaged by the stones of the floor. The phalanges of the toes were partially preserved. The skeleton, particularly its frontal part, was weakly colored with bright red ocher. Ocher concertation was near the ulnar epiphyses on the humeral bones and on the upper part of the femoral bones. The heel bones were colored more intensely. At the bottom, near the chest, there was an ocher spot measuring 2 × 2 cm. The burial contained no inventory.

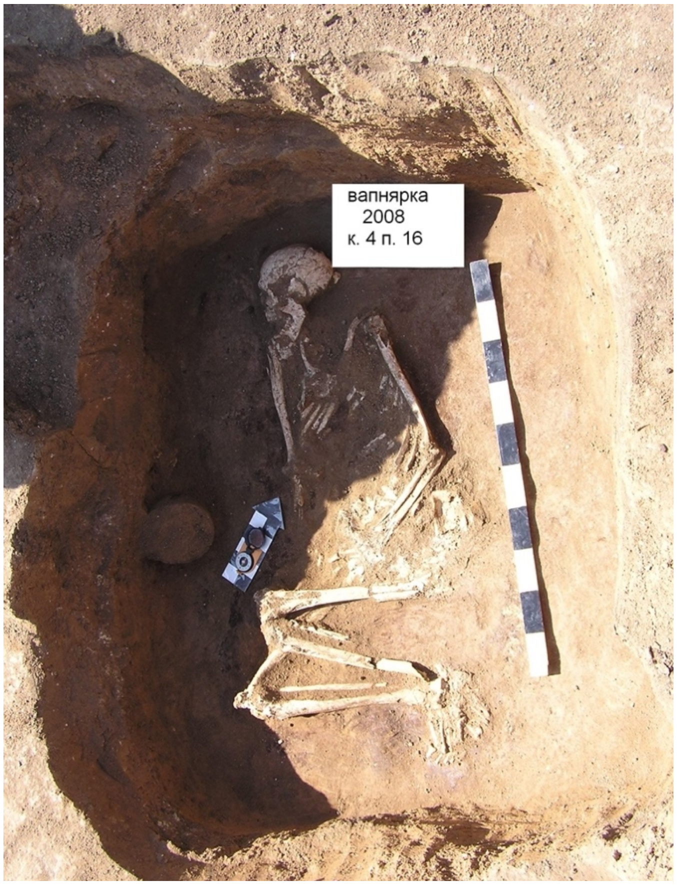

*Vapnyarka Kurgan 4 Burial 16. Photo by S. Ivanova, 2008.*

### I12619

V4.16, Vapnyarka Kurgan 4, Burial 16, male (50-55), 2876-2631 calBCE (weighted mean of 4050±60 BP (Ki-15014)[22] and 4180±30 (BETA- 441245)[5])

Burial 16 (Yamna archaeological complex) was located at 4.6 m to the west of the central benchmark at a depth of 0.93 m (stone floor level). The grave was covered transversely with three large polygonal limestone slabs torn in layers and slightly hewn. One of them, measuring 1.2 × 0.8 m, covered the pit in the headland and lay almost horizontally, partly on shoulders or a ledge. The second slab, 0.7 × 0.7 m, was located in the center of the grave and laid almost on top of the skeleton, destroying it. The third slab, 0.6 × 0.6 m, laid tilted at the eastern end of the pit. The slabs were partially coated with green clay. The grave was trapezoidal in plan with dimensions 1.45-1.50 × 1.00 (at the head) — 1.30 m (at the feet), 0.55 m deep from the floor. The northern corner was beveled to the bottom. In the north-western corner, a step (or part of the steps) measuring 0.8 × 0.5 m and 0.2-0.3 m deep was found, on which the floor slab was laid. The pit was filled with re-deposited yellow continental loam. At the bottom, brown rot from an organic mat remained.

The massive skeleton was laid contracted on the right side with the head to the northwest. The right arm was straight, the left was bent at the elbow, and the hands were near the knees. The legs were bent at a right angle to the body. A fragment of a goat's horn was found near the skull. A small fragment of a bronze artefact (possibly a string from an ankle bracelet, as there were traces of green oxide on the large tibia near the ankle), was located. A truncated-conical jar-shaped vessel with two horizontal handles-stops crossed by through vertical holes, containing a small subtriangular stone and a lump of green clay, was located near the left ankle, under the western wall, opposite the right elbow. The vessel, 15.5 × 16.5 × 8.4 cm, was decorated with a cord ornament consisting of three horizontal lines under the crowns and a vertical "herringbone" on the body. The surface was pink, with gray spots, partially peeled off.

##### *Verteba Cave*

Bilche Zolote, Chortkiv District, Ternopil Region (48.7889, 25.8715)

The Verteba Cave archaeological site is described in [6,7,9,183]

DNA data on I1926, 1V1a-H1, I2110, 4.V4a-H4, I2111, 5.V5a-H5, I3151, 6 V6a-H6 originally published in[8,9]

### I7584

VCS1.CL.05, Excavation Site 1, Individual 1, calcaneus (left side), female (subadult, >6), 3708-3543 calBCE (4875±25 BP, PSUAMS-10769)

### I7923

RL2.05, Excavation Site 2, Individual 1, radius (left side), male (adult), 3789-3649 calBCE (4955±30 BP, PSUAMS-8692)

### I7586

VCS1.TBL.05, Excavation Site 5, Individual 2, cuboid (left side), female (adult), 3766-3642 calBCE (4920±25 BP, PSUAMS-9527)

### I1929

9 V9a-A22.2, Verteba Cave Site 6, Individual 2, female, 4910±400 BP (Ki-14308[6]), 3900-3500 calBCE. Archaeological context was used for calibrated date due to a large margin of error in the existing $^{14}$C date.

### I7921

A22.P3, Excavation Site 6, Individual 3, female (subadult, >6), third phalanx, 3900-3500 BCE

I13064
VRT-VNWL3, Excavation Site 7, left third metacarpal/ VRT-VSE.22N, right third metacarpal, Individual 2, male (adult), 3648-3528 calBCE (4820±30 BP, PSUAMS-7807)

I2109
3 V3a-H3, Excavation Site 7, Individual 3.18.1, male (18-22), cranial bone, 3709–3537 calBCE (4863±33 BP, OxA-26202[7]

Additional analyses undertaken on the material from Excavation Site 7 revealed that the following three specimens came from the same individual:
I1926, 1V1a-H1, male (35-45), cranial fragment, 3769-3648 calBCE (4938±22 BP, weighted mean of 4890±30 BP, Beta-432808, and 4985±30 BP, OxA-25991) [7–9]
I7922, TV2.6.1, individual 1, thoracic vertebrae
I13066, VRT-VTV3, thoracic vertebrae

*Vynohradne Kurgan 3*
Tokmak District, Zaporizzya Region (47.221, 35.57)

The kurgan group 0.5 km to the west of the village of Vynohradne was located on a high promontory of the right bank of the Molochna River. Kurgan 3 was located at the highest point of the promontory, rising 80 m above the floodplain. The kurgan was excavated by S. Pustovalov and Y. Rassamakin in 1982.

The height of Kurgan 3 was 8.05 m, the diameter was 100 m. The mound had the shape of a truncated cone with a wide and relatively flat square on the top. Under the mound, a unique sanctuary of complex construction was discovered. Under the general embankment, 49 burials of the Eneolithic, Yamna, Catacomb, Babyne and Zrubna cultures were discovered[122].

The embankment of Kurgan 3 had a complex structure. Under the sod surface of the kurgan there were eight separate mounds, which were later united by a multilayered common embankment. A sanctuary was erected on top of the eighth embarkment.

Several stages can be traced in the functioning of the sanctuary. Initially, a 0.7 m × 1.5 m altar made of large clay blocks was erected in the center of the kurgan. The altar was surrounded from the south and east and possibly from the north by a path covered crosswise with reeds. Fragments of animal bones were found on the path. The path was later covered with clay and the altar was enlarged in the northern direction. As a result, a square approximately 20 m in diameter was formed with an altar in the center. Radial embankments were built on the slopes of the eighth embankment during the first stage of sanctuary functioning.

During the second stage, the space between the radial embarkments was filled with sod chernozem blocks, forming a continuous platform of approximately oval shape, 35-45 m in diameter, with an altar in the center. Two ramp entrances were attached to the platform, one from the east, the other from the west. Each of the ramps had two entrances, opposite each other, one from the north and one from the south.

In the last, third stage of the sanctuary's existence, all the ramps were filled in and the kurgan expanded to the north. Only one 10 m-wide entrance from the east remained.

According to aerial photography, 13 straight radial dark lines (roads) ranging in length from 1,200 m to 4,566 m - extend from the kurgan into the steppe to the north, northwest, west, and southwest. These radial lines are crossed by several identical lines running from north to south. The reconstructed number of ramparts coincides with the number of road lines that radially approach the kurgan from the side of the plateau[122].

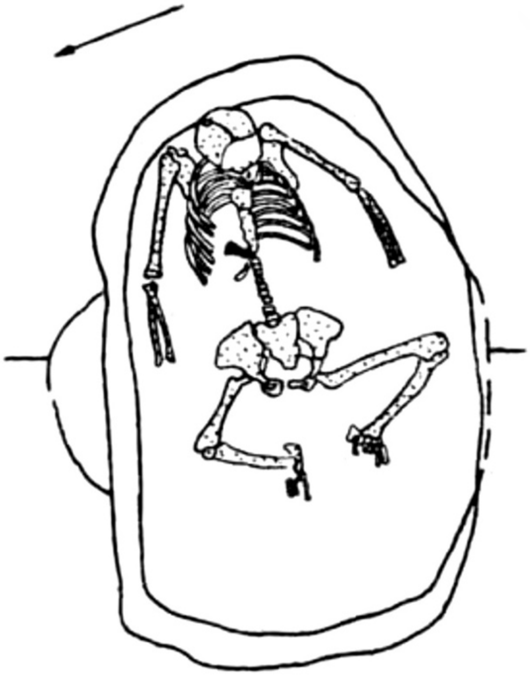

*Vynogradne Kurgan 3 Burial 15. Image from[19].*

### I1430

VIN1, Kurgan 3 Burial 15, male, adult, 4241-3951 calBCE (5230±60 BP, Ki-14726)

The main burial 15 was in the central of the eight mounds of Kurgan 3, 1.1 m from the ancient horizon. The burial chamber was surrounded by a ditch. The adult male at the bottom of the burial chamber was buried contracted on his back ("half-sitting"), with his head to the east. The skull, with a trepanation on the crown in the form of an oval hole, rested against the end wall of the pit, with the mandible touching the upper ribs. Legs were bent with knees up and at the same time spread apart in different directions (found *in situ* during the excavation). Arms were slightly bent at the elbows, somewhat apart. At the bottom of the chest laid a fragment of a large knife-like blade. The entire skeleton was painted with ochre[19].

The cultural affiliation of the Vynohradne Burial 3.15 is viewed as part of a continuation of Early Eneolithic steppe traditions of the Skeyla or Novodanylivka type [122,184,185].

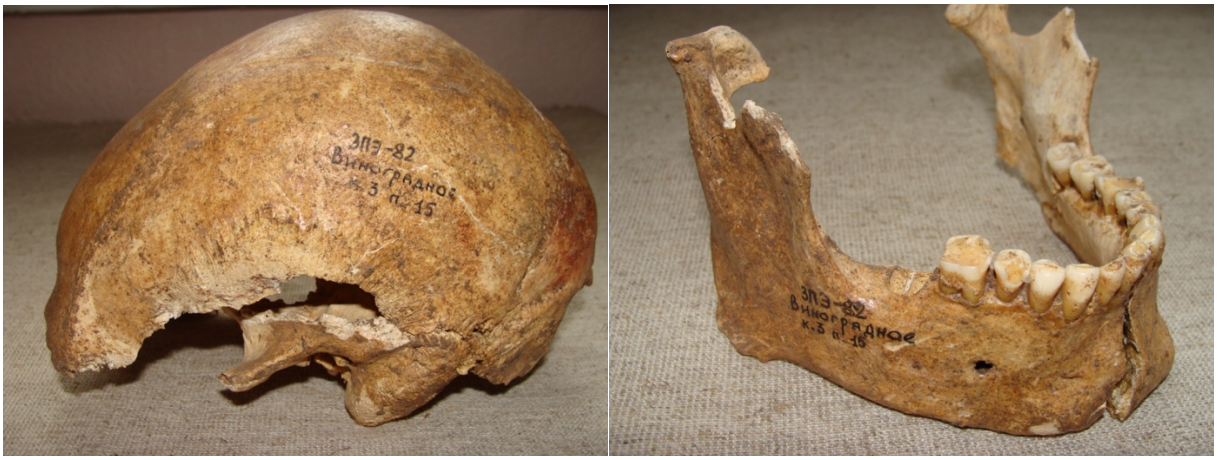

*Vynogradne Kurgan 3 Burial 15. Photo by I. D. Potekhina, 2012.*

*Yasynuvatka*

Vilnianskyi District, Zaporizhzhia Region (48.12, 35.09)

Yasynuvatka is one of the Mariupol-type cemeteries of extended burials in the Dnipro Valley. Yasynuvatka was discovered by A. B. Borodyansky and excavated by D. Ya. Telegin in 1978. The cemetery contained 68 burials. The earliest phase at the cemetery is represented by type A oval pits containing one to six skeletons. Multiple burials in the A-type pits were often on top of one another. Subsequent burials at the cemetery were placed in one large pit Б, 5.2 × 5.6m, the bottom of which was covered in red pigment.

V. N. Danilenko attributed Mariupol-type cemeteries to the Azov-Dnipro archaeological group. D. Y. Telegin considered these cemeteries to belong to several groups of the Dnipro-Donets archaeological complex (DDAC), extending from the Neolithic to the early Eneolithic. The latter included the late DDAC phase with Zasukha-type ceramics, synchronous with the early phase of Serednii Stih, represented by the Oleksandriya settlement and cemetery on the Oskil River [3].

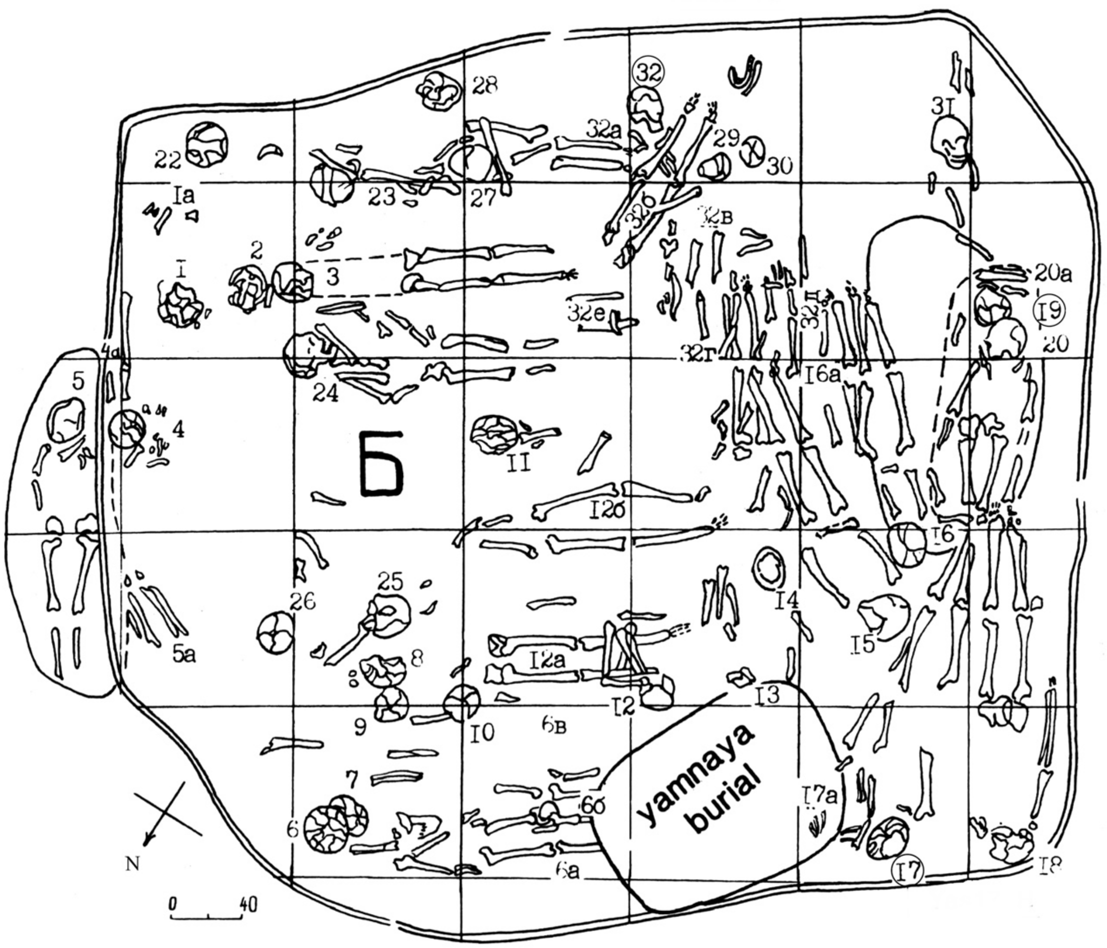

*Yasynuvatka cemetery, pit Б. Image from [3].*

## I27982

Burial 23.1, Pit Б-1, single cranium, male (40-45), 5700-5100 BCE

The Bioarchaeological Stores of the Institute of Archaeology of the National Academy of Sciences of Ukraine in Kyiv have individuals labeled Burial 23 Skeleton 1 and Burial 23 Skeleton 2. However, D. Y. Telegin's sketches of the burial pit only show a single cranium associated with Burial 23.

### I27983

Burial 24, male (30-40), 5700-5100 BCE

A male (30-40) burial in the red fill of Pit Б, extended supine, head to the NE (Б-1 type burial). Arms were slightly bent at elbows, wrists on the pelvis.

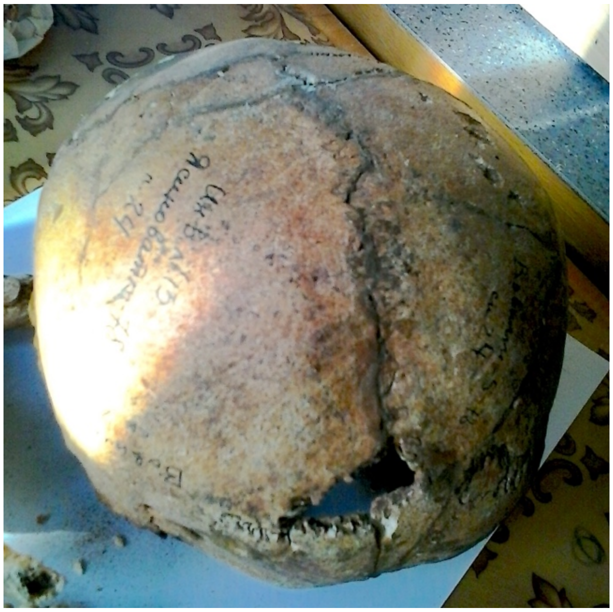

*Yasynuvatka Burial 24, occipital view of the cranium. Photo: A. G. Nikitin, 2014.*

### I27990

Burial 44, Pit Б-1 type burial (single cranium) in Pit Б, male, 5700-5100 BCE

### I27992

Burial 50, female, 5700-5100 BCE

An adult was buried in a boat-shaped grave under the north wall of the collective burial pit Б. The burial position is extended supine, head to NE, and face to the right. The tips of the feet and the cranium were above the 20-30 cm above the pelvis.

### I27994

Burial 55, Pit A2, male (25-30), 5630-5484 calBCE (6650±30 BP, Beta-445802)

Burials 55-58, 58a, and 59 were deposited in an oval pit A2, under Burial 31.

#### References

1. Makarenko, M. O. *The Mariupil Burial-Place*. (The Allukrainian Academy of Sciences, Kyiw, 1933).
2. Telegin, D. Ya. & Potekhina, I. D. *Neolithic Cemeteries and Populations in the Dnieper Basin*. (Archaeopress, Oxford, 1987).
3. Telegin, D. Y. *Neoliticheskiye Mogil'niki Mariupol'skogo Tipa*. (Naukova Dumka, Kiev, 1991).
4. Telegin, D. Ya., Potekhina, I. D., Lillie, M. & Kovaliukh, M. M. The chronology of the Mariupol-type cemeteries of Ukraine re-visited. *Antiquity* **76**, 356–363 (2002).
5. Mattila, T. M. *et al.* Genetic continuity, isolation, and gene flow in Stone Age Central and Eastern Europe. *Commun Biol* **6**, 793 (2023).

6. Nikitin, A. G., Sokhatsky, M. P., Kovaliukh, M. M. & Videiko, M. Y. Comprehensive Site Chronology and Ancient Mitochondrial DNA Analysis from Verteba Cave – a Trypillian Culture Site of Eneolithic Ukraine. *Interdisciplinaria Archaeologica. Natural Sciences in Archaeology.* **1**, 9–18 (2010).
7. Lillie, M. C. *et al.* First isotope analysis and new radiocarbon dating of Trypillia (Tripolye) farmers from Verteba Cave, Bilche Zolote, Ukraine. *Documenta Praehistorica* **44**, 306–324 (2018).
8. Mathieson, I. *et al.* The genomic history of southeastern Europe. *Nature* **555**, 197–203 (2018).
9. Nikitin, A. G. *et al.* Mitochondrial DNA analysis of Eneolithic Trypillians from Ukraine reveals Neolithic farming genetic roots. *PLoS One* **12**, e0172952 (2017).
10. Ward, G. K. & Wilson, S. R. Procedures for comparing and combining radiocarbon age determinations: a critique. *Archaeometry* **20**, 19–32 (1978).
11. Patokova, E. F., Petrenko, V. G., Burdo, N. B. & Polischuk, L. Y. *Pamyatniki Tripol'skoj Kul'tury v Severo-Zapadnom Prichernomorje*. (Naukova Dumka, Kiev, 1989).
12. Petrenko, V., Gerling, C. & Kaiser, E. Majaki – ein komplexes Denkmal der Usatovo-Kultur. Isotopendaten werfen neue Fragen auf. in *Eurasia Antiqua. Zeitschrift für Archäologie Eurasiens* (eds. Hansen, S. & Wagner, M.) vol. 21 45–74 (Rudolf Habelt Verlag, Bonn, 2018).
13. Ivanova, S. V., Petrenko, V. G. & Vetchinnikova, N. E. *Kurgans of Ancient Herdsmen from the South Bug and Dniester Interfluve*. (KP OGT, Odessa, 2005).
14. Telegin, D. Ya. *Dereivka. A Settlement and Cemetery of Copper Age Horse Keepers on the Middle Dnieper*. (British Archaeological Reports, Oxford, 1986).
15. Lillie, M., Budd, C., Potekhina, I. & Hedges, R. The radiocarbon reservoir effect: new evidence from the cemeteries of the middle and lower Dnieper basin, Ukraine. *J Archaeol Sci* **36**, 256–264 (2009).
16. Nikitin, A. G., Videiko, M., Patterson, N., Renson, V. & Reich, D. Interactions between Trypillian farmers and North Pontic forager-pastoralists in Eneolithic central Ukraine. *PLoS One* **18**, e0285449 (2023).
17. Wilde, S. *et al.* Direct evidence for positive selection of skin, hair, and eye pigmentation in Europeans during the last 5,000 y. *Proc Natl Acad Sci U S A* **111**, 4832–7 (2014).
18. Rassamakin, Y. Y. New radiocarbon dates to the absolute chronology of Eneolithic of the Black Sea steppe (preliminary analysis). *Arkheologicheskiy al'manakh* **20**, 289–296 (2009).
19. Rassamakin, Y. Y. Eneoliticheskiye pogrebeniya basseyna r. Molochnoy. in *Drevneyshiye skotovody stepey yuga Ukrainy* (ed. Shaposhnikova, O. G.) 31–43 (Naukova Dumka, Kiev, 1987).
20. Kotova, N. S. *Dereivskaya Kul'tura i Pamyatniki Nizhnemikhaylovskogo Tipa*. (Maidan, Kiev, Kharkov, 2013).
21. Serova, N. L. & Yarovoy, E. V. *Grigoriopolskie Kurgany*. (Shtiintsa, Kishinev, 1987).
22. Ivanova, S. V. New radiocarbon dates for the monuments in the North-Western coast of the Black Sea. *Arheologia* **3**, 69–75 (2010).

23. Yarovoy, E. V. Main Eneolithic burials of a mound near the village of Bursuceni in central Moldova (preliminary information). *Bulletin of the Moscow State Regional University (History and Political Science)* 54–68 (2019) doi:10.18384/2310-676X-2019-5-54-68.
24. Govedarica, B. & Manzura, I. The Giurgiuleşti cemetery in chronological and cultural context of southeastern and Eastern Europe. *Eurasia Antiqua* **22**, 1–39 (2016).
25. Todorova, H., Dimov, T. & Vaĭsov, I. Arkheologichesko prouchvane na praistoricheskiya nekropol pri s. Durankulak, Tolbukhinski okrŭg. in *XXVI Natsionalna konferentsiya po akhreologiya v Preslav* 17–18 (Sofia, 1981).
26. Kotova, N., Makhortykh, S. & Dzhos, V. New evidence on the interaction between the Yamnaya and Globular Amphora cultures. *Baltic-Pontic Studies* **25**, 317–341 (2021).
27. Nikitin, A. G., Ivanova, S., Kiosak, D., Badgerow, J. & Pashnick, J. Subdivisions of haplogroups U and C encompass mitochondrial DNA lineages of Eneolithic–Early Bronze Age Kurgan populations of western North Pontic steppe. *J Hum Genet* (2017) doi:10.1038/jhg.2017.12.
28. Ivanova, S. V. & Saveliev, O. K. Barrow near Sychavka Village of Odesa Oblast. *Archeologia* **3**, 70–82 (2011).
29. Agre, D. Arheologicheski razkopki na nadgrobna mogila v zemlishcheto na s. Popovo, obshtina Bolyarovo, Yambolska oblast. in *Arheologicheski otkritia i razkopki prez 2006 g.* 74–75 (Sofia, 2007).
30. Alexandrov, S. Fourth/third millennium BC barrow graves in North-East Bulgaria (120 years of investigations). in *Yamnaya Interactions. Proceedings of the International Workshop held in Helsinki, 25–26 April 2019* (eds. Heyd, V., Kulcsár, G. & Preda-Bălănică, B.) 271–314 (Budapest, 2021).
31. Tkachuk, T. M. The end of the stage CI and the beginning of the stage CII of the Tripolye culture in the Upper Dnestr region (according to material of the Bilshivtsi settlement). *Stratum Plus* **2**, 196–217 (2002).
32. Reimer, P. J. *et al.* The IntCal20 Northern Hemisphere Radiocarbon Age Calibration Curve (0–55 cal kBP). *Radiocarbon* **62**, 725–757 (2020).
33. Kadurin, S., Yanko-Hombach, V. & Smyntyna, O. The Ukraine: In Search of Submerged Late Palaeolithic Sites on the North-Western Black Sea Shelf. in 413–428 (2020). doi:10.1007/978-3-030-37367-2_21.
34. Larchenkov, E. & Kadurin, S. Paleogeography of the Pontic Lowland and northwestern Black Sea shelf for the past 25 k.y. in *Geology and Geoarchaeology of the Black Sea Region: Beyond the Flood Hypothesis* (Geological Society of America, 2011). doi:10.1130/2011.2473(06).
35. Svitoch, A. A. The Neoeuxinian basin of the Black Sea and the Khvalinian transgression of the Caspian Sea. *Quaternary International* **225**, 230–234 (2010).
36. Marin, I. N. & Palatov, D. M. Insights on the Existence of Ancient Glacial Refugee in the Northern Black/Azov Sea Lowland, with the Description of the First Stygobiotic Microcrustacean Species of the Genus Niphargus Schiödte, 1849 from the Mouth of the Don River. *Diversity (Basel)* **15**, 682 (2023).
37. Marret, F. *et al.* The Holocene history of the NE Black Sea and surrounding areas: An integrated record of marine and terrestrial palaeoenvironmental change. *Holocene* **29**, 648–661 (2019).

38. Dolukhanov, P. M. & Shilik, K. K. Environment, sea-level changes, and human migrations in the northern Pontic area during late Pleistocene and Holocene times. in *The Black Sea Flood Question: Changes in Coastline, Climate, and Human Settlement* 297–318 (Springer Netherlands, 2007). doi:10.1007/978-1-4020-5302-3_12.
39. Kiosak, D. Kamyane-Zavallia, the Easternmost Linear Pottery Culture Settlement Ever Excavated. *Sprawozdania Archeologiczne* **69**, 253–269 (2017).
40. Larina, O. V. Larina O.V. The Linear Pottery Culture of the area between rivers Prut and Dniester. *Stratum Plus* **2**, 10–140 (1999).
41. Tovkailo, M. The Neolithic period in Ukraine. in *Prehistoric Ukraine: From the First Hunters to the First Farmers* (eds. Lillie, M. C. & Potekhina, I. D.) 111–154 (Oxbow Books, 2020).
42. Rassamakin, Y. Y. Steppes of the Black Sea Region within a context of development of first agricultural societies . *Arkheolohiya* 3–26 (2004).
43. Petrenko, V. G. Problema «Tripol'ye i step'» i pamyatniki eneolita — ranney bronzy Severo-Zapadnogo Prichernomor'ya. *Materialy po arkheologii Severnogo Prichernomor'ya* **9**, 10–38 (2009).
44. Rassamakin, Y. Y. The Eneolithic of the Black Sea Steppe: Dynamics of Cultural and Economic Development 4500–2300 BC. in *Late Prehistoric Exploitation of the Eurasian Steppe* (eds. Levine, M., Rassamakin, Y., Kislenko, A. & Tatarintseva, N.) 59–182 (McDonald Institute Monographs, Cambridge, 1999).
45. Telegin, D. Y. *Seredn'o-Stohivs'ka Kul'tura Epokhy Midi*. (Naukova Dumka, Kyiv, 1973).
46. Nikitin, A. G. & Ivanova, S. Long-distance exchanges along the Black Sea coast in the Eneolithic and the steppe genetic ancestry problem. in *Steppe Transmissions* (eds. Preda-Bălănică, B. & Ahola, M.) 9–27 (Archaeolingua, Budapest, 2023). doi:10.33774/coe-2022-7m315.
47. Rassamakin, Y. Y. The Kvityanskaya Culture: History and Current State of the Problem. *Stratum Plus* **2**, 117–177 (2000).
48. Anthony, D. W. *The Horse, the Wheel, and Language: How Bronze-Age Riders from the Eurasian Steppes Shaped the Modern World*. (Princeton University Press, Princeton and Oxford, 2007).
49. Manzura, I. V. Eastern Europe at the dawn of the kurgan tradition. in *Ex Ungue Leonem. Sbornik statey k 90-letiyu L'va Samuilovicha Kleyna* (ed. Vishnyatsky, L. B.) 107–129 (Nestor-Istoriya, St. Petersburg, 2017).
50. Manzura, I. History Carved by the Dagger: the Society of the Usatovo Culture in the 4th Millennium BC. in *Repräsentationen Der Macht. Beiträge des Festkolloquiums zu Ehren des 65. Geburtstags von Blagoje Govedarica* (ed. Hansen, S.) 73–96 (Deutsches Archäologisches Insitut, Harrassovitz Verlag, Wiesbaden, 2020).
51. Lazaridis, I., Patterson, N., Anthony, D. & & others. *The Genetic Origin of the Indo-Europeans. in Submission*. (2024).
52. Rassamakin, Y. Y. Stepy Pivnichnoho Prychornomor"ya za doby midi. in *Ukrayina: khronolohiya rozvytku. Z davnikh chasiv do pizn'oyi antychnosti* vol. 1 202–2018 (Kyiv, 2008).

53. Mileto, S., Kaiser, E., Rassamakin, Y., Whelton, H. & Evershed, R. P. Differing modes of animal exploitation in North-Pontic Eneolithic and Bronze Age Societies. *STAR: Science & Technology of Archaeological Research* **3**, 112–125 (2018).
54. Rassamakin, Y. Y. & Evdokimov, G. L. Novyj pozhdneeneoliticheskij mogil’nik na iyge Khersonscheny v svete regional’nykh issledovanij stepnogo eneolita. *Donets’kyj Archeologichnyj Zbirnyk* **13/14**, 7–29.
55. Trifonov, V. A. Western boundaries of extension of the Maikop culture. *Izvestiya Samarskogo nauchnogo tsentra Rossiyskoy akademii nauk* 276–284 (2014).
56. Rassamakin, J. Ja. *Die nordpontische Steppe in der Kupferzeit : Gräber aus der Mitte des 5. Jts. bis Ende des 4. Jts. v. Chr. = Azovo-Pontijskie stepi v ėpochu medi : pogrebal’nye pamjatniki serediny V - konca IV tys. do n. e.* (P. von Zabern, Mainz, 2004).
57. Rassamakin, Y. Y. Between the Don and the Danube: The Phenomenon of the Zhivotilovka-Volchanskoe Type of Burials at the Turn of the Late Eneolithic and the Early Bronze Age in the Northern Black Sea Steppe. in *The Caucasus. Bridge between the urban centres in Mesopotamia and the Pontic steppes in the 4th and 3rd millennium BC. The transfer of knowledge and technologies between East and West in the Bronze Age* (eds. Giemsch, L. & Hansen, S.) 195–210 (Verlag Schnell & Steiner GmbH, Regensburg, 2021).
58. Rassamakin, Y. Y. Podveski bolgradskogo tipa kak spetsificheskiy indikator migratsionnykh protsessov na rubezhe eneolita-rannego bronzovogo veka v pontiyskikh stepyakh. in *XV Ural’skoye arkheologicheskoye soveshchaniye. Tezisy dokladov konferentsii* (ed. Morgunova, N. L.) 104–106 (Orenburg, 2001).
59. Kovaleva, I. F. Pogrebeniya zhivotilovskogo tipa v Prisamar’ye. in *Kurgannyye drevnosti stepnogo Podneprov’ya (III–I tys. do n.e.)* (ed. Kovaleva, I. F.) 46–54 (Dnepropetrovsk, 1978).
60. Demchenko, T. On the issue of the Bursuchenskaia cultural group identification within Gordinești - Maykopsk village phe- nomenon. in *Culturi, procese şi contexte în arheologie* (eds. Sîrbu, L., Telnov, N., Ciobanu, L., Sîrbu, G. & Kașuba, M.) 84–99 (Tipogr. ‘Garamont-Studio’’’, Chişinău, 2016).
61. Dergachev, V. A. Maykop in the late Tripolye culture. in *Antiquities of East Europe, South Asia and South Siberia in the context of connections and interactions within the Eurasian cultural space (new data and concepts). Volume II. Connections, contacts and interactions between ancient cultures of northern Eura* (eds. Polyakov, A. V. & Tkach, E. S.) 128–131 (IIMK RAN, Nevskaya Tipographia, St. Petersburg, 2019).
62. Immel, A. *et al.* Gene-flow from steppe individuals into Cucuteni-Trypillia associated populations indicates long-standing contacts and gradual admixture. *Sci Rep* **10**, 4253 (2020).
63. Penske, S. *et al.* Early contact between late farming and pastoralist societies in southeastern Europe. *Nature* **620**, 358–365 (2023).
64. Dergachev, V. A. *Late Tripolye and Maykop.* (Wydawnictwo Uniwersytetu Rzeszowskiego , Rzeszów, 2021).
65. Petrenko, V. G. Usatovskaya kul’tura. in *Drevniye kul’tury Severo-Zapadnogo Prichernomor’ya* (eds. Bruyako, I. V. & Samoylova, T. L.) 163–210 (SMIL, Odessa, 2013).
66. Diachenko, A. & Harper, T. K. The absolute chronology of Late Tripolye sites: a regional approach. *Sprawozdania Archeologiczne* **68**, 81–105 (2016).

67. Potekhina, I. D. Formuvannya eneolitychnoho naselennya Pivnichno-Zakhidnoho Prychornomor"ya za antropolohichnymy ta arkheohenetychnymy danymy mohyl'nykiv usativs'koyi kul'tury. in *Anotatsiyi dopovidey Mizhnarodnoyi konferentsiyi «Vid paleolitu do kozats'koyi Ukrayiny», prysvyachenoyi 100-litn'omu yuvileyu D.YA. Telehina* (eds. Zaliznyak, L. L. & Potekhina, I. D.) 91 (Oleh Filyuk, Kyiv, 2019).
68. Potekhina, I. D. Antropologicheskie materialy iz mogil'nika Mayaki. in *Pamyatniki Tripol'skoj kul'tury v Severo-Zapadnom Prichernomorje* (eds. Patokova, E. F., Petrenko, V. G., Burdo, N. B. & Polischuk, L. Y.) 125–133 (Naukova Dumka, Kiev, 1989).
69. Patokova, E. F. *Usatovskoye Poseleniye i Mogil'niki*. (Naukova Dumka, Kiev, 1979).
70. Manzura, I. The Mayaki cemetery as a source for social organization of the Usatovo culture. in *Culturi, Procese și Contexte în Arheologie* (eds. Sîrbu, L., Telnov, N., Ciobanu, L., Sîrbu, G. & Kașuba, M.) 63–83 (Chişinău, 2016).
71. Petrenko, V. G. & Kaiser, E. Kompleksnyy pamyatnik Mayaki: novyye izotopnyye daty i voprosy khronologii nalichnykh kul'tur. *Materialy po arkheologii Severnogo Prichernomor'ya* **12**, 31–61 (2011).
72. Nikitin, A. G. & Ivanova, S. The Origins and Chronology of the Usatove Culture. *Archaeologia Lituana* **23**, 148–156 (2022).
73. Dergachev, V. A. Kulturelle und historische Entwicklungen im Raum zwischen Karpaten und Dnepr. Zu den Beziehungen zwischen frühen Gesellschaften im nördlichen Südost- und Osteuropa. in *Das Karpatenbecken und die osteuropäische Steppe. Nomadenbewegungen und Kulturaustausch in den vorchristlichen Metallzeiten (4000–500 v. Chr.). Prähistorische Archäologie in Südosteuropa 12* (eds. Hänsel, B. & Machnik, J.) 27–64 (Marie Leidorf, Rahden/Westf., 1998).
74. Razumov, S. Transversal arrowheads of the Usatovo culture. in *Connections, contacts and interactions between ancient cultures of Northern Eurasia and civilizations of the East during the Palaeometal period (IV–I mil. BC)* 141–143 (Institute for the History of Material Culture Russian Academy of Sciences, 2019). doi:10.31600/978-5-907053-35-9-141-143.
75. Nikitin, A. G., Ivanova, S., Culleton, B. J., Potekhina, I. & Reich, D. New radiocarbon and stable isotope data from the Usatove culture site of Mayaky in Ukraine. *SSRN Electronic Journal* (2023) doi:10.2139/ssrn.4236123.
76. Cook, G. T. *et al.* Problems of dating human bones from the Iron Gates. *Antiquity* **76**, 77–85 (2002).
77. Weber, A. W. *et al.* Chronology of middle Holocene hunter–gatherers in the Cis-Baikal region of Siberia: Corrections based on examination of the freshwater reservoir effect. *Quaternary International* **419**, 74–98 (2016).
78. Cook, G. T. *et al.* A Freshwater Diet-Derived $^{14}$C Reservoir Effect at the Stone Age Sites in the Iron Gates Gorge. *Radiocarbon* **43**, 453–460 (2001).
79. Ramsey, C. B., Schulting, R., Goriunova, O. I., Bazaliiskii, V. I. & Weber, A. W. Analyzing Radiocarbon Reservoir Offsets Through Stable Nitrogen Isotopes and Bayesian Modeling: A Case Study Using Paired Human and Faunal Remains from the Cis-Baikal Region, Siberia. *Radiocarbon* **56**, 789–799 (2014).

80. Bonsall, C. *et al.* Radiocarbon and Stable Isotope Evidence of Dietary Change from the Mesolithic to the Middle Ages in the Iron Gates: New Results from Lepenski Vir. *Radiocarbon* **46**, 293–300 (2004).
81. Bonsall, C., Macklin, M., Payton, R. & Boroneanţ, A. Climate, floods and river gods. *Before Farming* **2002**, 1–15 (2002).
82. Bonsall, C. *et al.* Stable isotopes, radiocarbon and the Mesolithic-Neolithic transition in the Iron Gates. *Documenta Praehistorica* **27**, 19–132 (2000).
83. Cook, G. T. *et al.* The Mesolithic–Neolithic Transition in the Iron Gates, Southeast Europe: calibration and dietary issues. in *Chronology and Evolution within the Mesolithic of North-West Europe* (eds. Crombé, P., Van Strydonck, M., Sergant, J., Boudin, M. & Bats, M.) 497–515 (Cambridge Scholars Publishing, 2009).
84. Borić, D. & Price, T. D. Strontium isotopes document greater human mobility at the start of the Balkan Neolithic. *Proceedings of the National Academy of Sciences* **110**, 3298–3303 (2013).
85. Borić, D., French, C. & Dimitrijević, V. Vlasac revisited: formation processes, stratigraphy and dating. *Documenta Praehistorica* 261–287 (2008).
86. Chintalapati, M., Patterson, N. & Moorjani, P. The spatiotemporal patterns of major human admixture events during the European Holocene. *Elife* **11**, (2022).
87. Munchayev, R. M. *Kavkaz Na Zare Bronzovogo Veka: Neolit, Eneolit, Rannyaya Bronza*. (Nauka, Moscow, 1975).
88. Rigaud, S. *et al.* Exploring Hypotheses on Early Holocene Caspian Seafaring Through Personal Ornaments: A Study of Changing Styles and Symbols in Western Central Asia. *Open Archaeology* **9**, (2023).
89. Telegin, D. Y. Yeshche raz o vydelenii pamyatnikov novodanilovskogo tipa epokhi medi . in *Drevneyshiye obshchnosti zemledel'tsev i skotovodov Severnogo Prichernomor'ya (V tys. do n.e. – V v. n.e.). materialy mezhdunarodnoy konferentsii Kishinev 10-14 dekabrya 1990 g.* (ed. Yarovoj, E. V.) 60–61 (Kiev, 1991).
90. Videiko, M. Y. 'Shumers'kyy slid' v Trypil's'kiy arkhitekturi? *Trypillian Civilization journal* (2010).
91. Palaguta, I. An Assemblage of Anthropomorphic Figurines of the Neolithic and Copper Age Balkan-Carpathians Cultures: some Observations on the Structure of Images and its Development. in *Cucuteni Culture within the European Neo-Eneolithic Context: Proceedings of the International Colloquium „Cucuteni - 130. 15-17 October 2014, Piatra-Neamţ, Romania: In Memoriam dr. Dan Monah, In Memoriam dr. Gheorghe Dumitroaia* (eds. Preoteasa, C. & Nicola, C.-D.) 327–348 (Editura "Constantin Matasă", Piatra-Neamţ, 2016).
92. Rezepkin, A. Maykop Culture: Issue of Chronology and Cultural Relationships of the Ust-Dzheguta and Psekupsk Variants. *Slovenská archeológia* **68**, 515–526 (2020).
93. The British Museum Online Collection. https://www.britishmuseum.org/collection/search?object=pin&place=Middle%20East&view=grid&sort=object_name__asc&page=2#page-top.
94. *Treasures from the Royal Tombs of Ur*. (University of Pennsylvania, Philadelphia, 1998).
95. Dergachev, V. A. New hoards of metal objects of The Cucuteni-Trypillian Culture found in The Middle Dniester region (Ukraine). in *Culturi, procese şi contexte în arheologie* (eds.

Sîrbu, L., Telnov, N., Ciobanu, L., Sîrbu, G. & Kașuba, M.) 36–51 (Tipogr. 'Garamont-Studio', Chişinău, 2016).
96. Chernykh, L. A. & Daragan, M. N. *Kurgany Epokhi Eneolita-Bronzy Mezhdurech'ya Bazavluka, Solenoy, Chertomlyka : Monografiya. Kurgany Ukrainy.* vol. 4 (Oleg Filyuk, Kiev, 2014).
97. Petrenko, V. G., Sapozhnikov, I. V. & Sapozhnikova, G. V. Geometricheskiye mikrolity usatovskoy kul'tury. in *Drevneye Prichernomor'ye. Kratkiye soobshcheniya Odesskogo arkheologicheskogo obshchestva* 42–47 (Odessa, 1994).
98. Manzura, I., Nosova, L. & Petrenko, V. G. Zwei unbekannte Gräber der Usatovo-Kultur auf dem Gräberfeld von Majaki (Ukraine). *Altertum* **68**, 1–14 (2023).
99. Sinica, V. S., Razumov, S. N. & Telnov, N. P. *Archaeological Heritage of Pridnestrovie*. (GUIPP «Bend. tipografiya «Poligrafist», Tiraspol, 2016).
100. *Early Urbanism in Europe. The Trypillia Megasites of the Ukrainian Forest-Steppe*. (De Gruyter Poland Ltd, Warsaw/Berlin, 2020). doi:10.1515/9783110664959.
101. Ivanova, S. & Nikitin, A. G. The Yamna culture: Origin and migrations in the context of the Frontier Theory. *Notes of the History Faculty, Odessa I. I. Mechnikov National University* **31**, (2021).
102. Faifert, A. V. Early stage of the Yamanya culture from the Lower Don Region. (Voronezh State University, 2017).
103. Diaconescu, D. Step by Steppe: Yamnaya culture in Transylvania. *Praehistorische Zeitschrift* **95**, 17–47 (2020).
104. Palalidis, S. A GIS-based approach to the study of the Yamnaya cultural horizon (ca. 3300-2400 BC): preliminary results. in *InFieri. Incontri di Archeologia Sapienza Miscellanea degli atti II (2018-2019) e III (2020)* (eds. Abbondanzieri, E. & et al.) (Edizioni Quasar di Severino Tognon s.r.l., Roma, 2023).
105. Topal, D. Patterns of kurgan landscape between Dniester, Prut and Danube rivers. in *Connections, contacts and interactions between ancient cultures of Northern Eurasia and civilizations of the East during the Palaeometal period (IV–I mil. BC)* 29–32 (Institute for the History of Material Culture Russian Academy of Sciences, 2019). doi:10.31600/978-5-907053-35-9-29-32.
106. Dvorianinov, S. A., Petrenko, V. G. & Rychkov, N. A. K izucheniyu orientirovki yamnykh pogrebeniy. in *Drevnosti Severo-Zapadnogo Prichernomoria* (ed. Stanko, V. N.) 22–38 (Naukova Dumka, Kiev, 1981).
107. Subbotin, L. V., Razumov, S. N. & Sinika, V. S. *Semyonovka Barrows*. (Stratum Plus, Tiraspol, 2017).
108. Yarovoy, E. V. *Drevneyshie Skotovodcheskie Plemena Yugo-Zapada SSSR (Klassifikatsiya Pogrebalnogo Obriada)*. (Shtiintsa, Kishinev, 1985).
109. Ivanova, S. V. *Sotsial'naya Struktura Naseleniya Yamnoy Kul'tury Severo-Zapadnogo Prichernomor'ya*. (Druk, Odessa, 2001).
110. Dergachev, V. A. *Moldaviya i Sosednie Territorii v Epokhu Bronzy*. (Shtiintsa, Kishinev, 1986).
111. Nikolova. A. V. Khronologicheskaya klassifikatsiya pamyatnikov yamnoy kul'tury stepnoy zony Ukrainy. (Institute of Archaeology, National Academy of Sciences of Ukraine, Kyiv, 1992).

112. Ivanova, S. *Istoriya Naselennya Pivnichno-Zakhidnoho Prychornomor"ya Naprykintsi IV–III Tys. Do n. e.* . (Buk-Druk, Zhytomyr, 2021).
113. Ivanova, S. V. The social organisation of the Yamnaya culture tribes in the northwest Pontic area. *Stratum Plus* **2**, 388–402 (2000).
114. Włodarczak, P. Kultura złocka i problem genezy kultury ceramiki sznurowej w Małopolsce. in *Na pograniczu światów. Studia z pradziejów międzymorza bałtycko-pontyjskiego ofiarowane Profesorowi Aleksandrowi Kośko w 60 rocznicę urodzin* (eds. Bednarczyk, J., Czebreszuk, J., Makarowicz, P. & Szmyt, M.) 511–532 (Wydawnictwo Poznańskie, Poznań, 2008).
115. Rassamakin, Y. Y. & Nikolova, A. V. Carpathian Imports and Imitations in Context of Eneolithic and Bronze Age of the Black See Area. in *Import and Imitation in Archeology* (eds. Biehl, P. & Rassamakin, Y. Y.) 51–88 (Beier & Beran, Langenweibach, 2008).
116. Gening, V. V. & Korpusova, V. N. *Arkheologicheskiye Pamyatniki Krymskogo Prisivash'ya. Kurgany u s. Istochnoye i s. Bolotnoye*. (Institut Zoologii, Kiyev, 1989).
117. Bratchenko, S. N. *Donets'ka Katakombna Kul'tura Rann'oho Etapu*. (Shlyakh, Luhans'k, 2001).
118. Otroshchenko, V. V. Doba bronzy na terenakh Ukrayiny . in *Ukrayina. Khronolohiya rozvytku* (eds. Otroshchenko, V. V., Rassamakin, Y. Y. & Chernykh, L. A.) vol. 1 219–331 (KVSHTS, Kyiv).
119. Kaiser, E. Datarea absolută a culturii înmormântărilor în catacombe din nordul Mării Negre. **3(18)**, 59–70 (2009).
120. Sanzharov, S. N. *Poseleniya Neolita – Ranney Bronzy Severskogo Dontsa*. (SNU, Lugansk, 2000).
121. Lytvynenko, R. O. The Babyne culture circle. (Institute of Archaeology, National Academy of Sciences of Ukraine, Kyiv, 2009).
122. Pustovalov, S. Kurhanni svyatylyshcha doby rann'oyi bronzy stepovoyi chastyny Ukrayiny ta yikh astronomichna skladova. in *Ukrayins'ke nebo. Studiyi nad istoriyeyu astronomiyi v Ukrayini* (ed. Petruk, O.) 39–50 (Instytut prykladnykh problem mekhaniky i matematyky im. YA. S. Pidstryhacha NAN Ukrayiny, Lviv, 2014).
123. Drovosekova, O. V. Materialy epokhi eneolita i bronzovogo veka iz s. kapulovka Dnepropetrovskoy oblasti. in *Starozhytnosti stepovoho Prychornomor"ya i Krymu* (ed. Tolochko, P. P.) vol. 10 131–154 (Zaporizhzhia, 2002).
124. Teslenko, D. L., Kaiser, E. & Shalobudov, V. M. Kurgan sites in the Samara River Basin: Results of interdisciplinary research . *Archaeology and Early History of Ukraine* **46**, 146–177 (2023).
125. Ward, R. H., Frazier, B. L., Dew-Jager, K. & Pääbo, S. Extensive mitochondrial diversity within a single Amerindian tribe. *Proceedings of the National Academy of Sciences* **88**, 8720–8724 (1991).
126. Günther, T. *et al.* Population genomics of Mesolithic Scandinavia: Investigating early postglacial migration routes and high-latitude adaptation. *PLoS Biol* **16**, e2003703 (2018).
127. Mathieson, I. *et al.* Genome-wide patterns of selection in 230 ancient Eurasians. *Nature* **528**, 499–503 (2015).
128. Krause, J. *et al.* A Complete mtDNA Genome of an Early Modern Human from Kostenki, Russia. *Current Biology* **20**, 231–236 (2010).

129. Fu, Q. *et al.* The genetic history of Ice Age Europe. *Nature* **534**, 200–205 (2016).
130. Haak, W. *et al.* Massive migration from the steppe was a source for Indo-European languages in Europe. *Nature* **522**, 207–11 (2015).
131. Jones, E. R. *et al.* The Neolithic Transition in the Baltic Was Not Driven by Admixture with Early European Farmers. *Current Biology* **27**, 576–582 (2017).
132. Juras, A. *et al.* Maternal genetic origin of the late and final Neolithic human populations from present-day Poland. *Am J Phys Anthropol* **176**, 223–236 (2021).
133. Allentoft, M. E. *et al.* Population genomics of post-glacial western Eurasia. *Nature* **625**, 301–311 (2024).
134. Krzewińska, M. *et al.* Ancient genomes suggest the eastern Pontic-Caspian steppe as the source of western Iron Age nomads. *Sci Adv* **4**, (2018).
135. Włodarczak, P. Kurgan rites in the Eneolithic and Early Bronze Age Podolia in light of materials from the funerary ceremonial centre at Yampil. *Baltic-Pontic Studies* **22**, 246–283 (2018).
136. Knipper, C. *et al.* Female exogamy and gene pool diversification at the transition from the Final Neolithic to the Early Bronze Age in central Europe. *Proceedings of the National Academy of Sciences* **114**, 10083–10088 (2017).
137. Battaglia, V. *et al.* Y-chromosomal evidence of the cultural diffusion of agriculture in southeast Europe. *European Journal of Human Genetics* **17**, 820–830 (2009).
138. Lipson, M. *et al.* Parallel palaeogenomic transects reveal complex genetic history of early European farmers. *Nature* **551**, 368–372 (2017).
139. Dulias, K. *et al.* Ancient DNA at the edge of the world: Continental immigration and the persistence of Neolithic male lineages in Bronze Age Orkney. *Proceedings of the National Academy of Sciences* **119**, (2022).
140. Nikitin, A. G. *et al.* Interactions between earliest Linearbandkeramik farmers and central European hunter gatherers at the dawn of European Neolithization. *Sci Rep* **9**, 19544 (2019).
141. Paschou, P. *et al.* Maritime route of colonization of Europe. *Proceedings of the National Academy of Sciences* **111**, 9211–9216 (2014).
142. Fernández, E. *et al.* Ancient DNA Analysis of 8000 B.C. Near Eastern Farmers Supports an Early Neolithic Pioneer Maritime Colonization of Mainland Europe through Cyprus and the Aegean Islands. *PLoS Genet* **10**, e1004401 (2014).
143. Gaskevych, D. L. North-Pontic Impresso: Origin of the Neolithic Pottery with Comb Decoration in the South of Eastern Europe. *Stratum Plus* **2**, 213–251 (2010).
144. Burdo, N. B. Late Neolithic cultural elements from the Danube and Carpathian regions of Precucuteni – Trypillia A culture. *Documenta Praehistorica* **XXXVIII**, 357–371 (2011).
145. Underhill, P. A. *et al.* The phylogenetic and geographic structure of Y-chromosome haplogroup R1a. *European Journal of Human Genetics* **23**, 124–131 (2015).
146. Wang, C.-C. *et al.* Ancient human genome-wide data from a 3000-year interval in the Caucasus corresponds with eco-geographic regions. *Nat Commun* **10**, 590 (2019).
147. Lazaridis, I. *et al.* The genetic history of the Southern Arc: A bridge between West Asia and Europe. *Science (1979)* **377**, (2022).

148. González-Fortes, G. *et al.* Paleogenomic Evidence for Multi-generational Mixing between Neolithic Farmers and Mesolithic Hunter-Gatherers in the Lower Danube Basin. *Current Biology* **27**, 1801-1810.e10 (2017).
149. Gorbov, V. & Kolesnik, A. New Neolithic Settlement in Mariupol and Its Place in the System of Synchronous Monuments. *Science Journal of VolSU. History. Area Studies. International Relations* **21**, 16–31 (2016).
150. Mykhailova, N. Mykola Makarenko and Mariupol Neolithic Burial Ground The tragic fates of the archaeologist and the site. *Archaeologia Lituana* **24**, 19–33 (2024).
151. De Muynck, D., Huelga-Suarez, G., Van Heghe, L., Degryse, P. & Vanhaecke, F. Systematic evaluation of a strontium-specific extraction chromatographic resin for obtaining a purified Sr fraction with quantitative recovery from complex and Ca-rich matrices. *J Anal At Spectrom* **24**, 1498 (2009).
152. Ventresca Miller, A. R. *et al.* Re-evaluating Scythian lifeways: Isotopic analysis of diet and mobility in Iron Age Ukraine. *PLoS One* **16**, e0245996 (2021).
153. Gerling, C. *Prehistoric Mobility and Diet in the West Eurasian Steppes 3500 to 300 BC*. (De Gruyter, 2015). doi:10.1515/9783110311211.
154. Vajsov, I. Das Grab 982 und die Protobronzezeit in Bulgarien. in *Durankulak, Band II. Die prähistorischen Gräberfelder* (ed. Todorova, H.) 159–176 (Publishing House Anubis Ltd., Sofia, 2002).
155. Alexandrov, S. Forth-Second Millennium BC Barrow Graves in North Bulgaria. in *The River and the Time* (ed. Atanasov, V.) 27–39 (Tutrakan, 2010).
156. Suvorov, A. Modelling the Yamnaya Expansion Through Radiocarbon Dates. (University of Helsinki, Helsinki, 2021).
157. Agulnikov, S. & Popovich, S. *Raskopki Kurganov v Rayone Shtefan-Vode, Respublika Moldova: (1989, 1990, 1991)*. (Agenția Națională Arheologică, Chisinau, 2022).
158. Agulnikov, S. M. Issledovanie kurganov v s. Kotiuzhen’. *Archeologicheskie Issledovaniya v Moldove v 1986 g.* 104–120 (1992).
159. Popovici, S. & Ceban, I. Results of the Archaeological Investigations on the Tripolye C2 Site at Cunicea Prișanscaia-Gora. *Arheologia Moldovei* **37**, 205–217 (2014).
160. Dergachev, V. A. Raskopki v Danchenakh i nekotoryye voprosy izucheniya pamyatnikov pozdnego Tripol’ya i Katakombnoy kul’tury. in *Arkheologicheskiye Issledovaniya v Moldavii v 1974-1976 gg.* 35–45 (Кишинев, 1981).
161. Derhachev, V. A. & Manzura, I. V. *Pogrebal’nyye Kompleksy Pozdnego Tripol’ya*. (Štiinca, Chișinău, 1991).
162. Haheu, V. & Kurciatov, S. Cimitirulplan Eneolitic de lingă satul Giurgiuleşti. *Revista Arkheologică* 101–114 (1993).
163. Lysenko, S. D., Razumov, S. N., Lysenko, S. S., Sinika, V. S. & Тельнов, Н. П. Burials from the Late Bronze Age from the barrows of the “DOT”, “SAD”, “VODOVOD” groups near the Glinoye village on the Left Bank of the Lower Dniester . *Proceedings in Archaeology and History of Ancient and Medieval Black Sea Region* 164–201 (2022) doi:10.53737/2713-2021.2021.54.48.004.
164. Beylekchi, V. S. Raskopki kurgana 3 v s. Mierkulesht’. *Archeologicheskie Issledovaniya v Moldove v 1986 g.* 72–87 (1992).

165. Dergachev, V. A. & Sava, E. N. Excavation of tumuli in the vicinity of the villages of Mereni and Chirca. *Stratum Plus* **2**, 526–562 (2002).
166. Manzura, I. V., Klochko, E. O. & Savva, E. N. *Kamenskie Kurgany*. (Shtiintsa, Kishinev, 1992).
167. Leviţki, O., Manzura, I. & Demcenko, T. Necropola tumulară de la Sărăteni. in *Bibliotheca Thracologica XVII* (Bucureşti, 1996).
168. Sava, E., Agulnikov, S. & Manzura, I. *Issledovaniya Kurganov v Budzhakskoy Stepi (1980-1985 Gg.)*. (Bons Offices, Chişinău, 2019).
169. Savva, E. N. Issledovanie kurgana u g. Tiraspolya. *Archeologicheskie Issledovaniya v Moldavii v 1983 g.* 44–59 (1983).
170. Makhortykh, S. V., Kotova, N. S., Dzhos, V. S. & Radchenko, S. B. New burial and ritual assemblages of Early Bronze Age located near the complex of Kamyana Mohyla. *Archaeology and Early History of Ukraine* **37**, 226–239 (2020).
171. Zbenovich, V. G. Issledovanie pozdnetripol'skogo poselenija usatovskogo tipa b s. Mayaki. in *Arkheologicheskie Issledovanija na Ukraine v 1965-1966 gg.* (ed. Tolochko, P. P.) 83–87 (Naukova Dumka, Kiev, 1967).
172. Zbenovich, V. G. *Pozdnetripol'skiye Plemena Severnogo Prichernomor'ya*. (Naukova Dumka, Kiev, 1974).
173. Zinkovsky, K. V. & Patokova, E. F. Issledovaniya Mayakskogo mogil'nika v 1974 g. . in *Arkheologicheskiye issledovaniya Severo-Zapadnogo Prichernomor'ya* 134–143 (Naukova Dumka, Kiev, 1978).
174. Zin'kovskiy, K. V. & Petrenko, V. G. Burials with traces of ochre in Usatovo burial grounds. *Sovetskaya Arkheologia* **4**, 24–39 (1987).
175. Neradenko, T. M. The settlement Molyukhiv Buhor in the works of D. Ya. Telegin and according to modern data. *Archaeology and Early History of Ukraine* **37**, 348–357 (2020).
176. Korobkova, G. F. & Shaposhnikova, O. G. *Poselenie Mikhailovka: Etalonnyj Pamyatnik Drevneyamnoj Kultury*. (Evropejskij Dom, St. Petersburg, 2005).
177. Kotova, N. S. & Videiko, M. Y. The Absolute Chronology of the Ukraine during the Eneolithic. in *Zwischen Karpaten und Ägäis. Neolithikum und ältere Bronzezeit.* (eds. Hansel, B. & Studenikova. E.) 121–134 (Verlag Marie Leidorf GmbH · ., Rahden/Westf, 2004).
178. Toshchev, G. N. Yeshche raz o stratigrafii Odesskogo kurgana. in *Kurgany v zonakh novostroyek Moldavii* 175–182 (Shtiintsa , Kishinev, 1984).
179. Biagi, P. & Kiosak, D. The Mesolithic of the northwestern Pontic region: New AMS dates for the origin and spread of the blade and trapeze industries in southeastern Europe. *Eurasia Antiqua* 21–41 (2010).
180. Rassamakin, Y. Y. Mohyl'nyky Ihren' (Ohrin') 8 ta Oleksandriya doby eneolitu: problemy datuvannya ta kul'turnoyi prynalezhnosti. *Archeologia* 26–48 (2017).
181. Nikitin, A. G. & Ivanova, S. The megalithic past of the Bronze Age kurgans of the North Pontic Region. in *19th Annual Meeting of the European Association of Archaeologists* 178–179 (Plzen, Czech Republic, 2013).
182. Ivanova, S. V. New radiocarbon dates for the monuments in the north-western coast of the Black Sea. *Archeologia* **3**, 69–75 (2010).

183. Nikitin, A. G. Bioarchaeological Analysis of Bronze Age Human Remains from the Podillya Region of Ukraine. *Interdisciplinaria Archaeologica. Natural Sciences in Archaeology.* **2**, 9–14 (2011).
184. Pustovalov, S. Zh. Molochans'ke svyatylyshche. *Problemy Arkheolohiyi Podniprov"ya* 104–118 (1999).
185. Rassamakin, Y. Y. Das Frühäneolithikum der Don-Dnepr-Region (zweites bis drittes Viertel des 5 Jts. V. Chr.): einige Proleme der Interpretaon von archäologischen Denkmälern . in *Repräsentationen der Macht. Beiträge des Festkolloquiums zu Ehren des 65. Geburtstages von Blagoje Govedarica* (ed. Hansen, S.) 27–43 (Harrassovitz Verlag, Wiesbaden, 2020).
186. Dergachev, V. A. & Manzura, I. V. *Pogrebal'nyye Kompleksy Pozdnego Tripol'ya*. (Shtiintsa, Kishinev, 1991).

### A genomic history of the North Pontic Region from the Neolithic to the Bronze Age

*Supplementary File S2*

#### Population genetic analysis of populations from the North Pontic and adjacent regions

**Overview**

The North Pontic region (NPR) forms the western end of the Eurasian steppe and connects that great expanse of flatlands to its east with the world of central and southeastern Europe to its west. We study Eneolithic and Bronze Age populations that lived north and west of the Black Sea to arrive at key insights about their population history. First, we identify single individuals at Zhivotilovka, Taraclia, and Csongrád that crossed the NPR from the east without any discernible admixture with native populations, suggesting that long-distance migration occurred even before it became common during the Yamna expansion of the Bronze Age. In the main, however, Pre-Yamna period populations of the NPR were formed by admixture of varied Caucasus-Lower Volga (CLV) people from the east with native populations of the NPR. The Trypillian farmers received a little ancestry from these CLV migrants, but also contributed about half of the ancestry of Usatove-related populations that were formed on the basis of even Trypillian-CLV admixture. CLV migrants also admixed with NPR hunter-gatherers (forming the Serednii Stih populations, and eventually the Yamna themselves). These NPR hunter-gatherers were not isolated from the farmers of Europe but also received substantial input from them. The unprecedented mobility of the Yamna expansion included many individuals without any admixture of the pre-Yamna populations, but also many of diverse additional ancestry: from European farmers (west of the NPR), Ukraine hunter-gatherers (from the NPR itself), but also Maykop and Steppe Maykop admixture (from east of the NPR). The Yamna were thus formed from proximal Caucasus-Lower Volga and North Pontic Region sources who themselves included ancestry from nearly every native group of the Eurasian steppe, and more distally from ancestors further afield in Southeastern Europe, the Caucasus, and Central Asia. Formed by the confluence of diverse ancestors in the NPR, the Yamna and their descendants then expanded, and often incorporated others in their society in the NPR, proving that their remarkable demographic success did not fully eclipse the people that preceded them.

**Modeling framework**

We used the same qpWave/qpAdm setup[1,2] modeling framework as in the linked paper (ref.[3] to study populations of the North Pontic region (NPR). Briefly, the set of Right outgroup populations is:

**Right:** OldAfrica[4-6], Russia_AfontovaGora3[7], CHG[8], Iran_GanjDareh_N[9], Italy_Villabruna[7], Russia_Sidelkino.SG[10], Turkey_N[11]

This includes an African outgroup, and representatives of "Ancient North Eurasians" (Afontova Gora 3), inland West Asian populations (CHG and Ganj Dareh), Western hunter-gatherers (Villabruna), Eastern hunter-gatherers (Sidelkino), and Anatolian farmers (Turkey_N).

The set of candidate source populations is:

**Sources:** Armenia_Aknashen_N[12], BPgroup[13] and ref.[3], CoreYamna (high data quality set of 104 individuals identified in ref.[3] and including individuals from ref.[3] and ref.[2,10,11,13-18]), GK2 (ref.[3]), GlobularAmphora[16,18], Lebyazhinka_HG[2,16], Maykop[13], PVgroup[13] (grouped in ref.[3]), Remontnoye (ref.[3]), Russia_Karelia[2,16,19], Serbia_IronGates_Mesolithic[16], TTK[20], Trypillia[21] and ref.[3], Ukraine_N[16] and ref.[3], YUN_CA[22]

The set of sources includes population sources for each of the clines involving Pontic-Caspian steppe populations:

- The Volga cline, formed on the basis of North Caucasus-Lower Volga Eneolithic populations (Berezhnovka-2/Progress-2 group / BPgroup or Progress-2/Vonyuchka-1 / PVgroup) admixing with Eastern European hunter-gatherers (EHG) (Russia_Karelia as a remote source of EHG, or Lebyazhinka_HG from the Middle Volga). We also include the Neolithic individual from Tutkaul (TTK) as the Volga cline includes ancestry from an eastern Siberian/Central Asian source that can be modeled with TTK as a source.
- The Dnipro cline, formed with Ukraine Neolithic hunter-gatherer admixture (either Ukraine_N or GK2 from Golubaya Krinitsa on the Middle Don, itself ~2/3 Ukraine_N in ancestry) with a population of the Caucasus-Lower Volga cline (below), using either Core Yamna (as the sampled end point of the cline) or Remontnoye, a population midway on the Caucasus-Lower Volga Cline.
- The Caucasus-Lower Volga (CLV) cline, formed on the basis of largely "Caucasus Neolithic" and descended populations (Aknashen-related) and "Lower Volga Eneolithic" populations (BPgroup-related) with intermediate groups in the North Caucasus at Maykop (largely Caucasus Neolithic) and PVgroup (largely lower Volga Eneolithic) and north of the Manych Depression (at Remontnoye)

To these populations we add three groups representing European farmers: Trypillians (from Moldova and Ukraine), Globular Amphora (from Poland and Ukraine), and YUN_CA (Chalcolithic individuals from Yunatsite, Bulgaria). We also include Iron Gates hunter-gatherers from Serbia (also known as Balkan hunter-gatherers) to account for the possibility that populations of different hunter-gatherer ancestry than the three Neolithic groups could be contributing ancestry to the region.

The Core Yamna, one of the chosen sources, form a tight cluster in PCA and were determined in ref.[3] as comprising of Yamna or Afanasievo individuals that form a clade (using the same Right set as here) to a high degree of confidence (p=0.2 threshold and at least 400,000 autosomal SNPs) with the Samara Yamna (the Yamna population with the highest number of individuals that is part of the Yamna PCA cluster). This cluster does include some high quality individuals from north and west of the Black sea; we will analyze below Yamna individuals from the region that are *not* assigned to the Core Yamna, while also noting those that do.

As we did not want to assume that these 12 source populations admixed only in the combinations manifested in the three Pontic-Caspian clines, we allowed combinations (of $N$=1, 2, 3) populations in our modeling of the following set of Test populations:

**Test:** Bulgaria_C_ProtoYamna, Bulgaria_EBA_Yamna, Bulgaria_Riltsi_EBA_Yamna, Giurgiuleşti, GlobularAmphora, Igren_o, Moldova_Crasnoe_Eneolithic, Moldova_EBA_Yamna, Moldova_EBA_Yamna_o, Moldova_GlobularAmphora_Yamna, Mykhailivka_I32534, SShi, SSlo, SSmed, Trypillia, Ukraine_EBA_Catacomb, Ukraine_EBA_Catacomb_possible, Ukraine_EBA_Yamna, Ukraine_MBA_Catacomb_o1, Ukraine_MBA_MultiCordonedWare_Babine, Ukraine_N_I27992, Usatove, Usatove_EBA, Usatove_I20078, Usatove_Yamna, Zhivotilovka_I17973, Zhivotilovka_I17974

Some of these populations (in particular the Serednii Stih subsets SShi, SSmed, SSlo) were also considered in the linked manuscript ref.[3] but we wanted to see how they might be related in terms of their origin to the others from the NPR.

**An exploratory Principal Component Analysis**

We used smartpca[23] to perform principal component analysis (PCA) using the same set of populations to form the axes as in ref.[3] (Fig. S 1):

OberkasselCluster (set of trans-Alpine WHG individuals identified in[20]), Russia_Firsovo_N, Iran_HajjiFiruz_C[17], Iran_C_SehGabi[9], Iran_C_TepeHissar[24], Israel_C[25], Germany_EN_LBK[2,11,26,27]

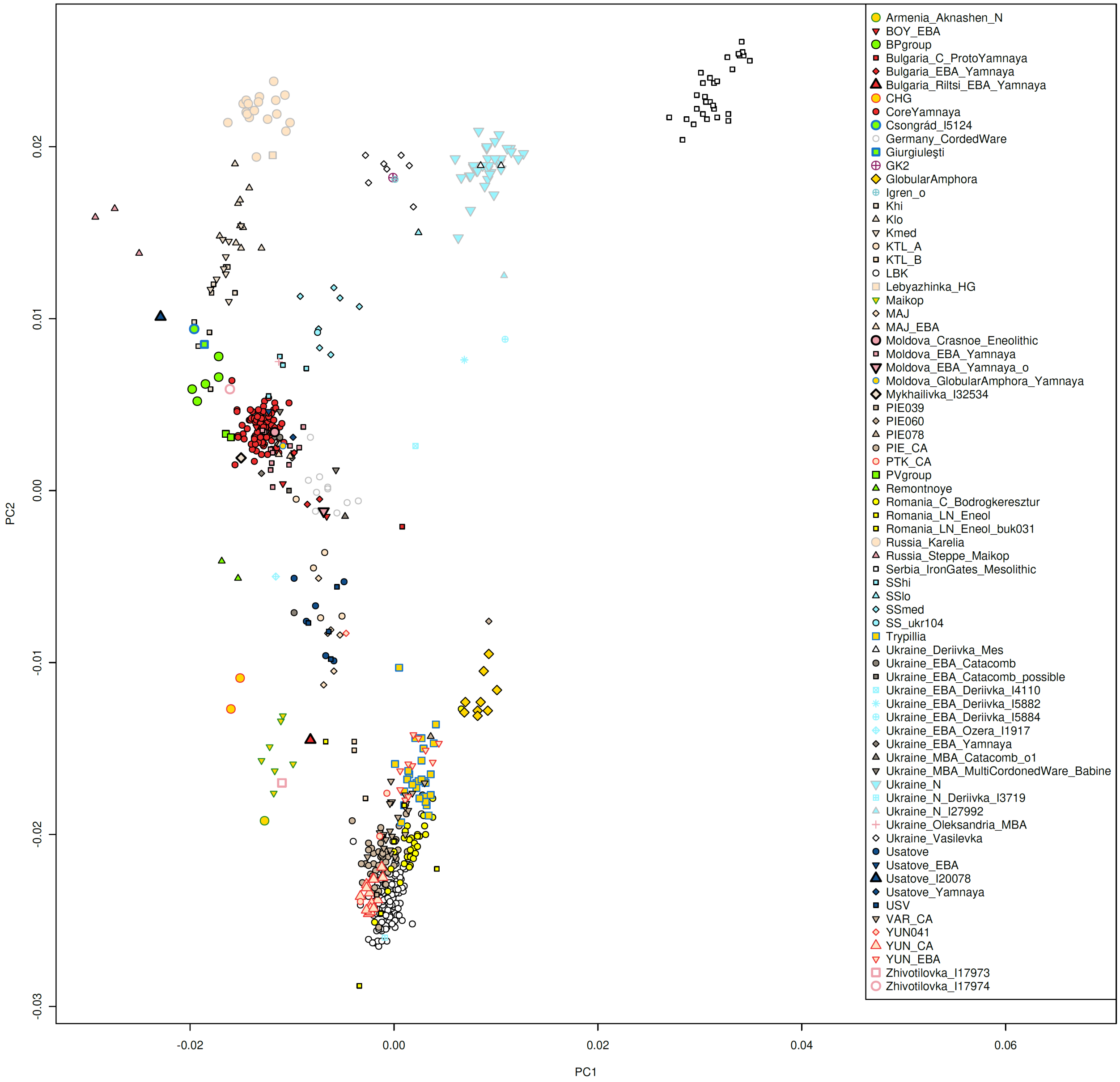

**Fig. S 1 An exploratory PCA.**

We observe that individuals from the North Pontic region and from southeastern Europe are mostly not along the three steppe clines, except, of course, the aforementioned Serednii Stih individuals that define the Dnipro cline. A Serednii Stih individual (SS_ukr104) published by Mattila et al.[28] clusters with the SSmed subset and forms a clade with it using qpWave (p=0.281).

Most individuals of interest are to the right and bottom of the PCA (with respect to the Dnipro cline), towards the cline formed by European farmers (LBK and YUN_CA) on one side and Balkan hunter-gatherers (Serbia Iron Gates Mesolithic). Clearly ancestry from European farmers of mixed hunter-gatherer ancestry played a role in their formation which we formally explore below. We highlight here a few individuals that are unusual in that they are shifted *away* from the farmer-hunter-gatherer cline of southeastern Europe. Two of these are an individual from Zhivotilovka (I17974) and an "Usatove" outlier (I20078) from Taraclia. Two other individuals (from Giurgiuleşti in Moldova and Csongrád in Hungary) are also similarly shifted and near the BPgroup (Lower Volga-North Caucasus Eneolithic group) at one end of the Volga cline.

The Core Yamna clearly played a key role in events of the Bronze Age. One individual of interest is Mykhailivka_I32534 from Ukraine and another Moldova_Crasnoe_Eneolithic (individual I20196) which cluster with the core Yamna itself. Note that in the very definition of the core Yamna exist individuals from Ukraine, Moldova, Romania, and Bulgaria (ref.[3]). But many other Yamna-related individuals deviate towards the European farmer cline, and as we will see, we find examples of Yamna ancestry combining with diverse pre-existing populations both west and east of the NPR.

**Populations that form a clade (N=1) with one of the sources.**

The following Test populations form a clade (using a p-value cutoff of 0.05) with one of the sources (Table S 1):

| Test | Source | P-value |
|---|---|---|
| Usatove_Yamna | CoreYamna | 0.178 |
| Ukraine_EBA_Catacomb | CoreYamna | 0.075 |
| Giurgiuleşti | BPgroup | 0.896 |
| Giurgiuleşti | PVgroup | 0.689 |
| Usatove_EBA | CoreYamna | 0.864 |
| Usatove_EBA | PVgroup | 0.067 |
| Moldova_Crasnoe_Eneolithic | CoreYamna | 0.683 |
| Igren_o | GK2 | 0.991 |
| Mykhailivka_I32534 | BPgroup | 0.163 |
| Mykhailivka_I32534 | CoreYamna | 0.684 |
| Mykhailivka_I32534 | PVgroup | 0.659 |

**Table S 1 Populations that can be modeled as simple clades (*N*=1) with one of the sources**

Usatove_Yamna is individual I12229_enhanced (3088-2911 calBCE) with a potentially Usatove cultural affiliation but a Yamna genetic profile and its label reflects these facts. Also cladal with the Core Yamna are the individuals of population "Ukraine_EBA_Catacomb" (p=0.075). The individual from Giurgiuleşti (I20072) is cladal with the Lower Volga-North Caucasus Eneolithic groups (BPgroup and PVgroup) and represents an example of long-range migration from the east into Moldova (4330-4058 calBCE). Usatove_EBA (individual I20079) is also cladal with the Yamna, later than the expansion of the Yamna and the aforementioned Usatove_Yamna individual (2571-2355 calBCE).

Moldova_Crasnoe_Eneolithic (individual I20196; 3352-3101 calBCE) is dated to the Yamna expansion and has a Yamna genetic profile. Igren_o (I27930) is analyzed in ref.[3] and appears to be of hunter-gatherer ancestry similar to the GK2 individual from Golubaya Krinitsa and unlike the Serednii Stih individuals of similar age from Igren.

**Mykhailivka_I32534**

Finally, the low coverage individual I32534 is a clade with Yamna, although we can also fit it as a clade with the Lower Volga-Caucasus Eneolithic groups BPgroup and PVgroup. This individual has a very old date of 3635-3383 calBCE which potentially establishes the early presence of genetically Yamna individuals in the Dnipro region which is consistent with the idea of Yamna being a late Serednii Stih population from the Dnipro-Don area.

To determine if some of the models of Table S 1 can be rejected, we used a model competition approach in which we tested the resilience of each model (with BPgroup, Core Yamna, and PVgroup as the source) when placing the alternative sources in the Right set. The results (Table S 2) indicate that only the Core

Yamna model is resilient to either BPgroup or PVgroup being placed on the Right; i.e., neither BPgroup nor PVgroup share any common genetic history with the I32534 that cannot be well-modeled as Core Yamna. By contrast, both BPgroup and PVgroup are rendered infeasible in at least one of their competition comparisons (and their p-values are low 0.1-0.11 for the other one). This comparison suggests that only the Core Yamna model is unambiguously feasible for this individual.

| | BPgroup | CoreYamna | PVgroup |
|---|---|---|---|
| BPgroup | | 5.5E-01 | 6.9E-05 |
| CoreYamna | 1.0E-01 | | 1.1E-01 |
| PVgroup | 4.2E-02 | 7.7E-01 | |

**Table S 2 Model competition for Mykhailivka_I32534.** Each column shows the p-value of a model when alternative models (rows) are placed on the Right set.

We were also concerned that the individual could fit as a clade with the Core Yamna even though it might not be "exactly" Core Yamna but may harbor other ancestry that is not, however, sufficient to cause the basic Core Yamna model to fail.

First, we considered whether the individual could harbor less NPR hunter-gatherer ancestry than the Core Yamna, as it is on that far end of the cline (Fig. S 1; beyond the main Core Yamna cluster and away from NPR hunter-gatherers like Ukraine_N and GK2). Adding either Ukraine_N (-6.2±7.9%; p=0.689) or GK2 (-3.5±9.9%; p=0.543) is consistent with this position, but the negative proportions of NPR ancestry are non-significant. Thus, we can say that it is *possible* that this early individual might have slightly less NPR hunter-gatherer ancestry than the Core Yamna cluster, this is *unproven*. To the limits of our resolution, the Mykhailivka individual is within the Core Yamna variation.

Second, we considered whether the individual might harbor some European farmer ancestry. This is a topic that we address below for other populations of Ukraine, Moldova, and Bulgaria, i.e., from the western portion of the Yamna range. However, when we add European farmer sources as a 2$^{nd}$ source, their inferred proportion is again negative and non-significant (-5.4±5.1% for Globular Amphora; -4.9±5.6% for Trypillia; -3.3±4.8% for YUN_CA). Thus there is no evidence at all for the presence of farmer ancestry in the Mykhailivka individual which appears, in this comparison as well, to be simply Core Yamna.

##### Admixture $f_3$-statistics

To test for the presence of admixture we computed $f_3$-statistics[29] of the form $f_3$(Source$_1$, Source$_2$; Test) for all Test populations with at least two individuals using inbreed: YES in qp3Pop. A significantly negative statistic of this form proves that admixture is present in the Test population, thus we show the lowest value of the Z-score of these statistics in Extended Data Table 1. Note that the pair of sources that minimizes the statistic are not necessarily those that are involved in the admixture. It is further possible that populations have admixture by more than two sources. The formal modeling in terms of sources using qpAdm will follow in this note. However, the simple $f_3$-statistics are useful as an initial test of the presence of admixture and complement the visual impressions of Fig. S 1 in which some populations appear to be "intermediate" between others, a situation that may arise in the presence of admixture. Our statistics prove -if significantly negative- that admixture took place; however, if they are not significantly negative or even positive it is still possible that the population is admixed, either between sources not available to us or because of post-admixture genetic drift which masks the signal of admixture.[29]

It is clear from Extended Data Table 1 that admixture is present in many of the Test populations. We next proceed to model this admixture.

| Test | A | B | P-value | A | B | S.E. |
|---|---|---|---|---|---|---|

| | | | | | | |
|---|---|---|---|---|---|---|
| Bulgaria_C_ProtoYamna | CoreYamna | GlobularAmphora | 0.099 | 45.3% | 54.7% | 2.7% |
| Bulgaria_EBA_Yamna | Armenia_Aknashen_N | Lebyazhinka_HG | 0.082 | 55.9% | 44.1% | 1.7% |
| Bulgaria_EBA_Yamna | BPgroup | GlobularAmphora | 0.812 | 70.8% | 29.2% | 1.5% |
| Bulgaria_EBA_Yamna | BPgroup | Trypillia | 0.246 | 70.4% | 29.6% | 1.5% |
| Bulgaria_EBA_Yamna | CoreYamna | GlobularAmphora | 0.310 | 83.3% | 16.7% | 1.6% |
| Bulgaria_EBA_Yamna | CoreYamna | Trypillia | 0.728 | 82.8% | 17.2% | 1.6% |
| Bulgaria_EBA_Yamna | CoreYamna | YUN_CA | 0.736 | 85.7% | 14.3% | 1.4% |
| Bulgaria_EBA_Yamna | GlobularAmphora | PVgroup | 0.458 | 26.2% | 73.8% | 1.9% |
| Bulgaria_EBA_Yamna | Lebyazhinka_HG | Maykop | 0.189 | 38.5% | 61.5% | 1.7% |
| Bulgaria_Riltsi_EBA_Yamna | Armenia_Aknashen_N | GlobularAmphora | 0.361 | 61.4% | 38.6% | 6.2% |
| Bulgaria_Riltsi_EBA_Yamna | Armenia_Aknashen_N | Trypillia | 0.212 | 56.4% | 43.6% | 7.0% |
| Bulgaria_Riltsi_EBA_Yamna | BPgroup | YUN_CA | 0.124 | 38.2% | 61.8% | 3.3% |
| Bulgaria_Riltsi_EBA_Yamna | CoreYamna | YUN_CA | 0.058 | 42.9% | 57.1% | 3.7% |
| Bulgaria_Riltsi_EBA_Yamna | GlobularAmphora | Maykop | 0.354 | 38.8% | 61.2% | 6.1% |
| Bulgaria_Riltsi_EBA_Yamna | Maykop | Trypillia | 0.538 | 55.8% | 44.2% | 6.3% |
| Bulgaria_Riltsi_EBA_Yamna | Maykop | YUN_CA | 0.103 | 63.7% | 36.3% | 5.4% |
| Bulgaria_Riltsi_EBA_Yamna | PVgroup | YUN_CA | 0.335 | 40.4% | 59.6% | 3.4% |
| Bulgaria_Riltsi_EBA_Yamna | Remontnoye | Trypillia | 0.075 | 42.9% | 57.1% | 5.3% |
| Bulgaria_Riltsi_EBA_Yamna | Remontnoye | YUN_CA | 0.558 | 50.3% | 49.7% | 4.5% |
| Moldova_EBA_Yamna | BPgroup | GlobularAmphora | 0.300 | 78.9% | 21.1% | 1.1% |
| Moldova_EBA_Yamna | CoreYamna | GlobularAmphora | 0.511 | 93.0% | 7.0% | 1.0% |
| Moldova_EBA_Yamna | CoreYamna | Trypillia | 0.545 | 92.8% | 7.2% | 1.0% |
| Moldova_EBA_Yamna | CoreYamna | YUN_CA | 0.361 | 94.0% | 6.0% | 0.9% |
| Moldova_EBA_Yamna | GK2 | Remontnoye | 0.672 | 29.6% | 70.4% | 2.1% |
| Moldova_EBA_Yamna_o | BPgroup | GlobularAmphora | 0.473 | 53.0% | 47.0% | 2.4% |
| Moldova_EBA_Yamna_o | BPgroup | Trypillia | 0.321 | 52.1% | 47.9% | 2.4% |
| Moldova_EBA_Yamna_o | CoreYamna | GlobularAmphora | 0.265 | 62.4% | 37.6% | 2.7% |
| Moldova_EBA_Yamna_o | CoreYamna | Trypillia | 0.523 | 61.3% | 38.7% | 2.8% |
| Moldova_EBA_Yamna_o | CoreYamna | YUN_CA | 0.191 | 67.9% | 32.1% | 2.4% |
| Moldova_EBA_Yamna_o | GlobularAmphora | PVgroup | 0.831 | 44.7% | 55.3% | 2.7% |
| Moldova_EBA_Yamna_o | PVgroup | Trypillia | 0.136 | 54.5% | 45.5% | 2.8% |
| Moldova_GlobularAmphora_Yamna | BPgroup | GlobularAmphora | 0.052 | 74.5% | 25.5% | 2.2% |
| Moldova_GlobularAmphora_Yamna | CoreYamna | GlobularAmphora | 0.180 | 88.0% | 12.0% | 2.5% |
| Moldova_GlobularAmphora_Yamna | CoreYamna | Trypillia | 0.057 | 88.6% | 11.4% | 2.6% |
| Moldova_GlobularAmphora_Yamna | GK2 | Remontnoye | 0.395 | 33.3% | 66.7% | 3.7% |
| Moldova_GlobularAmphora_Yamna | Lebyazhinka_HG | Maykop | 0.091 | 44.1% | 55.9% | 2.5% |
| Moldova_GlobularAmphora_Yamna | Remontnoye | Ukraine_N | 0.388 | 69.1% | 30.9% | 3.1% |
| SShi | CoreYamna | GK2 | 0.134 | 83.7% | 16.3% | 3.0% |
| SShi | CoreYamna | Lebyazhinka_HG | 0.075 | 85.8% | 14.2% | 2.6% |
| SShi | CoreYamna | Russia_Karelia | 0.058 | 87.5% | 12.5% | 2.3% |
| SShi | GK2 | PVgroup | 0.088 | 30.3% | 69.7% | 3.2% |
| SShi | PVgroup | Ukraine_N | 0.135 | 74.0% | 26.0% | 2.5% |
| SSlo | Armenia_Aknashen_N | GK2 | 0.204 | 15.4% | 84.6% | 3.8% |
| SSlo | BPgroup | Serbia_IronGates_Mesolithic | 0.063 | 59.5% | 40.5% | 3.4% |
| SSlo | BPgroup | Ukraine_N | 0.885 | 33.9% | 66.1% | 5.2% |
| SSlo | CoreYamna | GK2 | 0.142 | 31.4% | 68.6% | 7.5% |
| SSlo | CoreYamna | Ukraine_N | 0.750 | 37.8% | 62.2% | 5.7% |
| SSlo | GK2 | Maykop | 0.283 | 83.2% | 16.8% | 4.0% |
| SSlo | GK2 | PVgroup | 0.079 | 76.1% | 23.9% | 6.7% |
| SSlo | GK2 | Remontnoye | 0.314 | 78.1% | 21.9% | 5.3% |
| SSlo | PVgroup | Ukraine_N | 0.906 | 31.4% | 68.6% | 5.1% |
| SSlo | Remontnoye | Ukraine_N | 0.264 | 25.6% | 74.4% | 4.4% |
| SSmed | CoreYamna | GK2 | 0.520 | 60.6% | 39.4% | 3.2% |
| SSmed | CoreYamna | Ukraine_N | 0.309 | 65.5% | 34.5% | 2.3% |
| SSmed | GK2 | Remontnoye | 0.740 | 58.0% | 42.0% | 2.6% |
| Ukraine_EBA_Yamna | BPgroup | GlobularAmphora | 0.265 | 77.7% | 22.3% | 1.5% |
| Ukraine_EBA_Yamna | CoreYamna | GlobularAmphora | 0.853 | 91.9% | 8.1% | 1.6% |
| Ukraine_EBA_Yamna | CoreYamna | Trypillia | 0.697 | 91.9% | 8.1% | 1.6% |
| Ukraine_EBA_Yamna | CoreYamna | YUN_CA | 0.295 | 93.6% | 6.4% | 1.4% |
| Ukraine_EBA_Yamna | GK2 | Remontnoye | 0.781 | 30.9% | 69.1% | 2.6% |
| Ukraine_MBA_Catacomb_o1 | Armenia_Aknashen_N | CoreYamna | 0.083 | 42.4% | 57.6% | 9.3% |
| Ukraine_MBA_Catacomb_o1 | Armenia_Aknashen_N | GK2 | 0.301 | 63.5% | 36.5% | 6.0% |
| Ukraine_MBA_Catacomb_o1 | Armenia_Aknashen_N | Lebyazhinka_HG | 0.272 | 66.8% | 33.2% | 5.2% |
| Ukraine_MBA_Catacomb_o1 | Armenia_Aknashen_N | Russia_Karelia | 0.188 | 69.7% | 30.3% | 4.9% |
| Ukraine_MBA_Catacomb_o1 | Armenia_Aknashen_N | Ukraine_N | 0.145 | 65.5% | 34.5% | 5.6% |
| Ukraine_MBA_Catacomb_o1 | BPgroup | GlobularAmphora | 0.253 | 59.3% | 40.7% | 5.3% |

| | | | | | | |
|---|---|---|---|---|---|---|
| Ukraine_MBA_Catacomb_o1 | BPgroup | Trypillia | 0.410 | 56.4% | 43.6% | 5.3% |
| Ukraine_MBA_Catacomb_o1 | BPgroup | YUN_CA | 0.560 | 62.8% | 37.2% | 4.5% |
| Ukraine_MBA_Catacomb_o1 | CoreYamna | GlobularAmphora | 0.162 | 69.6% | 30.4% | 6.2% |
| Ukraine_MBA_Catacomb_o1 | CoreYamna | Trypillia | 0.264 | 66.2% | 33.8% | 6.4% |
| Ukraine_MBA_Catacomb_o1 | CoreYamna | YUN_CA | 0.458 | 71.7% | 28.3% | 5.1% |
| Ukraine_MBA_Catacomb_o1 | GK2 | Maykop | 0.227 | 30.8% | 69.2% | 6.1% |
| Ukraine_MBA_Catacomb_o1 | GlobularAmphora | PVgroup | 0.551 | 35.6% | 64.4% | 5.3% |
| Ukraine_MBA_Catacomb_o1 | GlobularAmphora | Remontnoye | 0.507 | 23.1% | 76.9% | 6.5% |
| Ukraine_MBA_Catacomb_o1 | Lebyazhinka_HG | Maykop | 0.226 | 27.5% | 72.5% | 5.4% |
| Ukraine_MBA_Catacomb_o1 | Maykop | Russia_Karelia | 0.171 | 75.1% | 24.9% | 5.0% |
| Ukraine_MBA_Catacomb_o1 | Maykop | Ukraine_N | 0.130 | 71.2% | 28.8% | 5.6% |
| Ukraine_MBA_Catacomb_o1 | PVgroup | Trypillia | 0.531 | 61.8% | 38.2% | 5.9% |
| Ukraine_MBA_Catacomb_o1 | PVgroup | YUN_CA | 0.454 | 68.0% | 32.0% | 4.9% |
| Ukraine_MBA_Catacomb_o1 | Remontnoye | Trypillia | 0.354 | 75.1% | 24.9% | 7.1% |
| Ukraine_MBA_Catacomb_o1 | Remontnoye | YUN_CA | 0.186 | 81.6% | 18.4% | 6.0% |
| Ukraine_MBA_MultiCordonedWare_Babine | CoreYamna | Serbia_IronGates_Mesolithic | 0.128 | 93.7% | 6.3% | 2.2% |
| Ukraine_MBA_MultiCordonedWare_Babine | CoreYamna | Ukraine_N | 0.053 | 90.9% | 9.1% | 3.5% |
| Ukraine_MBA_MultiCordonedWare_Babine | GK2 | Remontnoye | 0.153 | 36.1% | 63.9% | 3.6% |
| Ukraine_N_I27992 | Armenia_Aknashen_N | GK2 | 0.313 | 29.9% | 70.1% | 8.2% |
| Ukraine_N_I27992 | Armenia_Aknashen_N | Ukraine_N | 0.238 | 30.8% | 69.2% | 7.5% |
| Ukraine_N_I27992 | CoreYamna | Serbia_IronGates_Mesolithic | 0.073 | 66.5% | 33.5% | 8.5% |
| Ukraine_N_I27992 | GK2 | GlobularAmphora | 0.409 | 67.0% | 33.0% | 8.1% |
| Ukraine_N_I27992 | GK2 | Maykop | 0.366 | 68.9% | 31.1% | 8.9% |
| Ukraine_N_I27992 | GK2 | Trypillia | 0.535 | 69.0% | 31.0% | 7.8% |
| Ukraine_N_I27992 | GK2 | YUN_CA | 0.640 | 74.4% | 25.6% | 6.7% |
| Ukraine_N_I27992 | GlobularAmphora | Lebyazhinka_HG | 0.454 | 40.1% | 59.9% | 6.9% |
| Ukraine_N_I27992 | GlobularAmphora | Russia_Karelia | 0.243 | 46.3% | 53.7% | 6.6% |
| Ukraine_N_I27992 | GlobularAmphora | TTK | 0.107 | 53.3% | 46.7% | 5.8% |
| Ukraine_N_I27992 | Lebyazhinka_HG | Trypillia | 0.356 | 62.3% | 37.7% | 6.6% |
| Ukraine_N_I27992 | Lebyazhinka_HG | YUN_CA | 0.292 | 68.7% | 31.3% | 5.9% |
| Ukraine_N_I27992 | Maykop | Ukraine_N | 0.279 | 32.5% | 67.5% | 8.2% |
| Ukraine_N_I27992 | Remontnoye | Serbia_IronGates_Mesolithic | 0.084 | 57.0% | 43.0% | 7.4% |
| Ukraine_N_I27992 | Remontnoye | Ukraine_N | 0.262 | 38.5% | 61.5% | 9.6% |
| Ukraine_N_I27992 | Russia_Karelia | Trypillia | 0.171 | 56.0% | 44.0% | 6.2% |
| Ukraine_N_I27992 | Russia_Karelia | YUN_CA | 0.114 | 63.0% | 37.0% | 5.3% |
| Ukraine_N_I27992 | Trypillia | Ukraine_N | 0.111 | 29.3% | 70.7% | 7.8% |
| Ukraine_N_I27992 | Ukraine_N | YUN_CA | 0.155 | 75.8% | 24.2% | 6.6% |
| Usatove | PVgroup | Trypillia | 0.128 | 45.4% | 54.6% | 1.5% |
| Usatove_I20078 | BPgroup | TTK | 0.091 | 79.1% | 20.9% | 4.5% |
| Usatove_I20078 | CoreYamna | TTK | 0.606 | 65.2% | 34.8% | 3.6% |
| Usatove_I20078 | PVgroup | TTK | 0.072 | 72.2% | 27.8% | 4.4% |
| Zhivotilovka_I17974 | CoreYamna | Lebyazhinka_HG | 0.077 | 92.1% | 7.9% | 4.0% |
| Zhivotilovka_I17974 | CoreYamna | TTK | 0.616 | 89.0% | 11.0% | 3.4% |
| Zhivotilovka_I17974 | GK2 | PVgroup | 0.597 | 18.7% | 81.3% | 4.2% |
| Zhivotilovka_I17974 | Lebyazhinka_HG | PVgroup | 0.161 | 17.3% | 82.7% | 4.5% |
| Zhivotilovka_I17974 | PVgroup | Russia_Karelia | 0.135 | 85.1% | 14.9% | 3.8% |
| Zhivotilovka_I17974 | PVgroup | Serbia_IronGates_Mesolithic | 0.083 | 91.6% | 8.4% | 2.4% |
| Zhivotilovka_I17974 | PVgroup | Ukraine_N | 0.401 | 84.5% | 15.5% | 3.6% |

**Table S 3 Populations that can be modeled as 2-way mixtures clades (*N*=2) of the sources**

**Populations that can be modeled as mixtures of two of the sources (*N*=2)**

We next examined populations for which there are feasible models of 2-way admixture (Table S 3). We define feasible models as having p>0.05, standard errors ≤10%, and admixture proportions within two standard errors of the endpoints of the [0, 1] interval. Alternative models exist for many of these populations. We did not discuss further the Serednii Stih subsets (SShi, SSmed, SSlo) here, except to note that they are not modeled with European farmer populations as sources (YUN_CA or Trypillia) but with sources that are included in the analysis of the linked study ref.[3] to which we refer the reader. Here we can summarize that the Serednii Stih can be modeled with Core Yamna (the end point of the Dnipro cline) and Dnipro-Don hunter-gatherers (Ukraine_N or GK2), or with populations of the CLV cline (PVgroup, BPgroup, Remontnoye, or Maykop) and on the whole can be seen as the result of fusion of CLV cline migrants with local Dnipro-Don foragers that is generative of the Dnipro cline.

We use the model tournament approach of all-against-all to see if we can distinguish between some of the alternative models.

**Bulgaria_C_ProtoYamna (3500-3000 BCE)**

For this individual (I1456 / 3500-3000 BCE) from Durankulak only a single 2-way feasible model exists that involves a fairly even mix of Core Yamna (~45%) and Globular Amphora (~55%) ancestries (Table S 3). Models with Trypillian (p=1e-5) or YUN_CA (p=3e-13) alternative sources of European farmer ancestry both fail.

It has been postulated that the Globular Amphora was the source of farmer ancestry in the Corded Ware population of central-northern Europe on the basis of the fact that this European farmer population shares IBD segments with the Corded Ware.[30] The proportion of farmer ancestry in the Corded Ware and diverse derived populations is fairly even[31] and amounts to about ~1/4 as estimated in ref.[2] Thus, the Corded Ware represents a mixture of similar components as the Bulgarian "Proto-Yamna" individual, albeit in different proportions.

We applied the Core Yamna + Globular Amphora to Corded Ware individuals from Germany and obtained an estimate of 26.6±1.3% (p=0.034). For both the Proto-Yamna individual and the Corded Ware individual, we noticed that shared genetic drift with the Villabruna hunter-gatherer (from the Base set of outgroups) is underestimated (Z<-2.5), suggesting that there is an excess of hunter-gatherer ancestry (above and beyond what was part of the Globular Amphora population sample we have) in both. Adding Serbia Iron Gates hunter-gatherers as a 3$^{rd}$ source results in a successful fit in both Proto-Yamna (p=0.328) and Corded Ware (p=0.156) with a significant proportion of such ancestry (6.1±2.6% and 2.9±1.2% respectively). We also added Latvia_HG[16] from the Baltic region as the 3$^{rd}$ source inspired by the model of ref.[32] which includes an amount of northeastern European ancestry in the Corded Ware. This model also fits for both the Proto-Yamna (6.2±2.9%; p=0.262) and Corded Ware (3.2±1.3%; p=0.135) and thus the source of the Villabruna-related ancestry in the Corded Ware and Proto-Yamna cannot be well-determined geographically. As the territory of the Globular Amphora is located in-between the Balkans and the Baltic, we think it likely that the Corded Ware was formed by admixture with a local Globular Amphora-related population that had a slight excess of hunter-gatherer ancestry rather than an extra pulse of admixture from either the Balkans or the Baltic.

We were motivated to pursue the Corded Ware connection by the additional observation that the Bulgarian Proto-Yamna individual belonged to Y-haplogroup R-M417.[33] This lineage is typical of the Corded Ware populations[2,14,31,32] We offer the hypothesis that the Proto-Yamna individual is connected to the formation of the Corded Ware population as its autosomal ancestry points to the same sources (Core Yamna and Globular Amphora with a slight excess of hunter-gatherer ancestry) and was part of the same Y-chromosome lineage whose origin prior to the appearance of the Corded Ware is unclear.

Possibly, both the Proto-Yamna individual and the nascent Corded Ware were derived from a zone of interaction between Yamna-related and Globular Amphora-related populations which must have generated a cline of variable ancestry between the two sources: the Corded Ware emerged as an expansive archaeological culture in which ~1/4 of the Globular Amphora ancestry was incorporated while the Bulgarian Proto-Yamna individual had even more Globular Amphora ancestry.

As a final test, we fit the Proto-Yamna individual as Corded Ware + Globular Amphora (p=0.438) with an estimated 37.4±3.7% Globular Amphora. Given that Corded Ware itself has 26.6±1.3% of Globular Amphora ancestry, the total amounts to (1-0.374)*0.266+0.374≈54%, close to the directly estimated 54.7±2.7% Globular Amphora ancestry (Table S 3) from the Core Yamna + Globular Amphora model.

**Bulgaria_EBA_Yamna (3300-2000 BCE)**

For the Yamna from Bulgaria, we see that the only models that are feasible in model competition are the ones that involve Core Yamna as one source and a European farmer population as the other (Table S 4).

Models without Core Yamna as a source are feasible against the Base set of outgroups (Table S 3) but are not resilient to the addition of Core Yamna to the set of outgroups. Model competition does not allow us to decide between the different European farmer populations, but the models agree in deriving a similar proportion of such ancestry (~14-17%) from the farmer source. It is thus not clear whether the farmer ancestry was of local origin in Bulgaria or was acquired north of Bulgaria (where Trypillians and Globular Amphora-related populations lived).

| | BPgroup+GlobularAmphora | BPgroup+Trypillia | CoreYamna+GlobularAmphora | CoreYamna+Trypillia | CoreYamna+YUN_CA | Armenia_Aknashen_N+Lebyazhinka_HG | GlobularAmphora+PVgroup | Lebyazhinka_HG+Maykop |
|---|---|---|---|---|---|---|---|---|
| BPgroup+GlobularAmphora | | 3.18E-01 | 4.40E-01 | 8.22E-01 | 8.49E-01 | 4.05E-05 | 3.23E-03 | 1.93E-07 |
| BPgroup+Trypillia | 8.97E-01 | | 1.58E-01 | 8.22E-01 | 5.79E-01 | 2.68E-05 | 4.06E-03 | 5.85E-08 |
| CoreYamna+GlobularAmphora | 8.60E-16 | 2.32E-15 | | 7.19E-01 | 7.94E-01 | 1.47E-31 | 1.05E-15 | 3.87E-38 |
| CoreYamna+Trypillia | 3.71E-15 | 5.91E-16 | 1.17E-01 | | 5.63E-01 | 1.41E-34 | 3.54E-15 | 8.08E-38 |
| CoreYamna+YUN_CA | 2.79E-15 | 2.51E-15 | 4.50E-01 | 8.31E-01 | | 4.55E-31 | 4.13E-15 | 4.68E-37 |
| Armenia_Aknashen_N+Lebyazhinka_HG | 2.60E-01 | 1.51E-02 | 1.26E-01 | 1.72E-01 | 1.37E-01 | | 2.88E-02 | 1.57E-01 |
| GlobularAmphora+PVgroup | 7.03E-02 | 1.91E-02 | 2.61E-01 | 6.24E-01 | 7.94E-01 | 7.30E-05 | | 2.11E-04 |
| Lebyazhinka_HG+Maykop | 4.82E-01 | 8.42E-02 | 4.48E-01 | 7.65E-01 | 6.16E-01 | 8.09E-02 | 6.07E-02 | |

**Table S 4 Model competition for Bulgaria_EBA_Yamna.** Each column shows the p-value of a model when alternative models (rows) are placed on the Right set.

We fit the Core Yamna+YUN_CA model to individuals within the Bulgarian Yamna group (Table S 5). We see that this group included both individuals that were unadmixed Core Yamna as well as others with a significant amount of farmer ancestry. Thus, while we cannot be certain that the admixture between European farmers and Yamna occurred locally in Bulgaria, there were certainly core Yamna individuals present there, so the opportunity for such local admixture existed. These results are consistent with the penetration of southeastern Europe by Yamna well beyond the geographical transition zone of the steppe by Yamna individuals and their admixture with local farmers.[12]

| Test | P-value | CoreYamna | YUN_CA | S.E. |
|---|---|---|---|---|
| I1427 | 0.436 | 100.9% | -0.9% | 2.8% |
| I1448 | 0.639 | 77.7% | 22.3% | 3.1% |
| I18793 | 0.889 | 89.0% | 11.0% | 2.2% |
| I18794 | 0.696 | 78.3% | 21.7% | 2.6% |
| I18801_d | 0.628 | 99.0% | 1.0% | 3.5% |

**Table S 5 Admixture of Bulgarian Yamna individuals.**

#### Bulgaria_Riltsi_EBA_Yamna (3300-2500 BCE)

Multiple models apply to the Riltsi Yamna (Table S 3) which is analyzed separately from the other Yamna from Bulgaria. It is clear that in model competition, models with core Yamna as a source do not fit best (Table S 6), but rather two models that include local (YUN_CA) ancestry and ancestry from the CLV Cline (either PVgroup or Remontnoye) fit better. These models only lose a match to another model of Maykop+Trypillian ancestry. Since Maykop is itself part of the CLV Cline, we can conclude that the Riltsi Yamna individual (I1428; 3300-2500 BCE) are descended from CLV cline populations that admixed with European farmers and not from the Core Yamna as the other Yamna individuals from southeastern Europe.

| | Armenia_Aknashen_N+Trypillia | BPgroup+YUN_CA | CoreYamna+YUN_CA | Armenia_Aknashen_N+GlobularAmphora | GlobularAmphora+Maykop | Maykop+Trypillia | Maykop+YUN_CA | PVgroup+YUN_CA | Remontnoye+Trypillia | Remontnoye+YUN_CA |
|---|---|---|---|---|---|---|---|---|---|---|
| Armenia_Aknashen_N+Trypillia | | 4.46E-02 | 3.61E-02 | 9.91E-02 | 1.11E-02 | 7.01E-01 | 3.81E-04 | 6.25E-02 | 1.52E-01 | 1.24E-01 |
| BPgroup+YUN_CA | 6.26E-03 | | 7.10E-02 | 1.51E-02 | 3.23E-02 | 4.89E-02 | 1.92E-03 | 4.21E-01 | 1.66E-01 | 5.26E-01 |
| CoreYamna+YUN_CA | 3.34E-06 | 1.13E-01 | | 6.63E-06 | 3.98E-04 | 6.64E-04 | 8.09E-06 | 2.33E-01 | 9.77E-02 | 2.90E-01 |
| Armenia_Aknashen_N+GlobularAmphora | 3.16E-01 | 2.96E-01 | 1.64E-01 | | 4.62E-01 | 8.10E-01 | 2.27E-01 | 6.10E-01 | 2.19E-01 | 8.16E-01 |
| GlobularAmphora+Maykop | 5.57E-02 | 2.60E-02 | 1.60E-02 | 9.31E-02 | | 6.84E-01 | 1.55E-01 | 1.20E-01 | 2.21E-02 | 2.65E-01 |
| Maykop+Trypillia | 4.03E-02 | 5.66E-03 | 5.59E-03 | 3.79E-02 | 4.79E-03 | | 1.19E-04 | 1.53E-02 | 1.60E-02 | 4.14E-02 |
| Maykop+YUN_CA | 6.50E-02 | 1.51E-02 | 9.08E-03 | 1.57E-01 | 3.85E-01 | 6.29E-01 | | 8.37E-02 | 3.18E-02 | 2.10E-01 |
| PVgroup+YUN_CA | 9.23E-02 | 2.09E-01 | 1.05E-01 | 2.11E-01 | 3.21E-01 | 4.29E-01 | 5.61E-02 | | 2.16E-01 | 6.93E-01 |
| Remontnoye+Trypillia | 3.40E-01 | 4.53E-02 | 3.65E-02 | 2.01E-01 | 1.20E-02 | 6.87E-01 | 4.48E-04 | 6.06E-02 | | 6.97E-02 |
| Remontnoye+YUN_CA | 4.15E-01 | 1.83E-01 | 9.09E-02 | 6.40E-01 | 5.30E-01 | 7.66E-01 | 1.80E-01 | 4.37E-01 | 1.32E-01 | |

**Table S 6 Model competition for Bulgaria_Riltsi_EBA_Yamna.** Each column shows the p-value of a model when alternative models (rows) are placed on the Right set.

#### Moldova_EBA_Yamna (3400-2500BCE)

The model tournament for the Yamna of Moldova (Table S 7) shows that it is the models with Core Yamna and European farmer (Trypillian or YUN_CA) ancestry that fare best.

| | BPgroup+GlobularAmphora | CoreYamna+GlobularAmphora | CoreYamna+Trypillia | CoreYamna+YUN_CA | GK2+Remontnoye |
|---|---|---|---|---|---|
| BPgroup+GlobularAmphora | | 6.55E-01 | 7.40E-01 | 5.77E-01 | 4.25E-02 |
| CoreYamna+GlobularAmphora | 2.50E-29 | | 6.24E-01 | 4.61E-01 | 6.29E-52 |
| CoreYamna+Trypillia | 1.99E-29 | 5.47E-05 | | 1.79E-04 | 1.98E-53 |
| CoreYamna+YUN_CA | 8.04E-29 | 7.03E-03 | 4.15E-03 | | 3.02E-46 |
| GK2+Remontnoye | 3.80E-01 | 7.70E-01 | 8.41E-01 | 6.90E-01 | |

**Table S 7 Model competition for Moldova_EBA_Yamna.** Each column shows the p-value of a model when alternative models (rows) are placed on the Right set.

When we look at individuals (Table S 8) we see that they have inherited different amounts of European farmer ancestry. Four individuals are included in our Core Yamna definition so by definition we do not provide estimates of European farmer ancestry in them (10208, I10398, I12637, I7847). in addition, some of the other modeled Moldovan Yamna individuals are consistent with having no European farmer ancestry. Thus, Moldovan Yamna (like those from Bulgaria) included both unadmixed members of the core group as well as those that had acquired some European farmer ancestry.

| Test | P-value | CoreYamna | YUN_CA | S.E. |
|---|---|---|---|---|
| I10174 | 0.222 | 96.0% | 4.0% | 4.1% |
| I10206 | 0.600 | 83.7% | 16.3% | 2.9% |
| I10414 | 0.314 | 95.2% | 4.8% | 2.1% |
| I10415 | 0.605 | 97.4% | 2.6% | 4.6% |
| I10424 | 0.184 | 96.6% | 3.4% | 2.1% |
| I12507 | 0.358 | 95.5% | 4.5% | 2.2% |
| I12512 | 0.242 | 85.8% | 14.2% | 4.2% |
| I17743 | 0.593 | 93.1% | 6.9% | 2.4% |
| I17744 | 0.410 | 95.5% | 4.5% | 2.4% |
| I17745 | 0.228 | 88.9% | 11.1% | 2.5% |
| I20068 | 0.185 | 96.7% | 3.3% | 2.4% |
| I7848 | 0.015 | 93.0% | 7.0% | 2.1% |

**Table S 8 Admixture proportions for Moldova_EBA_Yamna.** Individuals 10208, I10398, I12637, I7847 are part of the Core Yamna group and used as a source here.

**Moldova_EBA_Yamna_o (2865-2576 calBCE)**

We analyze this outlier individual (I17747; 2865-2576 calBCE) separately as it has much farmer ancestry (Table S 3). The results of the model tournament (Table S 9) show that models with Core Yamna and European farmer (Trypillian or Globular Amphora) ancestry work best for this individual.

| | BPgroup+GlobularAmphora | BPgroup+Trypillia | CoreYamna+GlobularAmphora | CoreYamna+Trypillia | CoreYamna+YUN_CA | GlobularAmphora+PVgroup | PVgroup+Trypillia |
|---|---|---|---|---|---|---|---|
| BPgroup+GlobularAmphora | | 1.62E-02 | 2.34E-01 | 1.17E-01 | 1.12E-02 | 7.14E-01 | 7.89E-03 |
| BPgroup+Trypillia | 6.13E-01 | | 2.09E-01 | 5.32E-01 | 3.26E-01 | 7.86E-01 | 8.36E-02 |
| CoreYamna+GlobularAmphora | 3.20E-05 | 9.63E-06 | | 1.14E-01 | 7.78E-03 | 1.25E-05 | 1.06E-06 |
| CoreYamna+Trypillia | 4.12E-05 | 7.68E-05 | 2.88E-01 | | 2.71E-01 | 2.47E-05 | 3.49E-06 |
| CoreYamna+YUN_CA | 8.82E-05 | 1.59E-04 | 3.33E-01 | 4.92E-01 | | 3.06E-05 | 6.53E-06 |
| GlobularAmphora+PVgroup | 4.47E-01 | 1.77E-02 | 2.24E-01 | 1.34E-01 | 1.52E-02 | | 1.61E-02 |
| PVgroup+Trypillia | 5.75E-01 | 2.97E-01 | 1.95E-01 | 5.06E-01 | 3.29E-01 | 9.01E-01 | |

**Table S 9 Model competition for Moldova_EBA_Yamna_o.** Each column shows the p-value of a model when alternative models (rows) are placed on the Right set.

**Moldova_GlobularAmphora_Yamna (2906-2702 calBCE)**

This individual (I20076; 2906-2702 calBCE) is also analyzed separately. In the model competition (Table S 10) it is also inferred to be a mixture of core Yamna and European farmers (Trypillians or Globular Amphora).

| | BPgroup+GlobularAmphora | CoreYamna+GlobularAmphora | CoreYamna+Trypillia | GK2+Remontnoye | Lebyazhinka_HG+Maykop | Remontnoye+Ukraine_N |
|---|---|---|---|---|---|---|
| BPgroup+GlobularAmphora | | 1.59E-01 | 5.63E-02 | 4.67E-01 | 3.04E-03 | 4.05E-01 |
| CoreYamna+GlobularAmphora | 1.13E-13 | | 9.07E-02 | 1.31E-17 | 3.07E-28 | 1.06E-25 |
| CoreYamna+Trypillia | 1.85E-13 | 1.06E-01 | | 9.22E-19 | 1.76E-29 | 4.99E-28 |
| GK2+Remontnoye | 1.56E-01 | 3.88E-01 | 1.70E-01 | | 8.23E-03 | 4.49E-01 |
| Lebyazhinka_HG+Maykop | 1.05E-01 | 3.09E-01 | 1.27E-01 | 5.90E-01 | | 4.45E-01 |
| Remontnoye+Ukraine_N | 9.46E-03 | 2.28E-01 | 7.33E-02 | 3.40E-01 | 3.63E-03 | |

**Table S 10 Model competition for Moldova_GlobularAmphora_Yamna.** Each column shows the p-value of a model when alternative models (rows) are placed on the Right set.

**Ukraine_EBA_Yamna (3300-2500 BCE)**

For the Ukraine Yamna we see that the Remontnoye+GK2 and BPgroup+Globular Amphora models fail in competition that include Core Yamna as a source (Table S 11). The other models remain viable in competition to each other.

| | BPgroup+GlobularAmphora | CoreYamna+GlobularAmphora | CoreYamna+Trypillia | CoreYamna+YUN_CA | GK2+Remontnoye |
|---|---|---|---|---|---|
| BPgroup+GlobularAmphora | | 4.32E-01 | 2.90E-01 | 1.28E-01 | 4.30E-02 |
| CoreYamna+GlobularAmphora | 4.61E-27 | | 5.24E-01 | 1.81E-01 | 5.25E-41 |
| CoreYamna+Trypillia | 3.13E-27 | 7.59E-01 | | 2.34E-01 | 4.28E-39 |
| CoreYamna+YUN_CA | 4.13E-27 | 8.65E-01 | 7.53E-01 | | 8.51E-39 |
| GK2+Remontnoye | 4.11E-01 | 8.13E-01 | 7.45E-01 | 3.97E-01 | |

**Table S 11 Model competition for Ukraine_EBA_Yamna.** Each column shows the p-value of a model when alternative models (rows) are placed on the Right set.

We must note here that four Yamna individuals are included in the "Core Yamna" group definition: I12168, I20975, I2105, I3141_enhanced. Thus, in Ukraine too there were indeed representatives of the Core Yamna group without evidence of any farmer admixture, as well as individuals as those included under the Ukraine_EBA_Yamna genetic grouping (I12619, I12843, I11999, I12617) with some farmer admixture. Thus, the Yamna of Ukraine (in whose territory co-existed herder and settled farmer populations) was indeed home to Yamna of the core group (whose ancestry was from the Serednii Stih and consisted of the mix of eastern CLV migrants and Ukraine_N-related foragers) as well as Yamna who had acquired a fraction of European farmer ancestry. We fit the same model for the Ukraine_EBA_Yamna individuals that we did for Bulgarian ones (Table S 5) and show the results in Table S 12 (We also fit the model with the geographically more proximal Trypillia as a source). As in Bulgaria the Yamna of Ukraine included individuals of both the core group and those admixed with European farmers; the Yamna advance must have been rapid enough to ensure that at least individuals from the core group reached Bulgaria without any admixture from the farmer groups in the intervening space, but clearly some Yamna in both Ukraine (closer to the unknown origin of the core Yamna group) and Bulgaria had acquired some European farmer ancestry.

| Test | P-value | CoreYamna | YUN_CA | S.E. |
|---|---|---|---|---|
| I11999 | 0.026 | 95.9% | 4.1% | 2.1% |
| I12617 | 0.270 | 93.4% | 6.6% | 3.8% |
| I12619 | 0.207 | 92.6% | 7.4% | 2.3% |
| I12843 | 0.090 | 91.8% | 8.2% | 2.1% |
| **Test** | **P-value** | **CoreYamna** | **Trypillia** | **S.E.** |
| I11999 | 0.062 | 93.9% | 6.1% | 2.6% |
| I12617 | 0.188 | 93.6% | 6.4% | 4.7% |
| I12619 | 0.443 | 90.3% | 9.7% | 2.6% |
| I12843 | 0.038 | 91.1% | 8.9% | 2.6% |

**Table S 12 Admixture of Ukrainian Yamna individuals.** Individuals included in the Core Yamna group (I12168, I20975, I2105, I3141_enhanced) used as a source are not shown.

**Ukraine_MBA_Catacomb_o1 (2201-2032 calBCE)**

This individual (I13071; 2201-2032 calBCE) fits multiple models (Table S 3) and cannot be well resolved using the model tournament approach. In terms of the models with Core Yamna and European farmer ancestry that seem to describe well Bronze Age populations from the NPR, it seems to have ~1/3 of farmer ancestry.

**Ukraine_MBA_MultiCordonedWare_Babine (2400-1900BCE)**

The only feasible models for the Multi-Cordoned Ware/Babine population (Table S 13) involve Core Yamna and 9.1±3.6% Ukraine_N or 6.3±2.3% Serbia Iron Gates hunter-gatherer ancestry (Table S 3)

.

| | CoreYamna+Serbia_IronGates_Mesolithic | CoreYamna+Ukraine_N | GK2+Remontnoye |
|---|---|---|---|
| CoreYamna+Serbia_IronGates_Mesolithic | | 6.59E-02 | 1.04E-32 |
| CoreYamna+Ukraine_N | 2.04E-01 | | 9.49E-37 |
| GK2+Remontnoye | 1.90E-01 | 7.24E-02 | |

**Table S 13 Model competition for Ukraine_MBA_MultiCordonedWare_Babine.** Each column shows the p-value of a model when alternative models (rows) are placed on the Right set.

Note that for Multi-Cordoned Ware the model with Trypillian as a source (that worked for the Ukraine_EBA_Yamna) fails (p=0.027) and conversely for the Ukraine_EBA_Yamna the model with Serbia Iron Gates as a source fails (p=0.005).

We suspected that this is because in the Multi-Cordoned Ware the non-Yamna source is enriched in hunter-gatherer ancestry. This can be assessed directly by examining 3-way models with both European farmers and Ukraine_N/Serbian Iron Gates as sources (Table S 14) which confirms our intuition.

| | | | | | Proportions | | | Std. errors | | |
|---|---|---|---|---|---|---|---|---|---|---|
| **Test** | **A** | **B** | **C** | **P-value** | **A** | **B** | **C** | **A** | **B** | **C** |
| Ukraine_EBA_Yamna | CoreYamna | GlobularAmphora | Serbia_IronGates_Mesolithic | 0.783 | 91.3% | 7.8% | 0.9% | 1.7% | 1.7% | 1.5% |
| Ukraine_EBA_Yamna | CoreYamna | GlobularAmphora | Ukraine_N | 0.761 | 91.3% | 8.1% | 0.7% | 2.3% | 1.6% | 2.3% |
| Ukraine_EBA_Yamna | CoreYamna | Serbia_IronGates_Mesolithic | Trypillia | 0.840 | 90.5% | 2.0% | 7.6% | 1.8% | 1.4% | 1.6% |
| Ukraine_EBA_Yamna | CoreYamna | Serbia_IronGates_Mesolithic | YUN_CA | 0.776 | 90.9% | 3.0% | 6.1% | 1.9% | 1.4% | 1.4% |
| Ukraine_EBA_Yamna | CoreYamna | Trypillia | Ukraine_N | 0.740 | 89.7% | 7.9% | 2.4% | 2.5% | 1.6% | 2.2% |
| Ukraine_EBA_Yamna | CoreYamna | Ukraine_N | YUN_CA | 0.580 | 89.5% | 4.0% | 6.5% | 2.6% | 2.2% | 1.3% |
| Ukraine_MBA_MultiCordonedWare_Babine | CoreYamna | GlobularAmphora | Serbia_IronGates_Mesolithic | 0.148 | 91.6% | 3.0% | 5.3% | 2.7% | 2.5% | 2.4% |
| Ukraine_MBA_MultiCordonedWare_Babine | CoreYamna | GlobularAmphora | Ukraine_N | 0.107 | 88.4% | 4.1% | 7.5% | 3.7% | 2.3% | 3.7% |
| Ukraine_MBA_MultiCordonedWare_Babine | CoreYamna | Serbia_IronGates_Mesolithic | Trypillia | 0.142 | 91.1% | 5.8% | 3.0% | 2.8% | 2.2% | 2.5% |
| Ukraine_MBA_MultiCordonedWare_Babine | CoreYamna | Serbia_IronGates_Mesolithic | YUN_CA | 0.144 | 91.4% | 6.2% | 2.5% | 2.7% | 2.2% | 2.0% |
| Ukraine_MBA_MultiCordonedWare_Babine | CoreYamna | Trypillia | Ukraine_N | 0.094 | 87.7% | 4.0% | 8.3% | 3.9% | 2.3% | 3.5% |
| Ukraine_MBA_MultiCordonedWare_Babine | CoreYamna | Ukraine_N | YUN_CA | 0.090 | 87.6% | 9.1% | 3.2% | 4.0% | 3.5% | 1.9% |

**Table S 14 3-way models with varying hunter-gatherer ancestry for Bronze Age Ukraine.**

Thus, both the Yamna of Ukraine of the EBA and the later Multi-Cordoned Ware of the MBA were largely descended from the Core Yamna but the latter experienced gene flow from a population that seems to have had more hunter-gatherer ancestry.

It was previously observed that Bronze Age populations from neighboring Romania included those that had a high proportion of hunter-gatherer ancestry:[12]

*"The first cluster is made up mainly of individuals from Arman (Cârlomănești) who have very low amounts of CHG/EHG ancestry but substantial (~24%) Balkan hunter-gatherer ancestry as does an individual from Târgșoru Vechi (I7152) in the mid-3rd millennium BCE and an individual from Ploieşti Triaj (I10494/ROU_Ploieşti_B) at ~30%."*

These individuals, together with those of Ukraine both point to the existence of Chalcolithic populations of high hunter-gatherer ancestry that bequeathed it to those of the Bronze Age. We know of at least one such Chalcolithic individual from Gura Baciului of >50% hunter-gatherer ancestry,[12,34] which provides direct evidence for the existence of such populations during the Chalcolithic.

##### Ukraine_N_I27992 (5363-5216 calBCE)

We do not draw any conclusions from this Neolithic individual (I27992; 5363-5216 calBCE) as it has a high contamination rate mtcontam=[0.717,0.942]. It is not cladal with Ukrainian Neolithic hunter-gatherers but this would be a consequence of contamination and thus not evidence for early contact with the Ukrainian Neolithic population.

##### Usatove (4400-3600 BCE)

The Usatove population label includes individuals from Mayaky dated to the late 5$^{th}$ and early 4$^{th}$ millennia BCE. It can be modeled uniquely as a mixture of ~45% PVgroup and ~55% Trypillians (Table S 3). The next best model (however, p=0.002) involves ~56% Remontnoye ancestry and ~44% Trypillian ancestry. The model with BPgroup+Trypillians fails clearly (p<1e-4). We were curious about this as BPgroup and PVgroup are closely related populations within the North Caucasus-Lower Volga Eneolithic. We examined the 3-way model with BPgroup+Armenia_Aknashen_N ancestry substituting PVgroup, as PVgroup belongs on the CLV cline having some southern Caucasus Neolithic ancestry along that cline. Indeed, this model fits (p=0.393) with an estimated 14.4±3.1% Aknashen ancestry.

We also confirmed that the PVgroup+Trypillian model fits the population MAJ of ref.[22] a different Eneolithic sample set from Mayaky (p=0.231) as well as the USV Usatove-related population (p=0.083). The authors of ref.[22] observe that:

"*By contrast, MAJ and USV can be modelled as VAR_CA or Ukraine Trypillia (around 50%), Steppe Eneolithic (around 35%) and Caucasus Eneolithic/Maykop (around 15%) as minor third component.*"

Our results agree with this general conclusion.

The Usatove population at Mayaky is then modeled as a fairly even mix of CLV Cline migrants (as those of PVgroup, or alternatively a mix of BPgroup and Aknashen) with Trypillian farmers. We note that neither YUN_CA nor Globular Amphora work for the Usatove (p<1e-4).

An interpretation of this modeling is that Usatove is formed by westward migrations of CLV Cline people Iinto Ukraine. This migration is also evidenced in sites of the Serednii Stih culture which were formed on the basis of migrations of CLV Cline people admixing with Ukraine_N-related hunter-gatherers of the Dnipro-Don region. Thus, the CLV cline migrations westward admixed with both the hunter-gatherers of the Dnipro-Don region (creating the Serednii Stih populations along the "Dnipro Cline") and the Trypillian farmers to the west (creating the Usatove). Their different substratum (hunter-gatherer vs. farmer) is what separates Serednii Stih from Usatove. What unifies them is their common ancestry from the CLV cline.

**Usatove_I20078 (3340-3034 calBCE)**

This individual from Taraclia (3340-3034 calBCE) is chronologically later and an outlier with respect to the Mayaky individuals and is analyzed separately. Model competition (Table S 15) shows that only the model that involves Core Yamna and TTK ancestry remains feasible. TTK-related ancestry entered the ancestry of the Core Yamna via its CLV-cline component as such ancestry contributed to Lower Volga-North Caucasus Eneolithic populations (like BPgroup and PVgroup which also appear as sources for this individual). However, the fact that the Core Yamna+TTK model outcompetes the BPgroup/PVgroup+TTK models suggests that the individual shares common history with the Core Yamna and is not from a population derived independently from the CLV Cline with extra TTK-related ancestry.

| | BPgroup+TTK | CoreYamna+TTK | PVgroup+TTK |
|---|---|---|---|
| BPgroup+TTK | | 2.87E-01 | 1.81E-03 |
| CoreYamna+TTK | 2.10E-02 | | 1.15E-01 |
| PVgroup+TTK | 1.28E-03 | 2.83E-01 | |

**Table S 15 Model competition for Usatove_I20078.** Each column shows the p-value of a model when alternative models (rows) are placed on the Right set.

We were curious if the TTK-related ancestry in this individual could be from a source like the "Steppe Maykop"[13] which is modeled in the linked manuscript ref.[3] as having a substantial ancestry from a TTK-related "Central Asian/Siberian" source. Thus, we fit the Core Yamna+Russia_Steppe_Maykop to this individual (p=0.345) with a predicted 60.6±6.2% Steppe Maykop ancestry. The model still fits (p=0.432) in competition with the CoreYamna+TTK model while conversely the CoreYamna+TTK model fails (p<1e-4) when Steppe Maykop is added to the Right set. As the Steppe Maykop is both temporally and geographically closer to our individual of interest and wins against it, we can conclude that this individual is of likely mixed Yamna-Steppe Maykop origins.

**Zhivotilovka_I17974**

This individual cannot be well-resolved using the model tournament as several models remain feasible (Table S 16). The CoreYamna+TTK model (that can be fit successfully for the I20078 individual previously discussed) has the highest p-value (0.616) and low standard error (3.4%) and it is reasonable that alternative models cannot be rejected for this individual because of the much lower amount of TTK-related ancestry (~11% vs. ~35% for the I20078 individual). We also tested the CoreYamna+Steppe Maykop model (p=0.324) with an estimated 18.2±6.0% Steppe Maykop ancestry. A parsimonious interpretation of the data is that this individual (like I20078) was formed of the same Yamna+Steppe Maykop admixture process but with about ~1/3 of the Steppe Maykop ancestry found in I20078.

| | CoreYamna+Lebyazhinka_HG | CoreYamna+TTK | GK2+PVgroup | Lebyazhinka_HG+PVgroup | PVgroup+Russia_Karelia | PVgroup+Serbia_IronGates_Mesolithic | PVgroup+Ukraine_N |
|---|---|---|---|---|---|---|---|
| CoreYamna+Lebyazhinka_HG | | 6.50E-01 | 6.99E-01 | 1.52E-01 | 1.59E-01 | 1.87E-01 | 6.40E-01 |
| CoreYamna+TTK | 3.09E-02 | | 3.49E-01 | 1.08E-01 | 1.24E-01 | 5.15E-02 | 2.77E-01 |
| GK2+PVgroup | 2.56E-02 | 4.22E-01 | | 7.01E-02 | 6.35E-02 | 7.92E-02 | 1.80E-01 |
| Lebyazhinka_HG+PVgroup | 7.23E-02 | 5.11E-01 | 6.18E-01 | | 1.24E-01 | 1.18E-01 | 5.02E-01 |
| PVgroup+Russia_Karelia | 9.14E-02 | 6.11E-01 | 5.77E-01 | 1.42E-01 | | 8.70E-02 | 5.28E-01 |
| PVgroup+Serbia_IronGates_Mesolithic | 1.17E-02 | 5.99E-01 | 3.69E-01 | 2.35E-01 | 2.04E-01 | | 1.76E-01 |
| PVgroup+Ukraine_N | 7.51E-02 | 5.89E-01 | 4.67E-01 | 2.45E-01 | 1.95E-01 | 1.25E-01 | |

**Table S 16 Model competition for Zhivotilovka_I17974.** Each column shows the p-value of a model when alternative models (rows) are placed on the Right set.

**Populations that cannot be well-modeled**

Three populations cannot be well-modeled according to our feasibility criteria: GlobularAmphora, Ukraine_EBA_Catacomb_possible, Zhivotilovka_I17973 even if we allow for *N*=3 sources. We examine them here separately.

**GlobularAmphora (3400-2600 BCE)**

If we relax our p-value threshold to 0.001, then the best 2-way model (p=0.004) for the Globular Amphora population involves 75.7±0.9% YUN_CA and 24.9±0.9% Serbia_IronGates_Mesolithic ancestry which point to this population having a higher proportion of hunter-gatherer ancestry than the Chalcolithic population from Yunatsite in Bulgaria. The obtained proportions are similar to the analysis of a subset of Globular Amphora individuals studied in ref.[16] Due to the relative homogeneity of both European farmers (largely descended from their Anatolian predecessors[11,35]) and European hunter-gatherers (largely descended from the WHG[36] or "Villabruna[7]/Oberkassel[20] cluster") alternative models could be feasible and the set of sources used may not be ideal for modeling the Globular Amphora population.

We were curious about the bad fit of the 2-way model and observed that it overestimates (Z=1.9-2.3) shared genetic drift with the two highland West Asian[37] population outgroups (CHG and Ganj Dareh Neolithic). As this ancestry differentiated the Neolithic people of the South Balkans from those of central-northern Europe[12,16,38] we tried an alternative model with LBK (Linearbandkeramik) as the farmer source. This is also a priori more plausible given that the Globular Amphora emerged in areas of central-eastern Europe previously occupied by LBK farmers. Indeed, this model fits well (p=0.319) with an estimated 24.7±0.9% LBK ancestry.

**Trypillians (4700-2700 BCE)**

The Trypillian population is heterogeneous in PCA and cannot be modeled well with any 1, 2, 3 sources. We wanted to investigate whether the more notable PCA outlier within this population (I20069; 3323-2935 calBCE from Dănceni) might have different ancestry from the population as a whole.

When we model I20069 separately, we do find that it has a significant amount of CLV ancestry (Table S 17) and is composed of ~1/4 PVgroup and ~3/4 Trypillia ancestry (using the remaining 27 Trypillians as a source).

| A | B | P-value | A | B | S.E. |
|---|---|---|---|---|---|
| BPgroup | Trypillia | 0.055 | 22.7% | 77.3% | 2.2% |
| PVgroup | Trypillia | 0.122 | 24.3% | 75.7% | 2.4% |
| Remontnoye | Trypillia | 0.064 | 30.2% | 69.8% | 3.1% |

**Table S 17 Modeling Trypillia_I20069.**

We can also model (p=0.161) the individual as a mixture of 53.7±5.5% Usatove and Trypillia (as Usatove itself is a mixture of PVgroup and Trypillia).

Nonetheless, even after excluding the individual we cannot model Trypillians (p<1e-5 even for *N*=3 models) successfully. To understand the latent structure within this population we turned to modeling each of the 28 individuals under the "Trypillia" label separately in terms of the same as sources as we used in the rest of this note (except, of course, Trypillia itself). By doing so we wanted to identify models that could be used to describe each individual separately and disclose any peculiarities of particular Trypillian individuals.

24 of 28 individuals can be modeled in our framework. The four exceptions (to which we will return below) are: VERT105B_wNonUDG.SG, VERT015_wNonUDG.SG, VERT035_wNonUDG.SG, VERT111_wNonUDG.SG from Verteba Cave.[21]

A single model (with BPgroup, YUN_CA, and Serbia Iron Gates Mesolithic ancestry) is feasible for 23 of the 24 remaining individuals. For the remaining individual (VERT103B_wNonUDG.SG) there are other feasible models. Three other models are qualitatively similar to the identified model and are feasible for 22/24 individuals; they differ from it in replacing BPgroup with PVgroup, Remontnoye, or Core Yamna. The unifying element of all these models is that they include some CLV ancestry although it is unclear which part of the CLV cline it is derived froTm (BPgroup/PVgroup/Remontnoye) or if it also includes some Dnipro-Don hunter-gatherer ancestry (as Core Yamna does).

We show the fit of the top model for all 28 individuals (Extended Data Table 3). We see that for many individuals there is no significant amount of BPgroup-related ancestry, but this kind of ancestry is not present only in the PCA outlier (I20069) but also in several other individuals from the population.

Of the five individuals that do not fit this model, three fit it at the p=0.01 level. We examined qpAdm output for all five individuals to identify the Right populations that cause the poor fit. Z-scores are between -2.4 and +2.2. For three individuals (VERT105B_wNonUDG.SG, VERT015_wNonUDG.SG, and VERT111_wNonUDG.SG) shared genetic drift with CHG is underestimated (Z<-2). For VERT103B_wNonUDG.SG, shared genetic drift with Ganj Dareh is overestimated (Z>2).

In conclusion, we think that Trypillians were mainly formed on the basis of the European farmer cline (represented by YUN_CA and Iron Gates hunter-gatherers) that received admixture from people that had CLV ancestry. We have seen above that Usatove was formed on the basis of CLV people absorbing Trypillian ancestry. We can synthesize the evidence from Usatove and Trypillians as follows: the movement of people with CLV ancestry into the NPR brought them into contact with European farmer populations; in Usatove groups ancestry from the two groups was fairly balanced and in Trypillian groups farmer ancestry predominated but some gene flow occurred from the CLV migrants as well. Individuals like I20069, a fairly even mix of "Usatove" and "Trypillian" groups, bridge the gap between those two differentiated populations.

**Ukraine_EBA_Catacomb_possible (2874-2630 calBCE)**

Individual Ukraine_EBA_Catacomb_possible (I11850; 2874-2630 calBCE) from the Odesa region cannot be modeled at the p=0.05 threshold but several models exist for it at the p=0.01 threshold (Table S 18), however no single model emerges as a winner in the model tournament (Table S 19). All successful models include some CLV Cline ancestry.

| A | B | P-value | A | B | S.E. |
|---|---|---|---|---|---|
| Armenia_Aknashen_N | CoreYamna | 0.013 | 31.6% | 68.4% | 7.4% |
| Armenia_Aknashen_N | Lebyazhinka_HG | 0.012 | 64.5% | 35.5% | 4.3% |
| Armenia_Aknashen_N | Russia_Karelia | 0.013 | 65.5% | 34.5% | 4.0% |
| BPgroup | Trypillia | 0.013 | 69.6% | 30.4% | 4.3% |
| BPgroup | YUN_CA | 0.016 | 73.7% | 26.3% | 3.8% |
| CoreYamna | Maykop | 0.011 | 67.0% | 33.0% | 8.2% |
| GK2 | Remontnoye | 0.016 | 12.8% | 87.2% | 6.6% |
| GlobularAmphora | PVgroup | 0.010 | 26.6% | 73.4% | 4.7% |
| PVgroup | Trypillia | 0.036 | 72.3% | 27.7% | 4.8% |
| PVgroup | YUN_CA | 0.027 | 76.4% | 23.6% | 4.1% |
| Remontnoye | Russia_Karelia | 0.011 | 91.1% | 8.9% | 5.3% |
| Remontnoye | Serbia_IronGates_Mesolithic | 0.013 | 93.2% | 6.8% | 4.0% |
| Remontnoye | Ukraine_N | 0.016 | 89.3% | 10.7% | 5.8% |

**Table S 18 Feasible models at p=0.01 level for Ukraine_EBA_Catacomb_possible.**

| | Armenia_Aknashen_N+CoreYamna | BPgroup+Trypillia | BPgroup+YUN_CA | CoreYamna+Maykop | GK2+Remontnoye | Armenia_Aknashen_N+Lebyazhinka_HG | GlobularAmphora+PVgroup | PVgroup+Trypillia | PVgroup+YUN_CA | Remontnoye+Russia_Karelia | Remontnoye+Serbia_IronGates_Mesolithic | Remontnoye+Ukraine_N | Armenia_Aknashen_N+Russia_Karelia |
|---|---|---|---|---|---|---|---|---|---|---|---|---|---|
| Armenia_Aknashen_N+CoreYamna | | 1.86E-05 | 2.70E-05 | 1.50E-02 | 2.86E-07 | 5.58E-15 | 3.49E-06 | 1.11E-05 | 1.17E-05 | 1.76E-07 | 5.56E-10 | 3.91E-09 | 4.33E-17 |
| BPgroup+Trypillia | 2.56E-03 | | 2.35E-02 | 2.34E-03 | 6.72E-03 | 4.57E-07 | 1.89E-02 | 4.76E-02 | 4.43E-02 | 4.42E-03 | 1.67E-03 | 2.76E-03 | 4.90E-07 |
| BPgroup+YUN_CA | 6.90E-04 | 8.20E-03 | | 4.48E-04 | 1.69E-03 | 3.39E-07 | 6.79E-03 | 3.23E-02 | 3.84E-02 | 1.19E-03 | 5.89E-04 | 8.29E-04 | 2.36E-07 |
| CoreYamna+Maykop | 1.35E-02 | 9.96E-06 | 1.08E-05 | | 4.14E-08 | 1.44E-14 | 2.51E-07 | 9.55E-07 | 7.39E-07 | 2.12E-08 | 7.26E-12 | 2.25E-10 | 5.67E-17 |
| GK2+Remontnoye | 3.09E-03 | 4.50E-03 | 6.43E-03 | 2.12E-03 | | 8.84E-05 | 2.70E-03 | 6.27E-03 | 5.72E-03 | 5.64E-03 | 3.21E-03 | 9.53E-03 | 1.47E-04 |
| Armenia_Aknashen_N+Lebyazhinka_HG | 2.70E-03 | 6.53E-03 | 6.54E-03 | 3.41E-03 | 5.28E-03 | | 2.39E-03 | 9.00E-03 | 5.85E-03 | 2.37E-03 | 1.59E-03 | 2.98E-03 | 1.54E-03 |
| GlobularAmphora+PVgroup | 2.88E-02 | 3.74E-02 | 3.44E-02 | 2.47E-02 | 4.27E-02 | 5.96E-03 | | 5.23E-02 | 3.72E-02 | 2.79E-02 | 3.13E-02 | 3.97E-02 | 5.63E-03 |
| PVgroup+Trypillia | 4.10E-02 | 2.50E-02 | 3.65E-02 | 2.36E-02 | 4.18E-02 | 4.68E-03 | 1.43E-02 | | 3.24E-02 | 2.33E-02 | 3.22E-02 | 3.95E-02 | 5.36E-03 |
| PVgroup+YUN_CA | 1.61E-02 | 1.81E-02 | 2.57E-02 | 7.47E-03 | 1.23E-02 | 3.49E-03 | 5.12E-03 | 2.49E-02 | | 8.12E-03 | 1.12E-02 | 1.32E-02 | 3.36E-03 |
| Remontnoye+Russia_Karelia | 5.47E-04 | 1.93E-03 | 3.02E-03 | 5.48E-04 | 7.59E-03 | 2.74E-05 | 5.16E-04 | 1.03E-03 | 9.44E-04 | | 5.15E-05 | 1.44E-03 | 1.61E-04 |
| Remontnoye+Serbia_IronGates_Mesolithic | 4.09E-03 | 5.37E-03 | 3.27E-03 | 2.53E-03 | 2.10E-02 | 1.08E-04 | 2.85E-03 | 5.11E-03 | 1.31E-03 | 2.02E-03 | | 2.29E-02 | 1.17E-04 |
| Remontnoye+Ukraine_N | 7.30E-04 | 1.85E-03 | 1.60E-03 | 7.39E-04 | 1.23E-02 | 1.00E-05 | 7.84E-04 | 1.30E-03 | 6.68E-04 | 1.80E-03 | 1.90E-03 | | 3.57E-05 |
| Armenia_Aknashen_N+Russia_Karelia | 1.02E-03 | 7.60E-03 | 8.34E-03 | 2.00E-03 | 1.00E-02 | 1.17E-03 | 1.56E-03 | 3.38E-03 | 2.35E-03 | 1.44E-02 | 9.01E-05 | 2.11E-03 | |

**Table S 19 Model competition for Ukraine_EBA_Catacomb_possible.** Each column shows the p-value of a model when alternative models (rows) are placed on the Right set.

**Zhivotilovka_I17973 (3354-3103 calBC)**

The second Zhivotilovka individual cannot be modeled in the same way as the first (I17974) as the Core Yamna+TTK model fails (p=6e-41). However, several models with mostly Aknashen Neolithic as one source and different European or steppe populations fit this population at the p=0.01 threshold (Table S 20).

| A | B | P-value | A | B | S.E. |
|---|---|---|---|---|---|
| Armenia_Aknashen_N | BPgroup | 0.012 | 91.2% | 8.8% | 4.1% |
| Armenia_Aknashen_N | CoreYamna | 0.012 | 90.5% | 9.5% | 4.7% |
| Armenia_Aknashen_N | GK2 | 0.010 | 94.5% | 5.5% | 2.8% |
| Armenia_Aknashen_N | Lebyazhinka_HG | 0.011 | 95.0% | 5.0% | 2.4% |
| Armenia_Aknashen_N | Remontnoye | 0.011 | 85.1% | 14.9% | 8.1% |
| Armenia_Aknashen_N | Russia_Karelia | 0.013 | 95.3% | 4.7% | 2.2% |
| Armenia_Aknashen_N | Serbia_IronGates_Mesolithic | 0.011 | 96.0% | 4.0% | 2.1% |
| Armenia_Aknashen_N | Ukraine_N | 0.012 | 94.6% | 5.4% | 2.5% |

**Table S 20 Feasible models at p=0.01 level for Zhivotilovka_I17973**

The admixture proportions of this individual point to a population of the Caucasus Neolithic (like Aknashen) with a little steppe ancestry. The Maykop population is a similar blend (it can be modeled as 86.2±2.9% Aknashen and 13.8±2.9% BPgroup as discussed in ref.[3]), however it does not suffice as a source for this individual (p=0.0025). Neither does Aknashen Neolithic by itself (p=0.0047). All other single sources fail convincingly (p<1e-32). Examining the qpAdm output of 2-way models we observed that shared genetic drift with all the outgroups was positive which suggests that the individual might have ancestry from a source that has genetic drift not shared with any of the outgroups. Provisionally, we can say that this individual has ancestry from the Caucasus based on its position in PCA and the Ffact that Maykop is its closest (albeit imperfect) single source: the precise origin of this individual remains unclear.

**Re-analysis of Early Bronze Age outliers from Ukraine from Mathieson et al. (2018) (ref.[16])**

We discuss separately four previously published individuals from ref.[16] for two of which we report additional data: Ukraine_EBA_Deriivka_I4110, Ukraine_EBA_Deriivka_I5882, Ukraine_EBA_Deriivka_I5884, Ukraine_EBA_Ozera_I1917.

Individual I1917 (Ozera; 3096-2913 calBCE) was described as having some Anatolian Neolithic-related ancestry in its original publication.[16] It can be modeled as a 2-way mixture in our framework of more proximate sources (Table S 21) all of which do indeed possess such Anatolian Neolithic-related ancestry.

Model competition identifies a single feasible model for this individual that involves half Core Yamna and half Maykop ancestry (Table S 22). This may have come about either by the migration of Yamna into the North Caucasus and its admixture with Maykop and successor populations followed by the individual's migration to Ukraine or, alternatively, the admixture may have taken place in Ukraine itself.

| A | B | P-value | A | B | S.E. |
|---|---|---|---|---|---|
| Armenia_Aknashen_N | CoreYamna | 0.089 | 41.7% | 58.3% | 4.1% |
| CoreYamna | Maykop | 0.345 | 50.1% | 49.9% | 4.5% |
| GK2 | Maykop | 0.118 | 26.7% | 73.3% | 2.6% |
| GlobularAmphora | Remontnoye | 0.101 | 12.3% | 87.7% | 3.4% |
| Lebyazhinka_HG | Maykop | 0.127 | 23.9% | 76.1% | 2.2% |
| Remontnoye | Serbia_IronGates_Mesolithic | 0.083 | 91.8% | 8.2% | 2.2% |

**Table S 21 Feasible 2-way models for Ukraine_EBA_Ozera_I1917.**

| | Armenia_Aknashen_N+CoreYamna | CoreYamna+Maykop | GK2+Maykop | GlobularAmphora+Remontnoye | Lebyazhinka_HG+Maykop | Remontnoye+Serbia_IronGates_Mesolithic |
|---|---|---|---|---|---|---|
| Armenia_Aknashen_N+CoreYamna | | 4.25E-01 | 7.28E-13 | 2.13E-03 | 4.84E-13 | 2.12E-02 |
| CoreYamna+Maykop | 4.18E-03 | | 1.76E-13 | 1.73E-04 | 1.33E-13 | 1.71E-03 |
| GK2+Maykop | 1.02E-02 | 4.14E-01 | | 4.30E-03 | 1.58E-01 | 8.49E-03 |
| GlobularAmphora+Remontnoye | 1.57E-01 | 4.18E-01 | 1.76E-02 | | 1.58E-02 | 9.29E-02 |
| Lebyazhinka_HG+Maykop | 6.36E-04 | 5.39E-02 | 6.21E-02 | 1.29E-03 | | 1.34E-03 |
| Remontnoye+Serbia_IronGates_Mesolithic | 1.43E-01 | 2.55E-01 | 1.59E-02 | 2.60E-02 | 4.08E-03 | |

**Table S 22 Model competition for Ukraine_EBA_Ozera_I1917.** Each column shows the p-value of a model when alternative models (rows) are placed on the Right set.

The other three individuals of interest (all from Deriivka) cannot be modeled as 2-way mixtures and were described as having steppe, hunter-gatherer, and Anatolian Neolithic ancestry in ref.[16]. We identified six 3-way models that are applicable to all three individuals (Table S 23) which indeed combine Anatolian Neolithic-related (via either YUN_CA, Trypillian, or Globular Amphora European populations) ancestry, hunter-gatherer ancestry (represented by Iron Gates hunter-gatherers) and steppe ancestry (either North Caucasus-Lower Volga Eneolithic BPgroup/PVgroup or Core Yamna).

Model competition for each of the three individuals (Table S 24; Table S 25; Table S 26) identifies a single model (with Core Yamna as the steppe source and Trypillians as the farmer source). The amount of Balkan hunter-gatherer ancestry in these three individuals is substantial (~23-44%) and thus provides more evidence (to that of the Multi-Cordoned Ware / Babine) for admixture with populations of high hunter-gatherer ancestry during the Early Bronze Age. They provide additional evidence for the Bronze Age persistence of high hunter-gatherer ancestry from the Chalcolithic also seen in Romania and remarked on in our discussion of the Babine MBA populations.

| | | | | | Proportions | | | Std. errors | | |
|---|---|---|---|---|---|---|---|---|---|---|
| **Test** | **A** | **B** | **C** | **P-value** | **A** | **B** | **C** | **A** | **B** | **C** |
| Ukraine_EBA_Deriivka_I4110 | BPgroup | Serbia_IronGates_Mesolithic | Trypillia | 0.202 | 37.5% | 24.5% | 38.0% | 2.8% | 2.6% | 2.7% |
| Ukraine_EBA_Deriivka_I4110 | BPgroup | Serbia_IronGates_Mesolithic | YUN_CA | 0.113 | 39.4% | 29.5% | 31.0% | 2.7% | 2.4% | 2.1% |
| Ukraine_EBA_Deriivka_I4110 | CoreYamna | GlobularAmphora | Serbia_IronGates_Mesolithic | 0.118 | 49.2% | 33.2% | 17.6% | 3.2% | 3.2% | 2.8% |
| **Ukraine_EBA_Deriivka_I4110** | **CoreYamna** | **Serbia_IronGates_Mesolithic** | **Trypillia** | **0.216** | **45.6%** | **22.5%** | **32.0%** | **3.2%** | **2.6%** | **3.0%** |
| Ukraine_EBA_Deriivka_I4110 | CoreYamna | Serbia_IronGates_Mesolithic | YUN_CA | 0.136 | 47.6% | 26.6% | 25.9% | 3.0% | 2.4% | 2.3% |
| Ukraine_EBA_Deriivka_I4110 | PVgroup | Serbia_IronGates_Mesolithic | Trypillia | 0.054 | 38.2% | 26.3% | 35.5% | 2.8% | 2.5% | 2.7% |
| Ukraine_EBA_Deriivka_I5882 | BPgroup | Serbia_IronGates_Mesolithic | Trypillia | 0.776 | 37.1% | 42.5% | 20.5% | 2.6% | 2.5% | 2.5% |
| Ukraine_EBA_Deriivka_I5882 | BPgroup | Serbia_IronGates_Mesolithic | YUN_CA | 0.780 | 38.0% | 45.2% | 16.8% | 2.5% | 2.4% | 2.0% |
| Ukraine_EBA_Deriivka_I5882 | CoreYamna | GlobularAmphora | Serbia_IronGates_Mesolithic | 0.918 | 46.8% | 15.2% | 38.0% | 2.9% | 2.7% | 2.7% |
| **Ukraine_EBA_Deriivka_I5882** | **CoreYamna** | **Serbia_IronGates_Mesolithic** | **Trypillia** | **0.899** | **45.4%** | **40.2%** | **14.5%** | **3.0%** | **2.5%** | **2.6%** |
| Ukraine_EBA_Deriivka_I5882 | CoreYamna | Serbia_IronGates_Mesolithic | YUN_CA | 0.919 | 46.1% | 42.0% | 11.9% | 2.9% | 2.5% | 2.1% |
| Ukraine_EBA_Deriivka_I5882 | PVgroup | Serbia_IronGates_Mesolithic | Trypillia | 0.547 | 37.9% | 44.0% | 18.1% | 2.7% | 2.6% | 2.5% |
| Ukraine_EBA_Deriivka_I5884 | BPgroup | Serbia_IronGates_Mesolithic | Trypillia | 0.108 | 29.6% | 45.2% | 25.2% | 2.5% | 2.5% | 2.4% |
| Ukraine_EBA_Deriivka_I5884 | BPgroup | Serbia_IronGates_Mesolithic | YUN_CA | 0.075 | 30.8% | 48.6% | 20.6% | 2.5% | 2.3% | 2.1% |
| Ukraine_EBA_Deriivka_I5884 | CoreYamna | GlobularAmphora | Serbia_IronGates_Mesolithic | 0.095 | 38.3% | 20.9% | 40.8% | 3.0% | 2.9% | 2.7% |
| **Ukraine_EBA_Deriivka_I5884** | **CoreYamna** | **Serbia_IronGates_Mesolithic** | **Trypillia** | **0.179** | **36.0%** | **43.5%** | **20.4%** | **2.9%** | **2.4%** | **2.6%** |
| Ukraine_EBA_Deriivka_I5884 | CoreYamna | Serbia_IronGates_Mesolithic | YUN_CA | 0.142 | 37.3% | 46.2% | 16.5% | 3.0% | 2.5% | 2.1% |
| Ukraine_EBA_Deriivka_I5884 | PVgroup | Serbia_IronGates_Mesolithic | Trypillia | 0.151 | 30.4% | 46.7% | 22.8% | 2.6% | 2.4% | 2.6% |

**Table S 23 3-way models applicable to all three Ukraine_EBA outlier individuals.** The model that is feasible for all three individuals in the tournament approach is highlighted in bold.

| | BPgroup+Serbia_IronGates_Mesolithic+Trypillia | BPgroup+Serbia_IronGates_Mesolithic+YUN_CA | CoreYamna+GlobularAmphora+Serbia_IronGates_Mesolithic | CoreYamna+Serbia_IronGates_Mesolithic+Trypillia | CoreYamna+Serbia_IronGates_Mesolithic+YUN_CA | PVgroup+Serbia_IronGates_Mesolithic+Trypillia |
|---|---|---|---|---|---|---|
| BPgroup+Serbia_IronGates_Mesolithic+Trypillia | | 1.14E-01 | 2.13E-01 | 3.28E-01 | 2.82E-01 | 5.35E-02 |
| BPgroup+Serbia_IronGates_Mesolithic+YUN_CA | 1.70E-01 | | 1.81E-01 | 3.40E-01 | 2.24E-01 | 5.48E-02 |
| CoreYamna+GlobularAmphora+Serbia_IronGates_Mesolithic | 1.81E-03 | 7.24E-04 | | 1.81E-01 | 1.09E-01 | 1.40E-04 |
| CoreYamna+Serbia_IronGates_Mesolithic+Trypillia | 1.28E-03 | 1.17E-03 | 1.28E-01 | | 1.76E-01 | 6.42E-05 |
| CoreYamna+Serbia_IronGates_Mesolithic+YUN_CA | 1.48E-03 | 5.35E-04 | 1.15E-01 | 2.31E-01 | | 8.93E-05 |
| PVgroup+Serbia_IronGates_Mesolithic+Trypillia | 2.79E-01 | 1.59E-01 | 1.22E-01 | 1.70E-01 | 1.64E-01 | |

**Table S 24 Model competition for Ukraine_EBA_Deriivka_I4110.** Each column shows the p-value of a model when alternative models (rows) are placed on the Right set.

| | BPgroup+Serbia_IronGates_Mesolithic+Trypillia | BPgroup+Serbia_IronGates_Mesolithic+YUN_CA | CoreYamna+GlobularAmphora+Serbia_IronGates_Mesolithic | CoreYamna+Serbia_IronGates_Mesolithic+Trypillia | CoreYamna+Serbia_IronGates_Mesolithic+YUN_CA | PVgroup+Serbia_IronGates_Mesolithic+Trypillia |
|---|---|---|---|---|---|---|
| BPgroup+Serbia_IronGates_Mesolithic+Trypillia | | 1.45E-06 | 1.32E-05 | 8.00E-01 | 3.42E-05 | 6.73E-02 |
| BPgroup+Serbia_IronGates_Mesolithic+YUN_CA | 3.17E-02 | | 1.06E-01 | 1.14E-01 | 8.17E-01 | 3.65E-03 |
| CoreYamna+GlobularAmphora+Serbia_IronGates_Mesolithic | 4.83E-01 | 4.37E-01 | | 8.92E-01 | 9.00E-01 | 1.57E-01 |
| CoreYamna+Serbia_IronGates_Mesolithic+Trypillia | 4.12E-01 | 1.76E-06 | 8.57E-06 | | 2.17E-05 | 1.38E-01 |
| CoreYamna+Serbia_IronGates_Mesolithic+YUN_CA | 1.61E-02 | 3.76E-01 | 8.82E-02 | 1.01E-01 | | 6.05E-03 |
| PVgroup+Serbia_IronGates_Mesolithic+Trypillia | 4.02E-01 | 7.03E-07 | 2.54E-05 | 9.56E-01 | 6.29E-05 | |

**Table S 25 Model competition for Ukraine_EBA_Deriivnka_I5882.** Each column shows the p-value of a model when alternative models (rows) are placed on the Right set.

| | BPgroup+Serbia_IronGates_Mesolithic+Trypillia | BPgroup+Serbia_IronGates_Mesolithic+YUN_CA | CoreYamna+GlobularAmphora+Serbia_IronGates_Mesolithic | CoreYamna+Serbia_IronGates_Mesolithic+Trypillia | CoreYamna+Serbia_IronGates_Mesolithic+YUN_CA | PVgroup+Serbia_IronGates_Mesolithic+Trypillia |
|---|---|---|---|---|---|---|
| BPgroup+Serbia_IronGates_Mesolithic+Trypillia | | 6.19E-03 | 2.19E-02 | 2.77E-01 | 4.85E-02 | 7.74E-02 |
| BPgroup+Serbia_IronGates_Mesolithic+YUN_CA | 1.75E-01 | | 2.37E-01 | 3.68E-01 | 2.27E-01 | 1.21E-01 |
| CoreYamna+GlobularAmphora+Serbia_IronGates_Mesolithic | 1.48E-02 | 9.75E-03 | | 1.16E-01 | 9.72E-02 | 1.49E-02 |
| CoreYamna+Serbia_IronGates_Mesolithic+Trypillia | 2.35E-02 | 2.93E-03 | 1.26E-02 | | 2.95E-02 | 1.55E-02 |
| CoreYamna+Serbia_IronGates_Mesolithic+YUN_CA | 4.08E-02 | 1.46E-02 | 1.56E-01 | 2.65E-01 | | 2.78E-02 |
| PVgroup+Serbia_IronGates_Mesolithic+Trypillia | 1.97E-01 | 1.46E-02 | 1.61E-02 | 1.34E-01 | 3.53E-02 | |

**Table S 26 Model competition for Ukraine_EBA_Deriivnka_I5884.** Each column shows the p-value of a model when alternative models (rows) are placed on the Right set.

**Re-analysis of data from Penske et al. (2023) (ref.[22])**

We also re-analyzed the data from Penske et al. which partially overlap with the data of our study with regard to the cultural distribution of the newly reported data. We have already mentioned in passing that the MAJ population (from Mayaky) and USV can fit similarly to our Usatove population from the same site. We now look at other populations from the study.

**Chalcolithic Southeastern Europe (4900-3800 BCE)**

We used YUN_CA (Yunatsite) in our modeling as a source. According to ref.[22]:
*"The genetic homogeneity observed in and across the four CA sites (PIE, YUN, PTK and VAR) of the fifth millennium BC matches the cultural homogeneity of the archaeological records and suggests an extended period of a relative stable sociopolitical network and absence of large-scale cultural and genetic transformations."*

We confirm that PTK_CA (Tell Petko Karavelovo) is a clade with YUN_CA (p=0.146).

PIE_CA (Pietrele) does not form a clade with YUN_CA (p<1e-8) although it fits with that population as a source better than with any alternative (p<1e-107). Several subsets of PIE were analyzed separately in ref.[22] PIE039 (the earliest dated individual from the site) is a clade with YUN_CA (p=0.805). The individual PIE060 has 32.6±2.0% Iron Gates hunter-gatherer ancestry (p=0.755) in agreement with the observations of ref.[22] Individual PIE078 is not a clade with YUN_CA (p<1e-15) but is a clade with Trypillia (p=0.374). Overall, we assess that due to the large number of individuals in the PIE_CA population and the existence of outlier individuals at the site this population is overall similar to YUN_CA but includes other diversity that makes it fail the clade test with YUN_CA.

VAR_CA (Varna) does not form a clade with YUN_CA or any other population (p<1e-32) but is well-modeled as a mixture of YUN_CA and 8.8±0.7% Ukraine_N ancestry (p=0.154).

**Kartal clusters A and B (4150-3400 BCE)**

Of the two clusters at Kartal (ca. 4150–3400 BC; clusters KTL_A and KTL_B) we can model well KTL_A as a mixture of Trypillians and 53.9±1.2% BPgroup ancestry (p=0.618). The model PVgroup+Trypillia fails marginally (p=0.043).

In ref.[22] KTL_A is modeled as Steppe Eneolithic (corresponding to BPgroup or PVgroup in our setup) and VAR_CA. Both these models fail in our setup (p=6e-6 for PVgroup and p=0.002 for BPgroup). Examining qpAdm output we see that these models underestimate (Z=-4.3 and Z=-3.8 respectively) shared genetic drift with the Villabruna (WHG-related outgroup).

We conclude that the best reconstruction of this population of early steppe migrants is a combination of half Lower Volga-North Caucasus Eneolithic (BPgroup or PVgroup) and a Trypillian-related source. This scenario makes geographical sense as we do find the eastern migrants in unmixed form at Giurgiulești and we also know that these eastern migrants did admix with Trypillians to form Usatove (and USV/MAJ) and KTL_A could be derived from the same type of mixture. Note, however, that Usatove has a clear preference (as we explained above) for a PVgroup source and definitely has Aknashen-related ancestry from the Caucasus; the model without any such ancestry is the one that works for KTL_A. Thus, KTL_A represents a southern extension of CLV Cline migrants into Southeastern Europe, derived in this case from the Lower Volga end of the cline.

To verify this claim, we add as a 3rd population either Aknashen Neolithic or Lebyazhinka_HG: BPgroup stands at the end of two clines: the CLV cline to its south and the upriver Volga cline. By adding either of these two sources we can estimate where along either of these clines the admixing population was derived from. These two models are generalizations of the basic BPgroup+Trypillia model that fits KTL_A allowing for one additional source of ancestry (Table S 27). The proportion of Lebyazhinka ancestry is low and non-significant (1.6±2.2%) and so is that of Aknashen ancestry (-2.6±1.9%). Thus, we see no evidence that a population other than the BPgroup was involved in the westward migration. We also add as a 3rd source either Ukraine_N or GK2 to represent Dnipro-Don hunter-gatherer ancestry; these two are low and non-significant (2.8±2.1% or 2.7±2.3%). Thus, we see no evidence that the westward migrants picked any Dnipro-Don hunter-gatherer ancestry from the intervening territory between Trypillians and the Lower Volga.

| | | | | Proportions | | | Std. errors | | |
|---|---|---|---|---|---|---|---|---|---|
| A | B | C | P-value | A | B | C | A | B | C |
| Armenia_Aknashen_N | BPgroup | Trypillia | 0.555 | -2.6% | 55.0% | 47.6% | 3.4% | 1.9% | 2.3% |
| BPgroup | GK2 | Trypillia | 0.631 | 51.6% | 2.7% | 45.7% | 2.3% | 2.3% | 1.3% |
| BPgroup | Lebyazhinka_HG | Trypillia | 0.476 | 52.2% | 1.6% | 46.2% | 2.6% | 2.2% | 1.3% |
| BPgroup | Trypillia | Ukraine_N | 0.749 | 51.9% | 45.3% | 2.8% | 2.0% | 1.4% | 2.1% |

**Table S 27 KTL_A is derived from a BPgroup+Trypillia mixture even when adding sources defining the CLV (Aknashen) or Volga (Lebyazhinka) clines or allowing for Dnipro-Don hunter-gatherer (Ukraine_N or GK2) ancestry.**

Kartal B (KTL_B) does not fit any 2-way models; the only feasible one with p>0.001 involves 29.2±2.5% Maykop and 70.8±2.5%% Trypillian ancestry (p=0.011). This model underestimates shared genetic drift with Afontova Gora 3 (Z=-3.5).

We examined 3-way models (Table S 28). These invariably involve a CLV cline population (Aknashen, Maykop, or Remontnoye), a Dnipro-Don population (Ukraine_N or GK2) or Globular Amphora and YUN_CA.

| | | | | Proportions | | | Std. errors | | |
|---|---|---|---|---|---|---|---|---|---|
| A | B | C | P-value | A | B | C | A | B | C |

| | | | | | | | | | |
|---|---|---|---|---|---|---|---|---|---|
| Armenia_Aknashen_N | Ukraine_N | YUN_CA | 0.096 | 26.3% | 16.4% | 57.2% | 2.7% | 1.4% | 2.5% |
| GK2 | Maykop | YUN_CA | 0.064 | 14.3% | 24.7% | 61.0% | 1.6% | 2.7% | 2.2% |
| GlobularAmphora | Remontnoye | YUN_CA | 0.294 | 27.3% | 29.6% | 43.1% | 5.0% | 2.0% | 4.5% |
| Maykop | Ukraine_N | YUN_CA | 0.207 | 26.8% | 14.2% | 59.0% | 2.4% | 1.4% | 2.1% |
| Remontnoye | Serbia_IronGates_Mesolithic | YUN_CA | 0.370 | 28.2% | 7.3% | 64.5% | 2.0% | 1.2% | 1.8% |

**Table S 28 3-way admixture models for KTL_B.**

The model tournament competition between these models (Table S 29) excludes the models that involve Aknashen and GK2 ancestry but is otherwise inconclusive.

| | GK2+Maykop+YUN_CA | GlobularAmphora+Remontnoye+YUN_CA | Maykop+Ukraine_N+YUN_CA | Remontnoye+Serbia_IronGates_Mesolithic+YUN_CA | Armenia_Aknashen_N+Ukraine_N+YUN_CA |
|---|---|---|---|---|---|
| GK2+Maykop+YUN_CA | | 1.08E-01 | 2.68E-01 | 6.78E-02 | 1.99E-02 |
| GlobularAmphora+Remontnoye+YUN_CA | 1.76E-01 | | 4.47E-01 | 5.16E-01 | 2.19E-01 |
| Maykop+Ukraine_N+YUN_CA | 3.34E-02 | 5.84E-02 | | 1.18E-02 | 8.47E-03 |
| Remontnoye+Serbia_IronGates_Mesolithic+YUN_CA | 1.80E-01 | 3.99E-01 | 2.00E-01 | | 4.18E-02 |
| Armenia_Aknashen_N+Ukraine_N+YUN_CA | 3.98E-02 | 4.25E-01 | 2.15E-01 | 2.78E-01 | |

**Table S 29 Model competition for KTL_B.** Each column shows the p-value of a model when alternative models (rows) are placed on the Right set.

What is clear from the modeling of KTL_B is that it (i) has European farmer ancestry as its primary component, and (ii) that it has CLV cline ancestry. The CLV Cline ancestry is not (in contrast to KTL_A) from the northern (lower Volga) BPgroup end; rather it is from intermediate populations such as Maykop or Remontnoye.

An interpretation of these findings is that at least two groups participated in the westward movement of CLV Cline people into the ancestors of the Kartal populations: one consisting of unadmixed lower Volga-related (BPgroup) into the ancestors of KTL_A and another, consisting of mixed lower Volga-Caucasus Neolithic (Remontnoye or Maykop-related) into the ancestors of KTL_B.

In ref.[22] KTL_B is modeled in alternative ways, as a distal mix of CHG/EHG/Anatolian Neolithic; as a mixture of VAR_CA or Trypillians, Steppe Eneolithic (BPgroup or PVgroup in our setup), and Maykop; or as a simpler model of 12% Trypillian and 88% Maykop. This simpler 2-way model underestimates shared genetic drift with Afontova Gora 3 as we mentioned above; the models of Table S 28 account for this shared history in alternative ways: either (i) via Ukraine_N ancestry (itself a mixture of EHG and WHG ancestries[16], with EHG being a mixture of Ancient North Eurasians like Afontova Gora3 and

WHG[2]) or (ii) via Remontnoye ancestry (itself a mixture of BPgroup and Maykop/Aknashen, thus receiving Afontova Gora 3-related ancestry via BPgroup).

##### Early Bronze Age (BOY_EBA and MAJ_EBA) (3300-2500 BCE)

We show fitting 2-way models for Boyanovo and Mayaky populations of the Early Bronze Age in Table S 30. The EBA from Boyanovo like previously published data[12] and data included in ref.[3]and analyzed above as Bulgaria_EBA_Yamna can be modeled as a mixture of Core Yamna and European farmers.

| Test | A | B | P-value | A | B | S.E. |
|---|---|---|---|---|---|---|
| MAJ_EBA | GK2 | Remontnoye | 0.511 | 32.9% | 67.1% | 3.0% |
| MAJ_EBA | Remontnoye | Ukraine_N | 0.263 | 69.6% | 30.4% | 2.6% |
| BOY_EBA | BPgroup | GlobularAmphora | 0.255 | 77.8% | 22.2% | 1.5% |
| BOY_EBA | CoreYamna | GlobularAmphora | 0.658 | 92.1% | 7.9% | 1.5% |
| BOY_EBA | CoreYamna | Trypillia | 0.396 | 92.4% | 7.6% | 1.5% |
| BOY_EBA | CoreYamna | YUN_CA | 0.211 | 93.8% | 6.2% | 1.3% |
| BOY_EBA | GK2 | Remontnoye | 0.952 | 32.3% | 67.7% | 2.5% |

**Table S 30 Modeling BOY_EBA and MAJ_EBA.**

The results of the model tournament (Table S 31) show as above (Table S 4) that models involving Yamna ancestry are the only feasible ones.

| | BPgroup+GlobularAmphora | CoreYamna+GlobularAmphora | CoreYamna+Trypillia | CoreYamna+YUN_CA | GK2+Remontnoye |
|---|---|---|---|---|---|
| BPgroup+GlobularAmphora | | 5.73E-01 | 3.78E-01 | 1.63E-01 | 8.25E-01 |
| CoreYamna+GlobularAmphora | 3.21E-11 | | 4.85E-01 | 2.64E-01 | 9.78E-21 |
| CoreYamna+Trypillia | 4.35E-11 | 7.74E-02 | | 4.12E-02 | 5.38E-22 |
| CoreYamna+YUN_CA | 6.70E-11 | 7.56E-01 | 4.86E-01 | | 1.88E-18 |
| GK2+Remontnoye | 2.00E-01 | 2.48E-01 | 1.64E-01 | 8.51E-02 | |

**Table S 31 Model competition for BOY_EBA.** Each column shows the p-value of a model when alternative models (rows) are placed on the Right set.

Curiously no models with Yamna ancestry appear as feasible for the EBA from Mayaky (MAJ_EBA) in Table S 30. The model with CoreYamna+GlobularAmphora fails at the p=0.05 threshold (p=0.019). We examined 3-way models with Core Yamna as one of the sources. These models involve both about half to two thirds Yamna ancestry but also Dnipro-Don (Ukraine_N or GK2) ancestry as well as Caucasus Neolithic-related (Aknashen or Maykop) ancestry.

| | | | | Proportions | Std. errors |
|---|---|---|---|---|---|

| A | B | C | P-value | A | B | C | A | B | C |
|---|---|---|---|---|---|---|---|---|---|
| Armenia_Aknashen_N | CoreYamna | GK2 | 0.054 | 21.6% | 55.5% | 22.9% | 5.0% | 9.8% | 5.5% |
| CoreYamna | GK2 | Maykop | 0.163 | 50.7% | 23.0% | 26.3% | 9.5% | 5.0% | 5.2% |
| CoreYamna | Maykop | Ukraine_N | 0.084 | 64.2% | 19.6% | 16.2% | 6.3% | 4.0% | 3.2% |

**Table S 32 Feasible 3-way models for MAJ_EBA with Core Yamna as one of the sources.**

We have seen above in our analysis of KTL_B that Maykop-related ancestry flowed westwards during the Eneolithic and so its presence in MAJ_EBA could be related to that earlier migration. The combination of Core Yamna with GK2/Ukraine_N in the models of Table S 32 defines populations of the "Dnipro Cline" to which the Serednii Stih belonged. We show in linked manuscript ref.[3] that the Yamna of the lower Don had absorbed Serednii Stih ancestry and can be modeled as a mixture of Core Yamna with Serednii Stih populations having more Dnipro-Don (Ukraine_N/GK2) ancestry. To test if either Serednii Stih or Don Yamna could be contributing ancestry to MAJ_EBA we included Serednii Stih subsets (SShi, SSmed, SSlo) and Don Yamna as additional sources of ancestry.

We do indeed find simpler 2-way models with these additional sources of ancestry (Table S 33) that involve either Don Yamna or Serednii Stih. However, only models with Don Yamna and CLV Cline populations are feasible in the model tournament competition (Table S 34); the model with ~81% Don Yamna and ~19% Maykop ancestry fares particularly well. Recall that we inferred Core Yamna + Maykop admixture in the ancestry of Ukraine_EBA_Ozera_I1917. The EBA population at Mayaky represents another such mixture of Don Yamna and Maykop-related ancestry.

| A | B | P-value | A | B | S.E. |
|---|---|---|---|---|---|
| Armenia_Aknashen_N | Russia_Don_EBA_Yamna | 0.265 | 15.8% | 84.2% | 2.9% |
| GK2 | Remontnoye | 0.511 | 32.9% | 67.1% | 3.0% |
| Maykop | SSmed | 0.069 | 29.3% | 70.7% | 3.2% |
| Maykop | Russia_Don_EBA_Yamna | 0.424 | 18.9% | 81.1% | 3.1% |
| Remontnoye | Ukraine_N | 0.263 | 69.6% | 30.4% | 2.6% |
| Remontnoye | SSmed | 0.288 | 43.8% | 56.2% | 4.6% |
| Remontnoye | SSlo | 0.421 | 58.5% | 41.5% | 4.3% |
| Remontnoye | Russia_Don_EBA_Yamna | 0.174 | 29.2% | 70.8% | 5.2% |

**Table S 33 Feasible 2-way models for MAJ_EBA when Don Yamna and Serednii Stih are included as additional proximate sources.**

The BOY_EBA (and Southeastern European EBA populations in general) thus contrast with the MAJ_EBA: the former were formed on the basis of Yamna admixing with European farmer populations they encountered as the went beyond the steppe and into the territory of settled agriculturalists in the west; the latter had ancestry from the Don Yamna with Maykop-related people from the North Caucasus in the east.

| | Maykop+SSmed | Maykop+Russia_Don_EBA_Yamna | Remontnoye+Ukraine_N | Remontnoye+SSmed | Remontnoye+SSlo | Remontnoye+Russia_Don_EBA_Yamna | Armenia_Aknashen_N+Russia_Don_EBA_Yamna | GK2+Remontnoye |
|---|---|---|---|---|---|---|---|---|
| Maykop+SSmed | | 5.20E-01 | 2.00E-01 | 5.80E-02 | 1.15E-01 | 1.17E-01 | 2.06E-01 | 5.05E-01 |
| Maykop+Russia_Don_EBA_Yamna | 4.99E-11 | | 5.56E-12 | 6.17E-08 | 5.43E-07 | 1.77E-01 | 2.00E-01 | 1.08E-06 |
| Remontnoye+Ukraine_N | 2.78E-03 | 3.11E-01 | | 1.85E-01 | 3.09E-01 | 1.34E-01 | 2.43E-01 | 3.96E-01 |
| Remontnoye+SSmed | 1.58E-02 | 4.84E-01 | 3.59E-01 | | 5.62E-01 | 1.37E-01 | 3.51E-01 | 6.08E-01 |
| Remontnoye+SSlo | 1.83E-02 | 5.37E-01 | 3.33E-01 | 3.89E-01 | | 2.66E-01 | 4.41E-01 | 6.83E-01 |
| Remontnoye+Russia_Don_EBA_Yamna | 2.19E-10 | 4.40E-01 | 2.04E-12 | 1.51E-07 | 3.07E-06 | | 3.35E-01 | 3.41E-07 |
| Armenia_Aknashen_N+Russia_Don_EBA_Yamna | 1.08E-10 | 4.78E-01 | 8.83E-13 | 1.04E-07 | 5.82E-06 | 1.30E-01 | | 4.81E-07 |
| GK2+Remontnoye | 2.70E-02 | 5.97E-01 | 3.71E-01 | 3.03E-01 | 5.58E-01 | 2.62E-01 | 4.91E-01 | |

**Table S 34 Model competition for MAJ_EBA.** Each column shows the p-value of a model when alternative models (rows) are placed on the Right set.

We were curious if the Maykop-related admixture in MAJ_EBA was driven by any particular individual, so we fit the model to the three individuals from the site, confirming that it was indeed present in all of them.

| Individual | P-value | Maykop | Russia_Don_EBA_Yamna | S.E. |
|---|---|---|---|---|
| MAJ004.merge.bam | 0.417 | 20.0% | 80.0% | 4.4% |
| MAJ017.merge.bam | 0.581 | 17.9% | 82.1% | 4.0% |
| MAJ019.merge.bam | 0.263 | 21.7% | 78.3% | 4.4% |

**Table S 35 Admixture of MAJ_EBA individuals**

**Timing the admixture of Caucasus-Lower Volga and European farmer groups**

Both Usatove-related (Usatove, MAJ, and USV) and Kartal (KTL_A and KTL_B) populations are inferred to be admixtures of Caucasus Lower Volga people with European farmers. We use DATES[39] to date this admixture in number of generations and convert this to a calendar date by multiplying with 28 years per generation.[40] Our estimates place this admixture in the second half of the 5$^{th}$ millennium BCE (Fig. S 2) when we use (YUN_CA, Trypillia, GlobularAmphora, LBK) as the “European farmer” source and (PVgroup, BPgroup, Khi, Csongrád_I5124, Giurgiulești) as the “Caucasus-Volga” source. Removing Trypillia from the “European farmer” set and estimating its own admixture time, we obtain a similar date. This supports the idea that the formation of Trypillians (with little CLV ancestry) and Usatove (with about half) were part of the same mid-5$^{th}$ millennium BCE process.

This timeframe corresponds also with the emergence and efflorescence of the Serednii Stih culture from which the Yamna ancestral population was formed

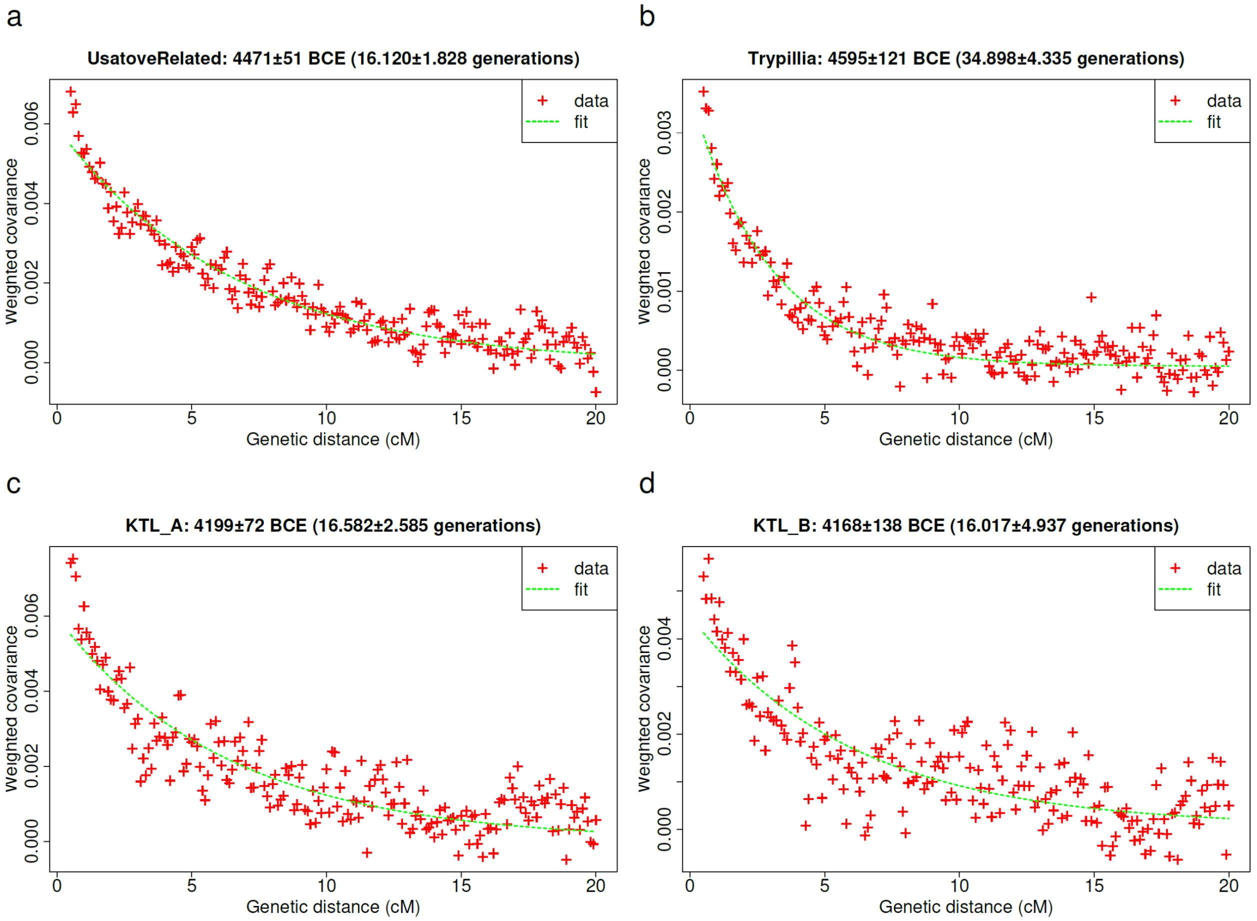

**Fig. S 2 DATES estimates of admixture timing of CLV and European farmer ancestry admixture.**

#### Summary of key findings

The discussion of this note suggests a complex landscape of interactions between people of the Pontic-Caspian steppe and the diverse descendants of European farmers and hunter-gatherers both before and after the formation and expansion of the Yamna archaeological horizon. We extract from our observations a list of key findings; we refer the reader to the preceding discussion for details of the modeling and possible alternative interpretations. The list below represents our synthesizing of the modeling of this note in a way that is not contradicted by the available evidence and should be evaluated critically in the future, both to confirm some of our conjectures and to propose alternative explanations for the available evidence.

1. We find relatively unadmixed representatives of the two expansive populations of the Eneolithic and Early Bronze Age: the Lower Volga-North Caucasus Eneolithic (BPgroup and PVgroup) and the Core Yamna (Table S 1). Of particular importance is the Eneolithic individual from Giurgiuleşti in Moldova which, together with the individual from Csongrád in Hungary discussed in ref.[3] proves the migration of unadmixed people from the Volga westwards. Both these migrants had no detectible ancestry from either the people of the Dnipro-Don region (through which they passed in their migratory route) or the European farmer populations that were settled near their sampling locations. We also find unadmixed Core Yamna either as individuals in

archaeologically uncertain contexts (Table S 1) and also among the Yamna of Bulgaria, Moldova, and Ukraine.

2. In Ukraine, Moldova, and Bulgaria, the Yamna included many individuals that had acquired European farmer ancestry (Table S 5;Table S 8)
3. An individual from Durankulak in Bulgaria is of particular interest as he possesses the "Corded Ware"-related R-M417 Y-haplogroup and a similar mix of Core Yamna and Globular Amphora ancestry as the Corded Ware (but with more ancestry from the Globular Amphora). This individual may stem from the admixture zone between the Core Yamna and the Globular Amphora from which the characteristic 3:1 blend[2,31] of the two components in the ancestry of the Corded Ware is derived.
4. There is some evidence of admixture between the expansive Yamna and populations of high hunter-gatherer ancestry in the Multi-Cordoned Ware / Babine. The existence of groups of both high and low hunter-gatherer ancestry has been observed in the Romanian Bronze Age[12]. The Yamna admixed with diverse farmer groups of both low hunter-gatherer ancestry (as the YUN_CA /Yunatsite of Bulgaria), higher such ancestry (as the Trypillians of Moldova-Ukraine), and higher still (as the Globular Amphora of Ukraine-Poland), and indeed other groups of even higher such ancestry as evidenced by both the Multi-Cordoned Ware / Babine and the Deriivka outliers of the Early Bronze Age.
5. The Yamna also admixed with the Steppe Maykop carrying this mix westwards to Taraclia (Table S 15) and Zhivotilovka (Table S 16). Yet, at Zhivotilovka we also find evidence of another individual whose ancestry is unclear but is connected to the Caucasus (Table S 20).
6. The Yamna also admixed with the Maykop or related populations as in Ukraine_EBA_Ozera_I1917 and MAJ_EBA. The latter population is the single example we know of the participation of migrations related to the Lower Don Yamna rather than the Core Yamna.
7. The CLV cline ancestry reached as far south as Bulgaria where a singleton individual from Riltsi possessed it in combination with local farmer ancestry (Table S 6).
8. The Usatove population was formed on the basis of North Caucasus Eneolithic ancestry (as in PVgroup) and Trypillians. It thus contrasts with the Serednii Stih which was formed in combination with Dnipro-Don hunter-gatherer ancestry. Different starting points along the CLV cline (in the east) combined with different local substrata populations (in the west) formed the contrasting genetic profiles of Serednii Stih and Usatove populations.
9. The Eneolithic population from Kartal included individuals well-described as mixtures of the Lower Volga Eneolithic (BPgroup) and Trypillians (Table S 27). Thus, we have evidence for the westward migration of both unadmixed people from the Volga and those who mixed with European farmer-descended populations. Kartal also included individuals with ancestry from more "southern" parts of the CLV Cline such as Maykop or Remontnoye admixing with European farmers (Table S 29). Westward migrations included people of "northern" Volga background (as in Csongrád and Giurgiuleşti) as well as of "southern" background (as in one of the Zhivotilovka individuals). In combination with European farmers these diverse CLV migrants formed diverse populations as in Kartal. At Mayaky in the Early Bronze Age, the Yamna migrants had absorbed some of the "southern" background ancestry; alternatively, Don Yamna acquired the "southern" (Maykop-related) ancestry in the east before migrating to Mayaky.

**The ancestral landscape: a compendium**

The key observation about the ancestral landscape of the people north and west of the Black Sea (in the North Pontic region and southeastern Europe) is their extreme diversity. Prior to the east-to-west migrations associated with Caucasus-Lower Volga (CLV) cline people, the people of the region had ancestry from local Dnipro-Don hunter-gatherers and European farmers with various amounts of Balkan hunter-gatherer (represented by the Iron Gates population from Serbia) ancestry. The CLV migrations were responsible for the emergence of the Yamna (likely in the Dnipro-Don region as part of the "Dnipro cline") and they were followed by the spread of the Yamna itself.

In Table 1 we summarize the analysis of this note presenting a "best guess" model for each of the populations we discussed.

#### Appendix I: Co-modeling populations with Dnipro-Don hunter-gatherer and European farmer ancestry

Populations of the Dnipro Cline can be modeled as mixtures of three sources: Aknashen-BPgroup (defining the CLV cline) and GK2 (as a representative of Dnipro-Don hunter-gatherers). However, we have seen that in the NPR admixture from European farmer descended populations (such as Trypillians) is important in the origins of many populations such as those at Mayaky and Kartal (Table 1).

To jointly model both the Core Yamna (and other populations of the Dnipro Cline) and populations with Trypillian/European farmer ancestry, we would like to fit a 4-source model. However, when we fit this model to the Core Yamna or Usatove, i.e., populations with Dnipro-Don hunter-gatherer and European farmer ancestry respectively, we observe that its standard errors are quite high (Table S 36).

| | | Proportions | | | | Std. errors | | | |
|---|---|---|---|---|---|---|---|---|---|
| Test | P-value | Aknashen | BPgroup | GK2 | Trypillia | Aknashen | BPgroup | GK2 | Trypillia |
| CoreYamna | 0.892 | 18.6% | 58.7% | 21.2% | 1.5% | 16.3% | 17.1% | 12.1% | 11.2% |
| Usatove | 0.416 | 24.4% | 25.8% | 7.9% | 41.8% | 45.7% | 49.3% | 34.5% | 31.0% |
| PeriPontic | 0.965 | 22.4% | 47.4% | 26.9% | 3.3% | 15.2% | 16.1% | 11.1% | 10.2% |

**Table S 36 4-source model with Right=Base.** High standard errors necessitate the introduction of an outgroup population to Base that can differentiate between the four sources.

##### Choosing additional Right populations

To reduce them we can add a new outgroup population to the Right set that can distinguish adequately between the four sources; however, if this outgroup population shares genetic drift with either the test population (e.g., Core Yamna) or the 4 sources in violation of the phylogenetic assumptions of qpAdm[2] it may result in the quintuple of populations (Core Yamna, Aknashen, BPgroup, GK2, Trypillians) being descended from five waves in relation to the Right set, i.e., the test population can no longer be modeled as the proposed 4-way admixture.

Thus, we want to find a phylogenetically informative outgroup to add to the Right set that both reduces the standard errors of the modeling (allowing us to estimate admixture proportions more accurately), but also does not violate the phylogenetic assumptions.

Moreover, this sought-after outgroup must be useful to reduce standard errors for both Core Yamna and Usatove and indeed among a variety of other populations that we wish to model. In order not to show preference for any of the targets, we combine all populations that can be feasibly modeled with the 3-source model, as well as those which also have Trypillian ancestry (Table 1) into a "PeriPontic" (circum-Black Sea) meta-population:

**"PeriPontic":** BOY_EBA, Bulgaria_EBA_Yamna, CoreYamna, GK1, Igren_o, KTL_A, Maykop, MAJ, MAJ_EBA, Moldova_Crasnoe_Eneolithic, Moldova_EBA_Yamna, Moldova_EBA_Yamna_o, Moldova_GlobularAmphora_Yamna, Mykhailivka_I32534, PVgroup, Remontnoye, Russia_Don_EBA_Yamna, SShi, SSlo, SSmed, SS_ukr104, Ukraine_EBA_Catacomb, Ukraine_EBA_Ozera_I1917, Ukraine_EBA_Yamna, Ukraine_Oleksandria_MBA, Ukraine_Vasilevka, Usatove, Usatove_EBA, Usatove_Yamna, USV, Zhivotilovka_I17974

Individuals of the "PeriPontic" population are quite diverse but they should all have (in our reconstruction) ancestry from the four sources or their subset. We confirm this by fitting the 4-source model on the PeriPontic meta-population whose overall admixture proportions reflect an average across its many diverse sub-populations (Table S 36).

In order to find such an outgroup we tried all population labels in our dataset not included in PeriPontic, the four sources or Base, plotting P-values and standard errors in Fig. S 3. We see that several outgroups give us leverage to estimate proportions accurately while remaining feasible. We show the top-20 (lowest standard error) models in Table S 37.

Choosing which outgroup to add is a subjective decision; ideally, we would like to avoid outgroups that are "too close" to the modeled and source populations. Such populations may be rejected if they violate the phylogenetic assumptions. For example, the considered outgroup "China_Xinjiang_Ayituohan_Afanasievo_BA.SG"[41] does so (p=2e-5), as it is a Yamna-Afanasievo related population from the eastern edge of the distribution of this genetic cluster and obviously shares post admixture genetic drift with "PeriPontic" which includes many Yamna individuals. Other violations may not be detected by a low p-value (which depends on data quality, and the amount of shared genetic drift) but will bias admixture inferences subtly.

Thus, we should also avoid populations whose estimates are far off from the "consensus" across the different choices. We decided to use the sum of the squared differences of inferred admixture proportions for each outgroup choice from the median (across all 20 outgroups) as a measure of proximity to the consensus (Table S 37). This measure identifies the Tarim_EMBA1 population from Xinjiang, China as the outgroup of choice. Standard errors are minimized for another population from the same region (China_Xinjiang_Xiaohe_BA.SG). Both these populations make good outgroups because of their distance to our region of interest and their description of possessing late "Ancient North Eurasian" (ANE) ancestry unrelated to the steppe migrations (of Afanasievo-related groups).[41]

We were curious about why these two outgroups could differentiate between the sources, so we estimated statistics of the form $f_4$(Added Outgroup, OldAfrica; $\text{Source}_1$, $\text{Source}_2$). These indicate (Table S 38) that they do so by strongly differentiating steppe sources (GK2 and BPgroup) from farmer ones (Aknashen and Trypillians).

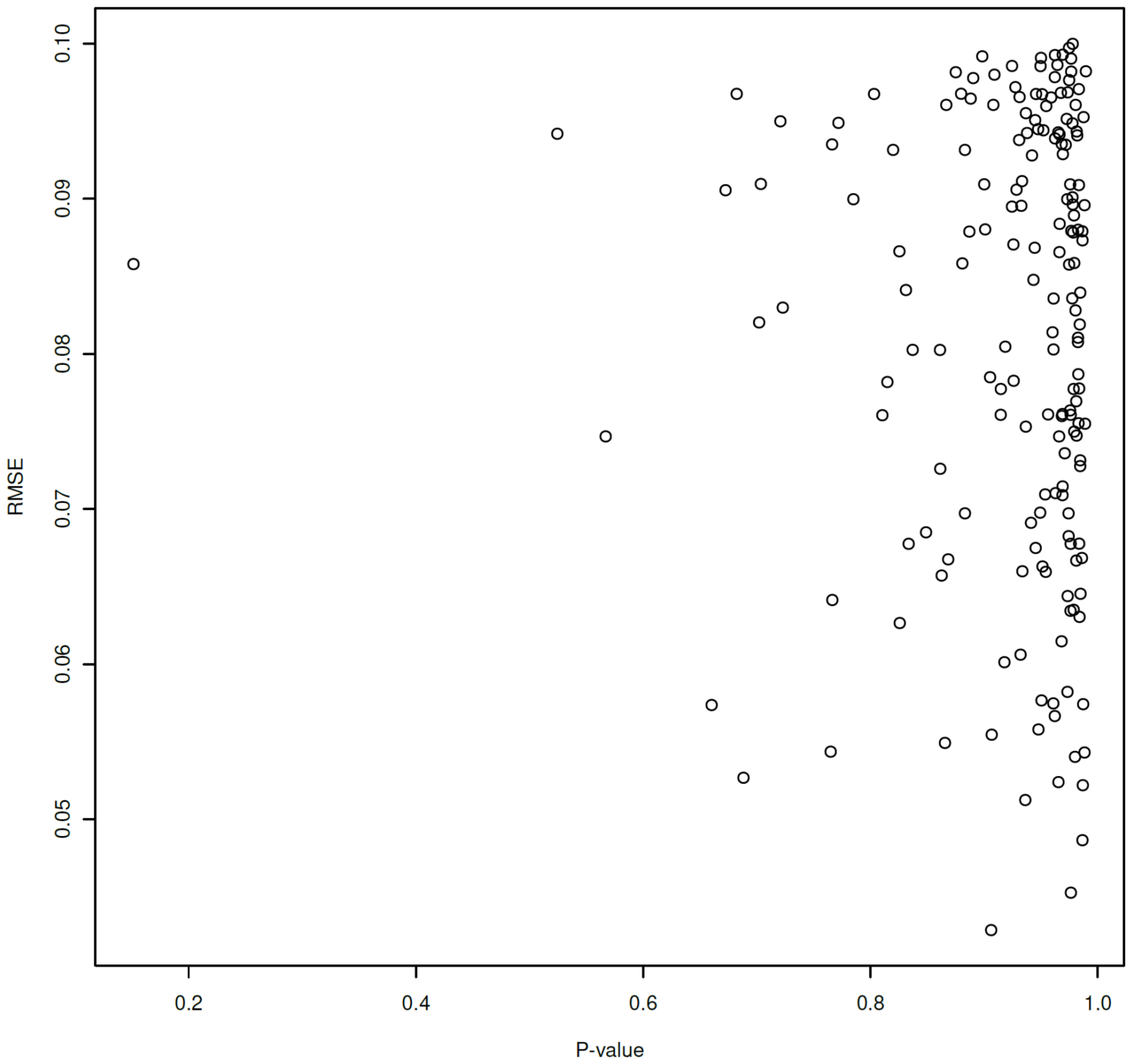

**Fig. S 3 P-value and Root Mean Square Error (RMSE) of 4-way model (Aknashen, BPgroup, GK2, Trypillia) when adding diverse outgroups to the Base set.** Models with <10% RMSE are shown.

| | | Proportions | | | | Std. errors | | | | RMSE | Proximity to Median |
|---|---|---|---|---|---|---|---|---|---|---|---|
| **Added outgroup** | **P-value** | **Aknashen** | **BPgroup** | **GK2** | **Trypillia** | **Aknashen** | **BPgroup** | **GK2** | **Trypillia** | | |
| **Tarim_EMBA1** | **0.962** | **19.5%** | **50.4%** | **25.0%** | **5.1%** | **6.3%** | **6.6%** | **4.9%** | **4.6%** | **5.7%** | **3.8E-06** |
| Russia_Khvalynsk_Eneolithic_brother.I6734 | 0.973 | 19.8% | 50.2% | 25.0% | 4.9% | 6.5% | 6.8% | 5.0% | 4.7% | 5.8% | 5.8E-06 |
| Ukraine_Deriivka_Mes | 0.948 | 20.2% | 50.0% | 24.9% | 5.0% | 6.6% | 6.5% | 4.4% | 4.4% | 5.6% | 4.0E-05 |
| Usatove_I20078 | 0.965 | 19.2% | 50.9% | 24.6% | 5.3% | 6.0% | 6.0% | 4.4% | 4.3% | 5.2% | 7.5E-05 |
| WHG | 0.961 | 19.5% | 50.8% | 24.3% | 5.4% | 6.8% | 6.6% | 4.5% | 4.7% | 5.7% | 8.2E-05 |
| Jordan_EBA | 0.936 | 18.8% | 51.0% | 24.5% | 5.7% | 5.7% | 6.1% | 4.4% | 4.0% | 5.1% | 1.8E-04 |
| China_Xinjiang_Beifang_Xiaohe_BA.SG | 0.907 | 18.6% | 51.6% | 24.0% | 5.7% | 6.2% | 6.5% | 4.7% | 4.5% | 5.5% | 4.1E-04 |
| Serbia_IronGates_Mesolithic | 0.951 | 21.0% | 49.1% | 25.6% | 4.3% | 6.9% | 6.7% | 4.4% | 4.6% | 5.8% | 4.2E-04 |
| Germany_Blatterhohle_MN | 0.918 | 18.3% | 52.0% | 23.7% | 6.0% | 7.0% | 6.9% | 4.9% | 4.9% | 6.0% | 7.2E-04 |
| Csongrád_I5124 | 0.987 | 21.4% | 48.4% | 26.3% | 3.9% | 6.5% | 6.6% | 4.9% | 4.7% | 5.7% | 9.8E-04 |
| Latvia_HG | 0.866 | 18.0% | 52.7% | 22.8% | 6.6% | 6.6% | 6.4% | 4.1% | 4.4% | 5.5% | 1.6E-03 |
| **China_Xinjiang_Xiaohe_BA.SG** | **0.906** | **17.4%** | **52.8%** | **23.3%** | **6.6%** | **4.7%** | **4.9%** | **3.8%** | **3.6%** | **4.3%** | **1.6E-03** |
| TUR_E_Arslantepe_ChL | 0.765 | 22.0% | 47.8% | 26.7% | 3.6% | 5.9% | 6.6% | 4.8% | 4.1% | 5.4% | 1.7E-03 |
| Russia_Khvalynsk_Eneolithic_brother.I6739 | 0.980 | 22.1% | 47.7% | 26.7% | 3.5% | 6.0% | 6.2% | 4.7% | 4.5% | 5.4% | 1.8E-03 |
| Kazakhstan_Kumsay_EBA | 0.987 | 22.2% | 47.6% | 26.7% | 3.4% | 5.4% | 5.5% | 4.3% | 4.1% | 4.9% | 2.0E-03 |
| Russia_Steppe_Maykop | 0.987 | 22.7% | 47.2% | 27.1% | 3.1% | 5.7% | 6.0% | 4.6% | 4.4% | 5.2% | 2.7E-03 |
| Russia_CaspianInland_EBA_Yamna_Yamna_o1 | 0.988 | 22.8% | 47.0% | 27.2% | 3.0% | 6.1% | 6.2% | 4.7% | 4.5% | 5.4% | 3.0E-03 |
| Khi | 0.976 | 23.6% | 46.2% | 27.7% | 2.5% | 5.0% | 5.0% | 4.0% | 4.0% | 4.5% | 4.6E-03 |
| Serbia_IronGates | 0.688 | 16.1% | 54.8% | 21.2% | 7.9% | 6.4% | 6.1% | 3.9% | 4.2% | 5.3% | 5.5E-03 |
| TUR_E_Arslantepe_ChL_brother.ART014 | 0.660 | 14.9% | 54.9% | 21.8% | 8.4% | 6.3% | 6.9% | 5.0% | 4.4% | 5.7% | 6.5E-03 |
| **Maximum difference** | | 8.7% | 8.7% | 6.5% | 5.9% | | | | | | |

**Table S 37 Adding outgroups to Base set.** We model the "PeriPontic" meta-population with diverse added outgroups to Base. Top-20 models of lowest RMSE are shown. Proximity to median is the sum of the squared difference between admixture proportions for each added outgroup to the median proportions across all 20 outgroups. The two chosen outgroups for addition are highlighted in bold and minimize the RMSE (China_Xinjiang_Xiaohe_BA.SG) and proximity to median (Tarim_EMBA1).

| Added outgroup | $Source_1$ | $Source_2$ | f4(Added outgroup, OldAfrica, $Source_1$, $Source_2$) | Z-score |
|---|---|---|---|---|
| Tarim_EMBA1 | Armenia_Aknashen_N | BPgroup | -0.009261 | -20.6 |
| Tarim_EMBA1 | Armenia_Aknashen_N | GK2 | -0.010226 | -17.2 |
| Tarim_EMBA1 | Armenia_Aknashen_N | Trypillia | 0.000128 | 0.3 |
| Tarim_EMBA1 | BPgroup | GK2 | -0.001062 | -2.3 |
| Tarim_EMBA1 | BPgroup | Trypillia | 0.009411 | 33.4 |
| Tarim_EMBA1 | GK2 | Trypillia | 0.010428 | 23.1 |
| China_Xinjiang_Xiaohe_BA.SG | Armenia_Aknashen_N | BPgroup | -0.009354 | -21.0 |
| China_Xinjiang_Xiaohe_BA.SG | Armenia_Aknashen_N | GK2 | -0.009771 | -16.5 |
| China_Xinjiang_Xiaohe_BA.SG | Armenia_Aknashen_N | Trypillia | 0.000043 | 0.1 |
| China_Xinjiang_Xiaohe_BA.SG | BPgroup | GK2 | -0.000284 | -0.6 |
| China_Xinjiang_Xiaohe_BA.SG | BPgroup | Trypillia | 0.009456 | 33.5 |
| China_Xinjiang_Xiaohe_BA.SG | GK2 | Trypillia | 0.009712 | 21.3 |

**Table S 38 Statistics showing how added Right outgroup populations differentiate between the populations of the 4-source model.**

#### Modeling Peri-Pontic populations

We modeled all Peri-Pontic sub-populations using the original Base set, augmenting it with either of the two identified outgroup populations, or with both (Table S 39).

| | | | Proportions | | | | Std. errors | | | | |
|---|---|---|---|---|---|---|---|---|---|---|---|
| **Test** | **Outgroup set** | **P-value** | **Aknashen** | **BPgroup** | **GK2** | **Trypillia** | **Aknashen** | **BPgroup** | **GK2** | **Trypillia** | **RMSE** |
| BOY_EBA | Base | 0.596 | 22.0% | 46.6% | 25.7% | 5.7% | 39.1% | 42.5% | 30.3% | 26.9% | 35.3% |
| BOY_EBA | +Tarim_EMBA1 | 0.717 | 24.7% | 43.9% | 27.5% | 3.9% | 13.5% | 14.5% | 11.0% | 10.0% | 12.4% |
| BOY_EBA | +Xiaohe_BA.SG | 0.763 | 23.9% | 44.7% | 26.9% | 4.4% | 8.7% | 9.0% | 6.9% | 6.7% | 7.9% |
| BOY_EBA | +both | 0.833 | 24.1% | 44.5% | 27.0% | 4.4% | 8.7% | 9.0% | 7.0% | 6.6% | 7.9% |
| Bulgaria_EBA_Yamna | Base | 0.828 | 25.0% | 40.0% | 23.2% | 11.8% | 34.8% | 36.9% | 26.1% | 24.1% | 31.0% |
| Bulgaria_EBA_Yamna | +Tarim_EMBA1 | 0.931 | 21.5% | 44.0% | 20.5% | 14.0% | 12.0% | 12.6% | 9.4% | 8.9% | 10.8% |
| Bulgaria_EBA_Yamna | +Xiaohe_BA.SG | 0.810 | 14.8% | 50.9% | 15.9% | 18.3% | 9.2% | 9.5% | 7.3% | 7.0% | 8.3% |
| Bulgaria_EBA_Yamna | +both | 0.817 | 14.8% | 51.1% | 15.5% | 18.6% | 13.0% | 13.5% | 10.5% | 9.9% | 11.8% |
| CoreYamna | Base | 0.892 | 19.3% | 58.0% | 21.7% | 1.0% | 12.9% | 13.7% | 9.6% | 8.7% | 11.4% |
| CoreYamna | +Tarim_EMBA1 | 0.892 | 15.8% | 61.7% | 19.3% | 3.2% | 6.5% | 6.8% | 5.1% | 4.8% | 5.9% |
| CoreYamna | +Xiaohe_BA.SG | 0.861 | 13.9% | 63.8% | 17.8% | 4.5% | 5.1% | 5.2% | 4.0% | 3.8% | 4.6% |

| | | | | | | | | | | | |
|---|---|---|---|---|---|---|---|---|---|---|---|
| CoreYamna | +both | 0.884 | 14.0% | 63.7% | 17.9% | 4.5% | 5.2% | 5.4% | 4.1% | 3.9% | 4.7% |
| GK1 | Base | 0.814 | 30.9% | 8.3% | 79.5% | -18.7% | 132.5% | 138.0% | 96.0% | 90.6% | 116.2% |
| GK1 | +Tarim_EMBA1 | 0.895 | 27.2% | 12.6% | 76.7% | -16.5% | 46.3% | 50.0% | 37.9% | 34.2% | 42.6% |
| GK1 | +Xiaohe_BA.SG | 0.871 | 18.6% | 21.3% | 70.9% | -10.8% | 15.8% | 16.6% | 12.6% | 11.9% | 14.4% |
| GK1 | +both | 0.806 | 19.6% | 20.5% | 71.1% | -11.2% | 16.3% | 17.5% | 13.3% | 12.1% | 15.0% |
| Igren_o | Base | 0.644 | 38.7% | -34.9% | 120.0% | -23.9% | 1545.3% | 1538.0% | 1058.0% | 1064.1% | 1323.4% |
| Igren_o | +Tarim_EMBA1 | 0.659 | 13.1% | -10.5% | 102.7% | -5.3% | 43.8% | 45.5% | 34.2% | 32.6% | 39.4% |
| Igren_o | +Xiaohe_BA.SG | 0.820 | -12.4% | 16.7% | 84.8% | 10.9% | 26.8% | 27.9% | 21.7% | 20.6% | 24.5% |
| Igren_o | +both | 0.539 | -9.9% | 14.2% | 85.4% | 10.4% | 46.7% | 48.4% | 37.6% | 35.8% | 42.5% |
| KTL_A | Base | 0.526 | 8.1% | 43.5% | 8.4% | 40.0% | 126.9% | 131.1% | 88.9% | 84.7% | 110.0% |
| KTL_A | +Tarim_EMBA1 | 0.446 | 18.3% | 32.8% | 15.6% | 33.4% | 20.3% | 21.6% | 16.0% | 14.7% | 18.4% |
| KTL_A | +Xiaohe_BA.SG | 0.344 | 26.5% | 24.3% | 21.4% | 27.8% | 9.3% | 9.3% | 7.1% | 7.1% | 8.3% |
| KTL_A | +both | 0.313 | 27.1% | 23.7% | 22.2% | 27.1% | 9.4% | 9.6% | 7.4% | 7.2% | 8.5% |
| Maykop | Base | 0.533 | 114.2% | -18.3% | 23.3% | -19.2% | 33.0% | 34.8% | 24.4% | 22.6% | 29.2% |
| Maykop | +Tarim_EMBA1 | 0.620 | 103.2% | -6.4% | 15.7% | -12.5% | 16.8% | 16.9% | 12.6% | 12.7% | 14.9% |
| Maykop | +Xiaohe_BA.SG | 0.679 | 106.4% | -9.9% | 17.7% | -14.1% | 12.9% | 13.0% | 10.1% | 10.2% | 11.6% |
| Maykop | +both | 0.688 | 107.8% | -11.0% | 19.1% | -15.8% | 14.5% | 14.4% | 11.1% | 11.4% | 12.9% |
| MAJ | Base | 0.106 | 10.1% | 37.5% | 0.1% | 52.3% | 94.8% | 99.0% | 69.9% | 65.8% | 83.7% |
| MAJ | +Tarim_EMBA1 | 0.146 | 19.2% | 27.8% | 6.5% | 46.5% | 28.9% | 30.5% | 22.7% | 21.1% | 26.1% |
| MAJ | +Xiaohe_BA.SG | 0.125 | 23.9% | 22.9% | 9.9% | 43.3% | 8.7% | 8.7% | 6.7% | 6.7% | 7.8% |
| MAJ | +both | 0.216 | 22.6% | 24.3% | 9.0% | 44.2% | 18.1% | 17.7% | 13.1% | 13.6% | 15.8% |
| MAJ_EBA | Base | 0.151 | 62.3% | 2.2% | 56.1% | -20.6% | 383.5% | 397.8% | 279.1% | 264.6% | 336.6% |
| MAJ_EBA | +Tarim_EMBA1 | 0.004 | 24.0% | 42.1% | 30.5% | 3.4% | 124.1% | 125.2% | 91.7% | 90.6% | 109.2% |
| MAJ_EBA | +Xiaohe_BA.SG | 0.015 | 21.1% | 45.3% | 27.4% | 6.2% | 22.9% | 23.4% | 17.7% | 17.3% | 20.5% |
| MAJ_EBA | +both | 0.007 | 46.2% | 17.4% | 49.4% | -13.0% | 892.4% | 993.0% | 764.2% | 663.6% | 837.7% |
| Moldova_Crasnoe_Eneolithic | Base | 0.530 | 49.5% | 26.3% | 41.7% | -17.6% | 330.5% | 336.8% | 237.8% | 231.7% | 288.5% |
| Moldova_Crasnoe_Eneolithic | +Tarim_EMBA1 | 0.266 | 17.4% | 60.4% | 18.8% | 3.3% | 40.7% | 43.9% | 31.8% | 28.7% | 36.8% |
| Moldova_Crasnoe_Eneolithic | +Xiaohe_BA.SG | 0.582 | 25.8% | 51.8% | 24.1% | -1.7% | 14.1% | 14.2% | 10.8% | 10.7% | 12.6% |
| Moldova_Crasnoe_Eneolithic | +both | 0.213 | 32.2% | 44.9% | 30.0% | -7.1% | 17.4% | 17.6% | 13.5% | 13.3% | 15.6% |
| Moldova_EBA_Yamna | Base | 0.737 | 22.5% | 49.4% | 23.1% | 5.0% | 21.9% | 23.3% | 16.6% | 15.2% | 19.6% |
| Moldova_EBA_Yamna | +Tarim_EMBA1 | 0.838 | 17.4% | 54.6% | 19.7% | 8.3% | 8.7% | 9.0% | 6.7% | 6.4% | 7.8% |
| Moldova_EBA_Yamna | +Xiaohe_BA.SG | 0.770 | 15.9% | 56.4% | 18.4% | 9.4% | 6.9% | 7.0% | 5.4% | 5.3% | 6.2% |
| Moldova_EBA_Yamna | +both | 0.868 | 16.0% | 56.1% | 18.6% | 9.3% | 6.7% | 6.8% | 5.2% | 5.1% | 6.0% |
| Moldova_EBA_Yamna_o | Base | 0.374 | -3.0% | 50.9% | 3.2% | 48.9% | 1017.0% | 1050.4% | 730.9% | 697.6% | 888.6% |
| Moldova_EBA_Yamna_o | +Tarim_EMBA1 | 0.265 | -2.2% | 50.6% | 4.2% | 47.4% | 42.9% | 43.7% | 32.7% | 31.9% | 38.2% |
| Moldova_EBA_Yamna_o | +Xiaohe_BA.SG | 0.458 | 3.3% | 44.4% | 8.2% | 44.1% | 15.5% | 15.5% | 12.0% | 12.0% | 13.9% |
| Moldova_EBA_Yamna_o | +both | 0.275 | 6.0% | 41.7% | 10.5% | 41.8% | 16.0% | 16.2% | 12.6% | 12.4% | 14.4% |
| Moldova_GlobularAmphora_Yamna | Base | 0.647 | 64.0% | -2.5% | 58.4% | -20.0% | 50.8% | 52.5% | 36.8% | 35.2% | 44.5% |
| Moldova_GlobularAmphora_Yamna | +Tarim_EMBA1 | 0.035 | 37.0% | 25.9% | 41.5% | -4.3% | 117.5% | 130.9% | 97.2% | 83.9% | 108.9% |
| Moldova_GlobularAmphora_Yamna | +Xiaohe_BA.SG | 0.195 | 31.6% | 31.8% | 35.8% | 0.7% | 17.3% | 17.9% | 13.6% | 13.1% | 15.6% |
| Moldova_GlobularAmphora_Yamna | +both | 0.056 | 39.9% | 22.7% | 43.6% | -6.2% | 21.2% | 21.7% | 16.4% | 15.8% | 19.0% |
| Mykhailivka_I32534 | Base | 0.261 | 41.5% | 42.6% | 33.5% | -17.7% | 268.0% | 280.0% | 195.6% | 183.8% | 235.7% |
| Mykhailivka_I32534 | +Tarim_EMBA1 | 0.426 | 25.3% | 59.3% | 22.1% | -6.6% | 73.2% | 74.8% | 54.3% | 52.8% | 64.6% |
| Mykhailivka_I32534 | +Xiaohe_BA.SG | 0.436 | 19.1% | 65.4% | 17.3% | -1.9% | 30.0% | 30.5% | 23.7% | 23.3% | 27.1% |
| Mykhailivka_I32534 | +both | 0.576 | 21.1% | 63.5% | 18.3% | -2.9% | 33.0% | 33.4% | 25.5% | 25.0% | 29.5% |
| PVgroup | Base | 0.767 | 26.8% | 75.9% | 8.6% | -11.3% | 33.8% | 35.3% | 24.4% | 23.0% | 29.6% |
| PVgroup | +Tarim_EMBA1 | 0.882 | 26.8% | 76.0% | 8.6% | -11.4% | 13.1% | 13.2% | 9.7% | 9.6% | 11.5% |
| PVgroup | +Xiaohe_BA.SG | 0.844 | 30.6% | 72.0% | 11.1% | -13.8% | 10.1% | 9.8% | 7.4% | 7.6% | 8.8% |
| PVgroup | +both | 0.849 | 31.9% | 70.7% | 12.2% | -14.8% | 13.1% | 12.8% | 9.3% | 9.6% | 11.3% |
| Remontnoye | Base | 0.480 | 29.8% | 70.1% | -10.7% | 10.7% | 151.8% | 169.8% | 120.5% | 102.5% | 138.7% |
| Remontnoye | +Tarim_EMBA1 | 0.656 | 28.0% | 72.3% | -12.1% | 11.9% | 14.8% | 15.1% | 11.1% | 10.9% | 13.1% |
| Remontnoye | +Xiaohe_BA.SG | 0.621 | 34.4% | 65.5% | -7.7% | 7.8% | 10.4% | 10.7% | 8.2% | 7.9% | 9.4% |
| Remontnoye | +both | 0.573 | 36.5% | 63.4% | -5.9% | 6.0% | 11.1% | 11.4% | 8.7% | 8.4% | 10.0% |
| Russia_Don_EBA_Yamna | Base | 0.733 | 32.8% | 23.3% | 52.4% | -8.5% | 20.9% | 22.2% | 15.5% | 14.1% | 18.5% |
| Russia_Don_EBA_Yamna | +Tarim_EMBA1 | 0.755 | 25.5% | 31.1% | 47.4% | -3.9% | 9.8% | 10.4% | 7.8% | 7.2% | 8.9% |
| Russia_Don_EBA_Yamna | +Xiaohe_BA.SG | 0.466 | 19.9% | 37.1% | 43.3% | -0.3% | 7.8% | 8.0% | 6.1% | 6.0% | 7.0% |
| Russia_Don_EBA_Yamna | +both | 0.484 | 19.9% | 37.2% | 43.1% | -0.2% | 7.8% | 8.0% | 6.1% | 5.8% | 7.0% |
| SShi | Base | 0.402 | 34.2% | 33.6% | 46.7% | -14.4% | 90.0% | 94.4% | 64.0% | 59.6% | 78.5% |
| SShi | +Tarim_EMBA1 | 0.464 | 36.8% | 30.7% | 48.6% | -16.1% | 15.8% | 16.4% | 12.1% | 11.5% | 14.1% |
| SShi | +Xiaohe_BA.SG | 0.571 | 26.5% | 41.8% | 41.0% | -9.3% | 10.9% | 10.9% | 8.3% | 8.2% | 9.7% |
| SShi | +both | 0.279 | 27.3% | 41.2% | 41.0% | -9.5% | 12.7% | 12.8% | 9.7% | 9.6% | 11.3% |
| SSlo | Base | 0.283 | 108.2% | -88.7% | 145.1% | -64.5% | 324.6% | 347.6% | 241.4% | 218.5% | 288.2% |
| SSlo | +Tarim_EMBA1 | 0.409 | 83.1% | -62.4% | 127.4% | -48.1% | 37.9% | 39.9% | 29.9% | 27.9% | 34.3% |
| SSlo | +Xiaohe_BA.SG | 0.312 | 57.1% | -34.3% | 108.6% | -31.3% | 24.0% | 25.0% | 19.5% | 18.4% | 21.9% |
| SSlo | +both | 0.257 | 60.8% | -38.4% | 110.9% | -33.2% | 31.4% | 33.1% | 25.3% | 23.7% | 28.7% |
| SSmed | Base | 0.864 | 36.5% | 7.2% | 70.2% | -13.9% | 52.0% | 55.6% | 38.0% | 34.5% | 45.9% |
| SSmed | +Tarim_EMBA1 | 0.909 | 26.9% | 17.6% | 63.3% | -7.8% | 15.3% | 15.9% | 11.8% | 11.3% | 13.7% |
| SSmed | +Xiaohe_BA.SG | 0.764 | 18.1% | 27.1% | 56.9% | -2.0% | 11.6% | 12.1% | 9.3% | 8.8% | 10.5% |
| SSmed | +both | 0.675 | 17.2% | 27.8% | 55.9% | -1.0% | 12.3% | 12.8% | 9.6% | 9.2% | 11.1% |
| SS_ukr104 | Base | 0.360 | 67.9% | -18.7% | 85.5% | -34.7% | 1199.9% | 1296.9% | 923.3% | 826.7% | 1079.1% |
| SS_ukr104 | +Tarim_EMBA1 | 0.529 | 27.7% | 25.5% | 54.1% | -7.3% | 25.7% | 26.9% | 20.1% | 18.8% | 23.1% |
| SS_ukr104 | +Xiaohe_BA.SG | 0.422 | 4.3% | 50.5% | 37.1% | 8.1% | 14.9% | 15.6% | 12.1% | 11.4% | 13.6% |
| SS_ukr104 | +both | 0.159 | 7.1% | 47.6% | 38.8% | 6.5% | 92.7% | 99.4% | 74.5% | 67.7% | 84.6% |
| Ukraine_EBA_Catacomb | Base | 0.613 | 36.3% | 32.4% | 32.9% | -1.6% | 329.0% | 325.5% | 229.3% | 233.0% | 283.3% |
| Ukraine_EBA_Catacomb | +Tarim_EMBA1 | 0.736 | 21.9% | 47.2% | 23.3% | 7.6% | 20.5% | 21.4% | 16.2% | 15.3% | 18.5% |
| Ukraine_EBA_Catacomb | +Xiaohe_BA.SG | 0.770 | 26.0% | 42.9% | 25.9% | 5.2% | 14.5% | 15.0% | 11.7% | 11.3% | 13.2% |
| Ukraine_EBA_Catacomb | +both | 0.740 | 30.4% | 38.2% | 29.9% | 1.5% | 15.6% | 16.2% | 12.4% | 11.9% | 14.2% |
| Ukraine_EBA_Ozera_I1917 | Base | 0.477 | 104.6% | -25.3% | 54.7% | -33.9% | 48.8% | 51.2% | 36.0% | 33.6% | 43.1% |
| Ukraine_EBA_Ozera_I1917 | +Tarim_EMBA1 | 0.096 | 60.2% | 21.8% | 23.7% | -5.7% | 545.7% | 547.6% | 390.5% | 388.6% | 474.6% |
| Ukraine_EBA_Ozera_I1917 | +Xiaohe_BA.SG | 0.043 | 54.4% | 28.3% | 18.6% | -1.2% | 80.8% | 80.6% | 64.4% | 64.6% | 73.1% |
| Ukraine_EBA_Ozera_I1917 | +both | 0.076 | 57.3% | 25.3% | 21.1% | -3.7% | 28.5% | 30.3% | 22.5% | 20.9% | 25.9% |
| Ukraine_EBA_Yamna | Base | 0.494 | 22.7% | 47.0% | 24.9% | 5.5% | 60.8% | 64.1% | 44.3% | 41.1% | 53.5% |
| Ukraine_EBA_Yamna | +Tarim_EMBA1 | 0.539 | 17.5% | 52.5% | 21.4% | 8.6% | 29.3% | 29.9% | 21.9% | 21.3% | 25.9% |
| Ukraine_EBA_Yamna | +Xiaohe_BA.SG | 0.337 | 8.5% | 62.0% | 14.8% | 14.7% | 8.7% | 8.8% | 6.7% | 6.5% | 7.8% |

| | | | | | | | | | | | |
|---|---|---|---|---|---|---|---|---|---|---|---|
| Ukraine_EBA_Yamna | +both | 0.310 | 8.2% | 62.5% | 14.2% | 15.1% | 12.1% | 12.3% | 9.0% | 8.8% | 10.7% |
| Ukraine_Oleksandria_MBA | Base | 0.666 | 9.8% | 61.0% | 25.1% | 4.1% | 95.2% | 101.0% | 69.4% | 63.6% | 83.9% |
| Ukraine_Oleksandria_MBA | +Tarim_EMBA1 | 0.666 | -19.1% | 91.7% | 5.0% | 22.4% | 28.4% | 28.6% | 21.2% | 21.0% | 25.1% |
| Ukraine_Oleksandria_MBA | +Xiaohe_BA.SG | 0.806 | 2.8% | 68.6% | 19.9% | 8.7% | 19.5% | 19.3% | 14.9% | 15.1% | 17.3% |
| Ukraine_Oleksandria_MBA | +both | 0.429 | 2.4% | 69.1% | 20.4% | 8.0% | 23.6% | 23.8% | 18.5% | 18.3% | 21.2% |
| Ukraine_Vasilevka | Base | 0.086 | 21.5% | -31.7% | 129.2% | -19.0% | 529.3% | 543.6% | 393.5% | 379.2% | 467.5% |
| Ukraine_Vasilevka | +Tarim_EMBA1 | 0.140 | 32.7% | -43.5% | 137.2% | -26.4% | 32.0% | 33.6% | 25.1% | 23.4% | 28.9% |
| Ukraine_Vasilevka | +Xiaohe_BA.SG | 0.168 | 13.9% | -23.5% | 123.7% | -14.0% | 20.9% | 21.9% | 16.7% | 15.7% | 19.0% |
| Ukraine_Vasilevka | +both | 0.090 | 14.0% | -23.2% | 122.6% | -13.3% | 21.5% | 22.5% | 17.0% | 16.1% | 19.5% |
| Usatove | Base | 0.416 | 23.0% | 27.4% | 6.8% | 42.9% | 45.6% | 44.6% | 30.4% | 31.3% | 38.6% |
| Usatove | +Tarim_EMBA1 | 0.520 | 19.3% | 31.2% | 4.3% | 45.3% | 10.7% | 11.0% | 8.4% | 8.1% | 9.6% |
| Usatove | +Xiaohe_BA.SG | 0.537 | 20.7% | 29.7% | 5.2% | 44.3% | 7.4% | 7.4% | 5.7% | 5.8% | 6.6% |
| Usatove | +both | 0.613 | 20.7% | 29.7% | 5.4% | 44.2% | 7.3% | 7.4% | 5.8% | 5.7% | 6.6% |
| Usatove_EBA | Base | 0.881 | 70.8% | 0.5% | 61.8% | -33.2% | 93.5% | 99.7% | 68.8% | 62.7% | 82.7% |
| Usatove_EBA | +Tarim_EMBA1 | 0.944 | 63.8% | 7.5% | 57.0% | -28.4% | 34.3% | 35.9% | 27.0% | 25.6% | 31.0% |
| Usatove_EBA | +Xiaohe_BA.SG | 0.931 | 65.7% | 6.0% | 57.9% | -29.5% | 24.1% | 25.1% | 19.9% | 19.0% | 22.2% |
| Usatove_EBA | +both | 0.951 | 64.9% | 6.4% | 57.6% | -29.0% | 25.9% | 26.5% | 20.8% | 20.3% | 23.5% |
| Usatove_Yamna | Base | 0.842 | 53.3% | 14.7% | 49.9% | -17.9% | 29.2% | 30.8% | 21.6% | 20.0% | 25.8% |
| Usatove_Yamna | +Tarim_EMBA1 | 0.815 | 62.2% | 5.0% | 56.5% | -23.8% | 20.9% | 21.6% | 16.0% | 15.3% | 18.7% |
| Usatove_Yamna | +Xiaohe_BA.SG | 0.874 | 38.7% | 30.4% | 39.5% | -8.6% | 14.4% | 14.6% | 11.1% | 10.9% | 12.9% |
| Usatove_Yamna | +both | 0.078 | 42.1% | 26.3% | 41.4% | -9.8% | 25.3% | 25.3% | 18.9% | 18.8% | 22.3% |
| USV | Base | 0.021 | -5.1% | 60.8% | -16.6% | 60.9% | 1757.8% | 2004.7% | 1390.5% | 1144.5% | 1608.7% |
| USV | +Tarim_EMBA1 | 0.044 | 22.3% | 29.3% | 7.5% | 40.9% | 25.3% | 26.1% | 19.5% | 18.7% | 22.6% |
| USV | +Xiaohe_BA.SG | 0.041 | 31.3% | 20.0% | 13.9% | 34.8% | 13.0% | 13.1% | 10.1% | 10.0% | 11.6% |
| USV | +both | 0.047 | 31.3% | 20.0% | 14.3% | 34.4% | 11.5% | 11.5% | 8.8% | 8.8% | 10.2% |
| Zhivotilovka_I17974 | Base | 0.133 | 38.1% | 41.5% | 38.6% | -18.1% | 345.6% | 374.6% | 259.5% | 230.5% | 308.3% |
| Zhivotilovka_I17974 | +Tarim_EMBA1 | 0.055 | -12.3% | 94.6% | 3.0% | 14.7% | 90.2% | 94.0% | 71.2% | 67.3% | 81.5% |
| Zhivotilovka_I17974 | +Xiaohe_BA.SG | 0.099 | -5.9% | 88.2% | 7.1% | 10.7% | 18.3% | 18.5% | 13.5% | 13.3% | 16.1% |
| Zhivotilovka_I17974 | +both | 0.064 | -2.7% | 84.6% | 10.0% | 8.1% | 19.1% | 19.8% | 14.7% | 14.0% | 17.1% |

**Table S 39 4-source modeling of Peri-Pontic populations with Trypillian ancestry as the 4$^{th}$ source.**

We plot the admixture proportions estimated using the Xiaohe BA outgroup (which have the lowest standard errors) in Extended Data Fig. 1. By jointly modeling them we can compare them using the same 4-source model which agrees qualitiatively with the results of the simpler 2- and 3-source models derived in this note and in ref.[3]. The three major groups apparent in our modeling are the CLV populations (Remontnoye and PVgroup) characterized by high BPgroup/Aknashen proportions; the Dnipro Cline populations (including several Yamna subsets) characterized by high but dampened amounts of the CLV cline ancestral sources complemented by substantial GK2-related ancestry; and the populations related to Usatove (including also MAJ and KTL_A) in which the Trypillian ancestry is highest.

We were concerned by the fact that KTL_A is shown in Extended Data Fig. 1 have both GK2 and Aknashen-related ancestry while it fits the much simpler 2-source BPgroup+Trypillia model as we discussed above. Given the high standard errors of this population we do not think there is strong evidence for this ancestry (Table S 39). Additionally, as we previously examined the BPgroup+Trypillia+(GK2 or Aknashen) 3-source models using the Base outgroup set and find that in both the amount of the 3$^{rd}$ source (GK2 or Aknashen) is low and non-significant (Table S 27). Thus, we think that the reconstruction of the ancestry of this population as a 2-source mixture of BPgroup and Trypillians reflects its most likely history ().

#### Appendix II: Co-modeling populations with Dnipro-Don hunter-gatherer and Steppe Maykop ancestry

We also wanted to co-model the two individuals with Steppe Maykop ancestry (Table 1) and thus add Steppe Maykop as a 4th source to the Aknashen-BPgroup-GK2 model. We repeat the analysis of Table S 39 for the set of populations that fit the 3-source model and do not have farmer ancestry and the two individuals with Steppe Maykop ancestry:

**"PeriPontic1":** CoreYamna, GK1, Igren_o, Maykop, MAJ_EBA, Moldova_Crasnoe_Eneolithic, PVgroup, Remontnoye, Russia_Don_EBA_Yamna, SShi, SSlo, SSmed, SS_ukr104, Ukraine_EBA_Catacomb, Ukraine_EBA_Ozera_I1917, Usatove_I20078, Usatove_Yamna, Zhivotilovka_I17974

Admixture with Steppe Maykop as the 4th source are shown in Table S 40 and visualized in Extended Data Fig. 2. The Steppe Maykop ancestry is significantly higher than zero for Usatove_I20078 even with the Base set of outgroups (55.6±5.3%), similar to the 60.6±6.2% estimated for the simpler Core Yamna + Steppe Maykop model for this individual. For Zhivotilovka_I17974 the Steppe Maykop ancestry is 18.2±6.0% for the Core Yamna + Steppe Maykop model; its nominal value is ~17-18% in Table S 40 but with a higher standard error. As we argued above, this individual too is likely to have Steppe Maykop-related ancestry, but the statistical evidence for it is weaker than is the case for Usatove_I20078.

| | | | Proportions | | | | Std. errors | | | | |
|---|---|---|---|---|---|---|---|---|---|---|---|
| Test | Outgroup set | P-value | Aknashen | BPgroup | GK2 | Steppe Maykop | Aknashen | BPgroup | GK2 | Steppe Maykop | RMSE |
| CoreYamna | Base | 0.851 | 20.4% | 57.6% | 22.8% | -0.8% | 2.0% | 6.4% | 1.8% | 5.2% | 4.3% |
| CoreYamna | +Tarim_EMBA1 | 0.826 | 20.7% | 55.1% | 22.4% | 1.8% | 1.9% | 5.4% | 1.7% | 3.7% | 3.5% |
| CoreYamna | +Xiaohe_BA.SG | 0.710 | 20.9% | 53.9% | 21.9% | 3.2% | 1.9% | 5.7% | 1.6% | 3.9% | 3.7% |
| CoreYamna | +both | 0.729 | 20.7% | 55.0% | 22.0% | 2.3% | 1.9% | 5.6% | 1.7% | 3.7% | 3.6% |
| GK1 | Base | 0.774 | 1.0% | 47.7% | 61.4% | -10.0% | 5.0% | 16.5% | 4.8% | 13.6% | 11.2% |
| GK1 | +Tarim_EMBA1 | 0.869 | 1.0% | 46.8% | 61.0% | -8.8% | 4.7% | 13.7% | 4.5% | 9.6% | 9.0% |
| GK1 | +Xiaohe_BA.SG | 0.847 | 1.4% | 44.4% | 60.9% | -6.6% | 4.8% | 13.7% | 4.3% | 9.5% | 8.9% |
| GK1 | +both | 0.851 | 0.8% | 46.6% | 60.7% | -8.0% | 4.7% | 13.5% | 4.5% | 9.1% | 8.8% |
| Igren_o | Base | 0.765 | 1.4% | 1.6% | 98.0% | -1.0% | 10.0% | 34.0% | 9.3% | 28.1% | 23.1% |
| Igren_o | +Tarim_EMBA1 | 0.839 | -0.2% | 10.3% | 99.4% | -9.6% | 9.4% | 26.9% | 8.9% | 18.3% | 17.5% |
| Igren_o | +Xiaohe_BA.SG | 0.854 | 1.9% | -4.5% | 96.5% | 6.0% | 10.1% | 29.4% | 8.5% | 20.0% | 19.0% |
| Igren_o | +both | 0.581 | -0.9% | 7.7% | 97.0% | -3.8% | 9.5% | 27.0% | 8.4% | 18.2% | 17.5% |
| Maykop | Base | 0.316 | 87.0% | 9.4% | 4.1% | -0.6% | 5.2% | 16.7% | 4.3% | 12.7% | 11.0% |
| Maykop | +Tarim_EMBA1 | 0.434 | 86.6% | 10.4% | 4.1% | -1.1% | 5.0% | 14.0% | 4.0% | 8.4% | 8.8% |
| Maykop | +Xiaohe_BA.SG | 0.374 | 86.2% | 14.6% | 5.2% | -6.1% | 5.1% | 14.8% | 3.9% | 8.9% | 9.2% |
| Maykop | +both | 0.267 | 86.8% | 10.9% | 5.3% | -3.0% | 5.0% | 14.4% | 3.9% | 8.7% | 9.0% |
| MAJ_EBA | Base | 0.130 | 35.2% | 20.7% | 33.5% | 10.6% | 4.7% | 15.5% | 3.9% | 12.3% | 10.4% |
| MAJ_EBA | +Tarim_EMBA1 | 0.107 | 36.9% | 9.9% | 32.1% | 21.1% | 4.4% | 12.2% | 3.5% | 8.0% | 7.8% |
| MAJ_EBA | +Xiaohe_BA.SG | 0.147 | 36.5% | 12.7% | 32.2% | 18.6% | 4.4% | 12.4% | 3.4% | 8.1% | 7.9% |
| MAJ_EBA | +both | 0.149 | 36.9% | 10.2% | 32.4% | 20.5% | 4.4% | 12.0% | 3.6% | 7.8% | 7.7% |
| Moldova_Crasnoe_Eneolithic | Base | 0.627 | 21.1% | 65.1% | 24.6% | -10.7% | 5.1% | 16.8% | 4.8% | 13.4% | 11.3% |
| Moldova_Crasnoe_Eneolithic | +Tarim_EMBA1 | 0.250 | 23.0% | 50.7% | 21.8% | 4.5% | 5.2% | 15.4% | 4.5% | 10.3% | 9.9% |
| Moldova_Crasnoe_Eneolithic | +Xiaohe_BA.SG | 0.575 | 21.8% | 58.6% | 22.5% | -2.9% | 4.9% | 14.5% | 4.2% | 9.5% | 9.2% |
| Moldova_Crasnoe_Eneolithic | +both | 0.180 | 22.9% | 52.5% | 23.2% | 1.4% | 5.3% | 16.0% | 4.2% | 10.4% | 10.1% |
| PVgroup | Base | 0.644 | 9.9% | 94.2% | -2.6% | -1.5% | 3.9% | 13.2% | 3.2% | 10.0% | 8.7% |
| PVgroup | +Tarim_EMBA1 | 0.734 | 9.2% | 97.8% | -2.1% | -4.8% | 3.7% | 11.2% | 3.2% | 7.3% | 7.1% |
| PVgroup | +Xiaohe_BA.SG | 0.568 | 8.7% | 101.7% | -1.3% | -9.1% | 3.8% | 11.7% | 3.0% | 7.6% | 7.4% |
| PVgroup | +both | 0.511 | 9.0% | 99.3% | -1.0% | -7.3% | 3.5% | 10.4% | 2.9% | 6.8% | 6.6% |
| Remontnoye | Base | 0.402 | 46.6% | 47.0% | -0.9% | 7.3% | 4.3% | 14.0% | 3.6% | 10.4% | 9.2% |
| Remontnoye | +Tarim_EMBA1 | 0.566 | 46.6% | 46.5% | -1.1% | 8.0% | 4.1% | 11.6% | 3.5% | 7.2% | 7.3% |
| Remontnoye | +Xiaohe_BA.SG | 0.568 | 46.6% | 47.8% | -0.7% | 6.3% | 4.1% | 11.6% | 3.3% | 7.4% | 7.4% |
| Remontnoye | +both | 0.610 | 46.9% | 46.2% | -0.6% | 7.6% | 3.9% | 10.8% | 3.3% | 6.7% | 6.8% |
| Russia_Don_EBA_Yamna | Base | 0.684 | 20.0% | 38.1% | 43.7% | -1.7% | 2.7% | 9.0% | 2.6% | 7.2% | 6.1% |
| Russia_Don_EBA_Yamna | +Tarim_EMBA1 | 0.772 | 20.2% | 36.4% | 43.3% | 0.1% | 2.5% | 7.2% | 2.4% | 5.0% | 4.7% |
| Russia_Don_EBA_Yamna | +Xiaohe_BA.SG | 0.581 | 20.6% | 33.7% | 42.7% | 2.9% | 2.7% | 7.8% | 2.2% | 5.3% | 5.0% |
| Russia_Don_EBA_Yamna | +both | 0.517 | 20.0% | 36.4% | 42.6% | 1.0% | 2.5% | 7.4% | 2.3% | 5.0% | 4.8% |
| SShi | Base | 0.426 | 10.4% | 66.3% | 32.5% | -9.3% | 3.5% | 11.7% | 3.2% | 9.4% | 7.9% |
| SShi | +Tarim_EMBA1 | 0.520 | 9.7% | 69.8% | 33.1% | -12.7% | 3.5% | 10.3% | 3.1% | 6.9% | 6.6% |
| SShi | +Xiaohe_BA.SG | 0.582 | 10.5% | 65.8% | 32.5% | -8.7% | 3.4% | 10.0% | 2.8% | 6.8% | 6.4% |
| SShi | +both | 0.444 | 9.5% | 69.9% | 32.3% | -11.8% | 3.6% | 10.5% | 3.0% | 6.9% | 6.7% |

| | | | | | | | | | | | |
|---|---|---|---|---|---|---|---|---|---|---|---|
| SSlo | Base | 0.385 | 1.8% | 58.9% | 83.2% | -43.9% | 8.5% | 29.8% | 8.0% | 24.6% | 20.2% |
| SSlo | +Tarim_EMBA1 | 0.477 | 3.8% | 48.1% | 81.9% | -33.8% | 7.6% | 22.6% | 7.3% | 15.2% | 14.6% |
| SSlo | +Xiaohe_BA.SG | 0.347 | 4.4% | 42.5% | 80.3% | -27.2% | 8.3% | 24.5% | 6.9% | 15.9% | 15.6% |
| SSlo | +both | 0.382 | 4.0% | 45.0% | 80.1% | -29.1% | 7.5% | 21.9% | 7.0% | 14.3% | 14.0% |
| SSmed | Base | 0.814 | 14.4% | 35.3% | 56.6% | -6.3% | 4.0% | 13.4% | 3.7% | 11.0% | 9.1% |
| SSmed | +Tarim_EMBA1 | 0.872 | 14.8% | 32.7% | 55.9% | -3.4% | 3.9% | 10.8% | 3.3% | 7.2% | 7.0% |
| SSmed | +Xiaohe_BA.SG | 0.736 | 15.4% | 28.7% | 55.0% | 0.9% | 3.7% | 10.7% | 3.2% | 7.3% | 6.9% |
| SSmed | +both | 0.628 | 15.1% | 30.6% | 55.2% | -0.8% | 3.7% | 10.7% | 3.2% | 7.4% | 6.9% |
| SS_ukr104 | Base | 0.394 | 16.4% | 36.9% | 45.9% | 0.8% | 5.8% | 19.9% | 5.2% | 16.4% | 13.5% |
| SS_ukr104 | +Tarim_EMBA1 | 0.553 | 15.7% | 40.6% | 46.4% | -2.8% | 5.1% | 15.0% | 4.7% | 10.4% | 9.8% |
| SS_ukr104 | +Xiaohe_BA.SG | 0.481 | 17.6% | 28.4% | 44.1% | 9.9% | 5.2% | 15.0% | 4.7% | 10.0% | 9.7% |
| SS_ukr104 | +both | 0.119 | 15.6% | 36.6% | 44.0% | 3.8% | 6.0% | 17.5% | 4.9% | 11.5% | 11.2% |
| Ukraine_EBA_Catacomb | Base | 0.747 | 37.1% | 19.2% | 30.7% | 13.0% | 6.4% | 22.0% | 5.6% | 17.1% | 14.6% |
| Ukraine_EBA_Catacomb | +Tarim_EMBA1 | 0.848 | 36.0% | 21.0% | 31.0% | 11.9% | 6.0% | 17.6% | 5.3% | 11.4% | 11.2% |
| Ukraine_EBA_Catacomb | +Xiaohe_BA.SG | 0.849 | 36.1% | 23.1% | 31.3% | 9.5% | 5.9% | 17.5% | 5.2% | 10.8% | 11.0% |
| Ukraine_EBA_Catacomb | +both | 0.818 | 35.5% | 23.3% | 31.8% | 9.3% | 6.0% | 17.3% | 5.2% | 10.8% | 10.9% |
| Ukraine_EBA_Ozera_I1917 | Base | 0.128 | 54.2% | 31.0% | 20.8% | -6.0% | 6.0% | 19.9% | 5.2% | 15.2% | 13.1% |
| Ukraine_EBA_Ozera_I1917 | +Tarim_EMBA1 | 0.100 | 56.5% | 16.2% | 18.4% | 8.9% | 5.9% | 17.3% | 4.9% | 10.7% | 10.9% |
| Ukraine_EBA_Ozera_I1917 | +Xiaohe_BA.SG | 0.074 | 56.8% | 14.6% | 17.5% | 11.0% | 6.0% | 17.3% | 4.9% | 10.6% | 10.9% |
| Ukraine_EBA_Ozera_I1917 | +both | 0.118 | 56.5% | 15.1% | 17.5% | 10.9% | 5.8% | 16.9% | 4.7% | 10.3% | 10.6% |
| Usatove_I20078 | Base | 0.740 | 5.8% | 36.6% | 1.9% | 55.6% | 5.0% | 16.8% | 4.3% | 13.5% | 11.3% |
| Usatove_I20078 | +Tarim_EMBA1 | 0.840 | 5.4% | 40.0% | 2.6% | 52.1% | 4.7% | 13.8% | 3.8% | 9.3% | 8.9% |
| Usatove_I20078 | +Xiaohe_BA.SG | 0.822 | 5.6% | 38.9% | 2.1% | 53.4% | 4.7% | 13.6% | 3.7% | 9.2% | 8.7% |
| Usatove_I20078 | +both | 0.861 | 5.3% | 40.5% | 2.3% | 51.9% | 4.6% | 13.1% | 3.6% | 8.9% | 8.4% |
| Usatove_Yamna | Base | 0.705 | 28.2% | 37.2% | 31.3% | 3.3% | 5.2% | 17.1% | 4.4% | 13.4% | 11.4% |
| Usatove_Yamna | +Tarim_EMBA1 | 0.462 | 26.3% | 48.9% | 33.8% | -9.1% | 5.4% | 15.0% | 4.2% | 9.8% | 9.6% |
| Usatove_Yamna | +Xiaohe_BA.SG | 0.819 | 27.8% | 39.4% | 31.5% | 1.4% | 5.2% | 14.4% | 4.3% | 9.4% | 9.2% |
| Usatove_Yamna | +both | 0.081 | 27.0% | 44.3% | 32.1% | -3.4% | 5.8% | 16.2% | 4.6% | 10.1% | 10.2% |
| Zhivotilovka_I17974 | Base | 0.173 | 13.8% | 57.0% | 18.2% | 11.1% | 5.1% | 16.6% | 4.4% | 12.7% | 11.0% |
| Zhivotilovka_I17974 | +Tarim_EMBA1 | 0.188 | 14.8% | 48.7% | 16.9% | 19.5% | 4.8% | 13.7% | 4.2% | 8.6% | 8.7% |
| Zhivotilovka_I17974 | +Xiaohe_BA.SG | 0.234 | 14.5% | 51.3% | 16.9% | 17.3% | 4.8% | 14.1% | 4.1% | 9.2% | 9.0% |
| Zhivotilovka_I17974 | +both | 0.255 | 15.1% | 48.2% | 17.3% | 19.4% | 5.0% | 13.9% | 4.2% | 8.6% | 8.8% |

**Table S 40 4-source modeling of Peri-Pontic populations with Steppe Maykop ancestry as the 4$^{th}$ source.**

The models of Extended Data Fig. 1 and Extended Data Fig. 2 also increase our confidence in the fact that the Core Yamna was indeed formed by the type of admixture of CLV and Dnipro-Don people inferred in ref.[3]. The fourth source, whether Trypillian (from the west) or Steppe Maykop (from the east) contributes a non-significant amount of ancestry to this population (4.5±3.8% and 3.2±3.9%). We cannot really exclude that some such ancestry was present in the Core Yamna population, but the bulk of its ancestry was indeed made up of the postulated CLV-Dnipro/Don mixture.

The Dnipro-Don area did indeed see diverse admixtures in which the CLV-Dnipro/Don mixture was primary and represented the coming together of CLV people (themselves of mixed Aknashen/BPgroup ancestry) with local Dnipro-Don (GK2 or Ukraine_N-related) hunter-gatherers and in which (or through which) both European farmer ancestry (in and from the west) and Steppe Maykop ancestry (from the east) flowed, creating the diverse tapestry of ancestry characterizing its Eneolithic and Bronze Age populations.

#### Appendix III: Do the core Yamna have Trypillian ancestry?

The fact that Trypillian ancestry exists in many populations of the NPR raises the question if it might also exist in low levels in the Core Yamna. While all populations of the Dnipro cline (such as the Serednii Stih and Yamna) were modeled without such ancestry in ref.[3], as the result of admixture between Caucasus-Lower Volga people with Dnipro-Don hunter-gatherer descendants, it is possible that some such ancestry may exist in them. As a further test of the possible presence of Trypillian ancestry in the Core Yamna we sought to increase power in the Base set by modifying it as follows, highlighting the newly introduced populations:

**Base1:** OldAfrica, CHG, Iran_GanjDareh_N, **Serbia_IronGates_Mesolithic, Tarim_EMBA1**, Russia_Sidelkino.SG, **Jordan_EBA**, Turkey_N, **Armenia_Aknashen_N, Trypillia**

This set includes more recent (and higher data quality) populations for both the WHG-related (Iron Gates instead of Villabruna) and Siberian-related (Tarim EMBA1 instead of Afontova Gora 3) ancestry, an additional EBA population from the Levant (Jordan_EBA) and both the Aknashen Neolithic and Trypillian populations. The Aknashen and Trypillia populations will, of course, not be used on the Right when they are included as sources on the left, but their inclusion there allows us to test 3-source models (with only one of the two populations including) for resilience when the other population is on the Right.

The inclusion of such more recent populations on the Base1 risks that they may have had opportunity to share common genetic history with either the sources or target population in violation of the qpAdm phylogenetic assumptions.[2] However, the hope is that they may be able to differentiate the sources more powerfully. This increased power can be assessed by examining whether the fitted model can be estimated with lower standard errors using Base1 vs. Base or the augmented Base sets with the addition of outgroup populations (Table S 39); we will empirically see this below. But, at the outset, we can test whether we can reject 3 waves for the four sources (Aknashen, BPgroup, GK2, Trypillia) using qpWave. We can do so using both the conservative Base set (p=0.0012) but much more distinctly using the Base1 set (p=4e-15). Thus, a priori, before examining the fit of the model to the core Yamna we may think that the Base1 set does indeed provide more power.

The 4-source model remains feasible (p=0.915) with a significant amount of Aknashen (15.6±4.3%) but not Trypillian (3.2±3.1%) ancestry. When Trypillia is excluded as a source, the model still fits (p=0.230) with 21.8±1.4% Aknashen-related ancestry, showing that it is resilient to the presence of Trypillian on the Base1 set; conversely, when Aknashen is excluded as a source and Trypillia is included, then the model does not fit albeit weakly (p=0.010), suggesting that Trypillia is not resilient to the presence of a Caucasus Neolithic population. Examining the qpAdm output reveals that the Trypillia model underestimates (Z=-2.0) shared genetic drift with Jordan_EBA, a Levantine population without known steppe ancestry,[9,12] but with Caucasus-related ancestry suggesting that it may share common history with Aknashen that is not well-modeled when Aknashen is placed on the right set.

The standard error of Trypillian as a 4$^{th}$ source is somewhat improved (3.1% vs. 4.6% for the best choice in Table S 39 and 8.7% for the Base set), but remains consistent with both some European farmer (Trypillian-related) ancestry and with a scenario in which no such ancestry is present. Trypillian-related ancestry in the Yamna is thus possible but unproven. However, using any of the augmented Base sets of Table S 39 or the alternative Base1 set, the proportion of Aknashen-related ancestry remains significant (15.6±4.3% corresponds to Z=3.6 for the 4-source model, and the simpler 3-source model remains feasible with Trypillian on the right set), therefore the presence of some such ancestry must be considered certain.

##### Appendix IV: Source of hunter-gatherer ancestry in the core Yamna along the UNHG-GK2-EHG continuum

In ref.[3] we noted that populations of the Dnipro-Don cline (Serednii Stih and Yamna) had Ukraine Neolithic (UNHG)-related hunter-gatherer ancestry which could be modeled with either UNHG (from the Dnipro) or a subset of Golubaya Krinitsa (GK2) from the Middle Don. As GK2 itself could be modeled as having 2/3 UNHG and 1/3 EHG ancestry, regardless of the source of the hunter-gatherer ancestry (UNHG or GK2) it would include a substantial amount of UNHG ancestry as would be expected, indeed, from the distribution of the Serednii Stih culture in the Dnipro-Don area where the Core Yamna itself must have emerged.

In ref.[3], we modeled Dnipro-Don populations as mixtures of Core Yamna (one end of the Dnipro-Don cline) and both UNHG and GK2 ancestries. What of the origin of the Core Yamna itself, which forms one end of the Dnipro-Don cline? In ref.[3], we presented proximal models for the origin of that population (as a mixture of Remontnoye and SShi populations), as well as more distal models, including the Aknashen-Bpgroup-GK2 model just discussed which derives their ancestry from a mixture of Caucasus-Lower Volga (CLV) cline people (Aknashen+Bpgroup) with Dnipro-Don hunter-gatherers. We would, however, want to investigate whether GK2 itself suffices as a hunter-gatherer source for the Core Yamna or the source of that ancestry may be slightly different from it along the UNHG-GK2-EHG cline.

We want to test the model Aknashen-Bpgroup-GK2-UNHG; while this model does fit the core Yamna (p=0.862), it can be estimated only with very high standard errors of 27.9±23.5% GK2 and -4.5±18.8% UNHG ancestry using the Base set. Using the Base1 set the model still fits (p=0.181) but standard errors are not improved (-12.7±78.9% UNHG ancestry!).

We thus attempt to find for the UNHG as a 4$^{th}$ source, in the same manner as we did for Trypillians as a 4$^{th}$ source, another outgroup which, when added to the Right might bring down standard errors. The top-20 populations with the smaller standard error are shown in Table S 41. Not surprisingly, Ukrainian Neolithic populations and individuals from this study and ref.[16,20,28] provide most traction towards the goal of lowering standard errors. In addition, Iron Gates Mesolithic from Romania, as well as diverse members of the "Oberkassel Cluster" (WHG-related populations of central-northern Europe identified in ref.[20]), are informative. The standard errors remain fairly high although we do observe that in general the proportion of GK2 ancestry is higher than UNHG ancestry and the latter is often negative and occasionally positive by small amounts.

| | | Proportions | | | | Std. errors | | | | RMSE |
|---|---|---|---|---|---|---|---|---|---|---|
| **Added outgroup** | **P-value** | **Aknashen** | **BPgroup** | **GK2** | **UNHG** | **Aknashen** | **BPgroup** | **GK2** | **UNHG** | |
| France_Mesolithic_1d.rel.PER503.SG | 0.891 | 21.2% | 55.6% | 26.0% | -2.8% | 2.1% | 4.8% | 12.1% | 9.5% | 8.1% |
| Spain_Canes.SG | 0.889 | 21.5% | 54.9% | 28.4% | -4.8% | 1.9% | 4.5% | 11.5% | 9.1% | 7.7% |
| Germany_Niedertiefenbach_Wartberg_LN.SG_brother.KH150622.SG.KH150620.SG | 0.914 | 21.6% | 54.8% | 28.9% | -5.3% | 2.1% | 4.4% | 11.3% | 9.2% | 7.7% |
| Russia_AfontovaGora2.SG | 0.910 | 21.5% | 55.0% | 28.4% | -5.0% | 2.0% | 4.3% | 11.3% | 9.2% | 7.7% |
| Russia_Ekaterinovka_Eneolithic_mother.I6068 | 0.877 | 21.4% | 55.4% | 26.1% | -3.0% | 2.2% | 4.3% | 10.7% | 8.9% | 7.4% |
| MesoFr | 0.887 | 21.4% | 55.3% | 27.2% | -3.9% | 2.0% | 4.5% | 10.9% | 8.5% | 7.3% |
| Latvia_MN_o3 | 0.886 | 21.5% | 55.4% | 26.8% | -3.6% | 2.1% | 4.2% | 10.6% | 8.9% | 7.3% |
| OberkasselCluster | 0.788 | 20.9% | 55.8% | 24.7% | -1.4% | 1.7% | 4.3% | 10.5% | 7.9% | 7.0% |
| ukr161 | 0.858 | 21.1% | 55.6% | 26.1% | -2.8% | 1.8% | 4.4% | 10.5% | 7.8% | 7.0% |
| Romania_IronGates_Mesolithic.SG | 0.889 | 21.5% | 54.9% | 28.3% | -4.8% | 1.8% | 4.3% | 10.3% | 7.9% | 6.9% |
| VO | 0.836 | 20.9% | 56.0% | 24.6% | -1.5% | 1.6% | 3.9% | 9.9% | 7.6% | 6.6% |
| Romania_IronGates_Mesolithic | 0.856 | 21.2% | 55.6% | 26.1% | -2.9% | 1.7% | 4.1% | 9.8% | 7.3% | 6.5% |
| Ukraine_N_1d.rel.I4114 | 0.834 | 21.2% | 55.6% | 26.3% | -3.0% | 1.8% | 3.9% | 9.6% | 7.6% | 6.5% |
| Ukraine_N_1d.rel.I5881 | 0.872 | 21.5% | 55.0% | 28.8% | -5.3% | 1.7% | 3.8% | 8.5% | 6.5% | 5.7% |
| Ukraine_N_father.or.son.I3718 | 0.838 | 21.4% | 55.2% | 27.4% | -4.0% | 1.7% | 3.8% | 8.2% | 6.3% | 5.6% |
| ukr162 | 0.580 | 20.2% | 57.6% | 19.1% | 3.1% | 1.5% | 3.4% | 8.1% | 6.2% | 5.4% |
| Ukraine_N_son.I1732 | 0.832 | 21.0% | 56.0% | 24.8% | -1.8% | 1.6% | 3.6% | 7.6% | 5.7% | 5.1% |
| ukr113 | 0.452 | 19.9% | 58.0% | 17.2% | 4.9% | 1.4% | 3.5% | 7.3% | 5.3% | 4.9% |
| Ukraine_N_dup.I4112 | 0.803 | 20.7% | 56.5% | 22.7% | 0.1% | 1.4% | 3.3% | 6.9% | 5.1% | 4.6% |
| ukr117 | 0.875 | 21.7% | 54.5% | 30.5% | -6.8% | 1.6% | 3.4% | 6.5% | 5.0% | 4.5% |

**Table S 41 Adding populations to Base set.** We add populations to the Right and examine the standard errors. Top-20 populations with lower standard errors are shown.

Conversely, we can test the model Aknashen-BPgroup-GK2-EHG, i.e., placing the hunter-gatherer source on the other side of GK2 along the UNHG-GK2-EHG cline. This model *can* be estimated fairly accurately with just the Base set (p=0.869) with 21.5±3.8% GK2 and 1.3±4.3% Russia_Karelia ancestry. The nominally positive proportion of EHG ancestry is not significantly different from zero as was the nominally negative proportion of UNHG ancestry in the previously considered model. Thus, significant deviations from GK2 towards either the UNHG or EHG side are supported by the data.

Finally, an alternative approach to placing the hunter-gatherer ancestry is to fit the Aknashen-BPgroup-UNHG-EHG model (using Russia_Karelia as the EHG source), i.e., do not use GK2 source but infer the position of the hunter-gatherer ancestry along the UNHG-EHG continuum. This model can be well fit and accurately estimated with just the Base set (p=0.759) with an estimated 13.0±1.9% UNHG and 8.4±2.7% EHG ancestry (thus the ratio of UNHG/(UNHG+EHG) is ~61%). GK2 itself can be fit as 66.6±4.6% UNHG and 33.4±4.6% EHG (p=0.556), or the aforementioned ~2/3 ratio. Thus, on the basis of this analysis it appears that GK2 is a reasonable approximation for the hunter-gatherer ancestry in the Core Yamna, although slight deviations to either the UNHG or EHG direction cannot be excluded from the data.

**Appendix V: European farmer ancestry in the Ukraine Neolithic hunter-gatherers**

We wanted to investigate the origins of Ukraine_N (UNHG) for two reasons.

The first, is the observation that in PCA (Fig. S 1), the direction of the EHG-UNHG-GK2 cline does not, when further projected tend towards the hunter-gatherers of the Balkans (BHG of Serbia Iron Gates) but is rather shifted "downwards", or, towards populations with farmer ancestry in the context of that PCA.

The second observation is that when we modeled Ukraine_N in the same setup as other populations of this note we can find only a single 2-source model that works (p=0.576) with 72.5±2.9% GK2 and 27.5±2.9% BHG ancestry. If Ukraine_N was part of a broader cline between EHG and BHG then it is surprising that it cannot also be modeled as a mixture of these two sources. However, using either Lebyazhinka or Karelia as the EHG source, this is not possible (p<1e-9) and qpAdm output suggests that these models underestimate shared genetic drift with Turkey_N (Z<-3.5). This is consistent with the shift in PCA towards farmers.

We thus explored 3-source models using all the same Sources as for other populations of this note, excluding only GK2: as the GK2+BHG model is feasible, there are many other generalized models of the form GK2+BHG+X with ~0% of X ancestry that are also feasible and our aim is to understand the position of Ukraine_N in the broader west-east/BHG-EHG continuum.

The feasible models (Table S 42) all include EHG-BHG sources (Lebyazhinka and Serbia Iron Gates), but they all include ~7-9% of farmer ancestry, with the source of this ancestry being unclear. Competition between the models of Table S 42 was inconclusive. We note that p-values for the European sources are higher than for the two Caucasus ones (Aknashen and Maykop) and this cannot be attributed to lower power to reject models as European sources all have larger sample sizes than those from the Caucasus.

| | | | | Proportions | | | Std. errors | | |
|---|---|---|---|---|---|---|---|---|---|
| A | B | C | P-value | A | B | C | A | B | C |
| Armenia_Aknashen_N | Lebyazhinka_HG | Serbia_IronGates_Mesolithic | 0.128 | 7.9% | 44.7% | 47.4% | 1.1% | 1.7% | 1.5% |
| GlobularAmphora | Lebyazhinka_HG | Serbia_IronGates_Mesolithic | 0.133 | 8.8% | 48.6% | 42.5% | 1.2% | 1.6% | 1.8% |
| Lebyazhinka_HG | Maykop | Serbia_IronGates_Mesolithic | 0.069 | 44.4% | 8.4% | 47.2% | 1.7% | 1.1% | 1.4% |
| Lebyazhinka_HG | Serbia_IronGates_Mesolithic | Trypillia | 0.257 | 47.8% | 44.0% | 8.2% | 1.7% | 1.7% | 1.2% |
| Lebyazhinka_HG | Serbia_IronGates_Mesolithic | YUN_CA | 0.198 | 48.2% | 45.0% | 6.8% | 1.6% | 1.6% | 1.0% |
| *LBK* | *Lebyazhinka_HG* | *Serbia_IronGates_Mesolithic* | *0.154* | *6.6%* | *48.7%* | *44.7%* | *1.0%* | *1.6%* | *1.7%* |

**Table S 42 3-source models for Ukraine_N.** All sources other than GK2 were considered. We manually tested the LBK+Lebyazhinka+Serbia Iron Gates model and include it in this table.

Given the sampling of Ukraine_N in the Dnipro area in some proximity to European farmer communities to its west, we think it likely that this 6$^{th}$ millennium BCE population was most probably interacted with European rather than Caucasus Neolithic populations.

We wanted to investigate whether the inferred farmer ancestry in Ukraine_N could be driven by individuals within that population as opposed to being a general feature of the population. Thus, we fit a model that included farmer ancestry to all 35 individuals under the Ukraine_N label, as well as relatives not included in the Ukraine_N label and outliers. We show this model with LBK (which is temporally more plausible, albeit genetically quite similar, to the later farmer groups, and also has the largest sample size) as the farmer source in Extended Data Table 2.

It is clear from Extended Data Table 2 individuals from Ukraine_N population have European farmer ancestry in general. An affinity to "Anatolian Neolithic farmer (ANF)" (and hence to the similar European farmers) was mentioned in ref.[20] but was said to include also Balkan hunter-gatherers, Baltic hunter-gatherers and Scandinavian hunter-gatherers.

We noticed that individual I31730 (Mariupol; 5474-5236 calBCE) does not fit the 3-source model (p=0.0037) which underestimates its shared genetic drift with CHG (Z=-2.9). Using CHG instead of LBK as a source, results in a successful fit (p=0.093) with an estimated 7.4±2.8% CHG-related ancestry. We also applied this alternative model to all individuals of Extended Data Table 2 that do not fit the model with LBK as a source (bottom of Extended Data Table 2) and discovered that one other individual (I1738; Vovnyhy, 5475-5320 calBC) could be modeled in such a way. Thus, while we can conclude that low amounts of European farmer ancestry entered the UNHG population (from the western neighbors of the NPR hunter-gatherers), it is possible that for at least some of them there was CHG-related ancestry as well (from the east). Such ancestry was also detected in the GK1 subset at Golubaya Krinitsa Neolithic in the Middle Don and at the Krivyansky Eneolithic (ref.[3]) in the Lower Don and may have thus extended further west into the Dnipro region.

Modeling the ancestry of European hunter-gatherers as a whole is beyond the scope of this note, but we observe that the farmer ancestry in the Ukraine_N is inferred with Balkan hunter-gatherers as one of the sources and therefore exceeds any that may have existed in that population. As for the northern groups, neither has any discernible LBK ancestry (the likely population transmitting such ancestry to northern Europe) with neither Motala[2,11,36] (0.3±1.2%), Stora Förvar (1.0±1.3%), or Latvia_HG (2.0±0.8%) having much evidence for it when the same model we applied to Ukraine_N is used (the true history for these northern populations probably involved native sources of WHG-related ancestry rather than the Balkan hunter-gatherers of the Iron Gates). As a control, we used the population from Ajvide in Sweden[42,43], a population of the Pitted Ware culture, which has a minority farmer ancestry component inferred to be 18.9±1.4% LBK ancestry, similar to the 22.1±1.1% inferred for another Pitted Ware population from Västerbjers.[44]

Thus, it appears that Ukraine Neolithic hunter-gatherers, similar to those of the Pitted Ware, incorporated some ancestry from their farmer neighbors while maintaining their overall hunter-gatherer genetic profile. Thus, while our investigation of the presence of European farmer ancestry in some populations like the core Yamna has failed to uncover some ancestry *on top of* the Ukraine Neolithic hunter-gatherer ancestry, some such ancestry may also have entered their population indirectly via populations with UNHG ancestry who had themselves European farmer ancestry.

**References**

1 Reich, D. *et al.* Reconstructing Native American population history. *Nature* **488**, 370-374, doi:10.1038/nature11258 (2012).

2 Haak, W. *et al.* Massive migration from the steppe was a source for Indo-European languages in Europe. *Nature* **522**, 207-211, doi:10.1038/nature14317 (2015).

3 Lazaridis, I., Patterson, N., Anthony, D. & others. The Genetic Origin of the Indo-Europeans. *in submission* (2024).

4 Skoglund, P. *et al.* Reconstructing Prehistoric African Population Structure. *Cell* **171**, 59-71.e21, doi:10.1016/j.cell.2017.08.049 (2017).

5 Wang, K. *et al.* Ancient genomes reveal complex patterns of population movement, interaction, and replacement in sub-Saharan Africa. *Science Advances* **6**, eaaz0183, doi:10.1126/sciadv.aaz0183.

6 Lipson, M. *et al.* Ancient DNA and deep population structure in sub-Saharan African foragers. *Nature* **603**, 290-296, doi:10.1038/s41586-022-04430-9 (2022).

7 Fu, Q. *et al.* The genetic history of Ice Age Europe. *Nature* **534**, 200-205, doi:10.1038/nature17993 (2016).

8 Jones, E. R. *et al.* Upper Palaeolithic genomes reveal deep roots of modern Eurasians. *Nat Commun* **6**, 8912, doi:10.1038/ncomms9912 (2015).

9 Lazaridis, I. *et al.* Genomic insights into the origin of farming in the ancient Near East. *Nature* **536**, 419-424, doi:10.1038/nature19310 (2016).

10 de Barros Damgaard, P. *et al.* The first horse herders and the impact of early Bronze Age steppe expansions into Asia. *Science* **360**, doi:10.1126/science.aar7711 (2018).

11 Mathieson, I. *et al.* Genome-wide patterns of selection in 230 ancient Eurasians. *Nature* **528**, 499-503, doi:10.1038/nature16152 (2015).

12 Lazaridis, I. *et al.* The genetic history of the Southern Arc: A bridge between West Asia and Europe. *Science* **377**, eabm4247, doi:10.1126/science.abm4247 (2022).

13 Wang, C.-C. *et al.* Ancient human genome-wide data from a 3000-year interval in the Caucasus corresponds with eco-geographic regions. *Nature Communications* **10**, 590, doi:10.1038/s41467-018-08220-8 (2019).

14 Allentoft, M. E. *et al.* Population genomics of Bronze Age Eurasia. *Nature* **522**, 167-172, doi:10.1038/nature14507 (2015).

15 Kumar, V. *et al.* Bronze and Iron Age population movements underlie Xinjiang population history. *Science* **376**, 62-69, doi:10.1126/science.abk1534 (2022).

16 Mathieson, I. *et al.* The genomic history of southeastern Europe. *Nature* **555**, 197-203, doi:10.1038/nature25778 (2018).

17 Narasimhan Vagheesh, M. *et al.* The formation of human populations in South and Central Asia. *Science* **365**, eaat7487, doi:10.1126/science.aat7487 (2019).

18 Patterson, N. *et al.* Large-scale migration into Britain during the Middle to Late Bronze Age. *Nature* **601**, 588-594, doi:10.1038/s41586-021-04287-4 (2022).

19 Mittnik, A. *et al.* Kinship-based social inequality in Bronze Age Europe. *Science* **366**, 731-734, doi:10.1126/science.aax6219 (2019).

20 Posth, C. *et al.* Palaeogenomics of Upper Palaeolithic to Neolithic European hunter-gatherers. *Nature* **615**, 117-126, doi:10.1038/s41586-023-05726-0 (2023).

21 Gelabert, P. *et al.* Genomes from Verteba cave suggest diversity within the Trypillians in Ukraine. *Scientific Reports* **12**, 7242, doi:10.1038/s41598-022-11117-8 (2022).

22 Penske, S. *et al.* Early contact between late farming and pastoralist societies in southeastern Europe. *Nature* **620**, 358-365, doi:10.1038/s41586-023-06334-8 (2023).

23 Patterson, N., Price, A. L. & Reich, D. Population Structure and Eigenanalysis. *PLOS Genetics* **2**, e190, doi:10.1371/journal.pgen.0020190 (2006).

24 Shinde, V. *et al.* An Ancient Harappan Genome Lacks Ancestry from Steppe Pastoralists or Iranian Farmers. *Cell* **179**, doi:10.1016/j.cell.2019.08.048 (2019).

25 Harney, É. *et al.* Ancient DNA from Chalcolithic Israel reveals the role of population mixture in cultural transformation. *Nature Communications* **9**, 3336, doi:10.1038/s41467-018-05649-9 (2018).

26 Lipson, M. *et al.* Parallel palaeogenomic transects reveal complex genetic history of early European farmers. *Nature* **551**, 368-372, doi:10.1038/nature24476 (2017).

27 Rivollat, M. *et al.* Ancient genome-wide DNA from France highlights the complexity of interactions between Mesolithic hunter-gatherers and Neolithic farmers. *Science Advances* **6**, eaaz5344, doi:10.1126/sciadv.aaz5344 (2020).

28 Mattila, T. M. *et al.* Genetic continuity, isolation, and gene flow in Stone Age Central and Eastern Europe. *Communications Biology* **6**, 793, doi:10.1038/s42003-023-05131-3 (2023).

29 Patterson, N. *et al.* Ancient Admixture in Human History. *Genetics* **192**, 1065-1093, doi:10.1534/genetics.112.145037 (2012).

30 Harald, R. *et al.* ancIBD - Screening for identity by descent segments in human ancient DNA. *bioRxiv*, 2023.2003.2008.531671, doi:10.1101/2023.03.08.531671 (2023).

31 Saag, L. *et al.* Genetic ancestry changes in Stone to Bronze Age transition in the East European plain. *Science Advances* **7**, eabd6535, doi:10.1126/sciadv.abd6535 (2021).

32 Papac, L. *et al.* Dynamic changes in genomic and social structures in third millennium BCE central Europe. *Science Advances* **7**, eabi6941, doi:doi:10.1126/sciadv.abi6941 (2021).

33 Underhill, P. A. *et al.* The phylogenetic and geographic structure of Y-chromosome haplogroup R1a. *European Journal of Human Genetics* **23**, 124-131, doi:10.1038/ejhg.2014.50 (2015).

34 González-Fortes, G. *et al.* Paleogenomic Evidence for Multi-generational Mixing between Neolithic Farmers and Mesolithic Hunter-Gatherers in the Lower Danube Basin. *Current Biology* **27**, 1801-1810.e1810, doi:10.1016/j.cub.2017.05.023 (2017).

35 Hofmanová, Z. *et al.* Early farmers from across Europe directly descended from Neolithic Aegeans. *Proceedings of the National Academy of Sciences* **113**, 6886-6891, doi:10.1073/pnas.1523951113 (2016).

36 Lazaridis, I. *et al.* Ancient human genomes suggest three ancestral populations for present-day Europeans. *Nature* **513**, 409-413, doi:10.1038/nature13673 (2014).

37 Lazaridis, I. *et al.* Ancient DNA from Mesopotamia suggests distinct Pre-Pottery and Pottery Neolithic migrations into Anatolia. *Science* **377**, 982-987, doi:10.1126/science.abq0762 (2022).

38 Lazaridis, I. *et al.* Genetic origins of the Minoans and Mycenaeans. *Nature* **548**, 214-218, doi:10.1038/nature23310 (2017).

39 Chintalapati, M., Patterson, N. & Moorjani, P. The spatiotemporal patterns of major human admixture events during the European Holocene. *eLife* **11**, e77625, doi:10.7554/eLife.77625 (2022).

40 Fenner, J. N. Cross-cultural estimation of the human generation interval for use in genetics-based population divergence studies. *Am J Phys Anthropol* **128**, 415-423, doi:10.1002/ajpa.20188 (2005).

41 Zhang, F. *et al.* The genomic origins of the Bronze Age Tarim Basin mummies. *Nature* **599**, 256-261, doi:10.1038/s41586-021-04052-7 (2021).

42 Skoglund, P. *et al.* Genomic diversity and admixture differs for Stone-Age Scandinavian foragers and farmers. *Science* **344**, 747-750, doi:10.1126/science.1253448 (2014).

43 Malmström, H. *et al.* The genomic ancestry of the Scandinavian Battle Axe Culture people and their relation to the broader Corded Ware horizon. *Proc Biol Sci* **286**, 20191528, doi:10.1098/rspb.2019.1528 (2019).

44 Coutinho, A. *et al.* The Neolithic Pitted Ware culture foragers were culturally but not genetically influenced by the Battle Axe culture herders. *American Journal of Physical Anthropology* **172**, 638-649, doi:https://doi.org/10.1002/ajpa.24079 (2020).

### A genomic history of the North Pontic Region from the Neolithic to the Bronze Age

***Supplementary File S3***

#### Unsupervised admixture modeling with summary individuals

Practitioners of ADMIXTURE analysis[1] know that as the number of ancestral populations (*K*) increases the algorithm often arrives at solutions in which a population of interest forms its own ancestral population. For example, the Kalash, a highly drifted genetic isolate from Pakistan form such a component[2] and are inferred to derive the majority of their ancestry from it with other populations deriving ancestry from this component.

This behavior is problematic for two reasons. First, the directionality of ancestry may be reversed; a highly drifted population may have the same types of ancestry as its neighbors but is shown to be "unadmixed" and contributing ancestry *to* them, rather than being admixed themselves. Second, while at lower *K* (before the emergence of the population-specific component) it may be possible to see what the ancestry of the highly drifted population is in terms of components shared with others, this ability is lost as soon as the population-specific cluster is inferred.

Thus, ADMIXTURE analysis relies on an element of "luck". The practitioner hopes that, for each population of interest, a population-specific component will *not* appear at a *K* value lower than the one for which all informative components of its ancestry have had the opportunity themselves to appear.

As a practical example, we show in Fig. S 1 an ADMIXTURE analysis of a set of populations that include diverse hunter-gatherers and Eneolithic/Bronze Age populations representing ancestry of the Pontic-Caspian steppe and of the Near East and Caucasus (from this study and ref.[3]). At *K*=2 populations of European hunter-gatherers (orange) are differentiated from those of the Near East (blue; both Anatolia and Iran/Caucasus). At *K*=3 European hunter-gatherers are split between Balkan hunter-gatherers (orange) and Eastern hunter-gatherers (yellow). The Core Yamna have divided affinities at these first *K*=2, 3 consistent with admixed origins of "north" (European hunter-gatherer) and "south" (Near Eastern) components (*K*=2), and their European component is predominantly EHG and not BHG (*K*=3).

At *K*=4, however, a population component (pink) specific to the Core Yamna appears, and this appears to contribute to much older and remote populations, such as Neolithic Iran, Caucasus hunter-gatherers and the Eneolithic people of both Khvalynsk and Serednii Stih. Clearly, we have a case here in which the pink "Yamna"-centered component is spurious; the algorithm set it to maximum global likelihood because there is a large number of homogeneous Yamna individuals in the dataset. Notice also the conflicting information regarding the hunter-gatherer admixture of the Core Yamna suggested by *K*=3 and *K*=4. The hunter-gatherer component at *K*=3 is almost entirely EHG (yellow); this ancestry is "absorbed" by the Yamna-specific component at *K*=4 which is then shown to require some extra BHG (orange) ancestry.

At *K*=5, an informative component does appear: Near Eastern populations, previously represented by the “blue” component are now split to Anatolian-related (blue) and Caucasus/Iran-related (green). It would be great if we could assess the ancestry of the Core Yamna in terms of these components, but we are unable to do so as the Yamna at *K*=5 continue to belong virtually completely from their population-specific “pink” component, which becomes even more specific to them as it swallows up the all the BHG-related ancestry of the preceding *K*=4 run and many Yamna individuals belong ~100% to it.

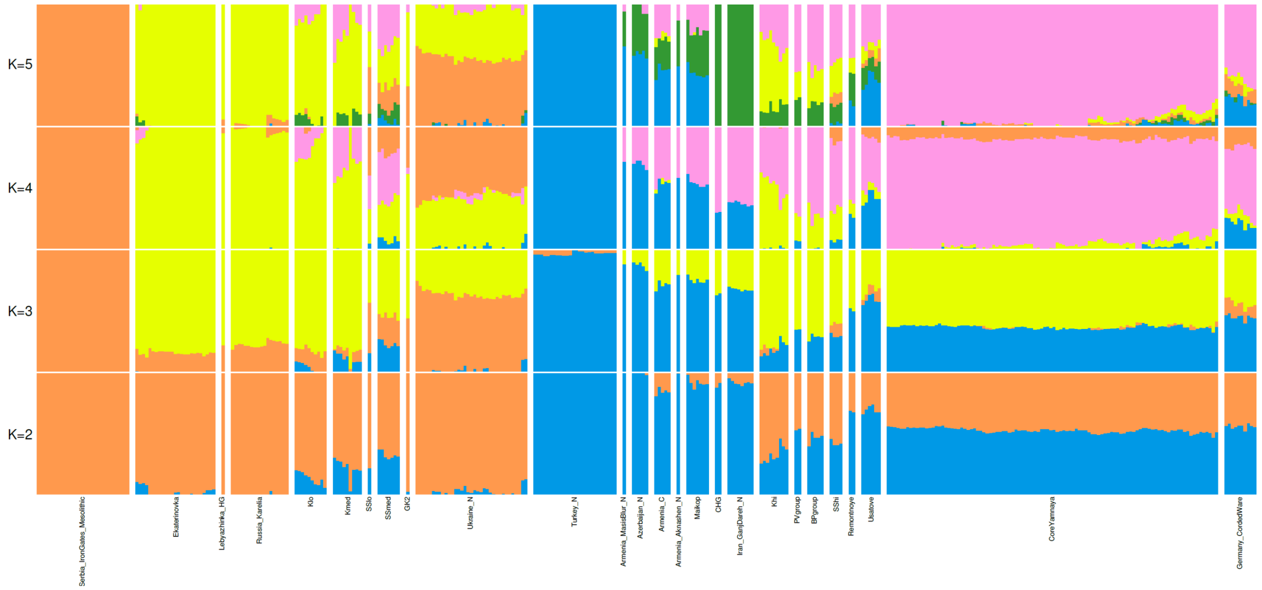

**Fig. S 1 Unsupervised ADMIXTURE analysis.**

A way to overcome this problem is to use a *single* individual from the entire population to represent it. We may create such an individual in different ways; a simple method (which we adopt here) is to simply take all individuals belonging to it and pick an allele at random for each genomic position creating a single pseudo-haploid individual. Another possible approach is to take a single high-quality individual from the population; the drawbacks of this method is that (a) it has slightly less data than using all the individuals, and (b) we are at some risk of picking an individual that systematically differs in some way from the population as a whole. A third approach addresses concern (b) by iterating over *all* individuals of the population.

Supervised ADMIXTURE

A possible objection to the procedure suggested in the previous paragraph is to use *supervised* ADMIXTURE analysis instead. There are a few drawbacks to that approach. First, it requires one to posit a priori what the source populations are. Second, any inhomogeneities in the source populations cannot be addressed: if a population has ancestral variation then by using it as a source in supervised ADMIXTURE then all its individuals are assigned 100% ancestry from its own source; there is no latitude to allow this proportion to vary to cover this variation. Third, it uses information only from the limited number postulated ancestral populations and cannot use data from other populations composed of the same kinds of ancestry.

Fourth, in our practice we notice that the numerical convergence of supervised ADMIXTURE is often to solutions as shown in Fig. S 2 in which we use the same dataset as in Fig. S 1 but use CHG, Turkey_N,

Russia_Karelia, and Serbia_IronGates_Mesolithic as the four sources in a supervised analysis. This solution shows the Core Yamna to have 84.9% (on average) of the CHG component which is much higher amount of "southern" ancestry inferred with either qpAdm or with the unsupervised ADMIXTURE of Fig. S 1 (37-53% in $K$=2, 3, prior to the appearance of the Yamna-centered component). Bias of supervised ADMIXTURE estimates in a different setting was also observed in ref.[4]

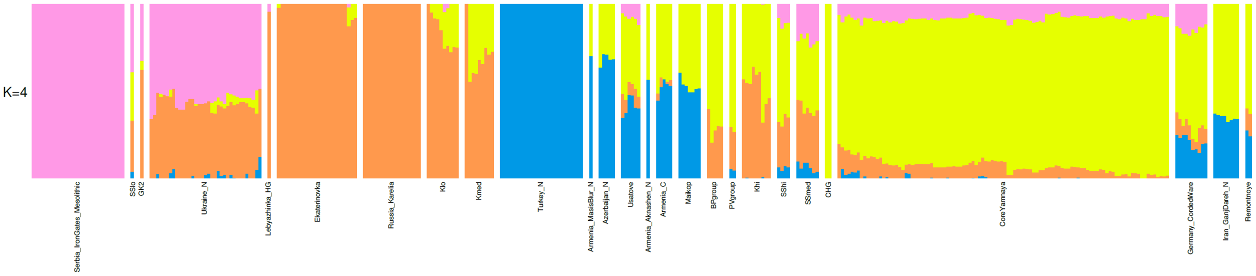

**Fig. S 2 Supervised ADMIXTURE analysis.**

Projection ADMIXTURE

ADMIXTURE can also be run in *projection mode*. This uses the allele frequency (*.P) files of an unsupervised ADMIXTURE analysis on one dataset to estimate the ancestry proportions of a second (test) dataset. An advantage of this method over supervised ADMIXTURE is that it uses more individuals than a pre-set number of source populations and makes no assumptions about their homogeneity.

Its main disadvantage is that the individuals used in the initial unsupervised analysis contribute to the allele frequencies, while the test individuals are treated differently (and do not contribute to the allele frequencies). This leaves room for potential bias, as the test individuals are not treated in the same manner as the ones used to compute the allele frequencies. Another potential problem is that the test individuals might have ancestry not represented in the unsupervised ADMIXTURE set; by projecting them using the allele frequencies computed from that set we are constraining them to be modeled using only the kinds of ancestry present there.

We used projection ADMIXTURE as follows: first, we computed allele frequencies using unsupervised ADMIXTURE on the populations of Fig. S 1 (excluding the Core Yamna), and then used these to project all the populations (including the Core Yamna). The results are shown in Fig. S 3.

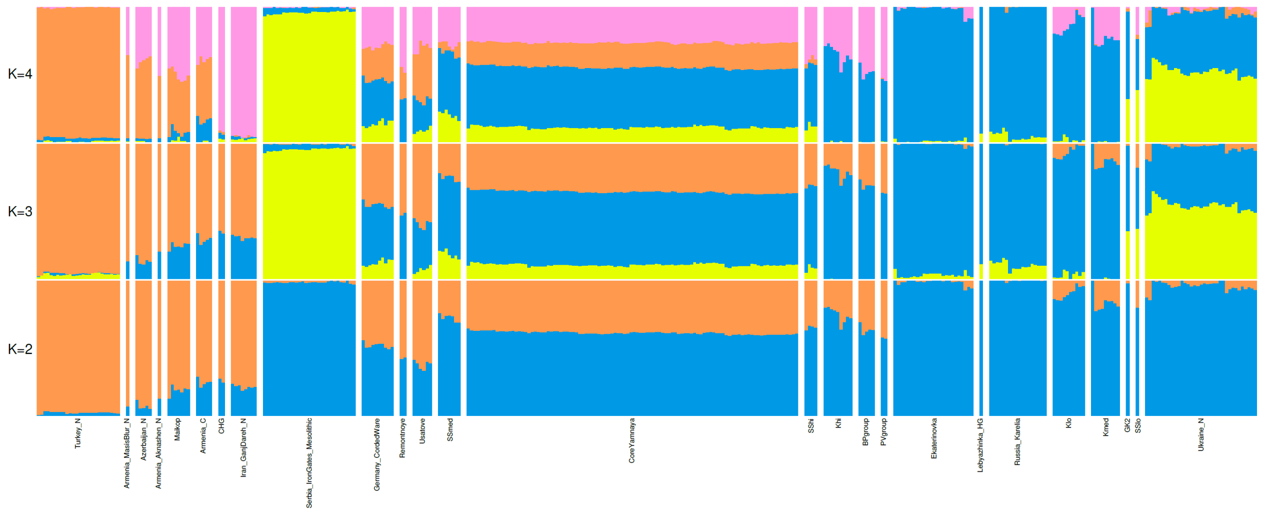

**Fig. S 3 Projection ADMIXTURE analysis.** Core Yamna are projected; other populations are used in an unsupervised ADMIXTURE analysis to compute allele frequencies.

<u>Unsupervised ADMIXTURE with a Core Yamna summary individual</u>

Thus, we adopt the single pseudo-haploid individual method as it does not have the concerns (a) or (b) of unsupervised ADMIXTURE or the potential bias problems of unsupervised/projection ADMIXTURE. The main drawback of the single pseudo-haploid individual method is that it collapses the population into a single individual, thus potentially masking any heterogeneity in it. However, we have validated the homogeneity of the Core Yamna in various ways (by the manner in which this population was assembled by showing that its high quality (at least 400K 1240k SNPs) individuals cluster with Samara Yamna in PCA and also are cladal with it to a high degree of certainty ($p \geq 0.2$) using qpWave[5]). Thus, we are confident that excessive heterogeneity or substantial outlier individuals do not exist among the 104 Core Yamna individuals.

The results of our method are shown in Fig. S 4. As is clear, there is no Yamna-specific cluster that appears in either $K$=3 or $K$=4 and thus we can assess the ancestry of the Core Yamna in terms of the other components. The averages of the populations of Fig. S 4 are shown in Table S 1.

<u>Comparison of unsupervised ADMIXTURE with qpAdm and with projection ADMIXTURE</u>

It is interesting to compare these inferred proportions with those inferred using qpAdm.[6] Recall that Core Yamna is modeled as either SShi+Remontnoye (SShi representing the Eneolithic Serednii Stih and Remontnoye the extra CLV ancestry) or (Aknashen+BPgroup)+GK2 (Aknashen+BPgroup representing the total CLV ancestry and GK2 the NPR hunter-gatherer ancestry). In Table S 2 we show this comparison which indicates that the two methods agree with each other within 0.3-3.1% for either of the qpAdm models and all $K$=4 components. This fairly close agreement provides independent confirmation using a non-$f$-statistics based method of the qpAdm reconstruction of the ancestry of the Core Yamna.

(Note that some slight differences may be inherent between the two methods as qpAdm is a supervised method in which the source populations are proposed a priori while in unsupervised ADMIXTURE they are discovered from the data and have imperfect correspondences to the sampled populations. In particular the EHG-centered component inferred by admixture is clear but may not be entirely corresponding to the sampled EHG as, for example, it is also dominated by the large sample size of individuals from Ekaterinovka which are inferred using qpAdm to have most (but not all) of their ancestry from EHG. Similarly, the CHG and Ganj Dareh Neolithic dominate the CHG-centered component, while it is known that these two populations are not identical to each other and CHG has more affinity to European hunter-gatherers.[7])

We include in Table S 2 the ADMIXTURE proportions obtained from projection ADMIXTURE (Fig. S 3). These differ from the unsupervised ADMIXTURE/qpAdm-inferred range substantially and especially for the Anatolian/CHG-related components, overestimating the former and underestimating the latter. So, while projection ADMIXTURE usefully shows the presence of all four components in the Core Yamna, there is reason to think that its different treatment of projected individuals introduces some bias.

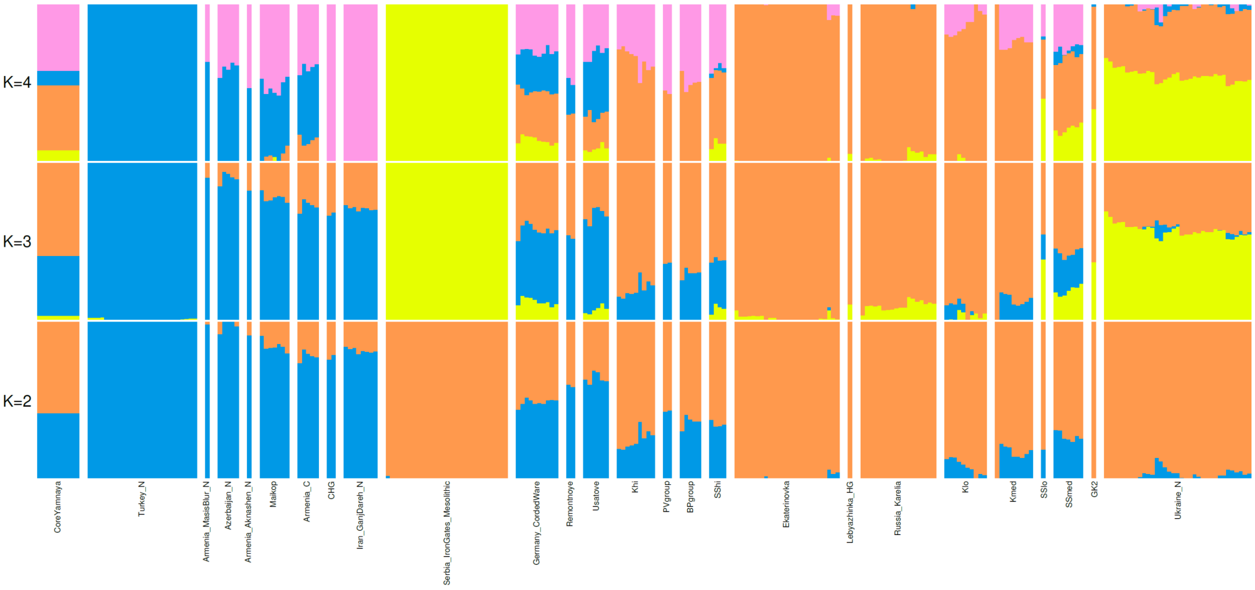

**Fig. S 4 Unsupervised ADMIXTURE with a single pseudo-haploid Core Yamna summary individual.** The summary individual is shown at the left side of the plot with a single wide bar.

| Population | *N* | Yellow BHG | Orange EHG | Blue Anatolian Neolithic | Pink CHG-Iran |
|---|---|---|---|---|---|
| Armenia_Aknashen_N | 1 | 0.0% | 0.0% | 46.8% | 53.2% |
| Armenia_C | 5 | 0.0% | 13.2% | 46.2% | 40.6% |
| Armenia_MasisBlur_N | 1 | 0.0% | 0.0% | 63.4% | 36.6% |
| Azerbaijan_N | 5 | 0.0% | 0.0% | 59.2% | 40.8% |
| BPgroup | 5 | 0.0% | 50.3% | 0.0% | 49.7% |
| CHG | 2 | 0.0% | 0.0% | 0.0% | 100.0% |
| CoreYamna | 1 | 6.8% | 41.7% | 9.2% | 42.3% |
| Ekaterinovka | 25 | 0.1% | 98.9% | 0.0% | 1.0% |
| Germany_CordedWare | 10 | 13.4% | 31.2% | 25.2% | 30.2% |
| GK2 | 1 | 33.1% | 65.6% | 1.4% | 0.0% |
| Iran_GanjDareh_N | 8 | 0.0% | 0.0% | 0.0% | 100.0% |
| Khi | 9 | 0.0% | 64.7% | 0.0% | 35.3% |
| Klo | 10 | 0.7% | 87.0% | 0.0% | 12.4% |
| Kmed | 9 | 0.0% | 78.0% | 0.0% | 22.0% |
| Lebyazhinka_HG | 1 | 4.6% | 95.4% | 0.0% | 0.0% |
| Maykop | 7 | 0.4% | 3.0% | 44.0% | 52.6% |
| PVgroup | 2 | 0.0% | 44.1% | 0.0% | 55.9% |
| Remontnoye | 2 | 0.0% | 30.1% | 20.9% | 49.0% |
| Russia_Karelia | 18 | 2.5% | 97.4% | 0.2% | 0.0% |
| Serbia_IronGates_Mesolithic | 29 | 100.0% | 0.0% | 0.0% | 0.0% |
| SShi | 4 | 11.1% | 45.4% | 2.9% | 40.6% |
| SSlo | 1 | 40.0% | 37.8% | 2.0% | 20.2% |

| | | | | | |
|---|---|---|---|---|---|
| SSmed | 7 | 20.6% | 46.0% | 5.5% | 27.9% |
| Turkey_N | 26 | 0.0% | 0.0% | 100.0% | 0.0% |
| Ukraine_N | 35 | 54.9% | 42.4% | 1.6% | 1.1% |
| Usatove | 6 | 8.1% | 21.2% | 39.5% | 31.2% |
| Armenia_Aknashen_N | 1 | 0.0% | 0.0% | 46.8% | 53.2% |

**Table S 1 Average proportions of populations of Fig. S 4**

| Population | BHG | EHG | Anatolian | CHG |
|---|---|---|---|---|
| CoreYamna | 6.8% | 41.7% | 9.2% | 42.3% |
| 0.263*Remontnoye+0.737*SShi | 8.2% | 41.4% | 7.6% | 42.8% |
| 0.207*Aknashen+0.568*BPgroup+0.225*GK2 | 7.4% | 43.3% | 10.0% | 39.2% |
| CoreYamna (projection ADMIXTURE) | 10.9% | 44.4% | 18.4% | 26.3% |

**Table S 2 Comparing ADMIXTURE and qpAdm admixture proportions.** We show the *K*=4 admixture proportions of Core Yamna from Table S 1 and the inferred *K*=4 proportions for the two alternative qpAdm models of Core Yamna.

Unsupervised ADMIXTURE analysis with summary individuals from diverse populations

We were encouraged by the preceding results to use the same method with summary individuals from populations other than those included in the unsupervised ADMIXTURE analysis. This would allow us to assess their ancestry in terms of the same model as we did for the Core Yamna summary individual. Thus, we repeated the ADMIXTURE analysis, creating summary individuals for each population of interest. The results are shown in Fig. S 5.

Naturally, the solution for some of the summary individuals may not be a valid representation of its ancestry. For example, when using OldAfrica or Han.DG as the test populations, this is coerced into the ancestral components generated by the West Eurasian individuals used in the ADMIXTURE analysis.

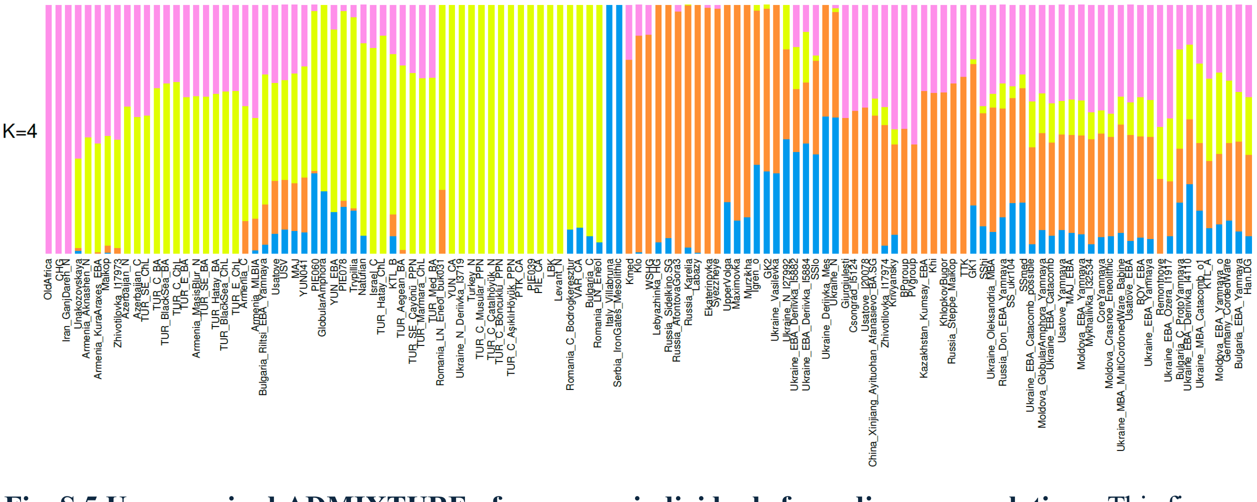

**Fig. S 5 Unsupervised ADMIXTURE of summary individuals from diverse populations.** This figure summarizes the results of Table S 1 as well as additional unsupervised ADMIXTURE runs in which the Core Yamna summary individual was replaced by a single summary individual for each of 97 additional populations.

Removing populations for which the 4-source model is inapplicable

To address the problem of whether individuals are correctly modeled in terms of the four ancestral components, we took the following approach. We performed three principal component analyses, including a "World PCA", a "West Eurasian" PCA[8] and a "Steppe" PCA (as in the main PCA of the paper). In each of these PCA we project the summary individuals for each population as well as summary individuals for the four admixture components; the latter can be generated by examining the *.P (allele frequency) output of ADMIXTURE and picking an allele at random for each position.

These results are shown for OldAfrica and Core Yamna in Fig. S 6. Note that in all these plots the ADMIXTURE components are represented by the run of Fig. S 4; for each ADMIXTURE run, the *.P files differ from each other, even if they converge to a similar solution. Each plot shows the "convex" (red) individual as a summary of the population given the estimated admixture proportions. It also shows the "summary" (blue) individual itself. Notice that in both "World" and "West Eurasian" PCA plots, the Old African "summary" individual is well beyond the "convex" one: this proves that the modeling of Old Africa in terms of the 4-source model (which is ~100% of the CHG-related component; Fig. S 5) is inadequate: Old Africa is squeezed into the components of this model even though in fact it has quite different ancestry. By contrast, the "convex" and "summary" individuals for the Core Yamna are in proximity to each other, and within the convex hull of the four components: the Core Yamna can indeed be modeled in terms of the four components and their modeling ("convex") is in proximity to their actual population ("summary").

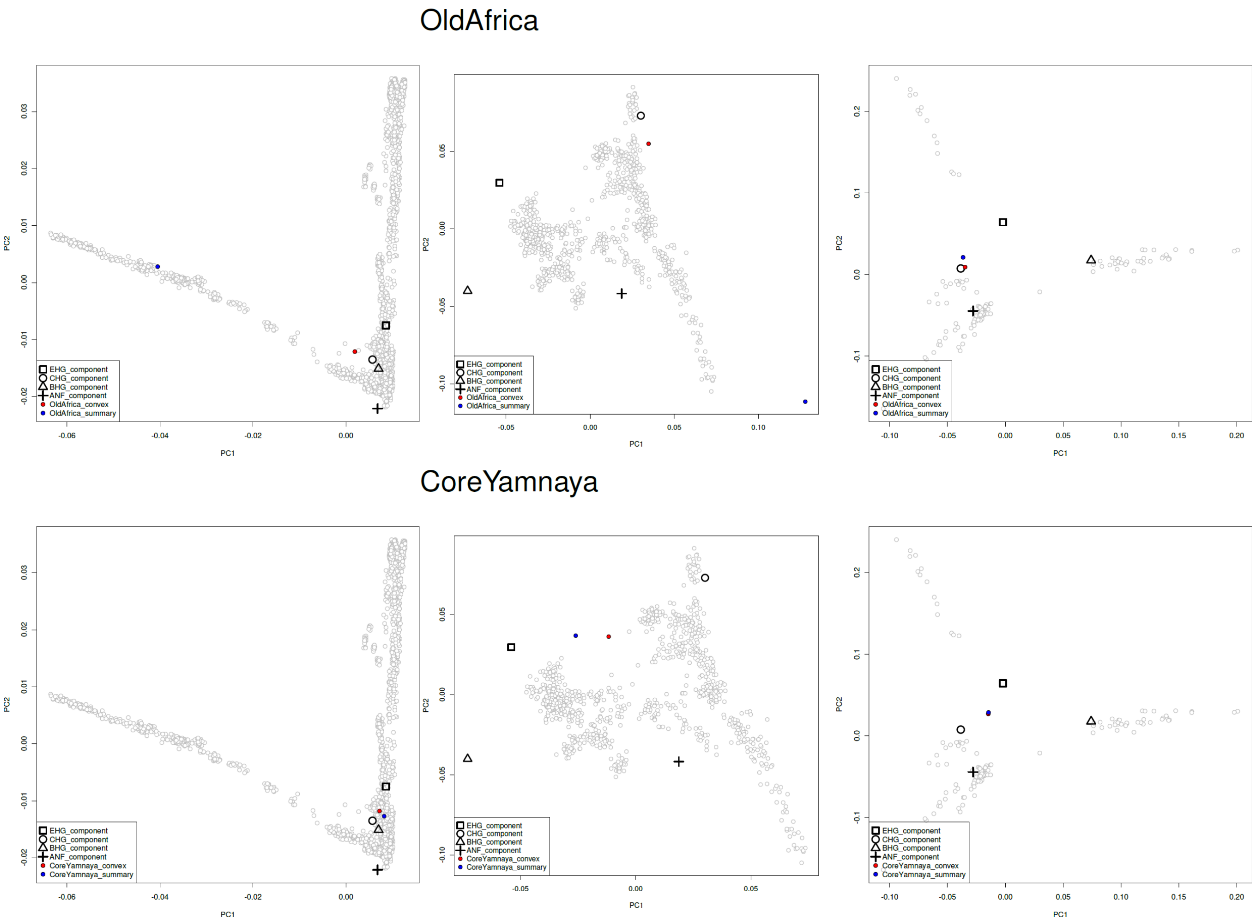

**Fig. S 6 Projection of summary individuals and ADMIXTURE components onto World (left), West Eurasian (middle), and Steppe (right) PCA.**

The approach we took here could be generalized to include other statistics; the basic idea is simple: we use the summary individual and the convex combination of the ADMIXTURE components to see if they agree with each other.

The three PCAs capture different aspects of genetic variation of the populations of interest and are a sufficient filter for our purposes of removing from Fig. S 5 those populations which cannot be well-modeled in terms of the four sources. This can be done either by examining the PCAs visually and making a subjective assessment or in an automated way. For “Old Africa” common sense and the visual assessment of Fig. S 6 are in agreement that this is a population outside the 4-source model; this may be less clear for other populations. We thus adopted an automated approach as follows.

First, we accept populations whose summary individual is *within* the convex hull of the four ancestral components. From the point of view of the three PCAs, the summary individual *can* be modeled as a convex combination of the four components.

This criterion may be too stringent, however, for populations that occupy the edges of the convex hull. For example, if a population is a BHG/EHG mixture in reality then its summary representation should be exactly on the BHG-EHG edge of the convex hull. However, because of stochastic variation it may, in fact be either within, or slightly outside the convex hull. An example of precisely such a population is Upper Volga (Fig. S 7).

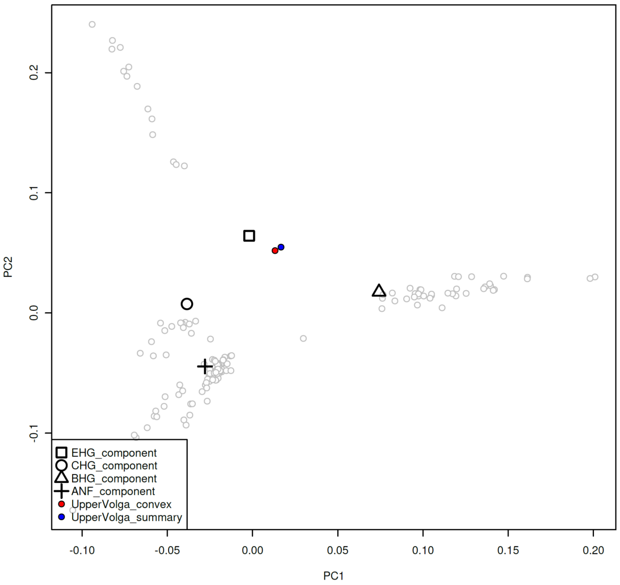

**Fig. S 7 Upper Volga in the "Steppe" PCA.** The summary and convex individuals of this population are close to each other, but the summary (blue) individual is slightly outside the convex hull of the four components.

We can address this issue empirically by examining the minimum distance of a summary individual to the convex hull. This is the minimum distance to the (four) edges of the convex hull. We adapted pseudo-code for this problem (https://stackoverflow.com/a/6853926) to R and can thus compute the minimum distance for all populations that fall outside the convex hull. We show these in Fig. S 8

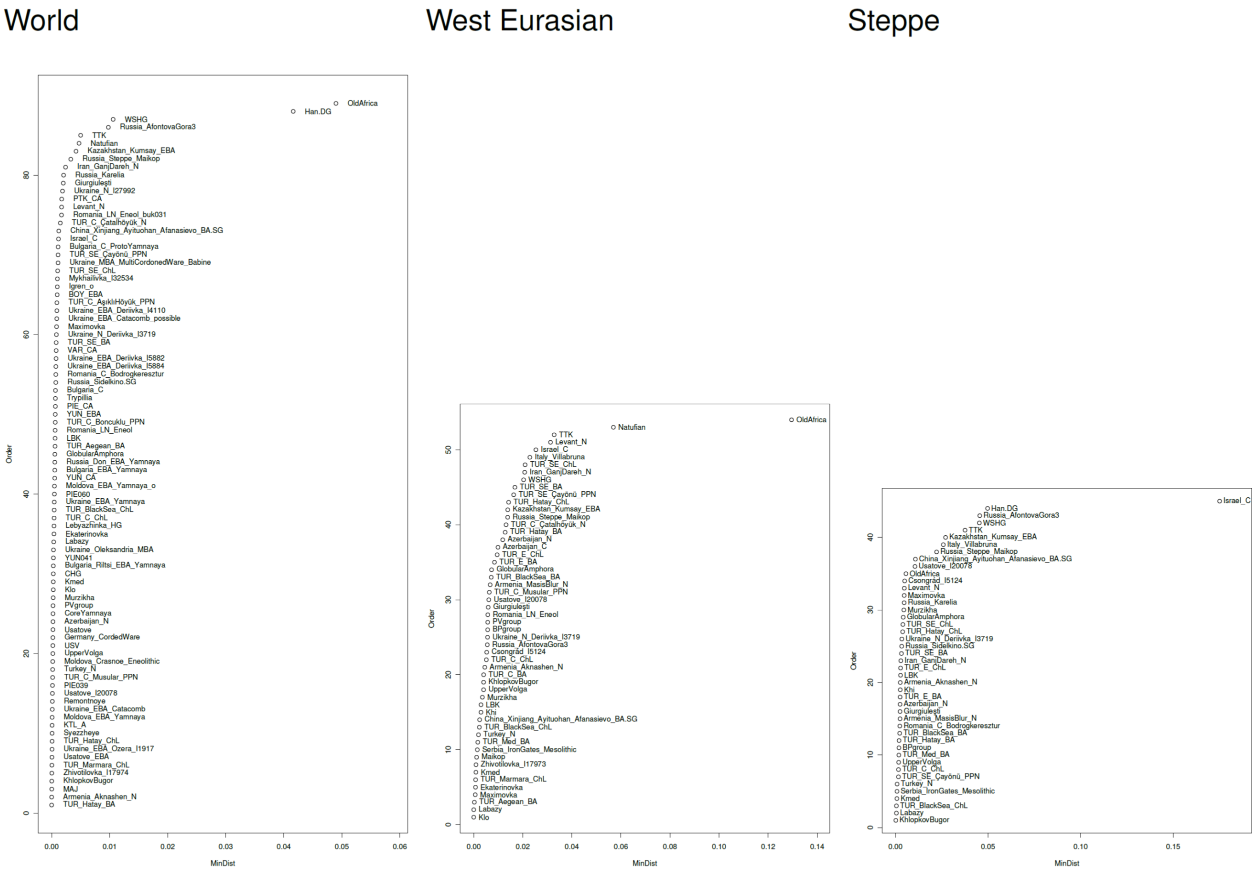

**Fig. S 8 Minimum distance of summary individuals to convex hull of four admixture components for the three PCAs.** The minimum distance of points that fall outside the convex hull, even by a small amount are shown.

The most egregious outliers are, as we might expect, Old Africa and Han Chinese in the World PCA. Natufians are a clear outlier in the West Eurasian PCA; this makes sense as the 4-source model does not have a Levantine source and is thus not suitable for ancient Levantine populations, of which the Epipaleolithic Natufians are most representative.[7,9] Israel_C is an outlier for the Steppe PCA; this too could be because of the Levantine ancestry of this population[10] but may also be a consequence of the fact that Israel_C is one of the populations used to form the axes of the Steppe PCA, in contrast to the "World" and "West Eurasian" PCAs that are formed on the basis of modern populations and are thus "neutral" with respected to all the ancient ones.

Other populations that are visibly at the top of the minimum distance plots include those of Siberia (such as Afontova Gora 3) or having ancestry related to it (such as the Neolithic TTK from Tajikistan[11] or the Steppe Maykop[12]).

Our final decision was to omit several populations aggressively that are in the top of Fig. S 8 and to keep those that form the undifferentiated set of low-MinDist populations at the bottom. The omitted populations are:

Azerbaijan_C, Azerbaijan_N, China_Xinjiang_Ayituohan_Afanasievo_BA.SG, Han.DG, Iran_GanjDareh_N, Israel_C, Italy_Villabruna, Kazakhstan_Kumsay_EBA, Levant_N, Natufian, OldAfrica, Russia_AfontovaGora3, Russia_Steppe_Maykop, TTK, TUR_C_Çatalhöyük_N, TUR_E_BA,

TUR_E_ChL, TUR_Hatay_BA, TUR_Hatay_ChL, TUR_SE_BA, TUR_SE_Çayönü_PPN, TUR_SE_ChL, Usatove_I20078, WSHG

Omitting these populations from Fig. S 5 we plot the remaining ones in Fig. S 9.

We highlight some contrasts in these results that correspond to the findings of our formal qpWave/qpAdm analysis.

1. The CLV Cline: its northern end (BPgroup) is a mix of CHG/EHG related components; its southern end (Aknashen) is a mix of CHG/Anatolian ancestry. Intermediate populations (Maykop and Remontnoye) have all three components.
2. The Volga Cline: Khvalynsk subsets (Khi, Klo, Kmed) and Ekaterinovka have more EHG ancestry than the southern end of the cline (BPgroup). Migrants from the Volga cline across the NPR at Giurgiuleşti and Csongrád represent a similar balance of components to the BPgroup and low-EHG end of the Volga Cline.
3. The Dnipro Cline: Serednii Stih subsets (SShi, SSlo, SSmed) and Yamna have some BHG-related ancestry not found in the CLV Cline, but found in NPR hunter-gatherers and their descendants such as Ukraine_N and GK2. Thus, the Dnipro Cline could be formed on the basis of CLV migrants admixing with NPR hunter-gatherers as our other analyses indicate.
4. The second major admixture process in the NPR was what formed the Usatove/USV/MAJ-related populations; these visibly differ from the Serednii Stih/Yamna in having much Anatolian-related ancestry.
5. The proximal source of the Anatolian-related ancestry in the Usatove could be the Trypillians; these also have a little CHG/EHG-related ancestry, consistent with our reconstruction that they absorbed some CLV migrants as well, but to a lesser degree than the Usatove
6. Populations of Armenia provide an interesting control for our reconstruction, paralleling what is known of the history of the country[13]. First, Aknashen/Masis Blur differ from each other in being more/less CHG-admixed and both lack any steppe influence. The Areni-1 population (Armenia_C) receives EHG-related ancestry. This disappears in the Kura Araxes period, and re-appears after the Middle Bronze Age (Armenia_MLBIA) in which it is accompanied by a smidgeon of BHG-related ancestry. That the Armenia_MLBIA population had input from the Yamna is certain because of Y-chromosome evidence[13]. The ADMIXTURE results nicely parallel this, by showing that the early steppe influence (at Areni-1) was CLV-like, while the later, post-2500BCE one included also the BHG-related ancestry present in the Yamna.

Finally, we highlight the limitations of this approach with the concrete example of the Central Anatolian Bronze Age population; we modeled this as a mixture of Çayönü Pre-Pottery Neolithic and steppe populations like BPgroup. However, in Fig. S 9 it is modeled as Anatolian and CHG components without any steppe ancestry. Notice that the Çayönü population was excluded Fig. S 8 because its predicted position is outside the convex hull of the four admixture components in the West Eurasian PCA. We can see this in Fig. S 10. Çayönü is indeed outside the convex hull of the four ancestral components; however, TUR_C_BA is indeed on the line connecting Çayönü with BPgroup; thus, the PCA is consistent with its modeling as such a mixture.

In conclusion, our development of unsupervised ADMIXTURE analysis of summary individuals in this note provides a powerful new way of assessing ancestry that complements the results of qpAdm. The four sources of the unsupervised ADMIXTURE model lead to similar conclusions as our qpAdm analysis about the formation of populations of the steppe and North Pontic region. However, they should always be interpreted with caution as even populations of much different ancestry (like "Han") will be forced to have their ancestry from the sources of the model (Fig. S 5). This violation can be detected by comparing summary individuals with their representation in a different statistic, such as PCA (Fig. S 8). However, even that procedure may miss cases (such as TUR_C_BA) which are seemingly well-modeled by unsupervised ADMIXTURE but in fact may possess ancestry other than what can be expressed in the unsupervised model (such as Çayönü-related ancestry; Fig. S 10).

Thus, we end this note by cautioning the reader that the unsupervised ADMIXTURE procedure developed here provides a powerful new tool of ancestry analysis that overcomes the problem of spurious population-centered components, but its results should be interpreted in the context of other lines of evidence (such as PCA and qpAdm) to arrive at a consistent reconstruction.

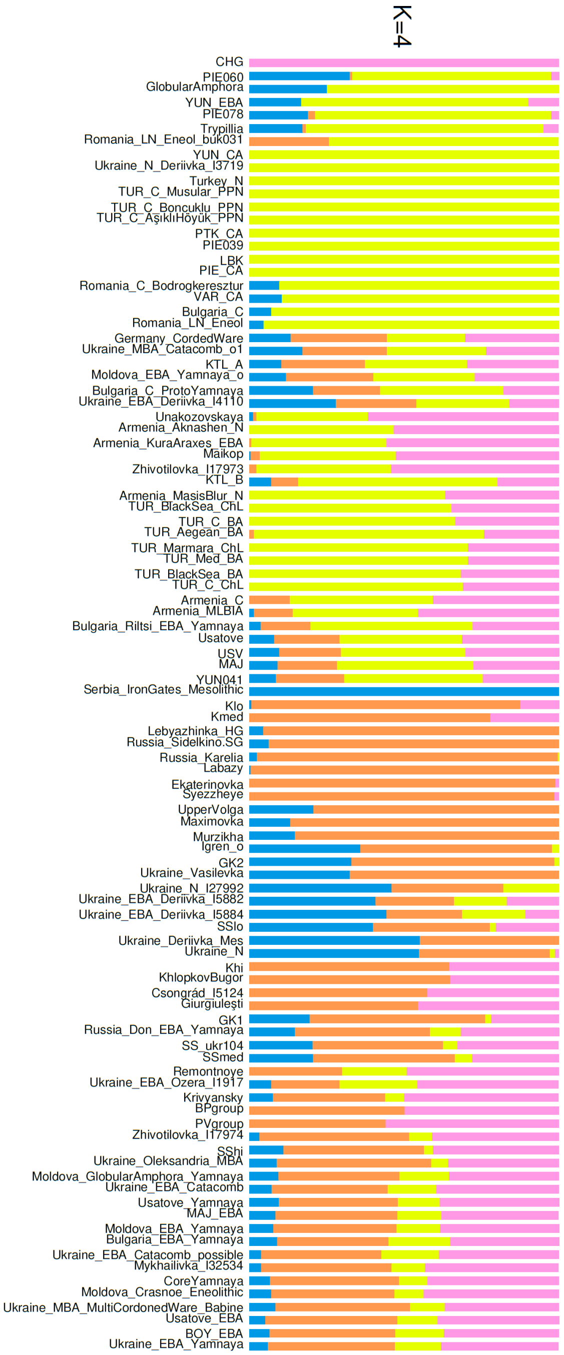

**Fig. S 9 Unsupervised ADMIXTURE of summary individuals from diverse populations (clean version).**

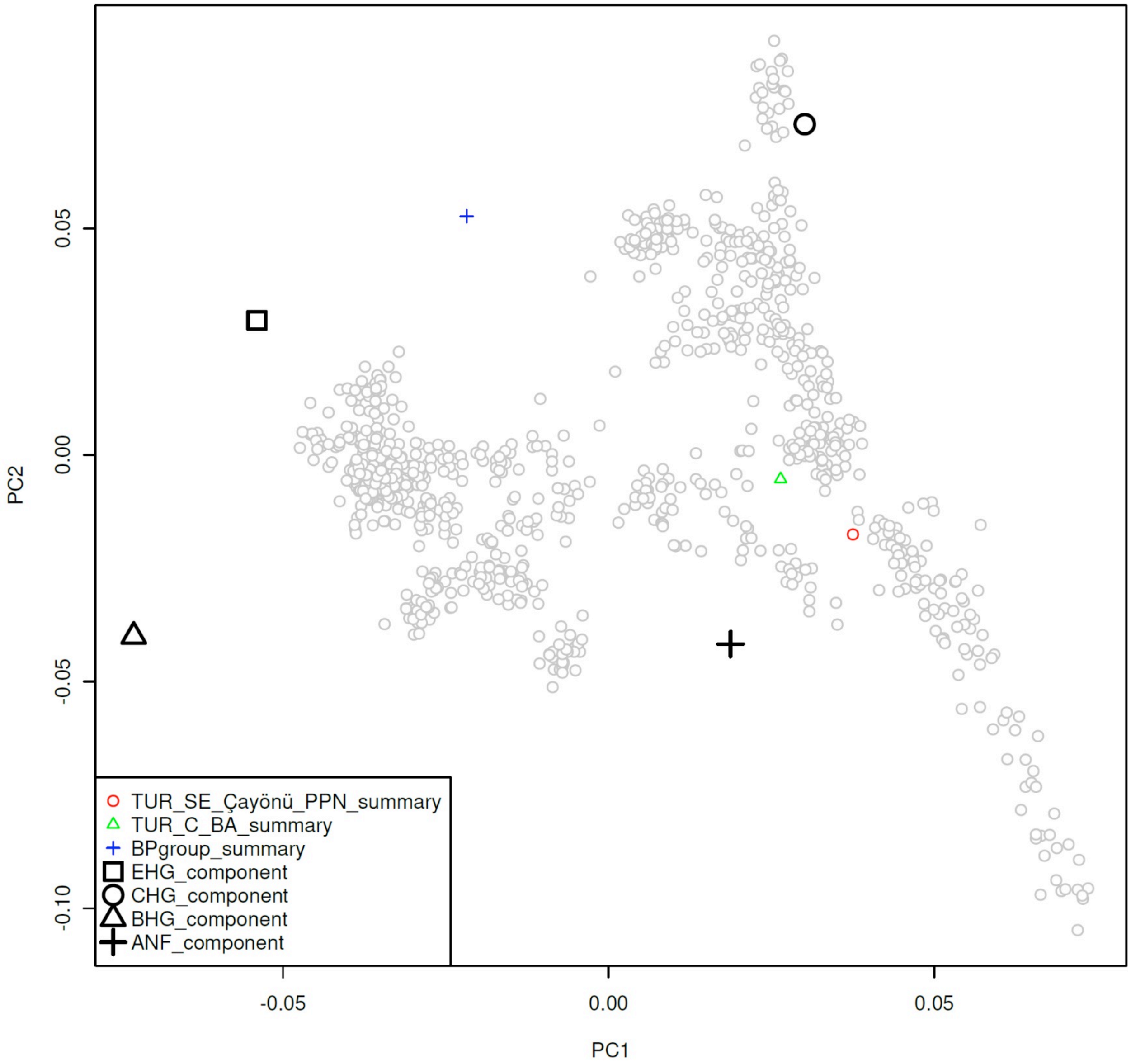

**Fig. S 10 Modeling TUR_C_BA.** TUR_C_BA is on the line between Çayönü and BPgroup; this corresponds to the qpAdm model we have developed for this population. When modeled in terms of the 4-source model it is modeled as a mixture of Anatolian (cross) and CHG (circle) ancestral populations.

**References**

1 Alexander, D. H., Novembre, J. & Lange, K. Fast model-based estimation of ancestry in unrelated individuals. *Genome Research* **19**, 1655-1664, doi:10.1101/gr.094052.109 (2009).

2 Ayub, Q. *et al.* The Kalash Genetic Isolate: Ancient Divergence, Drift, and Selection. *The American Journal of Human Genetics* **96**, 775-783, doi:https://doi.org/10.1016/j.ajhg.2015.03.012 (2015).

3 Lazaridis, I., Patterson, N., Anthony, D. & others. The Genetic Origin of the Indo-Europeans. *in submission* (2024).

4 Lazaridis, I. & Reich, D. Failure to replicate a genetic signal for sex bias in the steppe migration into central Europe. *Proceedings of the National Academy of Sciences* **114**, E3873-E3874, doi:10.1073/pnas.1704308114 (2017).

5 Reich, D. *et al.* Reconstructing Native American population history. *Nature* **488**, 370-374, doi:10.1038/nature11258 (2012).
6 Haak, W. *et al.* Massive migration from the steppe was a source for Indo-European languages in Europe. *Nature* **522**, 207-211, doi:10.1038/nature14317 (2015).
7 Lazaridis, I. *et al.* Genomic insights into the origin of farming in the ancient Near East. *Nature* **536**, 419-424, doi:10.1038/nature19310 (2016).
8 Lazaridis, I. *et al.* Ancient human genomes suggest three ancestral populations for present-day Europeans. *Nature* **513**, 409-413, doi:10.1038/nature13673 (2014).
9 Lazaridis, I. *et al.* Ancient DNA from Mesopotamia suggests distinct Pre-Pottery and Pottery Neolithic migrations into Anatolia. *Science* **377**, 982-987, doi:10.1126/science.abq0762 (2022).
10 Harney, É. *et al.* Ancient DNA from Chalcolithic Israel reveals the role of population mixture in cultural transformation. *Nature Communications* **9**, 3336, doi:10.1038/s41467-018-05649-9 (2018).
11 Posth, C. *et al.* Palaeogenomics of Upper Palaeolithic to Neolithic European hunter-gatherers. *Nature* **615**, 117-126, doi:10.1038/s41586-023-05726-0 (2023).
12 Wang, C.-C. *et al.* Ancient human genome-wide data from a 3000-year interval in the Caucasus corresponds with eco-geographic regions. *Nature Communications* **10**, 590, doi:10.1038/s41467-018-08220-8 (2019).
13 Lazaridis, I. *et al.* The genetic history of the Southern Arc: A bridge between West Asia and Europe. *Science* **377**, eabm4247, doi:10.1126/science.abm4247 (2022).
